# CASZ1 regulates the rate at which outer hair cells mature and is required for hearing

**DOI:** 10.1101/2025.07.04.663233

**Authors:** Yoko Nakano, Elizabeth C. Driver, Susan C. Wiechert, Braulio Peguero, Erich T. Boger, Chantal Allamargot, Rebecca Hipp, Angelika Doetzlhofer, Robert J. Morell, Matthew W. Kelley, Botond Bánfi

## Abstract

The transcriptional activator ATOH1 is a master regulator of the development of mechanosensory hair-cells (HCs) in the ear. We report that the ATOH1 target gene *Casz1* encodes a transcription factor that regulates the rate of outer HC (OHC) maturation by gene repression. Genetic deletion of *Casz1* during (but not after) development of the mouse cochlea caused: hearing loss; abnormal organization of mechanosensory stereocilia bundles in OHCs; abnormally low F-actin density in OHC cuticular plates; progressive loss of OHCs; and mild morphological alterations in inner HCs. RNA sequencing revealed that *Casz1* deletion delayed downregulation of genes expressed in immature OHCs, including the actin regulator-encoding gene *Coro2a*, and accelerated upregulation of genes expressed in mature OHCs. *Coro2a* knockdown restored the density of cuticular plate F-actin in *Casz1* mutant OHCs. Our data indicate that CASZ1 regulates transcriptional and morphological maturation of OHCs, and that CASZ1 in maturing HCs is necessary for hearing.

## INTRODUCTION

Hair cells (HCs) are mechanosensory receptor cells in the hearing and balance organs. In the hearing organ (organ of Corti), HCs are located in four rows. These rows form ∼2.6 parallel turns in the cochlea, a snail shell-like part of the inner ear. The innermost HC row consists of inner HCs (IHCs). The other HC rows consist of outer HCs (OHCs). Sound-induced vibrations transmitted to the cochlea from the middle ear cause deflections of microvillus-like structures (stereocilia) on the apical side of IHCs and OHCs. These deflections regulate the activity of mechanoelectrical transduction (MET) channel complexes in HCs (1). In IHCs, the opening and closing of MET channels regulate the release of the neurotransmitter glutamate at synapses formed with auditory neurons (2–4). In OHCs, the opening and closing of MET channels drive changes in cell length, and these oscillations in length generate forces that amplify sound-induced vibrations and cause ∼50 dB gain in hearing sensitivity (5–8).

Approximately 100 stereocilia are located in a single bundle on the apical side of each OHC and IHC (9). In each of these bundles, stereocilia of 3 different heights form staircase-like rows. The protein core of the stereocilium is composed of F-actin and actin-binding proteins. This core is anchored to the cuticular plate, an F-actin-rich dense matrix that underlies the apical plasma membrane in HCs (10,11). Defects in the organization of stereocilia bundles and abnormal thinning of the cuticular plate have been associated with hearing loss in several mutant mouse lines (12–17).

In humans, hearing loss is the most common sensory deficit (18). It can be caused by genetic alterations or environmental insults that irreversibly damage cochlear HCs (19). In humans and other mammals, the damaged HCs are replaced by protrusions of adjacent non-HCs, resulting in the formation of permanent scars in the hearing organ (20,21). In fish and birds, however, HC death stimulates adjacent non-HCs to re-enter the cell cycle and trans-differentiate into HCs, resulting in scarless repair of the sensory epithelium and restored hearing after ototoxic insults and noise trauma (22–26). This observation sparked the development of various approaches to trans-differentiate non-HCs to HCs in the mammalian cochlea (27–30).

Fundamental for the design of HC-regenerating approaches is the identification of regulators of HC development. In several studies, ATOH1 was identified as the master transcriptional regulator of HC development based on the complete absence of HCs in *Atoh1* gene-deleted mice and the ability of ectopically expressed ATOH1 to trans-differentiate non-HCs to HCs in the not fully developed cochlea (31–37). ATOH1 activates transcription directly by binding to E-box sequence motifs in its target genes, but it also regulates gene expression indirectly by upregulating target genes that encode transcription factors such as POU4F3 and GFI1 (38–42). In the context of a fully developed cochlea, ectopic ATOH1 expression is ineffective in trans-differentiating non-HCs to HCs, and trans-differentiation efficiency is greatly improved by the combinatorial expression of ATOH1, POU4F3, and GFI1 (43–46). However, the HC-like cells produced by this approach in the developed cochlea lacked organized stereocilia bundles, indicating that not all significant barriers to HC regeneration were overcome (46). Thus, the improvement of HC-regenerating approaches will require the identification of additional regulators of HC development.

We analyzed previously generated ATOH1 CUT&RUN data, single-cell RNA sequencing (scRNA-seq) data, and functional annotations of genes to identify additional ATOH1 target genes that encode HC-expressed transcription factors (41,47,48). In this study, we selected three of these genes (*Casz1*, *Neurod6*, and *Runx1t1*) for genetic deletion in mice, and found that of the three *Casz1* was the most important for hearing. Genetic deletion of *Casz1* in the cochlea was associated with hearing loss and with abnormally high expression of genes that are transcribed predominantly in OHCs in the organ of Corti. These gene expression defects were largely transient during the first postnatal week, and they were caused by delayed downregulation of immature OHC-expressed genes and accelerated upregulation of mature OHC-expressed genes. Cochlear deletion of *Casz1* was also associated with abnormal regulation of F-actin density in OHC cuticular plates, defective organization of OHC stereocilia bundles, progressive loss of OHCs, and mild alterations in stereocilium thickness in IHCs. Our data indicate that CASZ1-dependent gene repression regulates the rate at which OHC maturation progresses, and that CASZ1 inactivation causes morphological defects in OHCs and loss of hearing.

## RESULTS

### Conditional deletion of *Casz1* causes hearing loss

We searched for novel regulators of HC maturation among transcription factors that are encoded by ATOH1 target genes and expressed in mouse cochlear HCs, using three datasets. These were data from CUT&RUN analysis of ATOH1 binding sites in cochlear HCs, Gene Ontology (GO) annotations, and data from scRNA-seq analysis of the organ of Corti (Figure 1A) (41,47,48). The CUT&RUN data indicated that ∼3000 genes are ATOH1 targets in cochlear HCs (Table S1) (41). Of these, 276 are annotated with ‘transcription regulator activity’ in the GO database (Table S1) (48). Within this subset, only 20 were considered genes of interest because their expression levels are >8-fold higher in maturing HCs versus non-HCs (Figure 1B) on postnatal day 1 (P1), based on the scRNA-seq data (Table S1) (47). An 8-fold difference in expression was used as our cut-off for genes of interest because the upregulation of ATOH1 target genes in HCs is expected to be much more dramatic than that in non-HCs (39), and P1 was the selected time because *Atoh1* expression is the highest in cochlear HCs at ∼P1 (47,49). Of the 20 genes of interest, 3 were reported to be crucial for HC maturation before the completion of this study (i.e., A*toh1*, *Pou4f3*, and *Gfi1*) and were thus excluded from further analysis (50). An additional 7 genes of interest were excluded because previous testing of their effects on hearing had not revealed a role in the inner ear (Figure 1B) (Table S1). Of the 10 remaining genes of interest, we selected the 3 that are most highly expressed in P1 HCs (Figure 1B): neurogenic differentiation 6 (*Neurod6*), castor zinc finger 1 (*Casz1*), and RUNX1 translocation partner 1 (*Runx1t1*).

**Figure 1.**
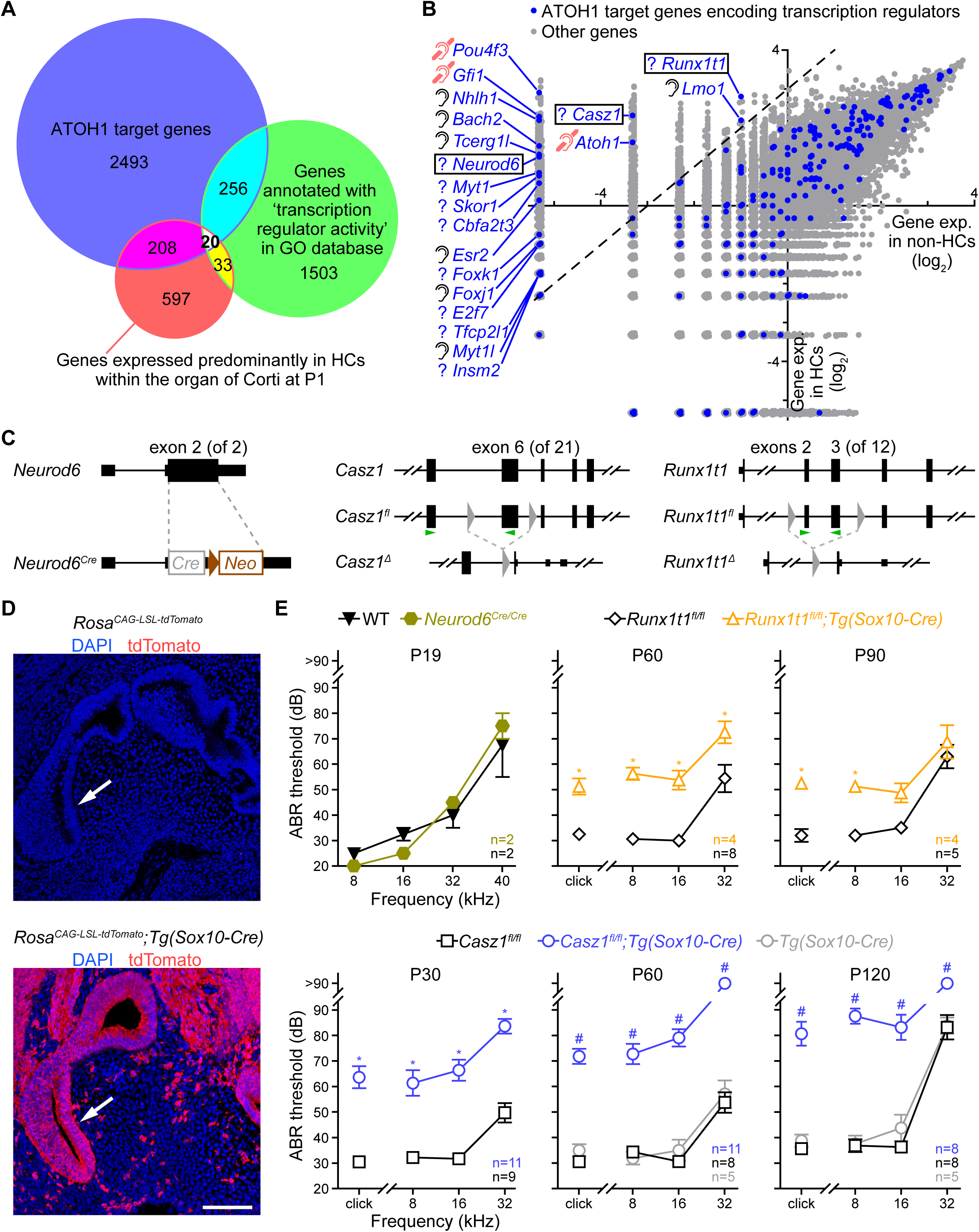
Conditional deletion of *Casz1* causes hearing loss in mice. (A) Venn diagram of shared and unique features of the indicated 3 groups of genes. The definition of predominantly hair-cell (HC)-expressed genes (red circle) is based on >8-fold higher expression in HCs versus non-HCs in the organs of Corti of postnatal day (P) 1 mice. Numbers of genes in the unique and overlapping segments of the diagram are indicated. Gene names are listed in Table S1. GO, Gene Ontology. (B) Gene expression levels in cochlear HCs versus non-HCs in P1 mice, based on scRNA-seq data from a previous study (Kolla et al., 2020). Dashed line marks the boundary of 8-fold difference in expression. Blue dots identify ATOH1 target genes that are annotated with the GO term ‘transcription regulator activity’ (those above the dashed line are additionally identified by gene symbols) and gray dots identify other genes. At left of the gene symbols, a deafness symbol (pink) indicates that the gene is crucial for hearing; an ear symbol (black) indicates that the gene is not required for hearing; and a question mark indicates that the effect of gene on hearing was not described before this study was completed. Black frames identify the genes that were selected for deletion. (C) Schematic of wild-type, exon-floxed (*^fl^*), exon-deleted (*^Δ^*), and Cre knock-in (*^Cre^*) alleles of the indicated genes. Depicted are protein-coding regions of exons (tall rectangles), non-coding regions of exons (short rectangles), introns (horizontal lines), loxP sites (gray triangles), a Cre-encoding sequence (gray rectangle), and a neomycin resistance gene (brown arrow and rectangle). Dashed gray lines indicate regions affected by the exon-deleting recombination events. Exon numbers are shown for the targeted exons. Parenthesized numbers indicate the total number of exons in the indicated genes. Green arrowheads show positions of qRT-PCR primers designed for testing exon deletions in Figure S1B and S1C. (D) DAPI staining (blue) and tdTomato immunostaining (red) of inner ear sections from E13 mice of the indicated genotypes. Arrows point to the cochlea. Scale bar, 100 µm. (E) Auditory brainstem response (ABR) thresholds of mice of the indicated genotypes, tested using broadband (click) and pure tone (8, 16, and 32 kHz) sounds on the indicated postnatal days. Values are mean ± SEM. The number of mice tested in each group is indicated (n). Two-group comparisons: Kolmogorov-Smirnov test with Holm-Sidak correction *p < 0.05. Three-group comparisons: two-way ANOVA (p < 0.0001 for genotype factor) and Dunnett’s post hoc test #p < 0.05 (reference group, *Casz1^fl/fl^*). WT, wild-type.

To test whether *Neurod6*, *Casz1*, and *Runx1t1* are necessary for hearing, we used mice that are homozygous for inactivated versions of the three genes. In the case of *Neurod6*, the gene was inactivated in the germline by replacing the protein coding region of the gene with a Cre-encoding sequence (*Neurod6^Cre/Cre^*) (Figure 1C) (51). In the cases of *Casz1* and *Runx1t1*, inactivation in the germ line is lethal (52,53). For this reason, we used conditional *Casz1* and *Runx1t1* alleles. The *Casz1* conditional allele harbored loxP sites flanking exon 6 (*Casz1^fl^*) (52), and the *Runx1t1* conditional allele harbored loxP sites flanking exons 2 and 3 (*Runx1t1^fl^*) (Figure 1C). To delete these exons, we used the *Tg(Sox10-Cre)* transgene because the *Sox10* promoter is active in the entire otic vesicle before HC marker proteins are upregulated (54–56). Using the Cre reporter allele *Rosa^CAG-LSL-tdTomato^*, we confirmed that *Tg(Sox10-Cre)* causes recombination of loxP sites in the entire cochlear epithelium by embryonic day (E) 13 (Figure 1D). By analyzing tdTomato expression in P0 *Rosa^+/CAG-LSL-tdTomato^*;*Tg(Sox10-Cre)* mice, we reconfirmed that the Cre reporter was activated in HCs and neighboring epithelial cells (Figure S1A). Using qRT-PCR with PCR primers that anneal to the floxed exons, we verified that the expression of these exons is much lower (∼3%) in organ of Corti samples from *Casz1^fl/fl^*;*Tg(Sox10-Cre)* and *Runx1t1^fl/fl^*;*Tg(Sox10-Cre)* mice versus control mice (Figure S1B, S1C).

We next tested the hearing of *Neurod6^Cre/Cre^, Casz1^fl/fl^*;*Tg(Sox10-Cre)*, and *Runx1t1^fl/fl^*;*Tg(Sox10-Cre)* mice by measuring auditory brainstem responses (ABRs) to sound stimuli. The *Neurod6^Cre/Cre^* mice were on the CD-1 genetic background, which is associated with early-onset hearing loss (57), and thus we tested ABRs at P19. The two conditional knock-out (cKO) mouse lines were on the C57BL/6 genetic background, which is not associated with hearing loss before P90–P120 (58,59), and thus we tested ABRs at various times between P30 and P120 (Figure 1E). These tests revealed that ABR thresholds were significantly higher in the two cKO mouse lines versus control (*Casz1^fl/fl^*) mice (Figure 1E). In *Neurod6^Cre/Cre^* and wild-type (WT) mice, the ABR thresholds were similar (Figure 1E). Of the two cKO genotypes, *Casz1^fl/fl^*;*Tg(Sox10-Cre)* was associated with higher ABR thresholds (Figure 1E, Figure S1D). Further analysis of ABR data demonstrated that ABR thresholds in *Casz1^fl/fl^*;*Tg(Sox10-Cre)* mice were higher at P60 versus P30 and still higher at P120 (Figure 1E, Figure S1E). ABR thresholds in *Runx1t1^fl/fl^*;*Tg(Sox10-Cre)* mice did not differ at P60 versus P90 (Figure 1E, Figure S1F). Given that the hearing loss was more severe in *Casz1^fl/fl^*;*Tg(Sox10-Cre)* mice versus *Runx1t1^fl/fl^*;*Tg(Sox10-Cre)* mice, our subsequent experiments focused on *Casz1*.

### Expression of *Casz1* in immature HCs is required for hearing

We used single-molecule fluorescence in situ hybridization (smFISH) to determine the *Casz1* expression pattern in the cochlea at multiple times between E14 and P5. To identify immature HCs at the earliest phases of differentiation, we used *Ccer2* probes (47). To identify immature HCs at later phases of differentiation and spiral ganglion neurons (SGNs), we used *Pvalb* probes (60). This analysis revealed that *Casz1* expression was specific to HCs in the organ of Corti (Figure 1A). In these cells, *Casz1* was upregulated between E14 and P1 and downregulated between P1 and P5 (Figure 2A). In cochlear regions outside the organ of Corti, *Casz1* mRNA was detected in SGNs (Figure 2B, Figure S2A). Thus, within the cochlea, *Casz1* is expressed in immature HCs and SGNs.

**Figure 2.**
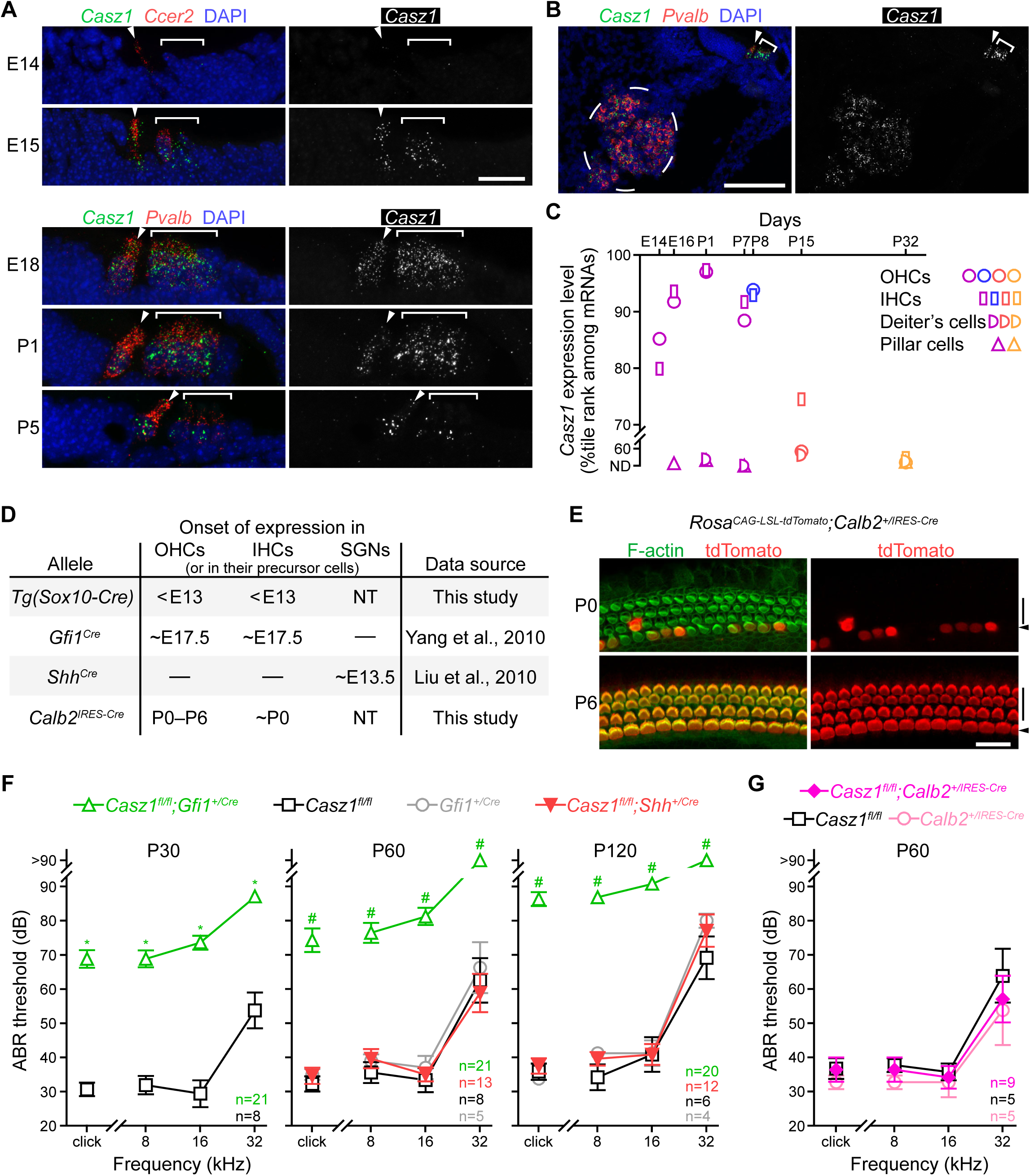
*Casz1* deletion specifically in immature HCs causes hearing loss. (A) Single-molecule fluorescence in situ hybridization (smFISH) of organ of Corti sections from WT mice with probes complementary to the *Casz1* mRNA (green and white as indicated), the HC marker mRNA *Ccer2* (red), and the HC and neuronal mRNA *Pvalb* (red). DAPI (blue) was used for counterstaining. Also indicated are the embryonic (E) and postnatal (P) days when mice were used for smFISH. Arrowheads, IHCs. Brackets, OHCs. Scale bar, 20 μm. (B) smFISH of a cochlear section from a WT mouse (P1) with probes complementary to *Casz1* mRNA (green and white as indicated) and *Pvalb* mRNA (red). DAPI (blue) was used for counterstaining. Dashed ellipse, spiral ganglion. Arrowhead, IHC. Bracket, OHCs. Scale bar, 100 μm. A larger area of the same section is shown in Figure S2A. (C) Percentile rank of *Casz1* expression relative to the expression levels of other mRNAs in OHCs, IHCs, Deiters’ cells, and pillar cells at the indicated times. Symbols represent data from RNA-seq studies by Kolla et al., 2020 (purple), Chessum et al., 2018 (blue), Ranum et al., 2019 (red), and Liu et al., 2018 (orange). ND, not detected. (D) Approximate times of gene expression onset for the indicated transgenic and knock-in alleles in OHCs, IHCs, and spiral ganglion neurons (SGNs). NT, not tested. ––, not expressed. (E) F-actin staining (green) and tdTomato immunostaining (red) of organ of Corti samples from P0 and P6 mice of the indicated genotype. Arrowheads, IHC rows. Vertical black lines, OHC rows. Scale bar, 10 µm. Lower magnification images are shown in Figure S2B. (F and G) ABR thresholds of mice of the indicated genotypes, tested using broadband (click) and pure tone (8, 16, and 32 kHz) sounds on the indicated postnatal days. Values are mean ± SEM. The number of tested mice in each group is indicated (n). Two-group comparison: Kolmogorov-Smirnov test with Holm-Sidak correction *p < 0.05. Three-group and four-group comparisons: two-way ANOVA (p < 0.0001 for genotype factor) and Dunnett’s post hoc test #p < 0.05 (reference group, *Casz1^fl/fl^*).

To evaluate *Casz1* expression in HCs over a longer period of time (E14–P32), we analyzed single-cell and sorted-cell RNA-seq data from four previous studies of mouse organ of Corti (Figure 2C) (47,61–63). In each RNA-seq dataset, we ranked genes based on expression levels and compared percentile ranks from different embryonic and postnatal times. This analysis revealed that the percentile rank of *Casz1* increased in HCs between E14 and P1, and that it decreased between P1 and P32 (Figure 2C). Thus, *Casz1* expression is much higher in immature HCs versus mature HCs.

Given that *Casz1* is expressed in SGNs, immature HCs, and (at low level) in mature HCs, we tested whether inactivation of the *Casz1^fl^* allele separately in these cell groups impaired hearing. To inactivate *Casz1^fl^* specifically in SGNs, we used the *Shh^Cre^* knock-in allele because *Shh^Cre^* drives loxP recombination specifically in immature SGNs within the cochlea (Figure 2D) (64). To inactivate *Casz1^fl^* in immature HCs, we used the *Gfi1^Cre^* knock-in allele because *Gfi1^Cre^* drives loxP recombination specifically in immature (∼E17.5) HCs and some immune cells in the inner ear (Figure 2D) (65,66). To inactivate *Casz1^fl^* in more mature HCs, we used the *Calb2^IRES-Cre^* knock-in allele because our analysis of *Rosa^+/CAG-LSL-tdTomato^;Calb2^+/IRES-Cre^* mice revealed that *Calb2^IRES-Cre^* causes loxP recombination in HCs during the first postnatal week (Figure 2E, Figure S2B). qRT-PCR tests confirmed that *Casz1* expression was abnormally low in the spiral ganglions of *Casz1^fl/fl^*;*Shh^+/Cre^* mice and in the organs of Corti of *Casz1^fl/fl^*;*Gfi1^+/Cre^* and *Casz1^fl/fl^*;*Calb2^+/IRES-Cre^* mice (Figure S2C, S2D). ABR testing of these mouse lines revealed that ABR thresholds were higher in *Casz1^fl/fl^*;*Gfi1^+/Cre^* mice versus control (*Casz1^fl/fl^*) mice (Figure 2F, Figure S2E). In *Casz1^fl/fl^*;*Shh^+/Cre^*, *Casz1^fl/fl^*;*Calb2^+/IRES-Cre^*, and *Casz1^fl/fl^* mice, the ABR thresholds were similar (Figure 2F, 2G). Thus, *Casz1* deletion in immature HCs causes hearing loss. In contrast, *Casz1* deletion in SGNs and in more mature (first postnatal week) HCs does not cause hearing loss.

### *Casz1* deletion in immature HCs causes malformation of stereocilia bundles and progressive OHC loss

We tested the effect of *Casz1* deletion on HC morphology in *Casz1^fl/fl^*;*Tg(Sox10-Cre)* and *Casz1^fl/fl^*;*Gfi1^+/Cre^* mice. Organ of Corti samples from these mice and control mice (*Casz1^fl/fl^*) were collected at various times (P6, P14, P30, and P60) and stained with fluorescently labeled phalloidin to visualize stereocilia bundles and other F-actin-rich structures. Microscopy analysis of these samples revealed that many stereocilia bundles were missing in the OHC rows in the cKO mouse lines at P30 and P60 (Figure 3A). In these areas, the pattern of F-actin staining was identical to the previously observed pattern of F-actin distribution at sites of OHC loss and epithelial scar formation (Figure 3B) (20,21,67). Quantification of OHC stereocilia bundles revealed that the loss of OHCs in the basal half of the cochlea started between P14 and P30, and that it continued between P30 and P60 (Figure 3C). In contrast, the number of IHCs was unchanged in the cKO mice at P60 (Figure S3A). These data indicate that CASZ1 inactivation in immature HCs causes progressive OHC loss.

**Figure 3.**
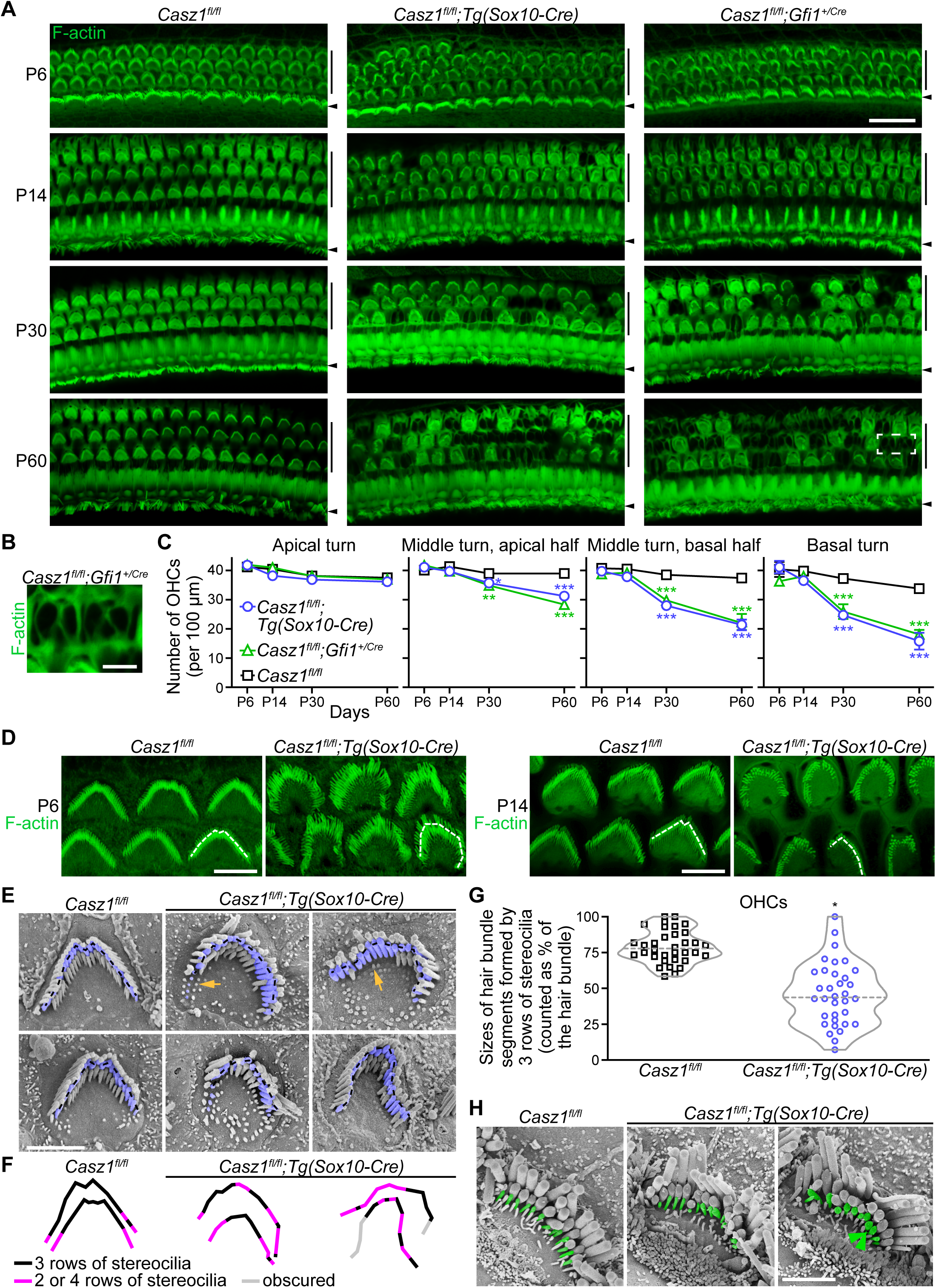
*Casz1* deletion in HCs causes malformation of OHC stereocilia bundles and loss of OHCs. (A) F-actin staining of organ of Corti samples from middle turn of the cochlea of mice of the indicated genotypes. Also indicated are the postnatal days (P) when samples were collected. Arrowheads, IHC rows. Vertical lines, OHC rows. Scale bar, 20 μm. Dashed rectangle indicate an area magnified in (B). (B) Scarred area in the organ of Corti magnified from (A). Scale bar, 5 μm. (C) Numbers of OHCs in the indicated turns of cochlea in *Casz1^fl/fl^;Tg(Sox10-Cre)*, *Casz1^fl/fl^;Gfi1^+/Cre^*, and control (*Casz1^fl/fl^*) mice at the indicated times. Values are mean ± SEM (n = 3 mice per time and genotype). Two-way ANOVA (genotype factor p = 0.17 in Apical turn graph and p < 0.0001 in other graphs), Dunnett’s post hoc test *p = 0.04, **p = 0.01, ***p < 0.001 (reference group, *Casz1^fl/fl^*). (D) F-actin staining of OHCs in organ of Corti samples from P6 and P14 mice of the indicated genotypes. Larger areas of the samples are shown in Figure S3B. Dashed white lines are drawn over the tips of second-row stereocilia of one OHC per image. Scale bar, 5 µm. (E) Scanning electron microscopy images of OHCs of mice of the indicated genotypes (P14). Second-row stereocilia are pseudo-colored blue. Dashed black lines are drawn over the tips of second-row stereocilia. Orange arrows indicate two stubby stereocilia. Scale bar, 2 µm. (F) Stacked display of lines drawn over the tips of second-row stereocilia in (E). Colors indicate the number of stereocilia rows along the bundle (3 rows, black; 2 or 4 rows, purple; third-row stereocilia obscured by other structures, gray). (G) Quantification of segments of stereocilia bundles formed by 3 rows of stereocilia in OHCs of mice of the indicated genotypes (P14). Dashed gray lines indicate medians (Welch’s test *p < 0.0001). (H) Scanning electron microscopy images of IHCs of P14 mice of the indicated genotypes. Third-row stereocilia are pseudocolored green. Two abnormally thick stereocilia are indicated (green arrowheads). Scale bar, 2 µm.

Further analysis of F-actin-stained OHCs revealed that prior to the OHC loss, many stereocilia bundles were misshapen in *Casz1^fl/fl^*;*Tg(Sox10-Cre)* and *Casz1^fl/fl^*;*Gfi1^+/Cre^* mice (Figure 3A). To visualize these defects at higher resolution, we used Airyscan microscopy on P6 and P14 samples from *Casz1^fl/fl^*;*Tg(Sox10-Cre)* mice. This analysis confirmed that most OHC stereocilia bundles were misshapen at both P6 and P14, with bundles lacking the W or V-like symmetrical shape that was present in the same tissue in *Casz1^fl/fl^* mice (Figure 3D, Figure S3B, S3C). Analysis of IHCs by Airyscan microscopy revealed that in P14 *Casz1^fl/fl^*;*Tg(Sox10-Cre)* mice, ∼35% of IHCs contained third-row stereocilia that were abnormally thick (Figure S3D, S3E).

We next used scanning electron microscopy to visualize stereocilia bundles at still higher resolution. This confirmed that in P14 *Casz1^fl/fl^*;*Tg(Sox10-Cre)* mice the shapes of most OHC stereocilia bundles were not W or V like (Figure 3E, 3F). Many of the misshapen stereocilia bundles included stubby stereocilia that were shorter than the distance between stereocilia rows (Figures 3E, Figure S3F). The scanning electron microscopy analysis also revealed that much larger segments of OHC stereocilia bundles consisted of not 3 but 1, 2, or 4 rows of stereocilia in *Casz1^fl/fl^*;*Tg(Sox10-Cre)* mice versus *Casz1^fl/fl^* mice at P14 (Figure 3E, 3F, 3G). Thus, CASZ1 is necessary for proper organization of OHC stereocilia bundles. Scanning electron microscopy analysis of IHCs confirmed that in P14 *Casz1^fl/fl^*;*Tg(Sox10-Cre)* mice some third-row stereocilia were abnormally thick (Figure 3H). These data indicate that *Casz1* deletion causes morphological defects in the stereocilia bundles of OHCs and IHCs.

### CASZ1 represses many OHC-specifically expressed genes

We analyzed gene expression in organs of Corti from P4 *Casz1^fl/fl^*;*Tg(Sox10-Cre)* and WT mice using RNA-seq; we thereby identified 1118 differentially expressed genes (Figure 4A; Table S2). Most of these genes (64%) were expressed at abnormally high levels in the *Casz1^fl/fl^*;*Tg(Sox10-Cre)* samples, and several OHC marker mRNAs (e.g., *Ocm*, *Cacng2*, and *Serpina1c*) were represented. To more systematically explore the effects of CASZ1 inactivation on HC-specific genes, we generated lists of predominantly OHC-expressed, predominantly IHC-expressed, and predominantly OHC *and* IHC (OHC–IHC) expressed genes based on previous scRNA-seq data (Table S3) (47). The predominantly OHC–IHC-expressed gene set did not overlap with the other two groups, and it included genes that are expressed at much higher levels in both IHCs and OHCs than non-HCs (Methods). Of the predominantly OHC-expressed genes, 49 out of 190 were differentially expressed in *Casz1^fl/fl^*;*Tg(Sox10-Cre)* versus WT organ of Corti. Among these genes, most (76%) were expressed at an abnormally high level in the *Casz1^fl/fl^*;*Tg(Sox10-Cre)* samples (Figure 4B). Of the predominantly IHC-expressed genes, 28 out of 121 were differentially expressed. Among these genes, 50% were expressed at an abnormally high level in the *Casz1^fl/fl^*;*Tg(Sox10-Cre)* samples (Figure 4B). Of the predominantly OHC–IHC-expressed genes, 135 out of 853 were differentially expressed. Among these genes, 74% were expressed at abnormally high level in the *Casz1^fl/fl^*;*Tg(Sox10-Cre)* samples (Figure S4A, Table S3). Thus, the CASZ1-dependent regulation of predominantly OHC-expressed and OHC–IHC-expressed genes is largely repressive. In contrast, the CASZ1-dependent regulation of predominantly IHC-expressed genes is an ∼equal mix of activation and repression.

**Figure 4.**
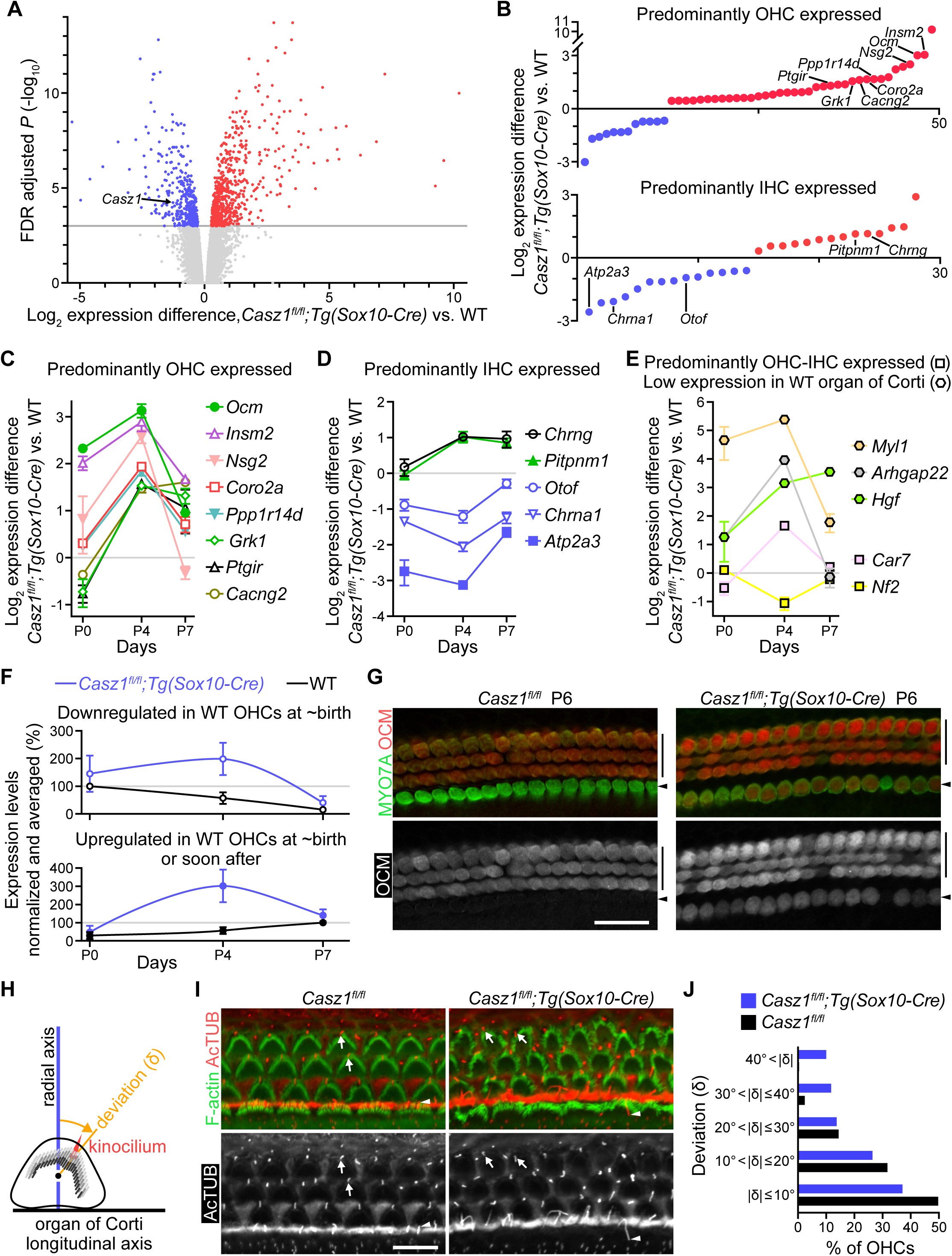
Conditional deletion of *Casz1* causes abnormally high expression of predominantly OHC-expressed genes and misorientation of stereocilia bundles in OHCs. (A) Volcano plot of gene expression differences in organ of Corti samples from *Casz1^fl/fl^;Tg(Sox10-Cre)* versus WT mice (P4), as revealed by RNA-seq. Colored dots represent genes that are expressed at abnormally low (blue) or high (red) levels. Gray dots represent genes that are not expressed at significantly different levels. Arrow indicates the difference in *Casz1* expression. Gray horizontal line indicates the cut-off for statistical significance. (B) Differential expression of predominantly OHC-expressed and predominantly IHC-expressed genes in organ of Corti samples from *Casz1^fl/fl^;Tg(Sox10-Cre)* versus WT mice (P4), as revealed by RNA-seq. Color coding is the same as in (A). Gene symbols indicate genes that are selected for further testing by qRT-PCR in (C and D). (C, D, and E) qRT-PCR data for differential expression of indicated genes in organs of Corti from *Casz1^fl/fl^;Tg(Sox10-Cre)* versus WT mice at P0, P4, and P7. Data are plotted separately for genes that are expressed predominantly in OHCs (C), predominantly in IHCs (D), predominantly in OHCs and IHCs (squares in E), or expressed at very low level in the WT organ of Corti (hexagons in E). Values are mean ± SEM (n = 3 mice / genotype). Gray horizontal lines indicate zero difference. (F) Normalized and averaged expression levels of two subgroups of predominantly OHC-expressed genes in organs of Corti of mice of the indicated genotypes at P0, P4, and P7, calculated using qRT-PCR data from (C). One subgroup (*Cacng2*, *Coro2a*, *Grk1*, *Insm2*, and *Ptgir*) is defined based on downregulation at around birth in WT OHCs (open circles). The other subgroup (*Nsg2*, *Ocm*, and *Ppp1r14d*) is defined based on upregulation at around birth in WT OHCs (closed circles). Maximum expression in the WT organ of Corti is the 100% expression level (gray lines). Values are mean ± SEM. (G) Immunofluorescence detection of OCM (red and white as indicated) and MYO7A (green) in organ of Corti samples from control (*Casz1^fl/fl^*) and *Casz1^fl/fl^*;*Tg(Sox10-Cre)* mice (P6). Arrowheads, IHC rows. Vertical lines, OHC rows. Scale bar, 20 µm. (H) Schematic representation of deviation angle (δ) formed between the radial axis of the organ of Corti (blue) and the line (orange) connecting the kinocilium (red triangle) base with the center of the cuticular plate (black dot) in an OHC. Also depicted are the apical surface (curved triangle) and stereocilia bundle (short gray lines) of the OHC. Horizontal black line represents the longitudinal axis of the organ of Corti. (I) F-actin staining (green) and acetylated α-tubulin (AcTUB) immunostaining (red and white as indicated) of organ of Corti samples from *Casz1^fl/fl^* and *Casz1^fl/fl^*;*Tg(Sox10-Cre)* mice (P6). Two OHC kinocilia (arrows) and one IHC kinocilium (arrowhead) are indicated in each panel. Scale bar, 10 µm. (J) OHC kinocilium deviation angles (δ) in organs of Corti of P6 mice of the indicated genotypes (n = 150 OHCs per genotype, Kolmogorov-Smirnov test p = 0.0009).

We selected 12 differentially expressed genes for qRT-PCR-based testing in organ of Corti samples from P4 *Casz1^fl/fl^*;*Tg(Sox10-Cre)*, WT, and *Casz1^fl/fl^*;*Gfi1^+/Cre^* mice. This analysis validated the alterations in gene expression in the *Casz1^fl/fl^*;*Tg(Sox10-Cre)* samples, and it revealed that 9 of the 12 genes were also differentially expressed in *Casz1^fl/fl^*;*Gfi1^+/Cre^* versus WT mice (Figure S4B). These data indicate that the alterations in cochlear gene expression are similar in the two *Casz1* cKO mouse lines.

We next tested whether the magnitude of alterations in gene expression varied during the first postnatal week. RNA was isolated from organs of Corti of *Casz1^fl/fl^*;*Tg(Sox10-Cre)* and WT mice at P0, P4, and P7, and the expression levels of 18 genes were tested using qRT-PCR. These tests revealed that the gene expression alterations were larger at P4 versus P0 and P7 (Figure 4C, 4D, 4E). Thus, during the first postnatal week the magnitude of alterations in gene expression changed rapidly in the *Casz1^fl/fl^*;*Tg(Sox10-Cre)* organ of Corti.

We processed further the qRT-PCR data from Figure 4C to evaluate the gene expression changes over time within genotype groups and, simultaneously, the gene expression differences between genotype groups (Figure 4F). The additional data processing involved sorting the genes into two subgroups based on previously reported timing of maximum expression in WT OHCs (i.e., ≤P1 vs. P7≤), averaging the magnitudes of gene expression alterations within the two subgroups, and calculating the over-time expression changes relative to the HC-specific control mRNA *Myo6* because previous RNA-seq data indicated that *Myo6* expression is steady during the first postnatal week in HCs (47,49). This analysis demonstrated that the *Casz1^fl/fl^*;*Tg(Sox10-Cre)* genotype was associated with delayed downregulation of immature OHC-expressed genes (i.e., *Cacng2*, *Coro2a*, *Grk1*, *Insm2*, and *Ptgir*) and accelerated upregulation of mature OHC-expressed genes (i.e., *Nsg2*, *Ocm*, and *Ppp1r14d*) (Figure 4F). Thus, CASZ1 regulates the timing and rates of some gene expression changes in the organ of Corti during the first postnatal week.

To test specifically the OHCs from *Casz1^fl/fl^;Tg(Sox10-Cre)* mice for some of the gene expression alterations that were identified in the organ of Corti, we collected OHCs from organs of Corti of P4 *Casz1^fl/fl^;Tg(Sox10-Cre)* mice and control (WT) mice using a combination of enzymatic digestion and selective pipetting of fluorescent dye (FM1-43) labeled HCs (Methods). The isolated cells were analyzed by qRT-PCR to determine expression levels of: the HC-specific control mRNA *Myo6*, the IHC-specific mRNA *Fgf8* (47), and 5 mRNAs whose expression is regulated by CASZ1 and that are expressed predominantly in OHCs (*Coro2a*, *Nsg2*, and *Ppp1r14d*) or both OHCs and IHCs (*Car7* and *Galnt9*). *Fgf8*-expressing samples were excluded from further analysis and gene expression was normalized to that of *Myo6*. The normalized data revealed that the expression of the 5 tested genes was 4–8-fold higher in *Casz1^fl/fl^;Tg(Sox10-Cre)* samples versus WT samples (Figure S4C). Thus, in immature (P4) OHCs CASZ1 acts as a repressor of some predominantly OHC-expressed genes and some predominantly OHC–IHC-expressed genes.

One of the genes that is expressed predominantly in OHCs and is upregulated to abnormally high levels in P4 *Casz1* cKO mice is *Ocm* (Figure 4C, Figure S4B). We used a previously validated anti-OCM antibody (68) and an immunofluorescence approach to visualize OCM in HCs of *Casz1^fl/fl^*;*Tg(Sox10-Cre)* mice at P6 and P30, and found that OCM expression was abnormally high in OHCs at P6 but not at P30 (Figure 4G, Figure S4D, S4E). In the IHCs, OCM expression was abnormally high at both P6 and P30 (Figure S4E). However, at both time points, the level of OCM expression was much lower in IHCs versus OHCs (Figure S4E). These data indicate that CASZ1 is a repressor of *Ocm* in both IHCs and OHCs, and that this repressor effect is brief in OHCs and longer lasting in IHCs.

We next analyzed GO annotations of genes that are differentially expressed in *Casz1^fl/fl^*;*Tg(Sox10-Cre)* versus WT organ of Corti samples and identified ‘glycoprotein’ and ‘cilium’ with the lowest p values (Figure S4F). In HCs, the only microtubule-based cilium is the kinocilium; the stereocilia are F-actin-based projections and thus structurally distinct from true cilia (69). In immature HCs, however, it is the location of the kinocilium that determines the orientation of the stereocilia bundle (69). In immature OHCs, the kinocilium is located close to the radial axis that runs through the center of the cuticular plate (Figure 4H) (69). Towards the end of HC maturation, the kinocilium is resorbed (70). To analyze the orientations of stereocilia bundles in OHCs, we visualized kinocilia in the cochlea of P6 *Casz1^fl/fl^*;*Tg(Sox10-Cre)* and *Casz1^fl/fl^* mice using an antibody against acetylated α-tubulin. We found that the position of kinocilium was abnormal in some OHCs in *Casz1^fl/fl^*;*Tg(Sox10-Cre)* mice (Figure 4I). We quantified and confirmed this defect by calculating the deviation angle that is formed between the radial axis and the line drawn from the base of the kinocilium to the center of the cuticular plate (Figures 4H, 4J). The visualization of acetylated α-tubulin also revealed that kinocilia were abnormally long and that they were resorbed abnormally late in OHCs and IHCs of *Casz1^fl/fl^*;*Tg(Sox10-Cre)* mice (Figure S4G, S4H, S4I, S4J, S4K). Thus, altered expression of kinocilium-related genes in *Casz1^fl/fl^*;*Tg(Sox10-Cre)* mice is associated with defects in positioning, regulation of length, and resorption of kinocilia.

### Elevated *Coro2a* expression is associated with reduced density of cuticular plate F-actin in OHCs

Deficiencies for kinociliary proteins have been reported to cause alterations in the orientation of stereocilia bundles, but not the loss of third-row stereocilia or the loss of OHCs in P0–P60 mice (71–77). We therefore searched for other groups of genes that could contribute to the OHC defects observed in *Casz1^fl/fl^*;*Tg(Sox10-Cre)* mice. Of the 1118 CASZ1-regulated genes that we identified, 35 are annotated with the GO term ‘perception of sound’ (Figure S5A, Table S2). However, previously reported manipulation of these genes either did not cause morphological defects in OHCs or caused morphological alterations that differ from those produced by the inactivation of *Casz1* (Table S4). Next, we searched for CASZ1-regulated genes that are annotated with ‘actin binding’ or ‘myosin complex’ GO terms because many actin-binding proteins and myosins are necessary for the formation of stereocilia bundles (10). This search identified 9 genes that are expressed at >2-fold higher or <0.5-fold lower levels in the organs of Corti of *Casz1^fl/fl^*;*Tg(Sox10-Cre)* versus WT mice at P4 (Figure 5A). Of these 9 genes of interest, 4 were excluded from further analysis based on data from previous gene deletion and overexpression studies (i.e., *Hpca*, *Nf2*, *Fscn2*, and *Nod2*) (Table S5). Another (*Cap1*) was excluded because *Cap1* expression is abnormally high in only one of the two *Casz1* cKO mouse lines that are characterized by hearing loss (Figure S4B). In addition, *Myh13* was deprioritized because its large size prevents gene delivery to HCs in a single adeno-associated virus (AAV). Thus, only the following three genes were prioritized: coronin 2a (*Coro2a*), carbonic anhydrase 7 (*Car7*), and myosin light chain 1 (*Myl1*).

**Figure 5.**
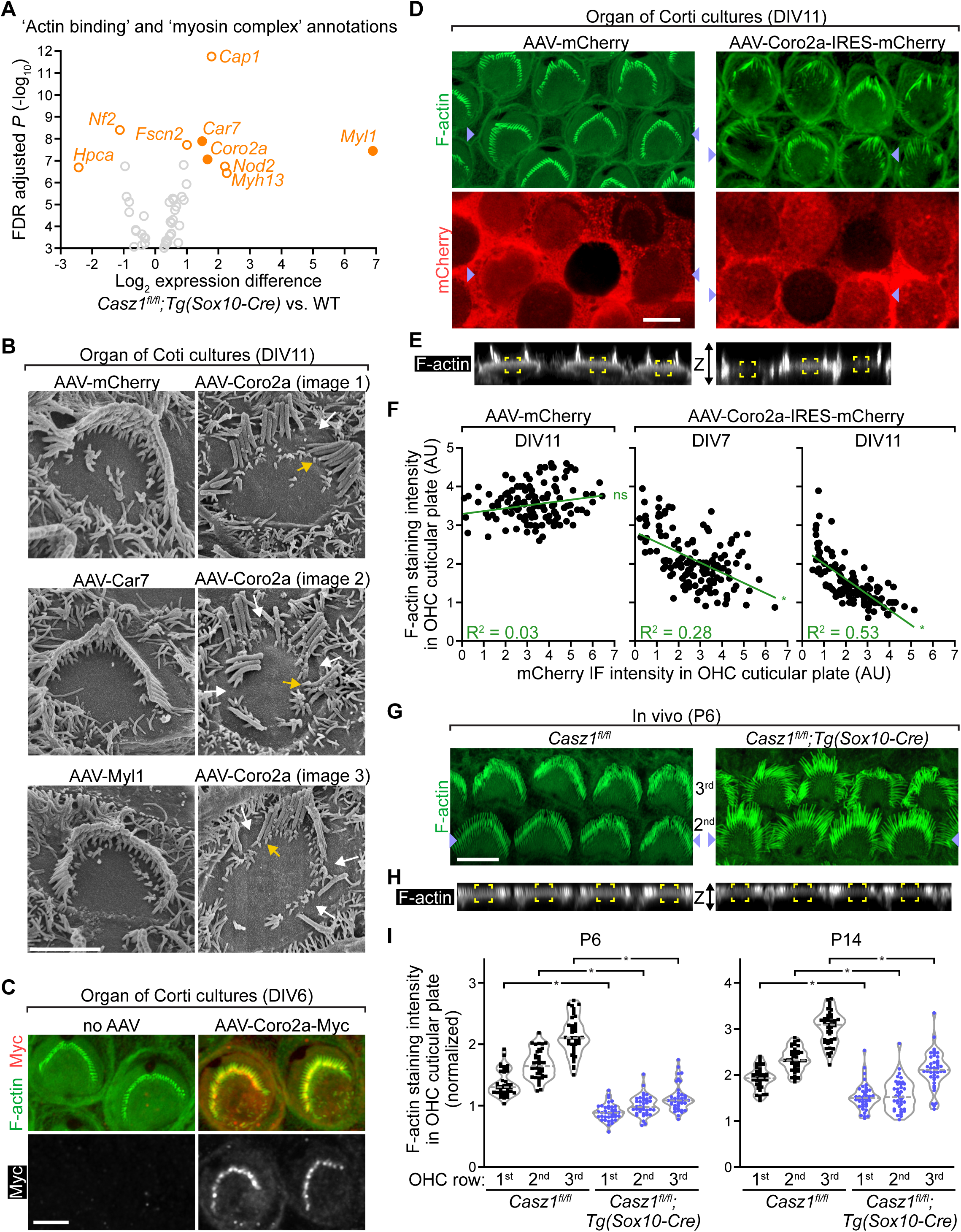
Overexpression of *Coro2a* causes degeneration of stereocilia bundles and cuticular plates in OHCs. (A) Volcano plot of a subset of gene expression differences between organs of Corti of *Casz1^fl/fl^*;*Tg(Sox10-Cre)* and WT mice, selected based on annotations of genes with GO terms ’actin binding’ and ’myosin complex’. Orange circles and gene symbols indicate larger than 2-fold expression differences; gray circles indicate less than 2-fold expression differences. Filled circles represent genes that were selected for overexpression in (B). (B) Scanning electron microscopy images of OHC stereocilia bundles in day in vitro (DIV) 11 organ of Corti cultures that were derived from WT mice (E17.5) and incubated with the indicated AAVs on DIV0. Arrows indicate large gaps between first-row stereocilia (white) and some abnormally short stereocilia (orange). Scale bar, 3 µm. (C) F-actin staining (green) and Myc-tag immunostaining (red and white as indicated) of DIV6 organ of Corti cultures that were derived from WT mice (E17.5) and incubated with the indicated AAV or no AAV on DIV0. Scale bar, 3 µm. (D) F-actin staining (green) and mCherry immunostaining (red) of OHCs in DIV11 organ of Corti cultures that were derived from WT mice (E17.5) and incubated with the indicated AAVs on DIV0. Blue arrowheads indicate places selected for visualization of F-actin in the XZ-plane in (E). Larger areas of the same samples are shown in Figure S5D. Scale bar, 5 μm. (E) Visualization of F-actin staining (white) in the XZ-plane between points that are indicated by blue arrowheads in (D). Yellow corners indicate 2 × 2 µm areas at the centers of cuticular plates of OHCs. (F) F-actin staining intensities (y-axes) and mCherry immunofluorescence (IF) intensities (x-axes) in the centers of OHC cuticular plates in DIV7 and DIV11 organ of Corti cultures that were incubated with the indicated AAVs as in (D). Each dot represents fluorescence intensities in a 2 × 2 µm area (in the XZ-plane) of one cuticular plate. Green lines represent best-fit linear regression. R^2^, goodness of fit (two-tailed t tests for deviation from zero *p < 0.0001; ns, not significant). AU, arbitrary units. (G) F-actin staining (green) of OHCs of P6 mice of the indicated genotypes. Blue arrowheads indicate places selected for visualization in the XZ-plane in (H). Numbers between the images indicate positions of OHC rows relative to pillar cells. Larger areas of the same samples are shown in Figure S5H. Scale bar, 5 μm. (H) Visualization of F-actin staining (white) in the XZ-plane between points that are indicated by blue arrowheads in (G). Yellow corners indicate 2 × 2 µm areas at the centers of cuticular plates. (I) Normalized intensities of F-actin staining in the middle areas of OHC cuticular plates in mice of the indicate genotypes at P6 and P14. Each symbol represents fluorescence intensity in a 2 × 2 µm area (in the XZ-plane) of one cuticular plate divided by the average fluorescence intensity in 2 × 2 µm apical regions (in the XZ-plane) of inner sulcus cells (P6) or pillar cells (P14) in the same image stack. Numbers below the plots indicate the OHC rows where the analyzed OHCs were located. Mann-Whitney tests, false discovery rate (FDR) adjusted *p < 0.0001.

Analysis of previously generated scRNA-seq data revealed that in WT organ of Corti the patterns of expression of the 3 prioritized genes differ (Figure S5B) (47). *Coro2a* is expressed in immature OHCs, but only briefly. *Car7* is upregulated in both OHCs and IHCs during the first postnatal week and is expressed in mature OHCs and IHCs. *Myl1* is expressed at very low levels in both mature and immature HCs. In our *Casz1^fl/fl^*;*Tg(Sox10-Cre)* mice, the expression levels of these genes in the organ of Corti was abnormally high (Figure 4C, 4E). We therefore overexpressed them in WT organ of Corti cultures using AAVs. AAV-Coro2a, AAV-Car7, AAV-Myl1, and AAV-mCherry (control) were produced in HEK293T cells and incubated with day in-vitro (DIV) 0 organ of Corti cultures that were derived from E17.5 mice. On DIV11, the OHC stereocilia bundles were analyzed for morphology. Scanning electron microscopy revealed that in AAV-Coro2a-incubated organ of Corti cultures many stereocilia bundles were abnormal (Figure 5B). Large gaps were between first-row stereocilia, and most second-row and third-row stereocilia were abnormally short (Figure 5B). Organ of Corti cultures incubated with the other AAVs lacked these phenotypes (Figure 5B). qRT-PCR analysis of a separate set of AAV-incubated organ of Corti cultures confirmed that the AAV-delivered genes were overexpressed (Figure S5C). Thus, overexpression of *Coro2a* is associated with degeneration of OHC stereocilia bundles.

Next, we tested the subcellular localization of overexpressed CORO2A. This test was facilitated by inserting a Myc tag sequence into the AAV-Coro2a genome (AAV-Coro2a-Myc). Organ of Corti cultures from WT mice (E17.5) were incubated with AAV-Coro2a-Myc on DIV0 and stained for Myc and F-actin on DIV6. Microscopy analysis of these samples revealed that CORO2A-Myc was located in stereocilia and cuticular plates of OHCs (Figure 5C). Thus, overexpressed CORO2A is located in F-actin rich structures in OHCs.

CORO2A has been reported to regulate actin turnover in various cancer cell lines, as well as in immune cells (78–81). Therefore, we examined the effect of CORO2A overexpression on stereocilia bundles in OHCs in organ of Corti cultures. To identify cells that overexpress AAV-delivered and non-tagged CORO2A, we subcloned an IRES sequence and an mCherry-encoding sequence into the AAV-Coro2a vector (AAV-Coro2a-IRES-mCherry). This AAV and control AAV (AAV-mCherry) were incubated with DIV0 organ of Corti cultures derived from E17.5 WT mice, and the organ cultures were stained for F-actin and mCherry at DIV7 and DIV11. Microscopy analysis of the AAV-Coro2a-IRES-mCherry-transduced organ of Corti cultures revealed that large gaps were in the stereocilia bundles of most mCherry-expressing OHCs at DIV11, but not at DIV7 (Figure 5D, S5D, S5E). In the DIV11 and DIV7 control cultures (transduced with AAV-mCherry), very few stereocilia bundles were fragmented by large gaps (Figure 5D, Figure S5D, S5E). Thus, overexpression of CORO2A caused degeneration of stereocilia bundles in OHCs by DIV11.

The same organ of Corti cultures were also analyzed for effects of AAV-Coro2a-IRES-mCherry on F-actin density in cuticular plates of OHCs. F-actin density was assessed based on F-actin staining intensity in the middle portion of cuticular plates along the Z axis (Figure 5E), and CORO2A expression was assessed based on mCherry immunostaining intensity in the same cells; there was an inverse correlation between the two staining intensities at both DIV7 and DIV11 (Figure 5E, 5F, Figure S5F, S5G). Such a correlation was not detected in AAV-mCherry-incubated cultures (Figure 5E, 5F). These data indicate that CORO2A overexpression causes loss of F-actin from the cuticular plate in OHCs, and that this loss precedes the degeneration of stereocilia bundles.

Given that the expression of *Coro2a* is abnormally high in OHCs of *Casz1^fl/fl^*;*Tg(Sox10-Cre)* mice (Figure S4C), we assessed the density of F-actin in OHC cuticular plates of these and control (*Casz1^fl/fl^*) mice at P6 and P14. In the P6 samples, the intensity of F-actin staining in OHC cuticular plates was normalized to F-actin staining intensity in the apical (microvilli-rich) region of inner sulcus cells, and in P14 samples it was normalized to F-actin staining intensity in the apical region of pillar cells. This analysis revealed that the intensity of F-actin staining was abnormally low in cuticular plates in *Casz1^fl/fl^*;*Tg(Sox10-Cre)* OHCs at both P6 and P14 (Figure 5G, 5H, 5I, Figure S5H). A similar analysis of IHC cuticular plates in P6 *Casz1^fl/fl^*;*Tg(Sox10-Cre)* and control mice did not reveal a difference in F-actin staining intensity (Figure S5I). These data indicate that F-actin density is abnormally low in the cuticular plates of OHCs (but not IHCs) in *Casz1^fl/fl^*;*Tg(Sox10-Cre)* mice.

### Knockdown of *Coro2a* in *Casz1* KO OHCs restores F-actin density in the cuticular plate

We used short hairpin RNAs (shRNAs) to reduce expression of *Coro2a* in organ of Corti cultures from *Casz1^fl/fl^;Tg(Sox10-Cre)* mice. Four shRNAs were designed to target *Coro2a* (shRNA*_Coro2a_*), and two control shRNAs were designed to target the HC marker mRNA *Calb1* (shRNA*_Calb1_*) (Figure 6A, Figure S6A, S6B). *Calb1* was selected as a control target because the encoded protein is not required for HC maturation (82). The efficacy of shRNAs was tested in HEK293T cells using Firefly luciferase (Fluc)-encoding reporter constructs, which were generated by subcloning the target regions (30-mer each) of shRNAs*_Coro2a_* and shRNAs*_Calb1_*between the start ATG and a T2A peptide-encoding sequence directly upstream of the Fluc-encoding sequence (Figure 6B, 6C) (Table S6). HEK293T cells were co-transfected with a reporter construct, an shRNA expression cassette, and a renilla luciferase (Rluc)-encoding control plasmid, and 36 h later Fluc and Rluc activities were measured. These tests revealed that each of the designed shRNAs inhibited expression of the target sequence-containing reporters (Figure 6B, 6C).

**Figure 6.**
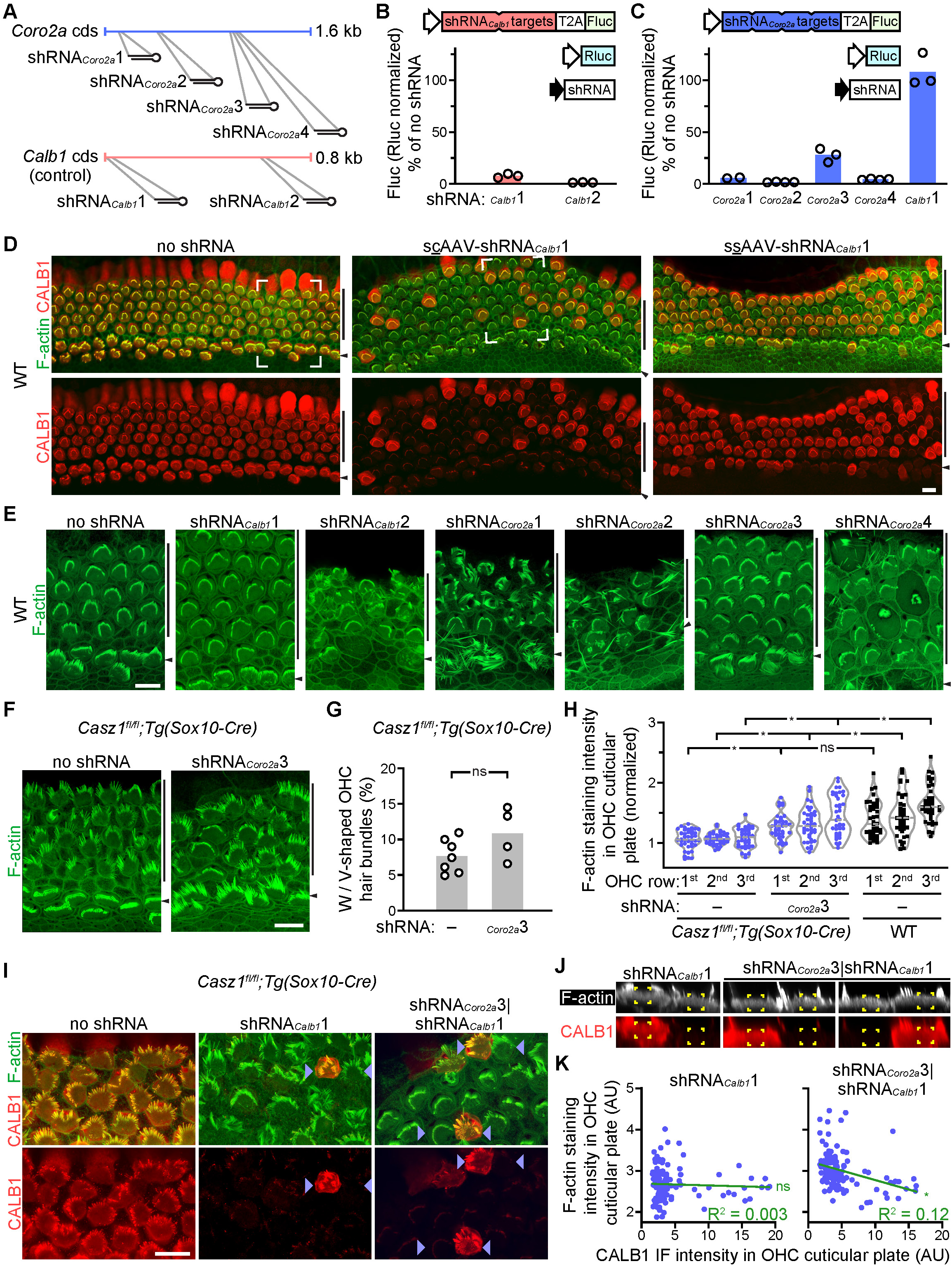
Knockdown of *Coro2a* restores F-actin density in the cuticular plates of *Casz1* KO OHCs. (A) Schematic of *Coro2a*-targeting shRNAs (shRNA*_Coro2a_*1–4), *Calb1*-targeting shRNAs (shRNA*_Calb1_*1 and shRNA*_Calb1_*2), and positions of shRNA complementary regions (gray lines) in the coding sequences (cds) of *Coro2a* (blue) and *Calb1* (red). The lengths of cds are shown in kilobases (kb). (B and C) Expression levels of a shRNA*_Calb1_* reporter construct (B) and a shRNA*_Coro2a_* reporter construct (C) in transfected HEK293T cells, evaluated based on enzymatic activity of the encoded Firefly luciferase (Fluc). Schematics at top show compositions of the transfected constructs, including the Fluc-encoding shRNA reporters, shRNA expression cassettes, and a Renilla luciferase-encoding (Rluc) reference construct. Control cell cultures (100% Fluc) were transfected with the luciferase-encoding constructs only (no shRNA). Circles represent data from separate experiments. Open arrow, thymidine kinase gene promoter. Filled arrow, U6 promoter. T2A indicates ribosomal skipping sequence. ‘shRNA targets’ (red and blue polygons) represent series of 30-base sequences that are complementary to the indicated shRNAs. (D) F-actin staining (green) and CALB1 immunostaining (red) of DIV9 organ of Corti cultures that were derived from WT mice (E17.5) and incubated with the indicated AAVs or no AAV (no shRNA) on DIV0. White corners indicate areas magnified in (E). scAAV, self-complementary AAV. ssAAV, single-stranded AAV. Vertical lines, OHC rows. Arrowheads, IHC rows. Scale bar, 10 μm. (E) F-actin staining of DIV9 organ of Corti cultures that were derived from WT mice (E17.5) and incubated with shRNA-delivering scAAVs or no AAV (no shRNA) on DIV0. The scAAV-delivered shRNAs are indicated. Vertical lines, OHC rows. Arrowheads, IHC rows. Scale bar, 10 μm. (F) F-actin staining of DIV9 organ of Corti cultures that were derived from *Casz1^fl/fl^;Tg(Sox10-Cre)* mice (E17.5) and incubated with scAAV-shRNA*_Coro2a_*3 or no AAV (no shRNA) on DIV0. Vertical lines, OHC rows. Arrowheads, IHC rows. Scale bar, 10 μm. (G) Quantification of W / V-shaped OHC stereocilia bundles in scAAV-shRNA*_Coro2a_*3-incubated and not AAV-incubated (–) organ of Corti cultures that were derived from *Casz1^fl/fl^;Tg(Sox10-Cre)* mice. The organ of Corti cultures were transduced and stained as in (F). Stereocilia bundle shapes were categorized based on criteria shown in Figure S6D. Circles represent data for organ of Corti cultures from separate mice. Bars indicate means (unpaired t test; ns, significant). (H) Normalized intensities of F-actin staining in middle areas of OHC cuticular plates in DIV9 organ of Corti cultures that were derived from E17.5 mice of the indicated genotypes and incubated with scAAV-shRNA*_Coro2a_*3 or no AAV (–) on DIV0. Each symbol represents fluorescence intensity in a 2 × 2 µm area (in the XZ-plane) of one cuticular plate divided by the average fluorescence intensity in 2 × 2 µm apical regions (in the XZ-plane) of inner sulcus cells in the same image stack. Fluorescence intensities are plotted separately for OHCs in different rows (n = 40 OHCs per row). Two-way ANOVA (genotype factor p < 0.0001) and Dunnett’s posthoc test *p < 0.05; ns, not significant. (I) F-actin staining (green) and CALB1 immunostaining (red) of DIV9 organ of Corti cultures that were derived from *Casz1^fl/fl^;Tg(Sox10-Cre)* mice (E17.5) and incubated with no AAV (no shRNA) or shRNA-delivering scAAVs on DIV0. The scAAV delivered shRNAs are indicated. Blue arrowheads indicate places selected for visualization in the XZ-plane in (J). Scale bar, 10 μm. (J) Visualization of F-actin staining (white) and CALB1 immunostaining (red) in the XZ-plane between points that are indicated by blue arrowheads in (I). Yellow corners indicate 2 × 2 µm areas at the centers of cuticular plates. (K) F-actin staining intensities (y-axes) and CALB1 immunofluorescence (IF) intensities (x-axes) in the centers of OHC cuticular plates in DIV9 organ of Corti cultures that were derived from *Casz1^fl/fl^;Tg(Sox10-Cre)* mice (E17.5) and incubated with shRNA-delivering scAAVs on DIV0. The scAAV delivered shRNAs are indicated. Each dot represents fluorescence intensities in a 2 × 2 µm area (in the XZ-plane) of one cuticular plate. Green lines represent best-fit linear regression. R^2^, goodness of fit (two-tailed t test for deviation from zero *p < 0.0001; ns, not significant). AU, arbitrary units.

Given the small size of the shRNA expression cassettes (∼450 bp), both single-stranded (ss) AAVs and self-complementary (sc) AAVs can be used as vectors for shRNA delivery. However, ssAAVs and scAAVs differ in their DNA packaging capacity (∼4.7 kb versus ∼2.3 kb) as well as their genome structure (83). In the case of scAAV, the genome is similar to that of double-stranded DNA, and second-strand DNA synthesis is not required for the expression of the delivered genes (83). Thus, the scAAV system can produce higher gene expression, with the magnitude of the difference depending on the cell type (83). We used the shRNA*_Calb1_*1 expression cassette to test the efficacies of ssAAV and scAAV in delivering an shRNA to organ of Corti cultures. Specifically, organ of Corti cultures from E17.5 WT mice were incubated with scAAV-shRNA*_Calb1_*1 and ssAAV-shRNA*_Calb1_*1 on DIV0, and CALB1 expression was assessed on DIV9. Microscopy analysis of these samples revealed that CALB1 expression was much lower in scAAV-shRNA*_Calb1_*1-incubated versus ssAAV-shRNA*_Calb1_*1-incubated organ of Corti cultures (Figure 6D, Figure S6C). Therefore, scAAV was selected for shRNA delivery.

To test the effects of each of the designed shRNAs on the morphology of WT HCs, organ of Corti cultures from E17.5 WT mice were incubated with the scAAV-shRNAs on DIV0 and used for F-actin staining on DIV9. Morphology of the stereocilia bundles was assessed based on the widths, lengths, and heights of the bundles and the sizes of gaps between stereocilia (Figure S6D and S6E). Two-thirds of the tested shRNAs caused degeneration of stereocilia bundles in both OHCs and IHCs, and the only non-damaging constructs were shRNA*_Calb1_*1 and shRNA*_Coro2a_*3 (Figure 6E, Figure S6F). Given that CALB1 is not required for the formation of stereocilia bundles and IHCs express very little *Coro2a* (47,82), we suggest that the HC-damaging shRNAs were toxic independent of the expression of the target mRNAs. This notion is consistent with previous studies reporting sequence-dependent but target-independent neurotoxicity and hepatotoxicity of other shRNAs (84,85).

We tested the effects of shRNA*_Coro2a_*3 on the morphology of stereocilia bundles and density of cuticular plate F-actin in the OHCs of *Casz1* cKO mice. Organ of Corti cultures from E17.5 *Casz1^fl/fl^;Tg(Sox10-Cre)* mice were incubated with scAAV-shRNA*_Coro2a_*3 on DIV0 and used for F-actin staining and qRT-PCR on DIV9. The F-actin staining revealed that scAAV-shRNA*_Coro2a_*3 largely restored the density of F-actin in the cuticular plates of OHCs but did not restore the W / V-like shape of OHC stereocilia bundles (Figure 6F, 6G, 6H), and the qRT-PCR analysis demonstrated that *Coro2a* expression was lower in scAAV-shRNA*_Coro2a_*3-incubated versus non-transduced organ of Corti cultures (Figure S6G). These data indicate that the knockdown of *Coro2a* resulted in upregulation of F-actin in cuticular plates of *Casz1* KO OHCs.

To evaluate association between shRNA*_Coro2a_*3 expression and F-actin density in cuticular plates, we visualized the shRNA*_Coro2a_*3-expressing OHCs by co-delivering shRNA*_Calb1_*1 with shRNA*_Coro2a_*3 (AAV-shRNA*_Coro2a_*3|shRNA*_Calb1_*1) and identifying CALB1 knocked-down cells. This approach was selected because the commonly used mCherry and EGFP expression cassettes are too large to be included alongside an shRNA expression cassette in the scAAV genome. DIV0 organ of Corti cultures from E17.5 *Casz1^fl/fl^;Tg(Sox10-Cre)* mice were incubated with scAAV-shRNA*_Coro2a_*3|shRNA*_Calb1_*1 or the control virus scAAV-shRNA*_Calb1_*1. On DIV9, these organ of Corti cultures were assessed for CALB1 and F-actin expression (Figure 6I). In organ of Corti cultures transduced with scAAV-shRNA*_Coro2a_*3|shRNA*_Calb1_*1, but not those transduced with scAAV-shRNA*_Calb1_*1, the intensity of CALB1 immunofluorescence correlated inversely with the intensity of F-actin staining in OHC cuticular plates (Figure 6J, 6K). These data indicate that the co-delivery of *Calb1-* and *Coro2a*-targeting shRNAs caused an increase in F-actin density in the cuticular plates of *Casz1* KO OHCs. Therefore, we suggest that CASZ1 regulates F-actin density in OHC cuticular plates by repressing *Coro2a*.

## DISCUSSION

Our study reveals that CASZ1 regulates the progression of transcriptional maturation of OHCs and is required for the long-term survival of OHCs. In the developing organ of Corti, CASZ1 fine-tunes the timing of gene expression changes by briefly repressing sets of genes during the first postnatal week. Some of these genes are expressed during an early phase of OHC maturation and downregulated during the first postnatal week, others are upregulated during the first postnatal week and expressed in mature OHCs. Genetic inactivation of CASZ1 de-represses these genes, thereby delaying downregulation of immature OHC-expressed genes and accelerating upregulation of mature OHC-expressed genes. At the morphological level, CASZ1 regulates the organization of stereocilia bundles in OHCs and the thickness of third-row stereocilia in IHCs. Based on these data, we suggest that CASZ1 fine-tunes the transcriptional maturation of OHCs by gene repression, and that CASZ1 activity in immature OHCs is important for the organization of stereocilia bundles and the survival of OHCs.

Our analysis of various *Casz1* cKO mouse lines revealed that CASZ1 in immature HCs is necessary for hearing but CASZ1 in mature HCs is not (Figure 2F, 2G). Therefore, we suggest that in immature HCs the inactivation of CASZ1 causes irreversible cellular damage. This notion is supported by the progressive nature of OHC loss and the failure of F-actin density to recover in OHC cuticular plates in *Casz1^fl/fl^;Tg(Sox10-Cre)* mice (Figure 3C, 5I). Our analysis of *Casz1* cKO mice also revealed that CASZ1 acts mainly as a repressor of genes that are expressed in OHCs or both OHCs and IHCs (Figure 4B, Figure S4A). The gene repressor complex NuRD have been identified as a CASZ1-interacting histone modifier in various cell lines and retinal progenitor cells (86–88), and the NuRD complex subunits CHD4 and KDM1A have been reported to be expressed in cochlear HCs of P0 mice (89). The mRNAs of other NuRD complex subunits have also been detected in the developing organ of Corti (89); however, the effect of this complex on cochlear gene expression has not been reported. We suggest that HC-specific inactivation of the NuRD complex will be necessary to evaluate its role in the repression of CASZ1 regulated genes in the organ of Corti.

During the writing of this manuscript, a complementary study on the cochlear function of CASZ1 was published (90). In that study, Sun and colleagues associate the *Casz1^fl/fl^*;*Atoh1^Cre/+^* genotype with trans-differentiation of IHCs to OHCs, loss of OHCs, and deafness. Our study also identified OHC loss and hearing loss (though milder) in the *Casz1* cKO mouse lines *Casz1^fl/fl^*;Tg(Sox10-Cre) and *Casz1^fl/fl^;Gfi1^+/Cre^*. The two studies differ in the evaluation of gene regulatory functions of CASZ1. Our gene expression data reveal that in immature OHCs CASZ1 represses several genes that are predominantly expressed in OHCs or both OHCs and IHCs, that this repression is incomplete and transient, and that it regulates the rate of change in gene expression in maturing OHCs. The study by Sun at al. (2025) did not reveal that CASZ1 regulates predominantly OHC-expressed genes and predominantly OHC–IHC-expressed genes in OHCs probably because P10 OHCs were analyzed for alterations in gene expression (90). In P10 OHCs, altered expression of metabolism-related genes was identified and interpreted to be indicative of OHC degeneration (90). These data support our model that in OHCs, CASZ1 represses briefly the predominantly OHC expressed and predominantly OHC–IHC-expressed genes around P4 but not 6 days later. Sun et al. (2025) also analyzed nearly-mature IHCs of *Atoh1^Cre/+^;Casz1^fl/fl^* mice (at P14) and concluded that in IHCs CASZ1 represses OHC-specifically expressed genes and upregulates IHC-specifically expressed genes (90). Our detection of OCM in IHCs of *Casz1* cKO mice at P4 and P30 supports the notion that CASZ1 is a repressor of predominantly OHC-expressed genes in IHCs. We note that OCM expression is much lower in *Casz1* KO IHCs versus WT OHCs, and that many *Casz1* KO IHCs do not express OCM at P30 (Figure S4D). Based on these data, we suggest that CASZ1 acts as a repressor of predominantly OHC-expressed genes in both OHCs and IHCs, and that the extent and duration of the repression is different in the two HC types.

Additional differences in the data reported by Sun et al. (2025) and our group relate to the extent of the morphological alterations in IHCs as well as the severity of hearing loss. Sun et al. (2025) showed that the *Casz1^fl/fl^*;*Atoh1^Cre/+^* genotype is associated with fusion of IHC stereocilia at P14, and with near-complete loss of hearing at P42 (90). In contrast, the *Casz1^fl/fl^;Tg(Sox10-Cre)* mice studied by our group did not exhibit either fusion of IHC stereocilia at P14 or a near-complete hearing loss at P60. We suggest that these differences stem from the use of different Cre alleles. In the *Atoh1^Cre^* allele, the Cre sequence replaces the endogenous *Atoh1* sequence. Thus, the *Casz1^fl/fl^*;*Atoh1^Cre/+^* mice are heterozygous for *Atoh1*, and deletion of a single *Atoh1* allele causes hearing impairment in mice ∼2 months after birth (91). Thus, we suggest that *Atoh1* heterozygosity is a modifier of IHC defects and hearing defects in *Casz1^fl/fl^*;*Atoh1^Cre/+^* mice. Accurate testing of this possibility will require the production of a new mouse line (e.g., *Casz1^fl/fl^;Tg(Sox10-Cre);Atoh1^+/-^*).

Our study is also distinct in that it identifies *Coro2a* as a CASZ1 repressed gene that encodes a regulator of the density of cuticular plate F-actin. shRNA-mediated knock-down of *Coro2a* largely restored the density of F-actin in the cuticular plates of OHCs of *Casz1^fl/fl^;Tg(Sox10-Cre)* mice, but it did not rescue the organization of the stereocilia bundle (Figure 6G, 6H). Therefore, we suggest that different alterations of gene expression are responsible for the reduced density of cuticular plate F-actin versus the defective organization of stereocilia bundles in *Casz1* KO OHCs. Although the cuticular plate is important for hearing and for providing structural support to stereocilia rootlets, a previous study demonstrated that the loss of ∼70% of F-actin from the cuticular plate does not alter severely the organization of stereocilia bundles (12). Specifically, inactivation of the cuticular plate protein LMO7 was shown to cause an ∼70% loss in cuticular-plate thickness but only mild asymmetry in the W-shape of stereocilia bundles in OHCs (12). Based on these data, we suggest that in *Casz1* KO OHCs the abnormally low density of cuticular plate F-actin is not the main cause of the defective organization of stereocilia bundles.

Our shRNA studies additionally revealed that most of the shRNAs*_Coro2a_*and half of the shRNAs*_Calb1_* caused degeneration of both IHCs and OHCs (Figure 6E). We suggest that these effects were independent of the target transcripts for two reasons. Firstly, CALB1 is not required for the maturation or survival of HCs (82). Secondly, *Coro2a* is not expressed in IHCs (Figure S5B) (47). Previous testing of various shRNAs in the central nervous system and liver demonstrated that many shRNAs cause cell death regardless of their targets, and several mechanisms have been proposed to account for shRNA toxicity, including competitive inhibition of microRNA processing, degradation of off-target transcripts, and a largely unexplored mechanism involving upregulation of RNAi pathway genes (92). In light of these prior data, we interpret the toxic effects of shRNAs on HCs in our experiments as a major side effect of the use of AAV-shRNAs in the organ of Corti.

Although the three ATOH1 target genes *Casz1*, *Pou4f3*, and *Gfi1* all play roles in the maturation of HCs, the survival of HCs differs greatly in *Casz1^fl/fl^*;Tg(Sox10-Cre) mice versus *Pou4f3^-/-^* and *Gfi1^-/-^* mice. In developing and young adult *Casz1^fl/fl^*;Tg(Sox10-Cre) mice, HC survival is impaired only in OHC rows and only after ∼P14 (Figure 3C). In contrast, the *Pou4f3^-/-^* and *Gfi1^-/-^* genotypes are associated with near-complete loss of OHCs by P4–P5 and degeneration or loss of IHCs by P32 (42,93–95). At the level of gene expression regulation in HC, POU4F3 and GFI1 are necessary for the upregulation of large sets of genes (41,94,96). In contrast, CASZ1 fine-tunes the timing and magnitude of gene expression changes. Although HC maturation is affected less robustly in *Casz1^fl/fl^*;Tg(Sox10-Cre) mice versus *Pou4f3^-/-^* and *Gfi1^-/-^* mice, we suggest that artificial reprogramming of non-HCs to long-term viable OHCs will require expression of CASZ1 in the transdifferentiating cells.

## METHODS

### Mice

All mouse procedures were approved by the University of Iowa Institutional Animal Care and Use Committee. Mice were housed in groups in temperature-controlled rooms (21±2 °C) with a 12 h / 12 h dark / light cycle. Food and water were available for the mice ad libitum. Animal husbandry and health status monitoring were provided by the Office of Animal Resources staff at the University of Iowa. The ages of tested mice are indicated in the figure legends. The sexes of pre-weaned mice were not determined. The sexes of weaned mice (P21 and older) were documented. The *Neurod6^Cre/Cre^* and *Casz1^fl/fl^* mouse lines were described previously (51,52). The *Runx1t1^fl/fl^* mouse line was generated for this study at GemPharmatech Co., Ltd. In these mice, *Runx1t1* exons 2 and 3 are flanked by loxP sites. The insertions of these loxP sites were verified by PCR and Sanger sequencing. To delete the loxP-flanked gene segments, we used the previously described mouse lines B6;CBA-Tg(Sox10-Cre)1*^Wdr^*/J (54), B6.Cg-*Shh^tm1(EGFP/Cre)Cjt^*/J (64), B6(Cg)-*Calb2^tm1(Cre)Zjh^*/J (97), and FVB/N-*Gfi1^+/Cre^* (65).

### Cell Lines and Tissue Culture

HEK293T cells (female) were obtained from Takara Bio and grown in Dulbecco’s Modified Eagle Medium (DMEM), supplemented with 10% fetal bovine serum, 100 U/mL penicillin, and 100 mg/mL streptomycin (all from Thermo Fisher Scientific). Organs of Corti were dissected from E17.5 mice, placed on Matrigel-coated Transwell permeable supports (Corning), and grown in neurobasal-A medium supplemented with B-27, N-2, 0.5 mM L-glutamine, and 100 U/mL penicillin (all from Thermo Fisher Scientific) (98). Half of the culture medium was exchanged for fresh medium every 3 days. All cell cultures were incubated at 37°C in humidified air atmosphere containing 5% CO_2_ until use.

### Hearing Tests

The ABR thresholds of mice were measured using a previously described open-field system, and both broadband and pure-tone stimuli (99).

### smFISH

Inner ears were collected from CD-1 mice and fixed with 4% paraformaldehyde in PBS (4% PFA) overnight at 4°C. Samples were cryoprotected using sequential incubations in 15% and 30% sucrose (in PBS), embedded into Optimal Cutting Temperature compound, frozen, and sectioned to 10 µm slices. The sections were pretreated and processed for smFISH using the ACD RNAscope Fluorescent Multiplex Reagent kit (versions 1 and 2) following the manufacturer’s instructions (47). smFISH probes for *Casz1*, *Ccer2*, and *Pvalb* detection were purchased from ACD (cat# 502461, 479181, 421931). DAPI was used as a counterstain to visualize cell nuclei.

### Immunofluorescence and Histochemistry

The antibodies used for this study are listed in Table S7. Whole-mount preparations of organs of Corti were fixed with 4% PFA for 2 h at room temperature, permeabilized with 0.5% Triton X-100 in PBS for 20 minutes, and incubated with 1% bovine serum albumin in PBS for 1.5 h. These samples were incubated with primary antibodies overnight at 4°C using the following antibody dilutions: anti-RFP antibody 1:350, anti-MYO7A antibody (mouse monoclonal) 1:15, anti-MYO7A antibody (rabbit polyclonal) 1:300, anti-acetylated α-tubulin antibody 1:1000, anti-OCM antibody 1:150, anti-Myc antibody 1:15, and anti-CALB1 antibody 1:100. For OCM immunostaining, an antigen retrieval step (incubation in pH 6.0 citrate buffer [10 mM] for 20 minutes at 95°C) was added between the fix and permeabilization steps. The binding of primary antibodies was visualized using Alexa 488-conjugated or Alexa 594-conjugated secondary antibodies (1:400 dilution). For F-actin staining, samples were fixed with 4% PFA for 2 h at room temperature, permeabilized with 0.5% Triton X-100 in PBS for 20 minutes, incubated with 1% bovine serum albumin in PBS for 1.5 h, and incubated with phalloidin-Alexa 488 (0.14 µM) for 20 minutes. Expression of *Tg(Sox10-Cre)* in the cochlea was examined using E13 *Rosa^+/CAG-LSL-tdTomato^;Tg(Sox10-Cre)* and control (*Rosa^+/CAG-LSL-tdTomato^*) mice. Heads of these mice were fixed with 4% PFA for 8 h at 4 °C, cryoprotected in 15% and 30% sucrose solutions, embedded into Optimal Cutting Temperature compound, frozen, and cryosectioned to 14 µm slices. The sections were incubated with 1% bovine serum albumin in PBS for 1.5 h and with an anti-RFP antibody (1:350 dilution) overnight at 4 °C. The binding of the primary antibody was visualized using Alexa 594-conjugated secondary antibody (1:400 dilution). Images of fluorescently labeled tissues were acquired using an LSM 880 confocal microscope (Figures 2E, 3A, 3B, 4G, 4I, S1A, S2B, and S4D) or an LSM980 confocal microscope in Airyscan mode (Figures 1D, 3D, 5C, 5D, 5E, 5G, 5H, 6D, 6E, 6F, 6I, 6J, S3B, S3D, S5D, S5F, S5G, and S5H) (Carl Zeiss). Images were analyzed using the ZEN lite 2012 software (Carl Zeiss).

### Scanning Electron Microscopy

Cochleae were dissected from mouse inner ears and fixed with 4% PFA overnight at 4°C. These samples were transferred into paraformaldehyde-free PBS, dissected further to expose the apical surface of HCs, and re-fixed with Karnovsky’s fixative overnight at 4°C. The re-fixed samples were stained using the osmium tetroxide-thiocarbohydrazide method, dehydrated, and critical-point dried before being mounted on aluminum stubs for imaging in a Hitachi S-4800 scanning electron microscope (100).

### RNA Extraction from the Organ of Corti

The floor of scala media was dissected from the cochleae of P0, P4 and P7 mice under a stereomicroscope (SMZ645; Nikon). These tissue pieces from P4 and P7 mice were incubated with 5 μM FM1-43 (Thermo Fisher Scientific) in HBSS for ∼15 s to fluorescently label HCs. The scala media floor pieces from P0 mice were not incubated with FM1-43 because P0 HCs are not selectively labeled by FM1-43 (101). All these tissue pieces were incubated with a mixture of thermolysin (0.2 mg/ml; Sigma-Aldrich) and DNase I (100 unit/ml; Worthington) in DMEM/F-12 for 5–6 minutes at 37°C to facilitate further dissection. The enzymatic digestion was quenched with an equal volume of ice cold DMEM/F-12 containing 10% FBS, and the tissue pieces were further dissected to remove cells that are not components of the organ of Corti. In the P4 and P7 samples, this final dissection was aided by the fluorescent visualization of FM1-43 labeled HCs. The dissected organ of Corti pieces were transferred into RLT lysis buffer (RNeasy Micro Kit; Qiagen), and RNA was isolated from the tissue lysates using the RNeasy Micro Kit (Qiagen).

### RNA-seq

Organ of Corti RNA was pooled from 7–9 mice to obtain a single sample with sufficiently high concentration for RNA-seq. In total, 39 WT mice (n = 5 samples) and 42 *Casz1^fl/fl^;Tg(Sox10-Cre)* mice (n = 5 samples) were used for RNA-seq. 100–320 ng of total RNA was reverse transcribed using Maxima H Minus Reverse Transcriptase (Thermo Fisher Scientific), oligo dT primer, and template switching oligo. Second strand cDNA was amplified using LA Taq DNA polymerase (Takara Bio) for 12 PCR cycles, purified on SPRIselect beads (Beckman Coulter), and eluted with Buffer EB (Qiagen). The concentration and size distribution of second strand cDNA were evaluated using Standard Sensitivity Large Fragment Analysis Kit and a Fragment Analyzer instrument (both from Advanced Analytical). 500 pg of second strand cDNA was tagmented, PCR amplified and indexed using the Nextera XT Library Prep Kit (Illumina). Following SPRIselect bead purification (Beckman Coulter), the size distribution and concentration of cDNA libraries were evaluated using Standard Sensitivity NGS Fragment Analysis Kit (Advanced Analytical) and Fragment Analyzer. All libraries were normalized, pooled and sequenced in a NextSeq2000 instrument (paired end mode; Illumina). Reads were filtered using Trimmomatic (sliding window trimming) to remove low quality sequences (102). Filtered reads were mapped to the mouse genome (GRCm38/mm10) using STAR aligner (v2.7.11a), and mapped reads were quantified using featureCounts (103,104). Genes were filtered out if expression level was not higher than 1 count per million in at least 3 of the 10 samples. Feature counts were compared between genotype groups using edgeR (105). Genes were considered differentially expressed if the adjusted p was lower than 0.001. GO annotations of differentially expressed genes were analyzed for overrepresented GO terms using the DAVID software (106).

### qRT-PCR Analysis

RNA samples were reverse transcribed using SuperScript IV (Thermo Fisher Scientific). The generated cDNA samples were analyzed by qPCR, using the SsoAdvanced Universal SYBR Green Supermix (Bio-Rad) and the primers listed in Table S6. One primer from each qRT-PCR primer pair annealed with an exon-exon junction in the amplified transcript. *Myo6* was used as reference mRNA. Relative mRNA expression values were determined using the ΔΔCT method (107). qRT-PCR analysis of AAV-delivered genes included a correction step based on the detection of AAV-delivered genes in not reverse-transcribed RNA samples (108). qRT-PCR assay efficiency was tested for each primer pair using threefold dilution series of organ of Corti cDNA as template (109). All qRT-PCR assays in this study had efficiencies between 83% and 112%.

### Gene Classification Based on Gene Expression Patterns in the Organ of Corti

Predominantly OHC-expressed, predominantly IHC-expressed, and predominantly OHC–IHC-expressed genes were identified based on organ of Corti scRNA-seq data that had been generated using P1 and P7 WT mice (47). Based on the scRNA-seq data, cell type-specific levels of gene expression were calculated using the Gene Expression Analysis Resource (gEAR) portal (110). Mean gene expression levels were compared among OHCs, IHCs, and types of non-HCs that are located in the organ of Corti (i.e., supporting cells, SCs). Genes were excluded from further analysis if expression levels were lower than 0.3 in OHCs, IHCs, and SCs at both P1 and P7. If the expression level of a gene was lower than 0.3 in OHCs, IHCs, and SCs at one time point only (e.g., P1), gene expression data from the other time point (e.g., P7) were analyzed for fold differences. If this analysis revealed that the expression of the gene was ≥2-fold higher in a HC type (i.e., ‘high expressor HC type’) versus the other HC type and SCs, the gene was classified to be expressed predominantly in the high expressor HC type. If the expression level of a gene was ≥0.3 in OHCs, IHCs, or SCs at both P1 and P7, the time of higher HC expression was identified (i.e., P1 or P7). If the expression level of this gene was ≥2-fold higher in a HC type (i.e., ‘high expressor HC type’) versus the other cell types at the time of higher expression (e.g., P7), expression levels at the other time point (e.g., P1) were also analyzed. If this analysis revealed that the expression level of the gene was ≥1-fold higher in the same (high expressor) HC type versus the other cell types, the gene was classified to be expressed predominantly in the high expressor HC type. If a gene was filtered out from the groups of predominantly OHC-expressed and predominantly IHC-expressed genes solely because the gene expression difference between OHCs and IHCs was less than 2-fold, the gene was classified as predominantly OHC–IHC expressed. The expression levels of these genes were ≥2-fold higher in OHCs and IHCs versus SCs at the time of higher expression, and they were ≥1-fold higher in OHCs and IHCs versus SCs at the time of lower expression (if above the 0.3 cut-off).

### OHC Collection and RNA Extraction

The floor of scala media was dissected from P4 mice and incubated with 5 μM FM1-43 (Thermo Fisher Scientific) in HBSS for ∼15 s to fluorescently label HCs in the dissected tissue. Right after FM1-43 labeling, the tissue pieces were incubated with a mixture of thermolysin (0.2 mg/ml; Sigma-Aldrich) and DNase I (100 unit/ml; Worthington) in DMEM/F-12 for 7–8 minutes at 37°C to facilitate separation of OHC rows from the rest of the tissue. The enzymatic digestion was quenched with an equal volume of ice-cold DMEM/F-12 containing 10% FBS, and OHC-containing groups of cells were released from the tissue pieces by gentle trituration. These groups of cells were selectively pipetted into low-binding barrier tips (MidSci) under a fluorescence microscope (Eclipse TE2000-U; Nikon) and transferred into DMEM/F-12 in ultra-low attachment culture dishes (Corning) as described previously (98). The cell groups were incubated with 0.25% trypsin-EDTA for 15 minutes at 37 °C, and gently triturated every 5 minutes. To inactivate trypsin, an equal volume of 10% FBS (in DMEM/F-12) was added. Single FM1-43-labeled cells were collected from the dishes using selective pipetting. RNA was extracted from the collected cells (∼150 cells per sample) using the RNeasy Micro Kit (Qiagen). RNA was reverse-transcribed using SuperScript IV (Thermo Fisher Scientific), and IHC contamination was tested using an qRT-PCR assay and primers that anneal with the IHC-specific transcript *Fgf8* and the HC-specific control transcript *Myo6*. *Fgf8*-expressing samples were excluded from further analysis.

### Production and Use of AAVs

AAVs (PHP.eB serotype) were produced in HEK293T cells using the triple transfection method (111). HEK293T cells were transfected with the helper plasmid pALD-X80 (Aldevron), the Rep/Cap plasmid pUCmini-iCAP-PHP.eB (112), and a shuttle plasmid using the TransIT-VirusGEN Transfection Reagent (Mirus Bio). Shuttle plasmids were generated by subcloning expression cassettes into ITR-containing plasmids from the Viral Vector Core Facility of the University of Iowa (FBAAVmcsBgHpA plasmid for ssAAV production and pscAAVmcsBgHpA plasmid for scAAV production). The protein-coding expression cassettes were composed of a CMV early enhancer/chicken β-actin (CAG) promoter, a protein coding region, and the bovine growth hormone polyadenylation signal. The encoded proteins were mouse CORO2A (RefSeq: NM_001164804.1), C-terminally Myc-tagged mouse CORO2A, mouse MYL1 (RefSeq: NM_001113387.1), mouse CAR7 (RefSeq: NM_053070.3), and mCherry. To generate an IRES-containing shuttle vector for the production of ssAAV-Coro2a-IRES-mCherry, picornavirus IRES was used (113). The shRNA expression cassettes consisted of a human U6 promoter, the DNA sequence for the shRNA, and a poly(dT) termination sequence. The shRNA sequences were selected using the Broad Institute GPP Web Portal. AAV particles were purified from triple-transfected HEK293T cultures using the minimal purification method (114). Organ of Corti cultures were incubated with the produced AAVs (∼3.7×10^10^ viral genome per Transwell insert) for 24 h on DIV0.

### Luciferase Assays

Firefly luciferase (Fluc)-encoding reporters were used to test the efficacy of shRNAs in transfected HEK293T cells. These shRNA reporters were constructed by subcloning the target sequences of the tested shRNAs and a few flanking nucleotides (30-basepair total length per shRNA target) between the start ATG and a T2A peptide-encoding sequence directly upstream of the Fluc-encoding sequence (Table S6). The translational reading frame was preserved from the start ATG to the 3’ end of the Fluc-encoding sequence (Table S6). The shRNA reporters were co-transfected with shRNA expression vectors and a renilla luciferase (Rluc)-encoding control plasmid (pGL4.74[hRluc/TK]; Promega) into HEK293T cells using Lipofectamin LTX and Plus Reagent (Thermo Fisher Scientific). Fluc and Rluc activities were measured ∼36 h after transfection, using the Dual-Luciferase Reporter Assay System (Promega).

## ACKNOWLEDGMENTS

The *Casz1^fl/fl^* mouse line used for this research project was obtained from the Mutant Mouse Resource and Research Center (MMRRC) at University of California at Davis, an NIH-funded strain repository, and was donated to the MMRRC by Frank Conlon, University of North Carolina at Chapel Hill. We thank Jian Zuo for providing the *Gfi1*^+/*Cre*^ mouse; Christine Blaumueller for critical review of the manuscript; and Viviana Gradinaru for providing the pUCmini-iCAP-PHP.eB plasmid. We acknowledge the use of the University of Iowa Central Microscopy Research Facility, a core resource supported by the University of Iowa Vice President for Research and the Carver College of Medicine. This project was supported by grants from NIDCD/NIH (R01DC014953 to B.B., and R01DC011571 to A.D.) and in part by funds from the NIDCD Division of Intramural Research/NIH (ZIC DC000086 to R.J.M., and DC000059 to M.W.K.). This work utilized computational resources of the NIH HPC Biowulf cluster (https://hpc.nih.gov).

## AUTHOR CONTRIBUTIONS

B.B., Y.N., M.W.K., R.J.M., and A.D. designed the study. R.J.M. and E.T.B. performed the RNA-seq. R.H. and E.C.D. performed the smFISH. C.A., Y.N., and B.B. performed the scanning electron microscopy. B.B., Y.N., and S.W. performed DNA cloning. Y.N. and B.P. performed the ABR assays. Y.N. performed all other experiments. All authors contributed to writing the manuscript.

## REFERENCES

1. Kazmierczak, P., and Müller, U. (2012). Sensing sound: molecules that orchestrate mechanotransduction by hair cells. Trends Neurosci 35, 220–229. 10.1016/j.tins.2011.10.007.

2. Peng, A.W., Salles, F.T., Pan, B., and Ricci, A.J. (2011). Integrating the biophysical and molecular mechanisms of auditory hair cell mechanotransduction. Nat Commun 2, 523. 10.1038/ncomms1533.

3. Gillespie, P.G., and Müller, U. (2009). Mechanotransduction by hair cells: models, molecules, and mechanisms. Cell 139, 33–44. 10.1016/j.cell.2009.09.010.

4. Meyer, A.C., and Moser, T. (2010). Structure and function of cochlear afferent innervation. Curr Opin Otolaryngol Head Neck Surg 18, 441–446. 10.1097/MOO.0b013e32833e0586.

5. Ashmore, J. (2008). Cochlear outer hair cell motility. Physiol Rev 88, 173–210. 10.1152/physrev.00044.2006.

6. LeMasurier, M., and Gillespie, P.G. (2005). Hair-cell mechanotransduction and cochlear amplification. Neuron 48, 403–415. 10.1016/j.neuron.2005.10.017.

7. Oghalai, J.S. (2004). The cochlear amplifier: augmentation of the traveling wave within the inner ear. Curr Opin Otolaryngol Head Neck Surg 12, 431–438. 10.1097/01.moo.0000134449.05454.82.

8. Dallos, P. (2008). Cochlear amplification, outer hair cells and prestin. Curr Opin Neurobiol 18, 370–376. 10.1016/j.conb.2008.08.016.

9. Barr-Gillespie, P.G. (2015). Assembly of hair bundles, an amazing problem for cell biology. Mol Biol Cell 26, 2727–2732. 10.1091/mbc.E14-04-0940.

10. Park, J., and Bird, J.E. (2023). The actin cytoskeleton in hair bundle development and hearing loss. Hear Res 436, 108817. 10.1016/j.heares.2023.108817.

11. Pollock, L.M., and McDermott, B.M., Jr. (2015). The cuticular plate: a riddle, wrapped in a mystery, inside a hair cell. Birth Defects Res C Embryo Today 105, 126–139. 10.1002/bdrc.21098.

12. Du, T.T., Dewey, J.B., Wagner, E.L., Cui, R., Heo, J., Park, J.J., Francis, S.P., Perez-Reyes, E., Guillot, S.J., Sherman, N.E., et al. (2019). LMO7 deficiency reveals the significance of the cuticular plate for hearing function. Nat Commun 10, 1117. 10.1038/s41467-019-09074-4.

13. Liu, Y., Qi, J., Chen, X., Tang, M., Chu, C., Zhu, W., Li, H., Tian, C., Yang, G., Zhong, C., et al. (2019). Critical role of spectrin in hearing development and deafness. Sci Adv 5, eaav7803. 10.1126/sciadv.aav7803.

14. Boussaty, E.C., Ninoyu, Y., Andrade, L.R., Li, Q., Takeya, R., Sumimoto, H., Ohyama, T., Wahlin, K.J., Manor, U., and Friedman, R.A. (2024). Altered Fhod3 expression involved in progressive high-frequency hearing loss via dysregulation of actin polymerization stoichiometry in the cuticular plate. PLoS Genet 20, e1011211. 10.1371/journal.pgen.1011211.

15. Haag, N., Schüler, S., Nietzsche, S., Hübner, C.A., Strenzke, N., Qualmann, B., and Kessels, M.M. (2018). The Actin Nucleator Cobl Is Critical for Centriolar Positioning, Postnatal Planar Cell Polarity Refinement, and Function of the Cochlea. Cell Rep 24, 2418–2431.e2416. 10.1016/j.celrep.2018.07.087.

16. Zhu, G.J., Huang, Y., Zhang, L., Yan, K., Qiu, C., He, Y., Liu, Q., Zhu, C., Morín, M., Moreno-Pelayo, M., et al. (2023). Cingulin regulates hair cell cuticular plate morphology and is required for hearing in human and mouse. EMBO Mol Med 15, e17611. 10.15252/emmm.202317611.

17. Chatterjee, P., Morgan, C.P., Krey, J.F., Benson, C., Goldsmith, J., Bateschell, M., Ricci, A.J., and Barr-Gillespie, P.G. (2023). GIPC3 couples to MYO6 and PDZ domain proteins, and shapes the hair cell apical region. J Cell Sci 136. 10.1242/jcs.261100.

18. Sheffield, A.M., and Smith, R.J.H. (2019). The Epidemiology of Deafness. Cold Spring Harb Perspect Med 9. 10.1101/cshperspect.a033258.

19. Wu, P.Z., O’Malley, J.T., de Gruttola, V., and Liberman, M.C. (2020). Age-Related Hearing Loss Is Dominated by Damage to Inner Ear Sensory Cells, Not the Cellular Battery That Powers Them. J Neurosci 40, 6357–6366. 10.1523/jneurosci.0937-20.2020.

20. Abrashkin, K.A., Izumikawa, M., Miyazawa, T., Wang, C.H., Crumling, M.A., Swiderski, D.L., Beyer, L.A., Gong, T.W., and Raphael, Y. (2006). The fate of outer hair cells after acoustic or ototoxic insults. Hear Res 218, 20–29. 10.1016/j.heares.2006.04.001.

21. Lee, S., Kurioka, T., Lee, M.Y., Beyer, L.A., Swiderski, D.L., Ritter, K.E., and Raphael, Y. (2021). Scar Formation and Debris Elimination during Hair Cell Degeneration in the Adult DTR Mouse. Neuroscience 453, 57–68. 10.1016/j.neuroscience.2020.11.041.

22. Corwin, J.T., and Cotanche, D.A. (1988). Regeneration of sensory hair cells after acoustic trauma. Science 240, 1772–1774. 10.1126/science.3381100.

23. Ryals, B.M., and Rubel, E.W. (1988). Hair cell regeneration after acoustic trauma in adult Coturnix quail. Science 240, 1774–1776. 10.1126/science.3381101.

24. Williams, J.A., and Holder, N. (2000). Cell turnover in neuromasts of zebrafish larvae. Hear Res 143, 171–181. 10.1016/s0378-5955(00)00039-3.

25. Adler, H.J., and Raphael, Y. (1996). New hair cells arise from supporting cell conversion in the acoustically damaged chick inner ear. Neurosci Lett 205, 17–20. 10.1016/0304-3940(96)12367-3.

26. Roberson, D.W., Alosi, J.A., and Cotanche, D.A. (2004). Direct transdifferentiation gives rise to the earliest new hair cells in regenerating avian auditory epithelium. J Neurosci Res 78, 461–471. 10.1002/jnr.20271.

27. Choi, S.W., Abitbol, J.M., and Cheng, A.G. (2024). Hair Cell Regeneration: From Animals to Humans. Clin Exp Otorhinolaryngol 17, 1–14. 10.21053/ceo.2023.01382.

28. Rai, V., Tu, S., Frank, J.R., and Zuo, J. (2021). Molecular Pathways Modulating Sensory Hair Cell Regeneration in Adult Mammalian Cochleae: Progress and Perspectives. Int J Mol Sci 23. 10.3390/ijms23010066.

29. McGovern, M.M., and Cox, B.C. (2025). Hearing restoration through hair cell regeneration: A review of recent advancements and current limitations. Hear Res 461, 109256. 10.1016/j.heares.2025.109256.

30. Wang, J., Zheng, J., Wang, H., He, H., Li, S., Zhang, Y., Wang, Y., Xu, X., and Wang, S. (2023). Gene therapy: an emerging therapy for hair cells regeneration in the cochlea. Front Neurosci 17, 1177791. 10.3389/fnins.2023.1177791.

31. Woods, C., Montcouquiol, M., and Kelley, M.W. (2004). Math1 regulates development of the sensory epithelium in the mammalian cochlea. Nat Neurosci 7, 1310–1318. 10.1038/nn1349.

32. Maricich, S.M., Xia, A., Mathes, E.L., Wang, V.Y., Oghalai, J.S., Fritzsch, B., and Zoghbi, H.Y. (2009). Atoh1-lineal neurons are required for hearing and for the survival of neurons in the spiral ganglion and brainstem accessory auditory nuclei. J Neurosci 29, 11123–11133. 10.1523/jneurosci.2232-09.2009.

33. Bermingham, N.A., Hassan, B.A., Price, S.D., Vollrath, M.A., Ben-Arie, N., Eatock, R.A., Bellen, H.J., Lysakowski, A., and Zoghbi, H.Y. (1999). Math1: an essential gene for the generation of inner ear hair cells. Science 284, 1837–1841. 10.1126/science.284.5421.1837.

34. Kelly, M.C., Chang, Q., Pan, A., Lin, X., and Chen, P. (2012). Atoh1 directs the formation of sensory mosaics and induces cell proliferation in the postnatal mammalian cochlea in vivo. J Neurosci 32, 6699–6710. 10.1523/jneurosci.5420-11.2012.

35. Izumikawa, M., Minoda, R., Kawamoto, K., Abrashkin, K.A., Swiderski, D.L., Dolan, D.F., Brough, D.E., and Raphael, Y. (2005). Auditory hair cell replacement and hearing improvement by Atoh1 gene therapy in deaf mammals. Nat Med 11, 271–276. 10.1038/nm1193.

36. Zheng, J.L., and Gao, W.Q. (2000). Overexpression of Math1 induces robust production of extra hair cells in postnatal rat inner ears. Nat Neurosci 3, 580–586. 10.1038/75753.

37. Liu, Z., Dearman, J.A., Cox, B.C., Walters, B.J., Zhang, L., Ayrault, O., Zindy, F., Gan, L., Roussel, M.F., and Zuo, J. (2012). Age-dependent in vivo conversion of mouse cochlear pillar and Deiters’ cells to immature hair cells by Atoh1 ectopic expression. J Neurosci 32, 6600–6610. 10.1523/jneurosci.0818-12.2012.

38. Krizhanovsky, V., Soreq, L., Kliminski, V., and Ben-Arie, N. (2006). Math1 target genes are enriched with evolutionarily conserved clustered E-box binding sites. J Mol Neurosci 28, 211–229. 10.1385/jmn:28:2:211.

39. Cai, T., Jen, H.I., Kang, H., Klisch, T.J., Zoghbi, H.Y., and Groves, A.K. (2015). Characterization of the transcriptome of nascent hair cells and identification of direct targets of the Atoh1 transcription factor. J Neurosci 35, 5870–5883. 10.1523/jneurosci.5083-14.2015.

40. Chonko, K.T., Jahan, I., Stone, J., Wright, M.C., Fujiyama, T., Hoshino, M., Fritzsch, B., and Maricich, S.M. (2013). Atoh1 directs hair cell differentiation and survival in the late embryonic mouse inner ear. Dev Biol 381, 401–410. 10.1016/j.ydbio.2013.06.022.

41. Jen, H.I., Singh, S., Tao, L., Maunsell, H.R., Segil, N., and Groves, A.K. (2022). GFI1 regulates hair cell differentiation by acting as an off-DNA transcriptional co-activator of ATOH1, and a DNA-binding repressor. Sci Rep 12, 7793. 10.1038/s41598-022-11931-0.

42. Hertzano, R., Montcouquiol, M., Rashi-Elkeles, S., Elkon, R., Yücel, R., Frankel, W.N., Rechavi, G., Möröy, T., Friedman, T.B., Kelley, M.W., and Avraham, K.B. (2004). Transcription profiling of inner ears from Pou4f3(ddl/ddl) identifies Gfi1 as a target of the Pou4f3 deafness gene. Hum Mol Genet 13, 2143–2153. 10.1093/hmg/ddh218.

43. Costa, A., Sanchez-Guardado, L., Juniat, S., Gale, J.E., Daudet, N., and Henrique, D. (2015). Generation of sensory hair cells by genetic programming with a combination of transcription factors. Development 142, 1948–1959. 10.1242/dev.119149.

44. McGovern, M.M., Hosamani, I.V., Niu, Y., Nguyen, K.Y., Zong, C., and Groves, A.K. (2024). Expression of Atoh1, Gfi1, and Pou4f3 in the mature cochlea reprograms nonsensory cells into hair cells. Proc Natl Acad Sci U S A 121, e2304680121. 10.1073/pnas.2304680121.

45. McGovern, M.M., Ghosh, S., Dupuis, C., Walters, B.J., and Groves, A.K. (2024). Reprogramming with Atoh1, Gfi1, and Pou4f3 promotes hair cell regeneration in the adult organ of Corti. PNAS Nexus 3, pgae445. 10.1093/pnasnexus/pgae445.

46. Iyer, A.A., Hosamani, I., Nguyen, J.D., Cai, T., Singh, S., McGovern, M.M., Beyer, L., Zhang, H., Jen, H.I., Yousaf, R., et al. (2022). Cellular reprogramming with ATOH1, GFI1, and POU4F3 implicate epigenetic changes and cell-cell signaling as obstacles to hair cell regeneration in mature mammals. Elife 11. 10.7554/eLife.79712.

47. Kolla, L., Kelly, M.C., Mann, Z.F., Anaya-Rocha, A., Ellis, K., Lemons, A., Palermo, A.T., So, K.S., Mays, J.C., Orvis, J., et al. (2020). Characterization of the development of the mouse cochlear epithelium at the single cell level. Nat Commun 11, 2389. 10.1038/s41467-020-16113-y.

48. Aleksander, S.A., Balhoff, J., Carbon, S., Cherry, J.M., Drabkin, H.J., Ebert, D., Feuermann, M., Gaudet, P., Harris, N.L., Hill, D.P., et al. (2023). The Gene Ontology knowledgebase in 2023. Genetics 224. 10.1093/genetics/iyad031.

49. Scheffer, D.I., Shen, J., Corey, D.P., and Chen, Z.Y. (2015). Gene Expression by Mouse Inner Ear Hair Cells during Development. J Neurosci 35, 6366–6380. 10.1523/jneurosci.5126-14.2015.

50. Costa, A., Powell, L.M., Lowell, S., and Jarman, A.P. (2017). Atoh1 in sensory hair cell development: constraints and cofactors. Semin Cell Dev Biol 65, 60–68. 10.1016/j.semcdb.2016.10.003.

51. Schwab, M.H., Bartholomae, A., Heimrich, B., Feldmeyer, D., Druffel-Augustin, S., Goebbels, S., Naya, F.J., Zhao, S., Frotscher, M., Tsai, M.J., and Nave, K.A. (2000). Neuronal basic helix-loop-helix proteins (NEX and BETA2/Neuro D) regulate terminal granule cell differentiation in the hippocampus. J Neurosci 20, 3714–3724. 10.1523/jneurosci.20-10-03714.2000.

52. Dorr, K.M., Amin, N.M., Kuchenbrod, L.M., Labiner, H., Charpentier, M.S., Pevny, L.H., Wessels, A., and Conlon, F.L. (2015). Casz1 is required for cardiomyocyte G1-to-S phase progression during mammalian cardiac development. Development 142, 2037–2047. 10.1242/dev.119107.

53. Calabi, F., Pannell, R., and Pavloska, G. (2001). Gene targeting reveals a crucial role for MTG8 in the gut. Mol Cell Biol 21, 5658–5666. 10.1128/mcb.21.16.5658-5666.2001.

54. Matsuoka, T., Ahlberg, P.E., Kessaris, N., Iannarelli, P., Dennehy, U., Richardson, W.D., McMahon, A.P., and Koentges, G. (2005). Neural crest origins of the neck and shoulder. Nature 436, 347–355. 10.1038/nature03837.

55. Breuskin, I., Bodson, M., Thelen, N., Thiry, M., Borgs, L., Nguyen, L., Lefebvre, P.P., and Malgrange, B. (2009). Sox10 promotes the survival of cochlear progenitors during the establishment of the organ of Corti. Dev Biol 335, 327–339. 10.1016/j.ydbio.2009.09.007.

56. Wakaoka, T., Motohashi, T., Hayashi, H., Kuze, B., Aoki, M., Mizuta, K., Kunisada, T., and Ito, Y. (2013). Tracing Sox10-expressing cells elucidates the dynamic development of the mouse inner ear. Hear Res 302, 17–25. 10.1016/j.heares.2013.05.003.

57. Shone, G., Raphael, Y., and Miller, J.M. (1991). Hereditary deafness occurring in cd/1 mice. Hear Res 57, 153–156. 10.1016/0378-5955(91)90084-m.

58. Zheng, Q.Y., Johnson, K.R., and Erway, L.C. (1999). Assessment of hearing in 80 inbred strains of mice by ABR threshold analyses. Hear Res 130, 94–107. 10.1016/s0378-5955(99)00003-9.

59. Johnson, K.R., Zheng, Q.Y., and Erway, L.C. (2000). A major gene affecting age-related hearing loss is common to at least ten inbred strains of mice. Genomics 70, 171–180. 10.1006/geno.2000.6377.

60. Hackney, C.M., Mahendrasingam, S., Penn, A., and Fettiplace, R. (2005). The concentrations of calcium buffering proteins in mammalian cochlear hair cells. J Neurosci 25, 7867–7875. 10.1523/jneurosci.1196-05.2005.

61. Chessum, L., Matern, M.S., Kelly, M.C., Johnson, S.L., Ogawa, Y., Milon, B., McMurray, M., Driver, E.C., Parker, A., Song, Y., et al. (2018). Helios is a key transcriptional regulator of outer hair cell maturation. Nature 563, 696–700. 10.1038/s41586-018-0728-4.

62. Liu, H., Chen, L., Giffen, K.P., Stringham, S.T., Li, Y., Judge, P.D., Beisel, K.W., and He, D.Z.Z. (2018). Cell-Specific Transcriptome Analysis Shows That Adult Pillar and Deiters’ Cells Express Genes Encoding Machinery for Specializations of Cochlear Hair Cells. Front Mol Neurosci 11, 356. 10.3389/fnmol.2018.00356.

63. Ranum, P.T., Goodwin, A.T., Yoshimura, H., Kolbe, D.L., Walls, W.D., Koh, J.Y., He, D.Z.Z., and Smith, R.J.H. (2019). Insights into the Biology of Hearing and Deafness Revealed by Single-Cell RNA Sequencing. Cell Rep 26, 3160–3171.e3163. 10.1016/j.celrep.2019.02.053.

64. Liu, Z., Owen, T., Zhang, L., and Zuo, J. (2010). Dynamic expression pattern of Sonic hedgehog in developing cochlear spiral ganglion neurons. Dev Dyn 239, 1674–1683. 10.1002/dvdy.22302.

65. Yang, H., Gan, J., Xie, X., Deng, M., Feng, L., Chen, X., Gao, Z., and Gan, L. (2010). Gfi1-Cre knock-in mouse line: A tool for inner ear hair cell-specific gene deletion. Genesis 48, 400–406. 10.1002/dvg.20632.

66. Matern, M., Vijayakumar, S., Margulies, Z., Milon, B., Song, Y., Elkon, R., Zhang, X., Jones, S.M., and Hertzano, R. (2017). Gfi1(Cre) mice have early onset progressive hearing loss and induce recombination in numerous inner ear non-hair cells. Sci Rep 7, 42079. 10.1038/srep42079.

67. Nakano, Y., Kim, S.H., Kim, H.M., Sanneman, J.D., Zhang, Y., Smith, R.J., Marcus, D.C., Wangemann, P., Nessler, R.A., and Bánfi, B. (2009). A claudin-9-based ion permeability barrier is essential for hearing. PLoS Genet 5, e1000610. 10.1371/journal.pgen.1000610.

68. Jan, T.A., Eltawil, Y., Ling, A.H., Chen, L., Ellwanger, D.C., Heller, S., and Cheng, A.G. (2021). Spatiotemporal dynamics of inner ear sensory and non-sensory cells revealed by single-cell transcriptomics. Cell Rep 36, 109358. 10.1016/j.celrep.2021.109358.

69. Tarchini, B. (2021). A Reversal in Hair Cell Orientation Organizes Both the Auditory and Vestibular Organs. Front Neurosci 15, 695914. 10.3389/fnins.2021.695914.

70. Leibovici, M., Verpy, E., Goodyear, R.J., Zwaenepoel, I., Blanchard, S., Lainé, S., Richardson, G.P., and Petit, C. (2005). Initial characterization of kinocilin, a protein of the hair cell kinocilium. Hear Res 203, 144–153. 10.1016/j.heares.2004.12.002.

71. Ross, A.J., May-Simera, H., Eichers, E.R., Kai, M., Hill, J., Jagger, D.J., Leitch, C.C., Chapple, J.P., Munro, P.M., Fisher, S., et al. (2005). Disruption of Bardet-Biedl syndrome ciliary proteins perturbs planar cell polarity in vertebrates. Nat Genet 37, 1135–1140. 10.1038/ng1644.

72. Ezan, J., Lasvaux, L., Gezer, A., Novakovic, A., May-Simera, H., Belotti, E., Lhoumeau, A.C., Birnbaumer, L., Beer-Hammer, S., Borg, J.P., et al. (2013). Primary cilium migration depends on G-protein signalling control of subapical cytoskeleton. Nat Cell Biol 15, 1107–1115. 10.1038/ncb2819.

73. Jones, C., Roper, V.C., Foucher, I., Qian, D., Banizs, B., Petit, C., Yoder, B.K., and Chen, P. (2008). Ciliary proteins link basal body polarization to planar cell polarity regulation. Nat Genet 40, 69–77. 10.1038/ng.2007.54.

74. Sipe, C.W., and Lu, X. (2011). Kif3a regulates planar polarization of auditory hair cells through both ciliary and non-ciliary mechanisms. Development 138, 3441–3449. 10.1242/dev.065961.

75. Jagger, D., Collin, G., Kelly, J., Towers, E., Nevill, G., Longo-Guess, C., Benson, J., Halsey, K., Dolan, D., Marshall, J., et al. (2011). Alström Syndrome protein ALMS1 localizes to basal bodies of cochlear hair cells and regulates cilium-dependent planar cell polarity. Hum Mol Genet 20, 466–481. 10.1093/hmg/ddq493.

76. Moon, K.H., Ma, J.H., Min, H., Koo, H., Kim, H., Ko, H.W., and Bok, J. (2020). Dysregulation of sonic hedgehog signaling causes hearing loss in ciliopathy mouse models. Elife 9. 10.7554/eLife.56551.

77. May-Simera, H.L., Petralia, R.S., Montcouquiol, M., Wang, Y.X., Szarama, K.B., Liu, Y., Lin, W., Deans, M.R., Pazour, G.J., and Kelley, M.W. (2015). Ciliary proteins Bbs8 and Ift20 promote planar cell polarity in the cochlea. Development 142, 555–566. 10.1242/dev.113696.

78. Marshall, T.W., Aloor, H.L., and Bear, J.E. (2009). Coronin 2A regulates a subset of focal-adhesion-turnover events through the cofilin pathway. J Cell Sci 122, 3061–3069. 10.1242/jcs.051482.

79. Huang, W., Ghisletti, S., Saijo, K., Gandhi, M., Aouadi, M., Tesz, G.J., Zhang, D.X., Yao, J., Czech, M.P., Goode, B.L., et al. (2011). Coronin 2A mediates actin-dependent de-repression of inflammatory response genes. Nature 470, 414–418. 10.1038/nature09703.

80. Rastetter, R.H., Blömacher, M., Drebber, U., Marko, M., Behrens, J., Solga, R., Hojeili, S., Bhattacharya, K., Wunderlich, C.M., Wunderlich, F.T., et al. (2015). Coronin 2A (CRN5) expression is associated with colorectal adenoma-adenocarcinoma sequence and oncogenic signalling. BMC Cancer 15, 638. 10.1186/s12885-015-1645-7.

81. Dhawan, K., Naslavsky, N., and Caplan, S. (2022). Coronin2A links actin-based endosomal processes to the EHD1 fission machinery. Mol Biol Cell 33, ar107. 10.1091/mbc.E21-12-0624.

82. Richard, E.M., Maurice, T., and Delprat, B. (2023). Calcium signaling and genetic rare diseases: An auditory perspective. Cell Calcium 110, 102702. 10.1016/j.ceca.2023.102702.

83. McCarty, D.M. (2008). Self-complementary AAV vectors; advances and applications. Mol Ther 16, 1648–1656. 10.1038/mt.2008.171.

84. de Solis, C.A., Holehonnur, R., Banerjee, A., Luong, J.A., Lella, S.K., Ho, A., Pahlavan, B., and Ploski, J.E. (2015). Viral delivery of shRNA to amygdala neurons leads to neurotoxicity and deficits in Pavlovian fear conditioning. Neurobiol Learn Mem 124, 34–47. 10.1016/j.nlm.2015.07.005.

85. Grimm, D., Streetz, K.L., Jopling, C.L., Storm, T.A., Pandey, K., Davis, C.R., Marion, P., Salazar, F., and Kay, M.A. (2006). Fatality in mice due to oversaturation of cellular microRNA/short hairpin RNA pathways. Nature 441, 537–541. 10.1038/nature04791.

86. Mattar, P., Jolicoeur, C., Dang, T., Shah, S., Clark, B.S., and Cayouette, M. (2021). A Casz1-NuRD complex regulates temporal identity transitions in neural progenitors. Sci Rep 11, 3858. 10.1038/s41598-021-83395-7.

87. Liu, Z., Lam, N., and Thiele, C.J. (2015). Zinc finger transcription factor CASZ1 interacts with histones, DNA repair proteins and recruits NuRD complex to regulate gene transcription. Oncotarget 6, 27628–27640. 10.18632/oncotarget.4733.

88. Liu, Z., Lam, N., Wang, E., Virden, R.A., Pawel, B., Attiyeh, E.F., Maris, J.M., and Thiele, C.J. (2017). Identification of CASZ1 NES reveals potential mechanisms for loss of CASZ1 tumor suppressor activity in neuroblastoma. Oncogene 36, 97–109. 10.1038/onc.2016.179.

89. Layman, W.S., Sauceda, M.A., and Zuo, J. (2013). Epigenetic alterations by NuRD and PRC2 in the neonatal mouse cochlea. Hear Res 304, 167–178. 10.1016/j.heares.2013.07.017.

90. Sun, Y., Ren, M., Zhang, Y., Li, S., Luo, Z., Sun, S., He, S., Wang, G., Zhang, D., Mansour, S.L., et al. (2025). Casz1 is required for both inner hair cell fate stabilization and outer hair cell survival. Science 388, eado4930. 10.1126/science.ado4930.

91. Xie, W.R., Jen, H.I., Seymour, M.L., Yeh, S.Y., Pereira, F.A., Groves, A.K., Klisch, T.J., and Zoghbi, H.Y. (2017). An Atoh1-S193A Phospho-Mutant Allele Causes Hearing Deficits and Motor Impairment. J Neurosci 37, 8583–8594. 10.1523/jneurosci.0295-17.2017.

92. Ahn, M., Witting, S.R., Ruiz, R., Saxena, R., and Morral, N. (2011). Constitutive expression of short hairpin RNA in vivo triggers buildup of mature hairpin molecules. Hum Gene Ther 22, 1483–1497. 10.1089/hum.2010.234.

93. Wallis, D., Hamblen, M., Zhou, Y., Venken, K.J., Schumacher, A., Grimes, H.L., Zoghbi, H.Y., Orkin, S.H., and Bellen, H.J. (2003). The zinc finger transcription factor Gfi1, implicated in lymphomagenesis, is required for inner ear hair cell differentiation and survival. Development 130, 221–232. 10.1242/dev.00190.

94. Matern, M.S., Milon, B., Lipford, E.L., McMurray, M., Ogawa, Y., Tkaczuk, A., Song, Y., Elkon, R., and Hertzano, R. (2020). GFI1 functions to repress neuronal gene expression in the developing inner ear hair cells. Development 147. 10.1242/dev.186015.

95. Xiang, M., Gao, W.Q., Hasson, T., and Shin, J.J. (1998). Requirement for Brn-3c in maturation and survival, but not in fate determination of inner ear hair cells. Development 125, 3935–3946. 10.1242/dev.125.20.3935.

96. Yu, H.V., Tao, L., Llamas, J., Wang, X., Nguyen, J.D., Trecek, T., and Segil, N. (2021). POU4F3 pioneer activity enables ATOH1 to drive diverse mechanoreceptor differentiation through a feed-forward epigenetic mechanism. Proc Natl Acad Sci U S A 118. 10.1073/pnas.2105137118.

97. Taniguchi, H., He, M., Wu, P., Kim, S., Paik, R., Sugino, K., Kvitsiani, D., Fu, Y., Lu, J., Lin, Y., et al. (2011). A resource of Cre driver lines for genetic targeting of GABAergic neurons in cerebral cortex. Neuron 71, 995–1013. 10.1016/j.neuron.2011.07.026.

98. Nakano, Y., Wiechert, S., Fritzsch, B., and Bánfi, B. (2020). Inhibition of a transcriptional repressor rescues hearing in a splicing factor-deficient mouse. Life Sci Alliance 3. 10.26508/lsa.202000841.

99. Nakano, Y., Kelly, M.C., Rehman, A.U., Boger, E.T., Morell, R.J., Kelley, M.W., Friedman, T.B., and Bánfi, B. (2018). Defects in the Alternative Splicing-Dependent Regulation of REST Cause Deafness. Cell 174, 536–548.e521. 10.1016/j.cell.2018.06.004.

100. Nakano, Y., Longo-Guess, C.M., Bergstrom, D.E., Nauseef, W.M., Jones, S.M., and Bánfi, B. (2008). Mutation of the Cyba gene encoding p22phox causes vestibular and immune defects in mice. J Clin Invest 118, 1176–1185. 10.1172/jci33835.

101. Meyers, J.R., MacDonald, R.B., Duggan, A., Lenzi, D., Standaert, D.G., Corwin, J.T., and Corey, D.P. (2003). Lighting up the senses: FM1-43 loading of sensory cells through nonselective ion channels. J Neurosci 23, 4054–4065. 10.1523/jneurosci.23-10-04054.2003.

102. Bolger, A.M., Lohse, M., and Usadel, B. (2014). Trimmomatic: a flexible trimmer for Illumina sequence data. Bioinformatics 30, 2114–2120. 10.1093/bioinformatics/btu170.

103. Dobin, A., Davis, C.A., Schlesinger, F., Drenkow, J., Zaleski, C., Jha, S., Batut, P., Chaisson, M., and Gingeras, T.R. (2013). STAR: ultrafast universal RNA-seq aligner. Bioinformatics 29, 15–21. 10.1093/bioinformatics/bts635.

104. Liao, Y., Smyth, G.K., and Shi, W. (2014). featureCounts: an efficient general purpose program for assigning sequence reads to genomic features. Bioinformatics 30, 923–930. 10.1093/bioinformatics/btt656.

105. Robinson, M.D., McCarthy, D.J., and Smyth, G.K. (2010). edgeR: a Bioconductor package for differential expression analysis of digital gene expression data. Bioinformatics 26, 139–140. 10.1093/bioinformatics/btp616.

106. Huang da, W., Sherman, B.T., and Lempicki, R.A. (2009). Systematic and integrative analysis of large gene lists using DAVID bioinformatics resources. Nat Protoc 4, 44–57. 10.1038/nprot.2008.211.

107. Livak, K.J., and Schmittgen, T.D. (2001). Analysis of relative gene expression data using real-time quantitative PCR and the 2(-Delta Delta C(T)) Method. Methods 25, 402–408. 10.1006/meth.2001.1262.

108. Laurell, H., Iacovoni, J.S., Abot, A., Svec, D., Maoret, J.J., Arnal, J.F., and Kubista, M. (2012). Correction of RT-qPCR data for genomic DNA-derived signals with ValidPrime. Nucleic Acids Res 40, e51. 10.1093/nar/gkr1259.

109. Bustin, S.A., Benes, V., Garson, J.A., Hellemans, J., Huggett, J., Kubista, M., Mueller, R., Nolan, T., Pfaffl, M.W., Shipley, G.L., et al. (2009). The MIQE guidelines: minimum information for publication of quantitative real-time PCR experiments. Clin Chem 55, 611–622. 10.1373/clinchem.2008.112797.

110. Orvis, J., Gottfried, B., Kancherla, J., Adkins, R.S., Song, Y., Dror, A.A., Olley, D., Rose, K., Chrysostomou, E., Kelly, M.C., et al. (2021). gEAR: Gene Expression Analysis Resource portal for community-driven, multi-omic data exploration. Nat Methods 18, 843–844. 10.1038/s41592-021-01200-9.

111. Ayuso, E., Mingozzi, F., and Bosch, F. (2010). Production, purification and characterization of adeno-associated vectors. Curr Gene Ther 10, 423–436. 10.2174/156652310793797685.

112. Chan, K.Y., Jang, M.J., Yoo, B.B., Greenbaum, A., Ravi, N., Wu, W.L., Sánchez-Guardado, L., Lois, C., Mazmanian, S.K., Deverman, B.E., and Gradinaru, V. (2017). Engineered AAVs for efficient noninvasive gene delivery to the central and peripheral nervous systems. Nat Neurosci 20, 1172–1179. 10.1038/nn.4593.

113. Nakano, Y., Wiechert, S., and Bánfi, B. (2019). Overlapping Activities of Two Neuronal Splicing Factors Switch the GABA Effect from Excitatory to Inhibitory by Regulating REST. Cell Rep 27, 860–871.e868. 10.1016/j.celrep.2019.03.072.

114. Konno, A., and Hirai, H. (2020). Efficient whole brain transduction by systemic infusion of minimally purified AAV-PHP.eB. J Neurosci Methods 346, 108914. 10.1016/j.jneumeth.2020.108914.

