## Supplemental Figure Legends for "CASZ1 regulates the rate at which outer hair cells mature and is required for hearing"

**Figure S1. In *Casz1*$^{fl/fl}$*;Tg(Sox10-Cre)* mice, hearing loss is progressive.**

(A) F-actin staining (green) and tdTomato immunostaining (red) of organ of Corti samples from P0 mice of the indicated genotypes. Arrowheads, IHC rows. Vertical black lines, OHC rows. Scale bar, 10 μm.

(B) qRT-PCR data for differences in *Casz1* expression levels in organs of Corti from *Casz1*$^{fl/fl}$*;Tg(Sox10-Cre)* mice versus WT mice (P4). Each symbol represents an expression difference between organs of Corti from one *Casz1*$^{fl/fl}$*;Tg(Sox10-Cre)* mouse versus organs of Corti pooled from 3 WT mice. Values are mean ± SEM. One sample t-test, theoretical mean = 0, *p<0.0001.

(C) qRT-PCR data for differences in *Runx1t1* expression levels in organs of Corti from *Runx1t1*$^{fl/fl}$*;Tg(Sox10-Cre)* mice versus WT mice (P4). Each symbol represents an expression difference between organs of Corti from one *Runx1t1*$^{fl/fl}$*;Tg(Sox10-Cre)* mouse versus organs of Corti pooled from 3 WT mice. Values are mean ± SEM. One sample t-test, theoretical mean = 0, *p=0.0072.

(D) Statistical analysis of auditory brainstem response (ABR) data shown in Figure 1E, for differences in ABR thresholds between mice of the indicated genotypes at P60. Values are mean ± SEM (Kolmogorov-Smirnov test with Holm-Sidak correction *p < 0.05).

(E and F) Statistical analysis of ABR data shown in Figure 1E, for age-dependent differences in ABR thresholds within the indicated genotype groups. Values are mean ± SEM. Three-group comparisons in (E): two-way ANOVA (p < 0.0001 for time factor) and Dunnett's post hoc test *p < 0.05 (reference group, P60). Two-group comparisons in (F): Kolmogorov-Smirnov test with Holm-Sidak correction (ns, not significant).

**Figure S2. The *Calb2*$^{IRES\text{-}Cre}$ allele drives loxP site recombination in OHCs during the first postnatal week.**

(A) Mouse (WT P1) cochlear section hybridized with *Casz1* complementary (green and white as indicated) and *Pvalb* complementary (red) smFISH probes and imaged at low magnification. Orange frame indicates the area shown in Figure 2B. DAPI staining, blue. Dashed ellipses, spiral ganglion. Arrowheads, IHCs. Brackets, OHCs. Scale bar, 100 μm.

(B) F-actin staining (green) and tdTomato immunostaining (red) of organ of Corti samples from P0 and P6 mice of the indicated genotypes. Arrowheads, IHC rows. Horizontal black lines, OHC rows. Scale bars, 20 µm.

(C) qRT-PCR data for *Casz1* expression differences in spiral ganglions from *Casz1*$^{fl/fl}$;*Shh*$^{+/Cre}$ mice versus control (*Casz1*$^{fl/fl}$) mice (P1). Each symbol represents expression level difference between spiral ganglions from one *Casz1*$^{fl/fl}$;*Shh*$^{+/Cre}$ mouse versus spiral ganglions pooled from 3 *Casz1*$^{fl/fl}$ mice. Values are mean ± SEM (one-sample t test *p = 0.008, theoretical mean = 0).

(D) qRT-PCR data for *Casz1* expression differences in organs of Corti from cKO mice of the indicated genotypes versus control (*Casz1*$^{fl/fl}$) mice (P8). Each symbol represents expression level difference between organs of Corti from one cKO mouse versus organs of Corti pooled from 4 *Casz1*$^{fl/fl}$ mice. Values are mean ± SEM (one-sample t tests, FDR-adjusted *p = 0.0002 and **p=0.0013, theoretical mean = 0).

(E) Statistical analysis of ABR data shown in Figure 2F, for age-dependent differences in ABR thresholds within the indicated genotype groups. Values are mean ± SEM. Two-way ANOVA ($p < 0.0001$ for time factor) and Dunnett's post hoc test *$p < 0.05$ (reference group, P60).

**Figure S3. Third-row stereocilia are abnormally thick in some IHCs of *Casz1*$^{fl/fl}$*;Tg(Sox10-Cre)* mice.**

(A) Numbers of IHCs in the indicated turns of the cochlea in control (*Casz1*$^{fl/fl}$), *Casz1*$^{fl/fl}$*;Tg(Sox10-Cre)*, and *Casz1*$^{fl/fl}$*;Gfi1*$^{+/Cre}$ mice (P60). Each symbol represents value for one mouse.

(B) F-actin staining of OHC rows in organ of Corti samples from P6 and P14 mice of the indicated genotypes. White corners indicate areas that are shown in Figure 3D. Scale bar, 10 µm.

(C) Stacked display of lines drawn over tips of second-row stereocilia of OHCs of control (*Casz1*$^{fl/fl}$) and *Casz1*$^{fl/fl}$*;Tg(Sox10-Cre)* mice (P14).

(D) F-actin staining of IHCs in organs of Corti from P6 and P14 mice of the indicated genotypes. Green corners indicate areas that are magnified in green frames. Arrowheads indicate third-row stereocilia in magnified areas. White scale bar, 5 µm. Green scale bar, 1 µm.

(E) Quantification of IHC stereocilia bundles that include 3 or more abnormally thick third-row stereocilia (≥220-nm wide) in P14 mice of the indicated genotypes. Stereocilia were visualized as in (D). Each symbol represents value for a single mouse (n = 40–50 IHCs analyzed per mouse).

(F) Numbers of stubby and missing stereocilia in the third row of stereocilia bundles in OHCs of control (*Casz1$^{fl/fl}$*) and *Casz1$^{fl/fl}$;Tg(Sox10-Cre)* mice (P14). The expected number of third-row stereocilia was equal to the number of second-row stereocilia in the same cell. Each symbol represents value for a single bundle of stereocilia. Horizontal gray lines indicate medians (Welch's test *p < 0.0001).

**Figure S4. Conditional deletion of *Casz1* causes gene expression alterations and kinocilium defects in HCs.**

(A) Differential expression of predominantly OHC–IHC-expressed genes in organ of Corti samples from *Casz1$^{fl/fl}$;Tg(Sox10-Cre)* versus WT mice (P4), as revealed by RNA-seq. Dots represent genes that are expressed at abnormally high (red) or low (blue) levels. *Nf2* and *Car7* indicate genes that were selected for further testing in Figure 4E.

(B) qRT-PCR data for differential expression of the indicated genes in organ of Corti samples from *Casz1$^{fl/fl}$;Tg(Sox10-Cre)* versus WT mice (blue bars) and from *Casz1$^{fl/fl}$;Gfi1$^{+/Cre}$* versus WT mice (green bars) (P4). Genes with similar expression patterns in the WT organ of Corti are grouped together as indicated. Each symbol represents expression level of an indicated gene in the organs of Corti of one cKO mouse relative to the expression level of the same gene in organs of Corti pooled from 3 WT mice. Gray line indicates zero expression difference (one-sample t tests, FDR-adjusted *p < 0.05, theoretical mean = 0).

(C) Differential expression of the indicated genes in OHCs from *Casz1$^{fl/fl}$;Tg(Sox10-Cre)* versus WT mice (P4), as determined by qRT-PCR. Each symbol represents expression of an indicated gene in one *Casz1$^{fl/fl}$;Tg(Sox10-Cre)* sample relative to the averaged expression of the same gene in 3 WT samples (one-sample t tests, FDR-adjusted *p < 0.05, theoretical mean = 0).

(D) Immunofluorescence detection of OCM (red and white as indicated) and MYO7A (green) in organ of Corti samples from P30 control (*Casz1$^{fl/fl}$*) and *Casz1$^{fl/fl}$;Tg(Sox10-Cre)* mice. Arrowheads, IHC rows. Vertical lines, OHC rows. Scale bar, 20 μm.

(E) Violin plots of OCM immunofluorescence (IF) intensities in OHCs and IHCs of mice of the indicated genotypes at P6 and P30. OCM IF intensities were normalized to MYO7A IF intensities in the same cells. Each symbol represents value for one OHC. Dashed gray lines indicate medians. Mann-Whitney tests, FDR-adjusted *p < 0.0001.

(F) Enrichment of GO annotations in the group of genes that are differentially expressed in organs of Corti of *Casz1$^{fl/fl}$;Tg(Sox10-Cre)* versus WT mice at P4.

(G) Lengths of kinocilia in OHCs and IHCs of control (*Casz1$^{fl/fl}$*) and *Casz1$^{fl/fl}$*;*Tg(Sox10-Cre)* mice (P6). Each symbol represents the length of a single kinocilium. Gray dashed lines indicate medians. Mann-Whitney tests, FDR-adjusted *p = 0.0002, **p < 0.0001.

(H, I, J, and K) Percentages of kinociliated OHCs (H, I) and IHCs (J, K) in the indicated turns of the cochlea in *Casz1$^{fl/fl}$*;*Tg(Sox10-Cre)* and *Casz1$^{fl/fl}$* mice at P10 (H, J) and P14 (I, K). Each symbol represents value for a single mouse (n = 33–40 IHCs and 95–120 OHCs per turn of cochlea per mouse). Gray lines indicate mean ± SEM. Two-way ANOVA genotype factor p<0.0001 (H and K) and p=0.0012 (J), Sidak's post hoc test *p=0.017, **p=0.0056, and ***p<0.0001.

**Figure S5. Overexpression of *Coro2a* causes loss of F-actin from the cuticular plate in OHCs.**

(A) Volcano plot of a subset of gene expression differences between organs of Corti of *Casz1$^{fl/fl}$*;*Tg(Sox10-Cre)* and WT mice, selected based on annotations of genes with the GO term 'sensory perception of sound'. The colors of dots indicate abnormally high (red) and abnormally low (blue) expression levels. Gray lines indicate 2-fold expression difference.

(B) Expression levels of *Coro2a*, *Car7* and *Myl1* in OHCs and IHCs of WT mice at the indicated times, as revealed by previous scRNA-seq analysis (Kolla et al., 2020). Depicted are median (red line), quartile 1–3 range (Q1–Q3, gray), and mean (circles) gene expression levels.

(C) qRT-PCR validation of overexpression of AAV-delivered genes in DIV10 organ of Corti cultures that were derived from WT mice (E17.5) and incubated with the indicated AAVs on DIV0. Each circle represents expression level of the indicated gene in one organ of Corti culture incubated with a gene of interest-delivering AAV (AAV-Coro2a, AAV-Car7, or AAV-Myl1) versus averaged expression level of the same gene in three organ of Corti cultures incubated with AAV-mCherry. Data were corrected for detection of *Coro2a*, *Car7*, and *Myl1* in not reverse-transcribed ($RT^-$) samples. Bars indicate means.

(D) F-actin staining (green) and mCherry immunostaining (red) of DIV11 organ of Corti cultures that were derived from WT mice (E17.5) and incubated with the indicated AAVs on DIV0. Blue corners indicate areas shown in Figure 5D. Scale bar, 10 μm.

(E) Quantification of stereocilia bundles that include >1 μm gaps between first-row stereocilia in mCherry-expressing OHCs in AAV-transduced organ of Corti cultures. The organ of Corti cultures were derived from WT mice (E17.5), incubated with the indicated AAVs on DIV0, and used for F-actin staining and mCherry immunostaining on DIV7 (white bars) or DIV11 (gray bars). Each circle represents value for one organ of Corti culture.

(F) F-actin staining (green) and mCherry immunostaining (red) of a DIV7 organ of Corti culture that was derived from a WT mouse (E17.5) and incubated with AAV-Coro2a-IRES-mCherry on DIV0. Blue arrowheads indicate the place selected for visualization in the XZ-plane in (G). Scale bar, 10 μm.

(G) Visualization of F-actin staining (green and white as indicated) and mCherry immunostaining (red) in the XZ-plane between points that are indicated by blue arrowheads in (F). Yellow corners indicate 2 × 2 μm areas at the centers of cuticular plates.

(H) F-actin staining (green) of organ of Corti samples from P6 mice of the indicated genotypes. Blue corners indicate areas shown in Figure 5G. Arrowheads, IHC rows. Vertical black lines, OHC rows. Scale bar, 10 μm.

(I) Normalized intensities of F-actin staining in the middle areas of IHC cuticular plates in mice of the indicate genotypes at P6. Each symbol represents fluorescence intensity in a 2 × 2 μm area (in the XZ-plane) of one cuticular plate divided by the average fluorescence intensity in 2 × 2 μm apical regions (in the XZ-plane) of inner sulcus cells in the same image stack. Mann-Whitney test; ns, not significant.

**Figure S6. The majority of tested shRNAs are toxic to both OHCs and IHCs.**

(A and B) Sequences of *Coro2a*-targeting shRNAs (A), *Calb1*-targeting shRNAs (B), and positions of guide strand complementary regions (gray connecting lines) in the coding sequences (cds) of *Coro2a* (A) and *Calb1* (B). Also indicated are the lengths of coding sequences, the predicted Dicer cleavage sites (purple arrowheads), the predicted base pair formations in the shRNAs (vertical lines), and the shRNA bases that are non-complementary to the target sequence in the predicted guide strand (underlined).

(C) CALB1 immunofluorescence (IF) intensities in OHC cuticular plates in DIV9 organ of Corti cultures that were derived from WT mice (E17.5) and incubated with the indicated AAVs on DIV0. CALB1 IF intensities were quantified relative to the average IF intensity in the two most brightly fluorescent OHC cuticular plates in the image. N = 120 OHCs per group.

(D) Graphical representation of structural alterations in OHC stereocilia bundles that are categorized as indicators of loss of W / V-like shape.

(E) Graphical representation of structural alterations in IHC stereocilia bundles that are categorized as indicators of degeneration.

(F) Shapes of stereocilia bundles in OHCs (circles, left y-axis) and IHCs (squares, right y-axis) in WT organ of Corti cultures (DIV9) that were incubated with scAAV-shRNAs or no AAV (–) on DIV0. The scAAV-delivered shRNAs are indicated. Symbol represent data for organ of Corti cultures from separate mice. One-way ANOVA $*p<0.0001$, Dunnett's post hoc test $*p <0.0001$; ns, not significant; control group, no shRNA (–). Dashed horizontal lines, means.

(G) *Coro2a* expression differences between scAAV-shRNA-incubated (transduced) and not AAV-incubated (non-transduced) organ of Corti cultures that were derived from *Casz1*$^{fl/fl}$*;Tg(Sox10-Cre)* mice (E17.5). The organ of Corti cultures were incubated with the indicated scAAV-shRNAs (or no AAV) on DIV0 and tested for *Coro2a* expression on DIV9. Circles represent data for organ of Corti cultures from separate mice (one sample t test $*p = 0.0041$, hypothetical mean = 0; ns, not significant). Dashed horizontal lines indicate means.
