## Supplementary figures and images for "CASZ1 regulates the rate at which outer hair cells mature and is required for hearing"

### Figure S1

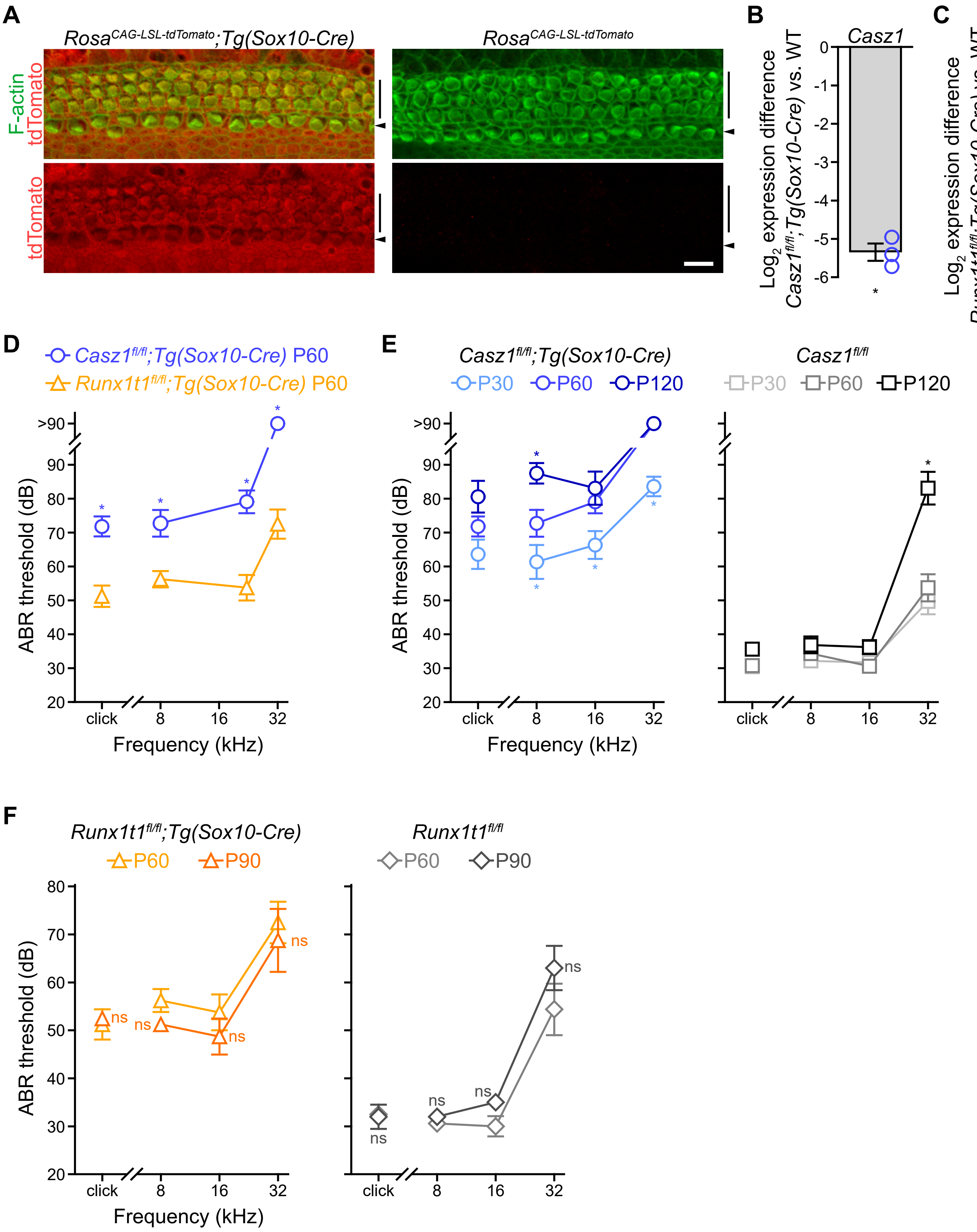

### Figure S2

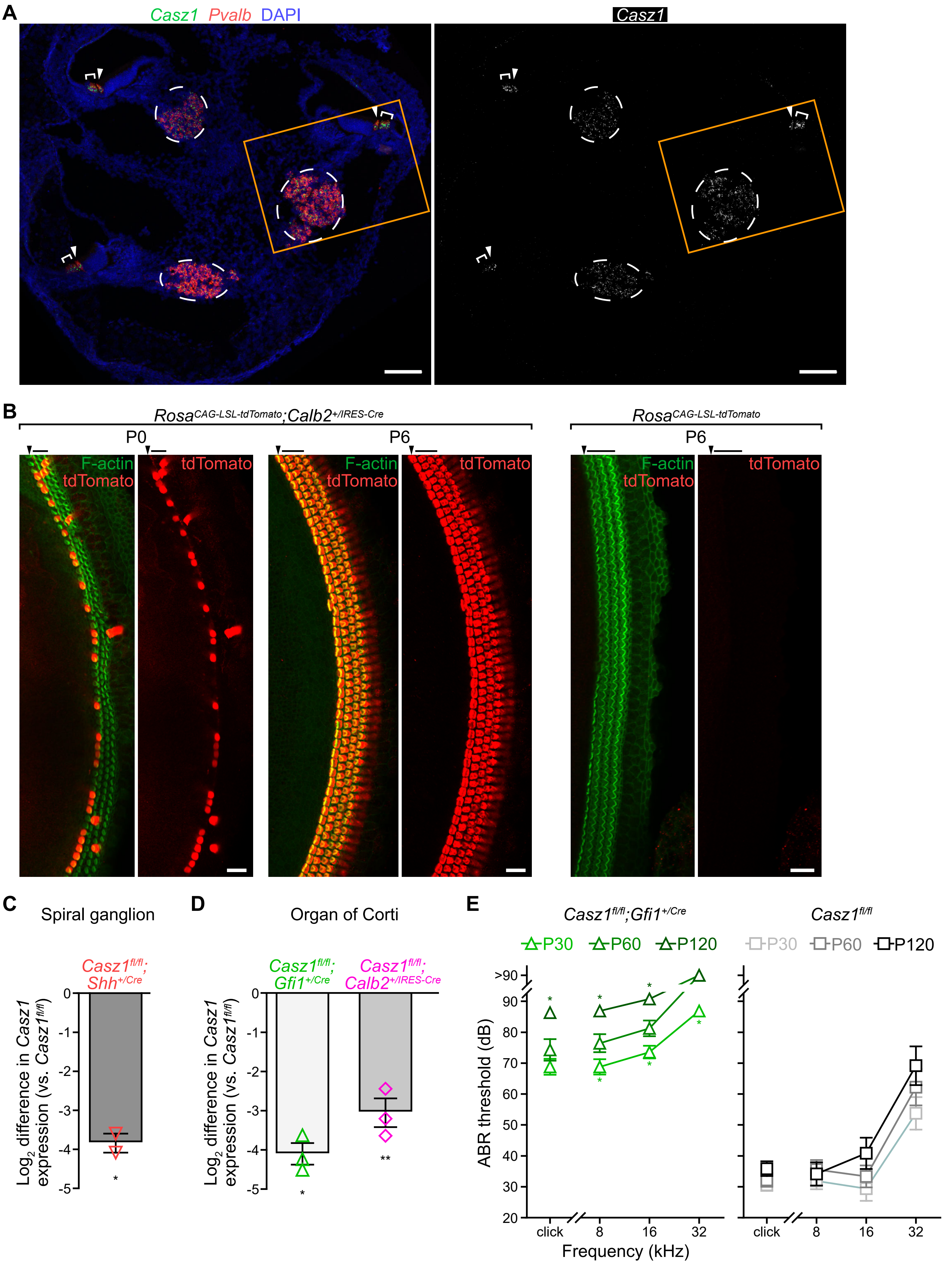

### Figure S3

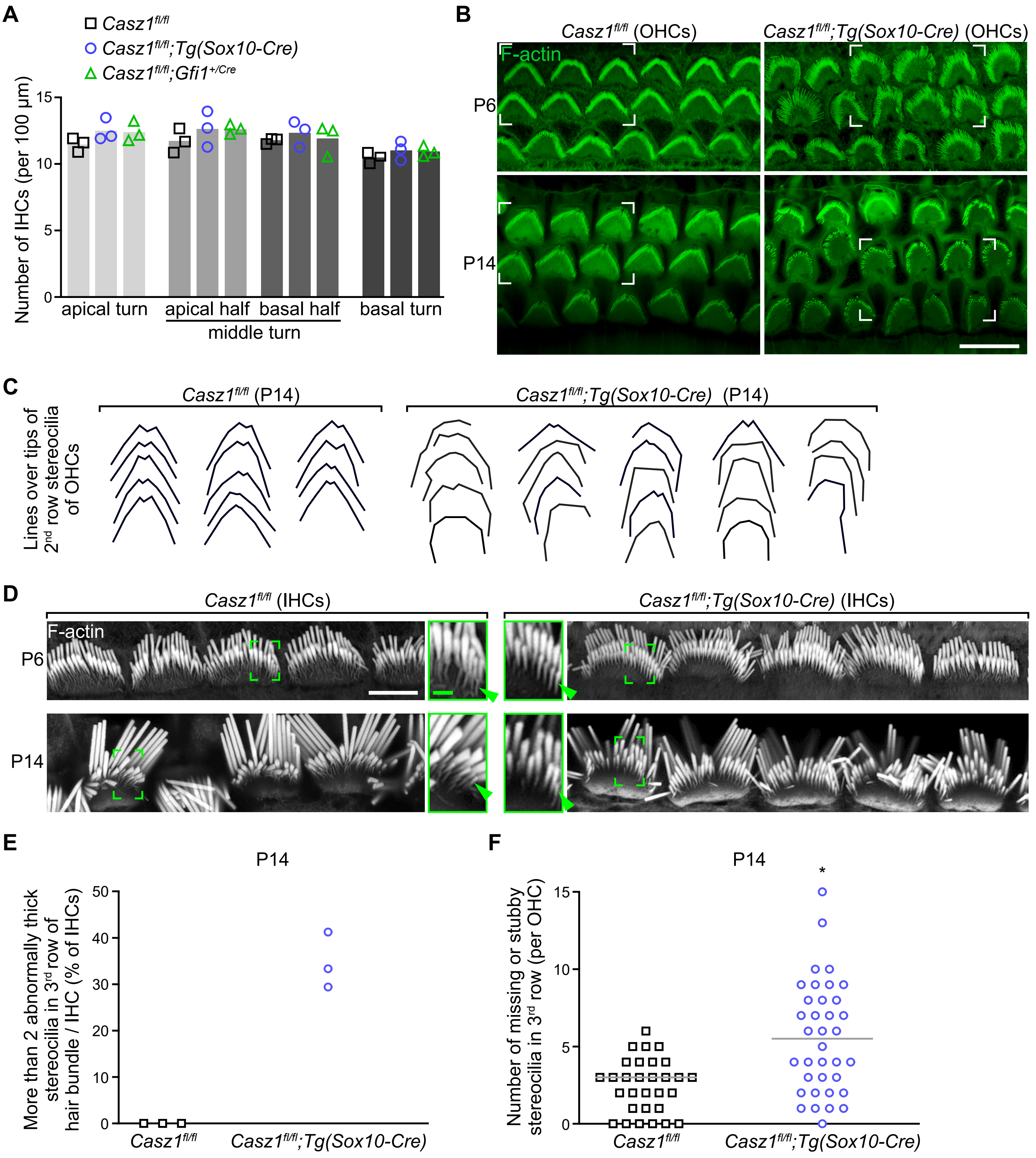

### Figure S4

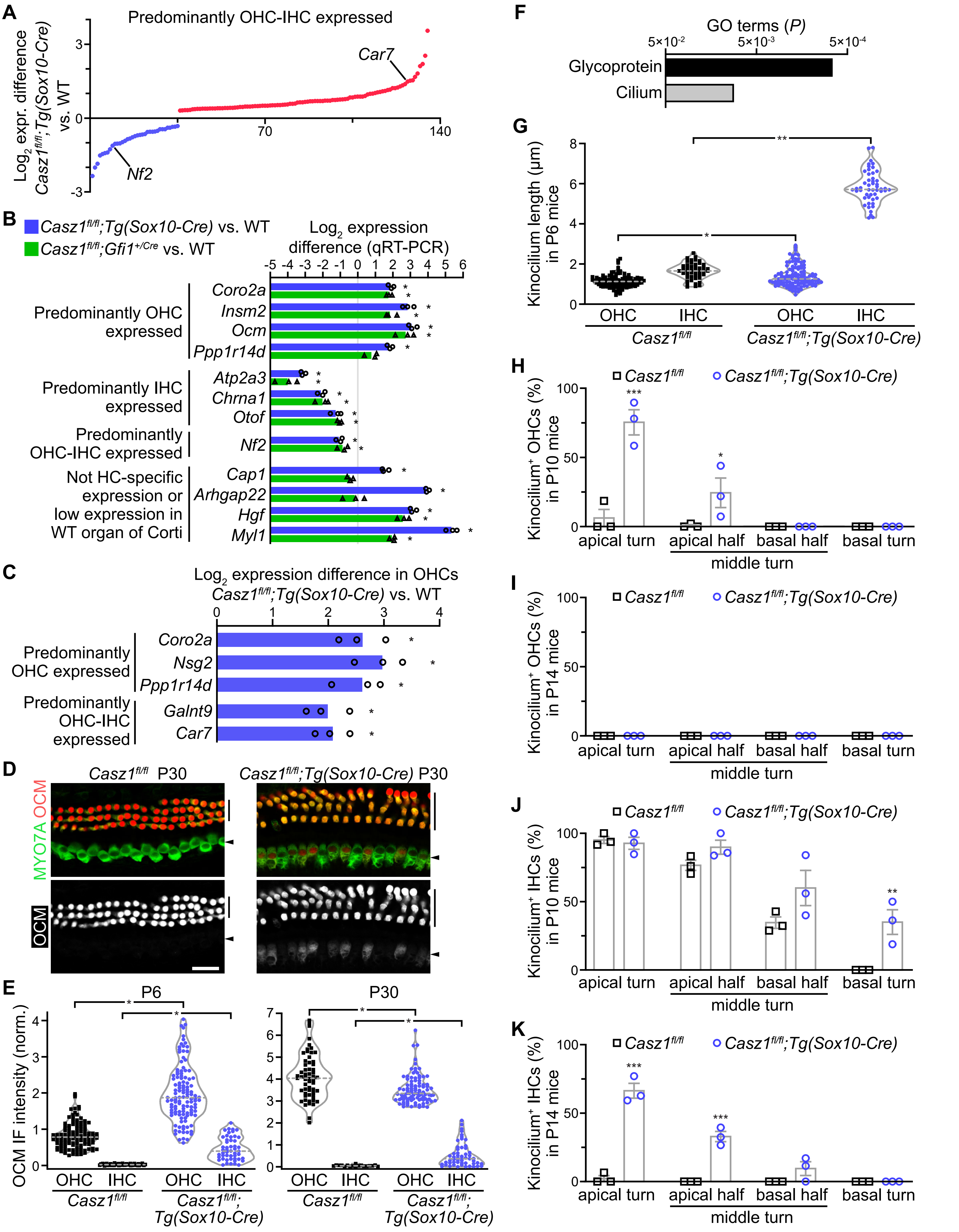

### Figure S5

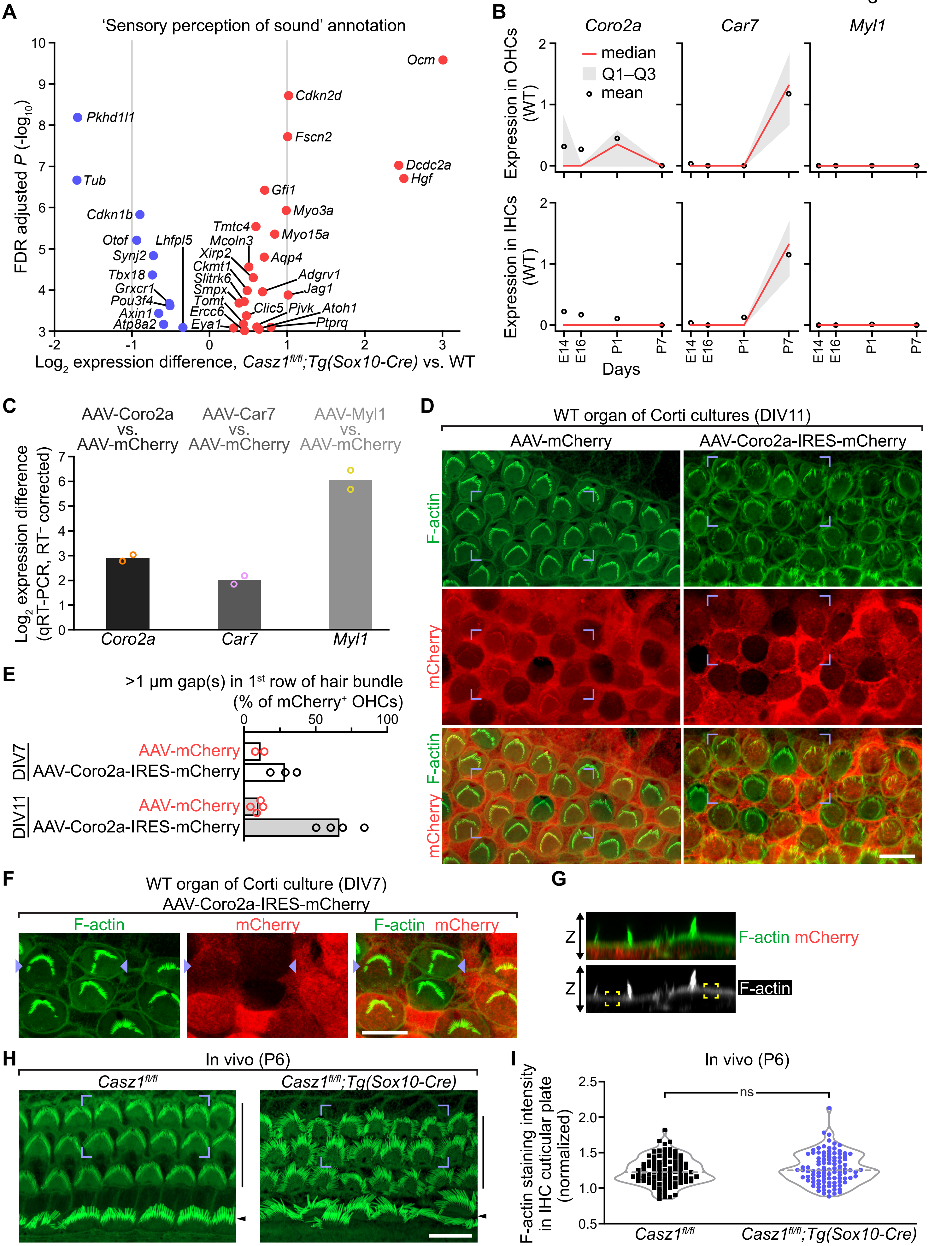

### Figure S6

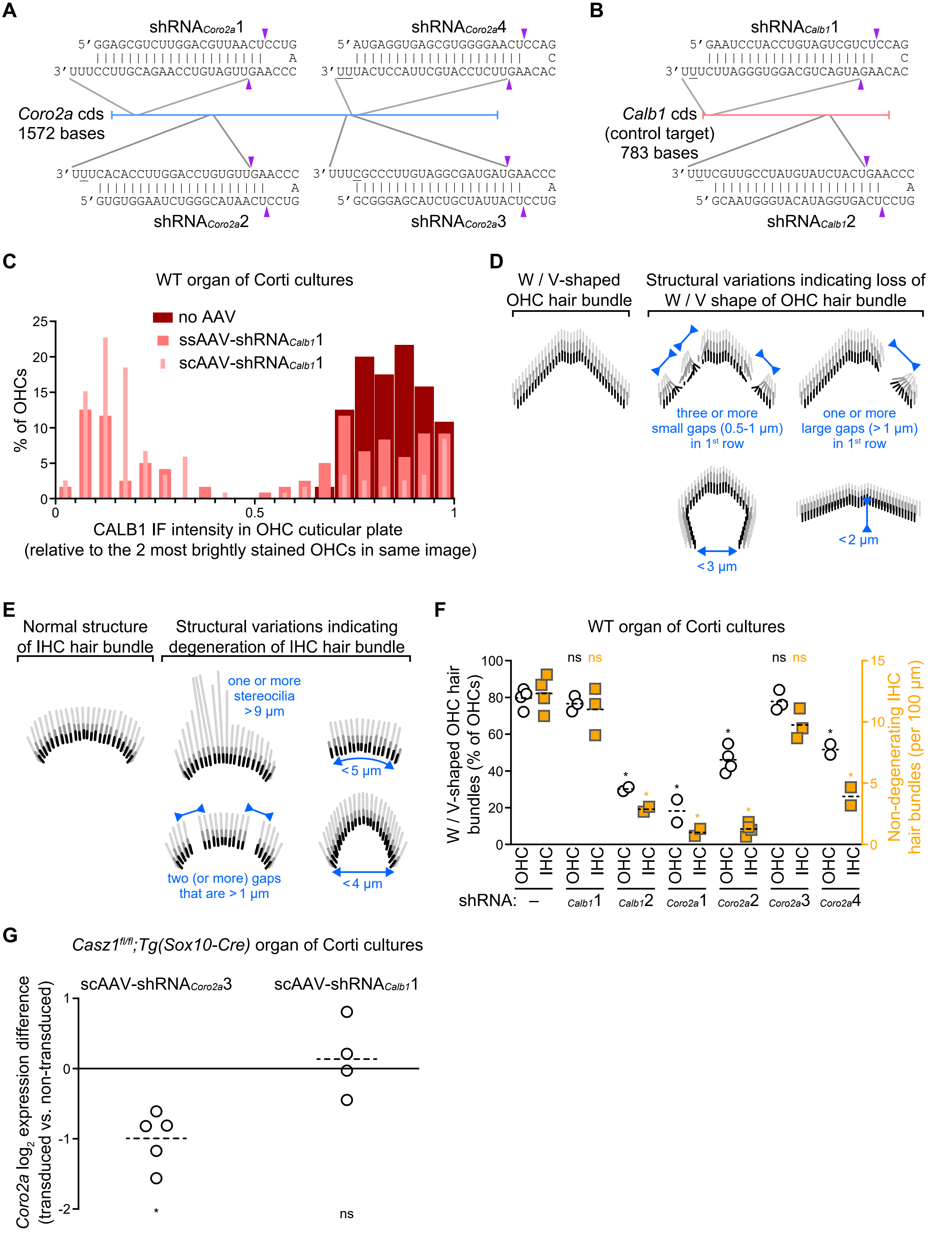
