## Supplementary material for "CASZ1 regulates the rate at which outer hair cells mature and is required for hearing": Table S1

**Table S1. Classification of genes based on identification as ATOH1 targets, functional annotations, and differential expression between HCs and non-HCs in the organ of Corti.**(500 genes are listed.)

| Gene symbol | Gene name | Entrez gene ID | MGI gene ID | Nearest gene neighbor of an ATOH1 binding site identified in PMID:35551236? y/n | Annotated with 'transcription regulator activity' GO term? y/n | Log2 fold expression difference in HCs versus non-HCs in organ of Corti at P1 (PMID: 32404924).*,** | Expressed >8-fold higher in hair cells versus supporting cells at P1, based on PMID: 32404924? y/n | Association between gene defect and hearing loss or HC loss? (Listed for top 20 genes.) |
|---|---|---|---|---|---|---|---|---|
| Atoh1 | atonal bHLH transcription factor 1 | 11921 | MGI:104654 | y | y | 4.95 | y | Prosensory cells do not mature into HCs in KO mice (PMID: 10364557). |
| Bach2 | BTB and CNC homology, basic leucine zipper | 12014 | MGI:894679 | y | y | 6.86 | y | Normal hearing in KO mice (PMID: 36305825). |
| Casz1 | castor zinc finger 1 | 69743 | MGI:1196251 | y | y | 5.64 | y | Hearing loss in Casz1 cKO mice reported after completing experiments for this study (PMID: 39883789). |
| Cbfa2t3 | CBFA2/RUNX1 translocation partner 3 | 12398 | MGI:1338013 | y | y | 5.91 | y | unknown |
| E2f7 | E2F transcription factor 7 | 52679 | MGI:1289147 | y | y | 4.32 | y | unknown |
| Esr2 | estrogen receptor 2 (beta) | 13983 | MGI:109392 | y | y | 5.46 | y | Normal hearing in KO mice (PMID: 18317592). |
| Foxj1 | forkhead box J1 | 15223 | MGI:1347474 | y | y | 4.32 | y | In humans, FOXJ1 deficiency is not associated with hearing loss (PMID: 31630787). *Foxj1* KO/cKO mice have not been characterized for hearing or HC loss. |
| Foxk1 | forkhead box K1 | 17425 | MGI:1347488 | y | y | 4.58 | y | unknown |
| Gfi1 | growth factor independent 1 transcription repressor | 14581 | MGI:103170 | y | y | 7.61 | y | HC loss and hearing loss in KO mice (PMID: 12441305). |
| Insm2 | insulinoma-associated 2 | 56856 | MGI:1930787 | y | y | 3.58 | y | unknown |
| Lmo1 | LIM domain only 1 | 109594 | MGI:102812 | y | y | 3.20 | y | Normal hearing in KO mice (PMID: 36305825). |
| Myt1 | myelin transcription factor 1 | 17932 | MGI:1100535 | y | y | 6.17 | y | unknown |
| Myt1l | myelin transcription factor 1-like | 17933 | MGI:1100511 | y | y | 3.58 | y | Normal hearing in KO mice (PMID: 36305825). |
| Neurod6 | neurogenic differentiation 6 | 11922 | MGI:106593 | y | y | 6.58 | y | unknown prior to this study |
| Nhlh1 | nescient helix loop helix 1 | 18071 | MGI:98481 | y | y | 7.52 | y | Normal hearing in KO mice (PMID: 36305825). |
| Pou4f3 | POU domain, class 4, transcription factor 3 | 18998 | MGI:102523 | y | y | 8.23 | y | HC loss and hearing loss in KO mice (PMID: 8637595). |
| Runx1t1 | RUNX1 translocation partner 1 | 12395 | MGI:104793 | y | y | 3.81 | y | unknown prior to this study |
| Skor1 | SKI family transcriptional corepressor 1 | 207667 | MGI:2443473 | y | y | 6.09 | y | unknown |
| Tcerg1l | transcription elongation regulator 1-like | 70571 | MGI:1917821 | y | y | 6.64 | y | Normal hearing in KO mice (PMID: 36305825). |
| Tfcp2l1 | transcription factor CP2-like 1 | 81879 | MGI:2444691 | y | y | 3.58 | y | unknown |
| Klf10 | Kruppel-like transcription factor 10 | 21847 | MGI:1101353 | y | y | -6.00 | n | |
| Hivep2 | human immunodeficiency virus type I enhancer | 15273 | MGI:1338076 | y | y | -4.58 | n | *Zero expression levels were converted to -5.32193 in log2 scale (i.e., 0.025 in linear scale) for ratio calculations |
| Zfp507 | zinc finger protein 507 | 668501 | MGI:1916378 | y | y | -4.58 | n | **Non-HCs are defined as the following cell types in the PMID: 32404924 dataset: LatSCa, LatSCb, MedSC, NSC (i), NSC (ii), and SC. |
| Srebf2 | sterol regulatory element bin | 20788 | MGI:107585 | y | y | -4.32 | n | |
| Carm1 | coactivator-associated argin | 59035 | MGI:1913208 | y | y | -4.32 | n | |
| Pou6f1 | POU domain, class 6, transc | 19009 | MGI:102935 | y | y | -4.00 | n | |
| Foxq1 | forkhead box Q1 | 15220 | MGI:1298228 | y | y | -3.58 | n | |
| Maf | MAF bZIP transcription facto | 17132 | MGI:96909 | y | y | -3.58 | n | |
| Mitf | melanogenesis associated t | 17342 | MGI:104554 | y | y | -3.58 | n | |
| Sp5 | trans-acting transcription fac | 64406 | MGI:1927715 | y | y | -3.58 | n | |
| Zfp963 | zinc finger protein 963 | 620419 | MGI:4867078 | y | y | -3.58 | n | |
| Fbxl19 | F-box and leucine-rich repea | 233902 | MGI:3039600 | y | y | -3.58 | n | |
| Klf12 | Kruppel-like transcription fac | 16597 | MGI:1333796 | y | y | -3.17 | n | |
| Zfp3 | zinc finger protein 3 | 193043 | MGI:99177 | y | y | -3.00 | n | |
| Tle2 | transducin-like enhancer of s | 21886 | MGI:104635 | y | y | -3.00 | n | |
| Sfmbt2 | aldolase 1 A, retrogene 1 | 353282 | MGI:2447794 | y | y | -3.00 | n | |
| Plagl1 | pleiomorphic adenoma gene | 22634 | MGI:1100874 | y | y | -2.81 | n | |
| Irx3 | Iroquois related homeobox 3 | 16373 | MGI:1197522 | y | y | -2.70 | n | |

| | | | | | | | |
|---|---|---|---|---|---|---|---|
| Nr4a3 | nuclear receptor subfamily 4 | 18124 | MGI:1352457 | y | y | -2.58 | n |
| Atf5 | activating transcription facto | 107503 | MGI:2141857 | y | y | -2.32 | n |
| Dach2 | dachshund family transcripti | 93837 | MGI:1890446 | y | y | -2.32 | n |
| Egr3 | early growth response 3 | 13655 | MGI:1306780 | y | y | -2.32 | n |
| Paqr8 | progestin and adipoQ recept | 74229 | MGI:1921479 | y | y | -2.32 | n |
| Hivep3 | human immunodeficiency vi | 16656 | MGI:106589 | y | y | -2.00 | n |
| Hsfy2 | heat shock transcription fact | 71066 | MGI:1918316 | y | y | -2.00 | n |
| Mef2c | myocyte enhancer factor 2C | 17260 | MGI:99458 | y | y | -2.00 | n |
| Snai3 | snail family zinc finger 3 | 30927 | MGI:1353563 | y | y | -2.00 | n |
| Tfap4 | transcription factor AP4 | 83383 | MGI:103239 | y | y | -2.00 | n |
| Rcor2 | REST corepressor 2 | 104383 | MGI:1859854 | y | y | -2.00 | n |
| Cenpj | glucoside xylosyltransferase | 219103 | MGI:2684927 | y | y | -2.00 | n |
| Mxd4 | Max dimerization protein 4 | 17122 | MGI:104991 | y | y | -1.58 | n |
| Zfp395 | zinc finger protein 395 | 380912 | MGI:2682318 | y | y | -1.58 | n |
| Dnajb1 | DnaJ heat shock protein fam | 81489 | MGI:1931874 | y | y | -1.52 | n |
| Zfp35 | zinc finger protein 35 | 22694 | MGI:99179 | y | y | -1.42 | n |
| Zfpm1 | zinc finger protein, multitype | 22761 | MGI:1095400 | y | y | -1.32 | n |
| Tgif2 | TGFB-induced factor homeo | 228839 | MGI:1915299 | y | y | -1.00 | n |
| Nrg1 | lipase, hepatic | 211323 | MGI:96083 | y | y | -1.00 | n |
| Tox3 | TOX high mobility group box | 244579 | MGI:3039593 | y | y | -0.86 | n |
| Cic | capicua transcriptional repre | 71722 | MGI:1918972 | y | y | -0.81 | n |
| Zfp143 | zinc finger protein 143 | 20841 | MGI:1277969 | y | y | -0.81 | n |
| Jdp2 | Jun dimerization protein 2 | 81703 | MGI:1932093 | y | y | -0.74 | n |
| Smad3 | SMAD family member 3 | 17127 | MGI:1201674 | y | y | -0.68 | n |
| Mycn | v-myc avian myelocytomato | 18109 | MGI:97357 | y | y | -0.65 | n |
| Rere | arginine glutamic acid dipep | 68703 | MGI:2683486 | y | y | -0.62 | n |
| Mxd1 | MAX dimerization protein 1 | 17119 | MGI:96908 | y | y | -0.58 | n |
| Tshz3 | teashirt zinc finger family me | 243931 | MGI:2442819 | y | y | -0.58 | n |
| Bmal1 | basic helix-loop-helix ARNT | 11865 | MGI:1096381 | y | y | -0.58 | n |
| Klf13 | Kruppel-like transcription fac | 50794 | MGI:1354948 | y | y | -0.52 | n |
| Ddx54 | DEAD box helicase 54 | 71990 | MGI:1919240 | y | y | -0.46 | n |
| Nfatc2 | nuclear factor of activated T | 18019 | MGI:102463 | y | y | -0.42 | n |
| Tead1 | TEA domain family member | 21676 | MGI:101876 | y | y | -0.36 | n |
| Fos | FBJ osteosarcoma oncogen | 14281 | MGI:95574 | y | y | -0.34 | n |
| Sox11 | SRY (sex determining regior | 20666 | MGI:98359 | y | y | -0.33 | n |
| Klf11 | Kruppel-like transcription fac | 194655 | MGI:2653368 | y | y | -0.32 | n |
| Nr6a1 | nuclear receptor subfamily 6 | 14536 | MGI:1352459 | y | y | -0.32 | n |
| Ncor2 | nuclear receptor co-represso | 20602 | MGI:1337080 | y | y | -0.26 | n |
| Rbpj | recombination signal binding | 19664 | MGI:96522 | y | y | -0.26 | n |
| Nfib | nuclear factor I/B | 18028 | MGI:103188 | y | y | -0.19 | n |
| Nr2f1 | nuclear receptor subfamily 2 | 13865 | MGI:1352451 | y | y | -0.17 | n |
| Tle3 | transducin-like enhancer of s | 21887 | MGI:104634 | y | y | -0.16 | n |
| Dcaf6 | DDB1 and CUL4 associated | 74106 | MGI:1921356 | y | y | -0.12 | n |
| C1d | C1D nuclear receptor co-rep | 57316 | MGI:1927354 | y | y | -0.10 | n |
| Irf2bp2 | interferon regulatory factor 2 | 270110 | MGI:2443921 | y | y | -0.10 | n |
| Klf7 | Kruppel-like transcription fac | 93691 | MGI:1935151 | y | y | -0.10 | n |
| Ebf1 | early B cell factor 1 | 13591 | MGI:95275 | y | y | -0.04 | n |
| Jun | jun proto-oncogene | 16476 | MGI:96646 | y | y | -0.02 | n |
| Id2 | inhibitor of DNA binding 2 | 15902 | MGI:96397 | y | y | -0.01 | n |
| Dbp | D site albumin promoter bind | 13170 | MGI:94866 | y | y | 0.00 | n |
| Patz1 | POZ (BTB) and AT hook con | 56218 | MGI:1891832 | y | y | 0.00 | n |
| Tshz1 | teashirt zinc finger family me | 110796 | MGI:1346031 | y | y | 0.00 | n |
| Zfp568 | zinc finger protein 568 | 243905 | MGI:2142347 | y | y | 0.00 | n |
| Zhx3 | zinc fingers and homeoboxe | 320799 | MGI:2444772 | y | y | 0.00 | n |
| Abhd2 | abhydrolase domain contain | 54608 | MGI:1914344 | y | y | 0.00 | n |
| Cbfb | core binding factor beta | 12400 | MGI:99851 | y | y | 0.00 | n |
| Trim32 | tripartite motif-containing 32 | 69807 | MGI:1917057 | y | y | 0.00 | n |
| Trim8 | tripartite motif-containing 8 | 93679 | MGI:1933302 | y | y | 0.00 | n |
| Id3 | inhibitor of DNA binding 3 | 15903 | MGI:96398 | y | y | 0.02 | n |
| Mrtfb | 5'-nucleotidase domain cont | 239719 | MGI:3050795 | y | y | 0.03 | n |
| Tcerg1 | transcription elongation regu | 56070 | MGI:1926421 | y | y | 0.04 | n |
| Naca | nascent polypeptide-associa | 17938 | MGI:106095 | y | y | 0.04 | n |
| Hbp1 | high mobility group box trans | 73389 | MGI:894659 | y | y | 0.07 | n |
| Litaf | LPS-induced TN factor | 56722 | MGI:1929512 | y | y | 0.08 | n |
| Nr2c2 | nuclear receptor subfamily 2 | 22026 | MGI:1352466 | y | y | 0.09 | n |
| Kdm3b | KDM3B lysine (K)-specific de | 277250 | MGI:1923356 | y | y | 0.09 | n |

| | | | | | | | |
|---|---|---|---|---|---|---|---|
| Thrb | thyroid hormone receptor be | 21834 | MGI:98743 | y | y | 0.09 | n |
| Btg2 | BTG anti-proliferation factor | 12227 | MGI:108384 | y | y | 0.10 | n |
| Nipbl | NIPBL cohesin loading facto | 71175 | MGI:1913976 | y | y | 0.10 | n |
| Sertad2 | SERTA domain containing 2 | 58172 | MGI:1931026 | y | y | 0.11 | n |
| Hsbp1 | heat shock factor binding pro | 68196 | MGI:1915446 | y | y | 0.12 | n |
| Aff3 | AF4/FMR2 family, member 3 | 16764 | MGI:106927 | y | y | 0.13 | n |
| Skil | SKI-like | 20482 | MGI:106203 | y | y | 0.13 | n |
| Smad7 | SMAD family member 7 | 17131 | MGI:1100518 | y | y | 0.14 | n |
| Zmynd8 | zinc finger, MYND-type cont | 228880 | MGI:1918025 | y | y | 0.14 | n |
| Trim27 | tripartite motif-containing 27 | 19720 | MGI:97904 | y | y | 0.16 | n |
| Sp1 | trans-acting transcription fac | 20683 | MGI:98372 | y | y | 0.18 | n |
| Isl1 | ISL1 transcription factor, LIM | 16392 | MGI:101791 | y | y | 0.19 | n |
| Jund | jun D proto-oncogene | 16478 | MGI:96648 | y | y | 0.19 | n |
| Atf4 | activating transcription facto | 11911 | MGI:88096 | y | y | 0.21 | n |
| Camta1 | calmodulin binding transcrip | 100072 | MGI:2140230 | y | y | 0.21 | n |
| Kdm4c | lysine (K)-specific demethyla | 76804 | MGI:1924054 | y | y | 0.21 | n |
| Sfr1 | SWI5 dependent recombina | 67788 | MGI:1915038 | y | y | 0.22 | n |
| Pknox1 | Pbx/knotted 1 homeobox | 18771 | MGI:1201409 | y | y | 0.26 | n |
| Nfkbiz | nuclear factor of kappa light | 80859 | MGI:1931595 | y | y | 0.26 | n |
| Nfia | nuclear factor I/A | 18027 | MGI:108056 | y | y | 0.27 | n |
| Sall3 | spalt like transcription factor | 20689 | MGI:109295 | y | y | 0.28 | n |
| Trp53bp1 | transformation related protei | 27223 | MGI:1351320 | y | y | 0.28 | n |
| Hivep1 | human immunodeficiency vi | 110521 | MGI:96100 | y | y | 0.30 | n |
| Foxp4 | forkhead box P4 | 74123 | MGI:1921373 | y | y | 0.32 | n |
| Sox4 | SRY (sex determining regior | 20677 | MGI:98366 | y | y | 0.32 | n |
| Lpin1 | lipin 1 | 14245 | MGI:1891340 | y | y | 0.32 | n |
| Hcfc2 | host cell factor C2 | 67933 | MGI:1915183 | y | y | 0.32 | n |
| Rybp | RING1 and YY1 binding pro | 56353 | MGI:1929059 | y | y | 0.32 | n |
| Klf9 | Kruppel-like transcription fac | 16601 | MGI:1333856 | y | y | 0.33 | n |
| Tob1 | transducer of ErbB-2.1 | 22057 | MGI:1349721 | y | y | 0.36 | n |
| Calcoco1 | calcium binding and coiled c | 67488 | MGI:1914738 | y | y | 0.37 | n |
| Kdm3a | lysine (K)-specific demethyla | 104263 | MGI:98847 | y | y | 0.39 | n |
| Peg3 | paternally expressed 3 | 18616 | MGI:104748 | y | y | 0.40 | n |
| Etv6 | ets variant 6 | 14011 | MGI:109336 | y | y | 0.40 | n |
| Fhl2 | four and a half LIM domains | 14200 | MGI:1338762 | y | y | 0.42 | n |
| Ubp1 | upstream binding protein 1 | 22221 | MGI:104889 | y | y | 0.44 | n |
| Tef | thyrotroph embryonic factor | 21685 | MGI:98663 | y | y | 0.44 | n |
| Tdrd3 | tudor domain containing 3 | 219249 | MGI:2444023 | y | y | 0.44 | n |
| Tsg101 | tumor susceptibility gene 10 | 22088 | MGI:106581 | y | y | 0.45 | n |
| Sall1 | spalt like transcription factor | 58198 | MGI:1889585 | y | y | 0.45 | n |
| Stat3 | signal transducer and activa | 20848 | MGI:103038 | y | y | 0.46 | n |
| Eid1 | EP300 interacting inhibitor o | 58521 | MGI:1889651 | y | y | 0.46 | n |
| Gtf2i | general transcription factor I | 14886 | MGI:1202722 | y | y | 0.50 | n |
| Wwox | WW domain-containing oxid | 80707 | MGI:1931237 | y | y | 0.51 | n |
| Psip1 | PC4 and SFRS1 interacting | 101739 | MGI:2142116 | y | y | 0.53 | n |
| Zfp644 | zinc finger protein 644 | 52397 | MGI:1277212 | y | y | 0.55 | n |
| Jmjd1c | jumonji domain containing 1 | 108829 | MGI:1918614 | y | y | 0.56 | n |
| Tfb1m | transcription factor B1, mitoc | 224481 | MGI:2146851 | y | y | 0.58 | n |
| Usf1 | upstream transcription factor | 22278 | MGI:99542 | y | y | 0.58 | n |
| Mnt | max binding protein | 17428 | MGI:109150 | y | y | 0.62 | n |
| Pbx1 | pre B cell leukemia homeob | 18514 | MGI:97495 | y | y | 0.63 | n |
| Atf7ip | activating transcription facto | 54343 | MGI:1858965 | y | y | 0.64 | n |
| Zmiz1 | phosphatidylinositol-3,4,5-tri | 328365 | MGI:3040693 | y | y | 0.64 | n |
| Eny2 | ENY2 transcription and expo | 223527 | MGI:1919286 | y | y | 0.66 | n |
| Actn1 | actinin, alpha 1 | 109711 | MGI:2137706 | y | y | 0.66 | n |
| Foxp1 | forkhead box P1 | 108655 | MGI:1914004 | y | y | 0.66 | n |
| Ssbp3 | single-stranded DNA binding | 72475 | MGI:1919725 | y | y | 0.67 | n |
| Nfatc4 | nuclear factor of activated T | 73181 | MGI:1920431 | y | y | 0.68 | n |
| Elf2 | E74-like factor 2 | 69257 | MGI:1916507 | y | y | 0.68 | n |
| Gtf2ird1 | general transcription factor I | 57080 | MGI:1861942 | y | y | 0.70 | n |
| Tbl1xr1 | transducin (beta)-like 1X-link | 81004 | MGI:2441730 | y | y | 0.73 | n |
| Xbp1 | X-box binding protein 1 | 22433 | MGI:98970 | y | y | 0.77 | n |
| Tfdp2 | transcription factor Dp 2 | 211586 | MGI:107167 | y | y | 0.78 | n |
| Zfp1 | zinc finger protein 1 | 22640 | MGI:99154 | y | y | 0.78 | n |
| Fiz1 | Flt3 interacting zinc finger pr | 23877 | MGI:1344336 | y | y | 0.78 | n |
| Relb | avian reticuloendotheliosis v | 19698 | MGI:103289 | y | y | 0.78 | n |

| | | | | | | | |
|---|---|---|---|---|---|---|---|
| Sin3a | transcriptional regulator, SIN | 20466 | MGI:107157 | y | y | 0.82 | n |
| Dmap1 | DNA methyltransferase 1-as | 66233 | MGI:1913483 | y | y | 0.85 | n |
| Psmd9 | proteasome (prosome, macr | 67151 | MGI:1914401 | y | y | 0.86 | n |
| Cebpg | CCAAT/enhancer binding pr | 12611 | MGI:104982 | y | y | 0.87 | n |
| Mtf2 | metal response element bin | 17765 | MGI:105050 | y | y | 0.88 | n |
| Zbtb5 | zinc finger and BTB domain | 230119 | MGI:1924601 | y | y | 0.92 | n |
| Hmga1 | high mobility group AT-hook | 15361 | MGI:96160 | y | y | 0.93 | n |
| Elk3 | ELK3, member of ETS onco | 13713 | MGI:101762 | y | y | 1.00 | n |
| Zfp319 | zinc finger protein 319 | 79233 | MGI:1890618 | y | y | 1.00 | n |
| Zfp696 | zinc finger protein 696 | 0004313 | MGI:2442738 | y | y | 1.00 | n |
| Camta2 | calmodulin binding transcrip | 216874 | MGI:2135957 | y | y | 1.00 | n |
| Nrip1 | nuclear receptor interacting | 268903 | MGI:1315213 | y | y | 1.03 | n |
| Zfp91 | zinc finger protein 91 | 109910 | MGI:104854 | y | y | 1.04 | n |
| Hif3a | hypoxia inducible factor 3, a | 53417 | MGI:1859778 | y | y | 1.08 | n |
| Tcf4 | transcription factor 4 | 21413 | MGI:98506 | y | y | 1.10 | n |
| Bcl11a | BCL11 transcription factor A | 14025 | MGI:106190 | y | y | 1.13 | n |
| Satb1 | special AT-rich sequence bi | 20230 | MGI:105084 | y | y | 1.14 | n |
| Mrtfa | myocardin related transcripti | 223701 | MGI:2384495 | y | y | 1.17 | n |
| Zfp865 | zinc finger protein 865 | 319748 | MGI:2442656 | y | y | 1.17 | n |
| Per2 | period circadian clock 2 | 18627 | MGI:1195265 | y | y | 1.17 | n |
| Zfp710 | zinc finger protein 710 | 209225 | MGI:1921747 | y | y | 1.21 | n |
| Dyrk1b | dual-specificity tyrosine pho | 13549 | MGI:1330302 | y | y | 1.22 | n |
| Nab1 | Ngfi-A binding protein 1 | 17936 | MGI:107564 | y | y | 1.25 | n |
| Zbtb7a | zinc finger and BTB domain | 16969 | MGI:1335091 | y | y | 1.26 | n |
| Pias4 | protein inhibitor of activated | 59004 | MGI:2136940 | y | y | 1.28 | n |
| Sox12 | SRY (sex determining regior | 20667 | MGI:98360 | y | y | 1.30 | n |
| Deaf1 | DEAF1, transcription factor | 54006 | MGI:1858496 | y | y | 1.30 | n |
| Cux2 | cut-like homeobox 2 | 13048 | MGI:107321 | y | y | 1.32 | n |
| Mef2d | myocyte enhancer factor 2D | 17261 | MGI:99533 | y | y | 1.32 | n |
| Zfp438 | zinc finger protein 438 | 240186 | MGI:2444919 | y | y | 1.32 | n |
| Zfp51 | zinc finger protein 51 | 22709 | MGI:99198 | y | y | 1.32 | n |
| Nfkb1 | nuclear factor of kappa light | 18033 | MGI:97312 | y | y | 1.32 | n |
| Wwc1 | WW, C2 and coiled-coil dom | 211652 | MGI:2388637 | y | y | 1.33 | n |
| Crebl2 | cAMP responsive element b | 232430 | MGI:1889385 | y | y | 1.42 | n |
| Nfatc1 | nuclear factor of activated T | 18018 | MGI:102469 | y | y | 1.42 | n |
| Lrrfip1 | leucine rich repeat (in FLII) i | 16978 | MGI:1342770 | y | y | 1.44 | n |
| Cdyl2 | chromodomain protein, Y ch | 75796 | MGI:1923046 | y | y | 1.49 | n |
| Zfp184 | zinc finger protein 184 (Krup | 193452 | MGI:1922244 | y | y | 1.55 | n |
| Hdac4 | histone deacetylase 4 | 208727 | MGI:3036234 | y | y | 1.58 | n |
| Trim62 | tripartite motif-containing 62 | 67525 | MGI:1914775 | y | y | 1.63 | n |
| Zfp667 | zinc finger protein 667 | 384763 | MGI:2442757 | y | y | 1.70 | n |
| Cbfa2t2 | CBFA2/RUNX1 translocatior | 12396 | MGI:1333833 | y | y | 1.79 | n |
| Emx2 | empty spiracles homeobox 2 | 13797 | MGI:95388 | y | y | 1.81 | n |
| Max | Max protein | 17187 | MGI:96921 | y | y | 1.81 | n |
| Zfp647 | zinc finger protein 647 | 239546 | MGI:3052806 | y | y | 1.81 | n |
| Pou2f1 | POU domain, class 2, transc | 18986 | MGI:101898 | y | y | 1.85 | n |
| Rxra | retinoid X receptor alpha | 20181 | MGI:98214 | y | y | 1.93 | n |
| Rara | retinoic acid receptor, alpha | 19401 | MGI:97856 | y | y | 2.00 | n |
| Cited4 | Cbp/p300-interacting transa | 56222 | MGI:1861694 | y | y | 2.00 | n |
| Satb2 | special AT-rich sequence bi | 212712 | MGI:2679336 | y | y | 2.00 | n |
| Hes6 | hairy and enhancer of split 6 | 55927 | MGI:1859852 | y | y | 2.12 | n |
| Zkscan16 | zinc finger with KRAB and S | 0004158 | MGI:3510405 | y | y | 2.58 | n |
| Dach1 | dachshund family transcripti | 13134 | MGI:1277991 | y | y | 2.75 | n |
| Zfp119b | zinc finger protein 119b | 240120 | MGI:2385323 | y | y | 2.91 | n |
| Zfp169 | zinc finger protein 169 | 67911 | MGI:1915161 | y | y | 2.91 | n |
| Mamstr | MEF2 activating motif and S | 74490 | MGI:1921740 | y | y | 3.00 | n |
| Esrrg | estrogen-related receptor ga | 26381 | MGI:1347056 | y | y | 3.00 | n |
| Mxd3 | Max dimerization protein 3 | 17121 | MGI:104987 | y | y | 3.00 | n |
| Atoh8 | atonal bHLH transcription fa | 71093 | MGI:1918343 | y | y | not in dataset | n |
| Barx2 | BarH-like homeobox 2 | 12023 | MGI:109617 | y | y | not in dataset | n |
| Bcl3 | B cell leukemia/lymphoma 3 | 12051 | MGI:88140 | y | y | not in dataset | n |
| Bhlha9 | basic helix-loop-helix family, | 320522 | MGI:2444198 | y | y | not in dataset | n |
| E2f8 | E2F transcription factor 8 | 108961 | MGI:1922038 | y | y | not in dataset | n |
| Ebf2 | early B cell factor 2 | 13592 | MGI:894332 | y | y | not in dataset | n |
| Foxo6 | forkhead box O6 | 329934 | MGI:2676586 | y | y | not in dataset | n |
| Hes2 | hes family bHLH transcriptio | 15206 | MGI:1098624 | y | y | not in dataset | n |

| | | | | | | | |
|---|---|---|---|---|---|---|---|
| Ikzf1 | IKAROS family zinc finger 1 | 22778 | MGI:1342540 | y | y | not in dataset | n |
| Lhx2 | LIM homeobox protein 2 | 16870 | MGI:96785 | y | y | not in dataset | n |
| Lhx8 | LIM homeobox protein 8 | 16875 | MGI:1096343 | y | y | not in dataset | n |
| Lmx1b | LIM homeobox transcription | 16917 | MGI:1100513 | y | y | not in dataset | n |
| Mafa | MAF bZIP transcription facto | 378435 | MGI:2673307 | y | y | not in dataset | n |
| Mecom | MDS1 and EVI1 complex loc | 14013 | MGI:95457 | y | y | not in dataset | n |
| Mef2b | myocyte enhancer factor 2B | 17259 | MGI:104526 | y | y | not in dataset | n |
| Meis2 | Meis homeobox 2 | 17536 | MGI:108564 | y | y | not in dataset | n |
| Myb | myeloblastosis oncogene | 17863 | MGI:97249 | y | y | not in dataset | n |
| Mycl | v-myc avian myelocytomato: | 16918 | MGI:96799 | y | y | not in dataset | n |
| Myf6 | myogenic factor 6 | 17878 | MGI:97253 | y | y | not in dataset | n |
| Myrf | myelin regulatory factor | 225908 | MGI:2684944 | y | y | not in dataset | n |
| Neurod1 | neurogenic differentiation 1 | 18012 | MGI:1339708 | y | y | not in dataset | n |
| Neurod2 | neurogenic differentiation 2 | 18013 | MGI:107755 | y | y | not in dataset | n |
| Notch4 | notch 4 | 18132 | MGI:107471 | y | y | not in dataset | n |
| Nr1h4 | nuclear receptor subfamily 1 | 20186 | MGI:1352464 | y | y | not in dataset | n |
| Otx2 | orthodenticle homeobox 2 | 18424 | MGI:97451 | y | y | not in dataset | n |
| Pitx1 | paired-like homeodomain tra | 18740 | MGI:107374 | y | y | not in dataset | n |
| Pou3f1 | POU domain, class 3, transc | 18991 | MGI:101896 | y | y | not in dataset | n |
| Runx2 | runt related transcription fac | 12393 | MGI:99829 | y | y | not in dataset | n |
| Shox2 | SHOX homeobox 2 | 20429 | MGI:1201673 | y | y | not in dataset | n |
| Skor2 | SKI family transcriptional co | 664805 | MGI:3645984 | y | y | not in dataset | n |
| St18 | suppression of tumorigenicit | 240690 | MGI:2446700 | y | y | not in dataset | n |
| Tal2 | T cell acute lymphocytic leuk | 21350 | MGI:99540 | y | y | not in dataset | n |
| Tcp10a | t-complex protein 10a | 21461 | MGI:98541 | y | y | not in dataset | n |
| Tfap2c | transcription factor AP-2, ga | 21420 | MGI:106032 | y | y | not in dataset | n |
| Uncx | UNC homeobox | 22255 | MGI:108013 | y | y | not in dataset | n |
| Usf3 | upstream transcription facto | 207806 | MGI:2685454 | y | y | not in dataset | n |
| Vsx2 | visual system homeobox 2 | 12677 | MGI:88401 | y | y | not in dataset | n |
| Zbtb18 | zinc finger and BTB domain | 30928 | MGI:1353609 | y | y | not in dataset | n |
| Zbtb7c | zinc finger and BTB domain | 207259 | MGI:2443302 | y | y | not in dataset | n |
| Zfp41 | zinc finger protein 41 | 22701 | MGI:99186 | y | y | not in dataset | n |
| Zfp78 | zinc finger protein 78 | 330463 | MGI:107783 | y | y | not in dataset | n |
| Pou2af1 | POU domain, class 2, assoc | 18985 | MGI:105086 | y | y | not in dataset | n |
| Kat6b | K(lysine) acetyltransferase 6 | 54169 | MGI:1858746 | y | y | not in dataset | n |
| Uri1 | URI1, prefoldin-like chapero | 19777 | MGI:1342294 | y | y | not in dataset | n |
| Prkn | parkin RBR E3 ubiquitin prot | 50873 | MGI:1355296 | y | y | not in dataset | n |
| Trim14 | tripartite motif-containing 14 | 74735 | MGI:1921985 | y | y | not in dataset | n |
| Dcc | dihydrofolate reductase | 13176 | MGI:94869 | y | y | not in dataset | n |
| Pkm | polycystin 1, transient recep | 18746 | MGI:97591 | y | y | not in dataset | n |
| Neurl1a | neuralized E3 ubiquitin prote | 18011 | MGI:1334263 | y | n | 3.09 | y |
| Zmat4 | zinc finger, matrin type 4 | 320158 | MGI:2443497 | y | n | 3.14 | y |
| Iqcg | IQ motif containing G | 69707 | MGI:1916957 | y | n | 3.14 | y |
| Ncald | neurocalcin delta | 52589 | MGI:1196326 | y | n | 3.17 | y |
| Sh2d4b | SH2 domain containing 4B | 328381 | MGI:1925182 | y | n | 3.17 | y |
| Clstn3 | calsyntenin 3 | 232370 | MGI:2178323 | y | n | 3.17 | y |
| Tmcc2 | transmembrane and coiled-c | 68875 | MGI:1916125 | y | n | 3.17 | y |
| Rbm24 | RNA binding motif protein 24 | 666794 | MGI:3610364 | y | n | 3.21 | y |
| Necab2 | N-terminal EF-hand calcium | 117148 | MGI:2152211 | y | n | 3.25 | y |
| Kcnn2 | potassium intermediate/sma | 140492 | MGI:2153182 | y | n | 3.25 | y |
| Srxn1 | sulfiredoxin 1 homolog (S. c | 76650 | MGI:104971 | y | n | 3.32 | y |
| Rab6b | RAB6B, member RAS oncog | 270192 | MGI:107283 | y | n | 3.32 | y |
| Septin3 | septin 3 | 24050 | MGI:1345148 | y | n | 3.32 | y |
| Fcho1 | FCH domain only 1 | 74015 | MGI:1921265 | y | n | 3.32 | y |
| Dgkh | diacylglycerol kinase, eta | 380921 | MGI:2444188 | y | n | 3.32 | y |
| Lrp8 | low density lipoprotein recep | 16975 | MGI:1340044 | y | n | 3.32 | y |
| Rims2 | regulating synaptic membrar | 116838 | MGI:2152972 | y | n | 3.39 | y |
| Ranbp17 | RAN binding protein 17 | 66011 | MGI:1929706 | y | n | 3.39 | y |
| Cnnm2 | cyclin M2 | 94219 | MGI:2151054 | y | n | 3.39 | y |
| Rbfox2 | RNA binding protein, fox-1 h | 93686 | MGI:1933973 | y | n | 3.41 | y |
| Palld | palladin, cytoskeletal associ | 72333 | MGI:1919583 | y | n | 3.42 | y |
| Rimbp2 | RIMS binding protein 2 | 231760 | MGI:2443235 | y | n | 3.46 | y |
| Gm2011 | predicted gene 2011 | 0003902 | MGI:3780180 | y | n | 3.46 | y |
| Acbd7 | acyl-Coenzyme A binding dc | 78245 | MGI:1925495 | y | n | 3.47 | y |
| Nfasc | neurofascin | 269116 | MGI:104753 | y | n | 3.52 | y |
| Espn | espin | 56226 | MGI:1861630 | y | n | 3.54 | y |

| | | | | | | | |
|---|---|---|---|---|---|---|---|
| Tmem229a | transmembrane protein 229A | 319832 | MGI:2442812 | y | n | 3.54 | y |
| Sema5b | sema domain, seven thromb | 20357 | MGI:107555 | y | n | 3.58 | y |
| Limk1 | LIM domain kinase 1 | 16885 | MGI:104572 | y | n | 3.58 | y |
| Ksr1 | kinase suppressor of ras 1 | 16706 | MGI:105051 | y | n | 3.58 | y |
| Sema6b | sema domain, transmembra | 20359 | MGI:1202889 | y | n | 3.58 | y |
| P2rx6 | purinergic receptor P2X, liga | 18440 | MGI:1337113 | y | n | 3.58 | y |
| Slc7a8 | solute carrier family 7 (cation | 50934 | MGI:1355323 | y | n | 3.58 | y |
| Nell2 | NEL-like 2 | 54003 | MGI:1858510 | y | n | 3.58 | y |
| Cd248 | CD248 antigen, endosialin | 70445 | MGI:1917695 | y | n | 3.58 | y |
| 33431G14Rik | RIKEN cDNA 4933431G14 g | 71265 | MGI:1918515 | y | n | 3.58 | y |
| Lrriq3 | leucine-rich repeats and IQ r | 74435 | MGI:1921685 | y | n | 3.58 | y |
| Vwc2 | von Willebrand factor C dom | 319922 | MGI:2442987 | y | n | 3.58 | y |
| Prrt3 | proline-rich transmembrane | 210673 | MGI:2444810 | y | n | 3.58 | y |
| Slc6a11 | solute carrier family 6 (neuro | 243616 | MGI:95630 | y | n | 3.58 | y |
| Kcnc1 | potassium voltage gated cha | 16502 | MGI:96667 | y | n | 3.58 | y |
| Scn5a | sodium channel, voltage-gat | 20271 | MGI:98251 | y | n | 3.58 | y |
| Pacsin1 | protein kinase C and casein | 23969 | MGI:1345181 | y | n | 3.64 | y |
| Vwa5b2 | von Willebrand factor A dom | 328643 | MGI:2681859 | y | n | 3.64 | y |
| Plce1 | phospholipase C, epsilon 1 | 74055 | MGI:1921305 | y | n | 3.68 | y |
| Stxbp1 | syntaxin binding protein 1 | 20910 | MGI:107363 | y | n | 3.70 | y |
| Pgf | placental growth factor | 18654 | MGI:105095 | y | n | 3.70 | y |
| Slc9a2 | solute carrier family 9 (sodiu | 226999 | MGI:105075 | y | n | 3.70 | y |
| Ankrd22 | ankyrin repeat domain 22 | 52024 | MGI:1277101 | y | n | 3.70 | y |
| Lrrc20 | leucine rich repeat containin | 216011 | MGI:2387182 | y | n | 3.70 | y |
| Pdzd7 | PDZ domain containing 7 | 0050304 | MGI:3608325 | y | n | 3.70 | y |
| Frmd6 | FERM domain containing 6 | 319710 | MGI:2442579 | y | n | 3.74 | y |
| Rprm | reprimo, TP53 dependent G | 67874 | MGI:1915124 | y | n | 3.77 | y |
| Dpp10 | dipeptidylpeptidase 10 | 269109 | MGI:2442409 | y | n | 3.81 | y |
| Cep250 | centrosomal protein 250 | 16328 | MGI:108084 | y | n | 3.81 | y |
| Slc37a4 | solute carrier family 37 (gluc | 14385 | MGI:1316650 | y | n | 3.91 | y |
| Il25 | interleukin 25 | 140806 | MGI:2155888 | y | n | 3.91 | y |
| Fbxl16 | F-box and leucine-rich repea | 214931 | MGI:2448488 | y | n | 3.91 | y |
| Eml1 | echinoderm microtubule ass | 68519 | MGI:1915769 | y | n | 4.00 | y |
| R3hdml | R3H domain containing-like | 0004389 | MGI:3650937 | y | n | 4.00 | y |
| Kcnab2 | potassium voltage-gated cha | 16498 | MGI:109239 | y | n | 4.00 | y |
| Tyrobp | TYRO protein tyrosine kinas | 22177 | MGI:1277211 | y | n | 4.00 | y |
| Adcy8 | adenylate cyclase 8 | 11514 | MGI:1341110 | y | n | 4.00 | y |
| Ddit4 | DNA-damage-inducible trans | 74747 | MGI:1921997 | y | n | 4.00 | y |
| Rab37 | RAB37, member RAS oncog | 58222 | MGI:1929945 | y | n | 4.00 | y |
| Ghsr | growth hormone secretagog | 208188 | MGI:2441906 | y | n | 4.00 | y |
| Pik3r5 | phosphoinositide-3-kinase re | 320207 | MGI:2443588 | y | n | 4.00 | y |
| 32416N19Rik | RIKEN cDNA 4732416N19 g | 320737 | MGI:2444634 | y | n | 4.00 | y |
| Slc37a1 | solute carrier family 37 (glyc | 224674 | MGI:2446181 | y | n | 4.00 | y |
| Pttg1ip2 | PTTG1IP family member 2 | 381716 | MGI:2686532 | y | n | 4.00 | y |
| Neurl1b | neuralized E3 ubiquitin prote | 240055 | MGI:3643092 | y | n | 4.00 | y |
| Adcy2 | adenylate cyclase 2 | 210044 | MGI:99676 | y | n | 4.00 | y |
| Tcp11 | t-complex protein 11 | 21463 | MGI:98544 | y | n | 4.09 | y |
| Trnp1 | TMF1-regulated nuclear pro | 69539 | MGI:1916789 | y | n | 4.14 | y |
| Lrrn3 | leucine rich repeat protein 3, | 16981 | MGI:106036 | y | n | 4.29 | y |
| Lnx2 | ligand of numb-protein X 2 | 140887 | MGI:2155959 | y | n | 4.29 | y |
| Kcnh2 | potassium voltage-gated cha | 16511 | MGI:1341722 | y | n | 4.32 | y |
| Lamc3 | laminin gamma 3 | 23928 | MGI:1344394 | y | n | 4.32 | y |
| Lrtm2 | leucine-rich repeats and tran | 211187 | MGI:2141485 | y | n | 4.32 | y |
| Synpo2 | synaptopodin 2 | 118449 | MGI:2153070 | y | n | 4.32 | y |
| Disc1 | disrupted in schizophrenia 1 | 244667 | MGI:2447658 | y | n | 4.32 | y |
| Ifnk | interferon kappa | 387510 | MGI:2683287 | y | n | 4.32 | y |
| Fat2 | FAT atypical cadherin 2 | 245827 | MGI:2685369 | y | n | 4.32 | y |
| Gm1043 | predicted gene 1043 | 381634 | MGI:2685889 | y | n | 4.32 | y |
| Snora33 | small nucleolar RNA, H/ACA | 0052907 | MGI:3819501 | y | n | 4.32 | y |
| Serpinb8 | serine (or cysteine) peptidas | 20725 | MGI:894657 | y | n | 4.32 | y |
| Itpr3 | inositol 1,4,5-triphosphate re | 16440 | MGI:96624 | y | n | 4.32 | y |
| Ryr2 | ryanodine receptor 2, cardia | 20191 | MGI:99685 | y | n | 4.32 | y |
| Lrrc10b | leucine rich repeat containin | 278795 | MGI:2685551 | y | n | 4.46 | y |
| Slc8a2 | solute carrier family 8 (sodiu | 110891 | MGI:107996 | y | n | 4.46 | y |
| Syt14 | synaptotagmin XIV | 329324 | MGI:2444490 | y | n | 4.49 | y |
| Mpped2 | metallophosphoesterase dor | 77015 | MGI:1924265 | y | n | 4.52 | y |

| | | | | | | | |
|---|---|---|---|---|---|---|---|
| Cacna1d | calcium channel, voltage-de | 12289 | MGI:88293 | y | n | 4.58 | y |
| Abcd2 | ATP-binding cassette, sub-fa | 26874 | MGI:1349467 | y | n | 4.58 | y |
| Syt7 | synaptotagmin VII | 54525 | MGI:1859545 | y | n | 4.58 | y |
| Angptl6 | angiopoietin-like 6 | 70726 | MGI:1917976 | y | n | 4.58 | y |
| Lrguk | leucine-rich repeats and gua | 74354 | MGI:1921604 | y | n | 4.58 | y |
| Ppp1r3e | protein phosphatase 1, regu | 105651 | MGI:2145790 | y | n | 4.58 | y |
| Nav3 | neuron navigator 3 | 260315 | MGI:2183703 | y | n | 4.58 | y |
| Gsg1l | GSG1-like | 269994 | MGI:2685483 | y | n | 4.58 | y |
| Mir339 | microRNA 339 | 723898 | MGI:3619354 | y | n | 4.58 | y |
| Cdc25b | cell division cycle 25B | 12531 | MGI:99701 | y | n | 4.64 | y |
| Fez1 | fasciculation and elongation | 235180 | MGI:2670976 | y | n | 4.67 | y |
| Serpine3 | serpin peptidase inhibitor, cl | 319433 | MGI:2442020 | y | n | 4.70 | y |
| Scn8a | sodium channel, voltage-gat | 20273 | MGI:103169 | y | n | 4.70 | y |
| Kcns3 | potassium voltage-gated cha | 238076 | MGI:1098804 | y | n | 4.75 | y |
| Spock2 | sparc/osteonectin, cwcv and | 94214 | MGI:1891351 | y | n | 4.75 | y |
| Clrn1 | clarin 1 | 229320 | MGI:2388124 | y | n | 4.81 | y |
| Nostrin | nitric oxide synthase traffick | 329416 | MGI:3606242 | y | n | 4.81 | y |
| Dll3 | delta like canonical Notch lig | 13389 | MGI:1096877 | y | n | 4.81 | y |
| Tsnaxip1 | translin-associated factor X | 72236 | MGI:1919486 | y | n | 4.81 | y |
| 310008I18R | RIKEN cDNA 1810008I18 g | 0050396 | MGI:1920875 | y | n | 4.81 | y |
| 30006D01R | RIKEN cDNA E130006D01 g | 269683 | MGI:2685527 | y | n | 4.81 | y |
| Drd3 | dopamine receptor D3 | 13490 | MGI:94925 | y | n | 4.81 | y |
| Tesc | tescalcin | 57816 | MGI:1930803 | y | n | 4.86 | y |
| Myo7a | myosin VIIA | 17921 | MGI:104510 | y | n | 4.95 | y |
| Naaladl2 | N-acetylated alpha-linked ac | 635702 | MGI:2685867 | y | n | 5.00 | y |
| Cyp46a1 | cytochrome P450, family 46, | 13116 | MGI:1341877 | y | n | 5.00 | y |
| Pak5 | p21 (RAC1) activated kinase | 241656 | MGI:1920334 | y | n | 5.00 | y |
| 30543E12R | RIKEN cDNA 4930543E12 g | 75239 | MGI:1922489 | y | n | 5.00 | y |
| Sla2 | Src-like-adaptor 2 | 77799 | MGI:1925049 | y | n | 5.00 | y |
| Rassf2 | Ras association (RalGDS/AI | 215653 | MGI:2442060 | y | n | 5.00 | y |
| Fstl4 | follistatin-like 4 | 320027 | MGI:2443199 | y | n | 5.00 | y |
| Galnt9 | polypeptide N-acetylgalactos | 231605 | MGI:2677965 | y | n | 5.00 | y |
| 30416C01R | RIKEN cDNA 5330416C01 g | 403201 | MGI:2685378 | y | n | 5.00 | y |
| Gm10248 | predicted gene 10248 | 791307 | MGI:3641931 | y | n | 5.00 | y |
| Dpp6 | dipeptidylpeptidase 6 | 13483 | MGI:94921 | y | n | 5.00 | y |
| Sstr2 | somatostatin receptor 2 | 20606 | MGI:98328 | y | n | 5.00 | y |
| Chgb | chromogranin B | 12653 | MGI:88395 | y | n | 5.13 | y |
| Ttc39b | tetratricopeptide repeat dom | 69863 | MGI:1917113 | y | n | 5.17 | y |
| Atp6v0a4 | ATPase, H+ transporting, lys | 140494 | MGI:2153480 | y | n | 5.17 | y |
| Plekha6 | pleckstrin homology domain | 240753 | MGI:2388662 | y | n | 5.17 | y |
| Obscn | obscurin, cytoskeletal calmo | 380698 | MGI:2681862 | y | n | 5.17 | y |
| Wdr25 | WD repeat domain 25 | 212198 | MGI:3045255 | y | n | 5.17 | y |
| Aknad1 | AKNA domain containing 1 | 329738 | MGI:3584453 | y | n | 5.17 | y |
| Ksr2 | kinase suppressor of ras 2 | 333050 | MGI:3610315 | y | n | 5.17 | y |
| Gpr156 | G protein-coupled receptor 1 | 239845 | MGI:2653880 | y | n | 5.32 | y |
| Sult4a1 | sulfotransferase family 4A, m | 29859 | MGI:1888971 | y | n | 5.32 | y |
| Fer1l6 | fer-1 like family member 6 | 631797 | MGI:3645398 | y | n | 5.32 | y |
| Fam124a | family with sequence similar | 629059 | MGI:3645930 | y | n | 5.32 | y |
| Miat | myocardial infarction associa | 330166 | MGI:2444886 | y | n | 5.36 | y |
| Calb2 | calbindin 2 | 12308 | MGI:101914 | y | n | 5.39 | y |
| Pdzrn3 | PDZ domain containing RIN | 55983 | MGI:1933157 | y | n | 5.43 | y |
| Dll1 | delta like canonical Notch lig | 13388 | MGI:104659 | y | n | 5.46 | y |
| Btbd17 | BTB domain containing 17 | 72014 | MGI:1919264 | y | n | 5.46 | y |
| Abtb3 | ankyrin repeat and BTB dom | 74007 | MGI:1921257 | y | n | 5.46 | y |
| Pnma2 | paraneoplastic antigen MA2 | 239157 | MGI:2444129 | y | n | 5.46 | y |
| Ttc34 | tetratricopeptide repeat dom | 242800 | MGI:2445205 | y | n | 5.46 | y |
| Rtn4rl1 | reticulon 4 receptor-like 1 | 237847 | MGI:2661375 | y | n | 5.46 | y |
| Syt13 | synaptotagmin XIII | 80976 | MGI:1933945 | y | n | 5.49 | y |
| Ppp1r14a | protein phosphatase 1, regu | 68458 | MGI:1931139 | y | n | 5.58 | y |
| Thsd7b | thrombospondin, type I, dom | 210417 | MGI:2443925 | y | n | 5.61 | y |
| Tmem91 | transmembrane protein 91 | 320208 | MGI:2443589 | y | n | 5.64 | y |
| Pcdh8 | protocadherin 8 | 18530 | MGI:1306800 | y | n | 5.70 | y |
| Dock10 | dedicator of cytokinesis 10 | 210293 | MGI:2146320 | y | n | 5.70 | y |
| Slc52a3 | solute carrier protein family 5 | 69698 | MGI:1916948 | y | n | 5.73 | y |
| Cbln1os | cerebellin 1 precursor protei | 0004029 | MGI:3780864 | y | n | 5.75 | y |
| Sting1 | stimulator of interferon respo | 72512 | MGI:1919762 | y | n | 5.88 | y |

| | | | | | | | |
|---|---|---|---|---|---|---|---|
| Hydin | HYDIN, axonemal central pa | 244653 | MGI:2389007 | y | n | 5.91 | y |
| Atp2b2 | ATPase, Ca++ transporting, | 11941 | MGI:105368 | y | n | 6.00 | y |
| Nefm | neurofilament, medium poly | 18040 | MGI:97314 | y | n | 6.07 | y |
| Dynlrb2 | dynein light chain roadblock- | 75465 | MGI:1922715 | y | n | 6.09 | y |
| Dll4 | delta like canonical Notch lig | 54485 | MGI:1859388 | y | n | 6.09 | y |
| Ablim2 | actin-binding LIM protein 2 | 231148 | MGI:2385758 | y | n | 6.09 | y |
| Gng8 | guanine nucleotide binding p | 14709 | MGI:109163 | y | n | 6.17 | y |
| Ecel1 | endothelin converting enzym | 13599 | MGI:1343461 | y | n | 6.17 | y |
| Tas1r1 | taste receptor, type 1, memb | 110326 | MGI:1927505 | y | n | 6.17 | y |
| Cacng5 | calcium channel, voltage-de | 140723 | MGI:2157946 | y | n | 6.17 | y |
| Myo16 | myosin XVI | 244281 | MGI:2685951 | y | n | 6.17 | y |
| Hecw1 | HECT, C2 and WW domain | 94253 | MGI:2444115 | y | n | 6.32 | y |
| Igsf21 | immunoglobulin superfamily | 230868 | MGI:2681842 | y | n | 6.32 | y |
| Grxcr1 | glutaredoxin, cysteine rich 1 | 433899 | MGI:3577767 | y | n | 6.38 | y |
| Mmp24 | matrix metallopeptidase 24 | 17391 | MGI:1341867 | y | n | 6.39 | y |
| Scn11a | sodium channel, voltage-gat | 24046 | MGI:1345149 | y | n | 6.39 | y |
| Ttc21a | tetratricopeptide repeat dom | 74052 | MGI:1921302 | y | n | 6.39 | y |
| Sgpp2 | sphingosine-1-phosphate ph | 433323 | MGI:3589109 | y | n | 6.39 | y |
| Cbln1 | cerebellin 1 precursor protei | 12404 | MGI:88281 | y | n | 6.39 | y |
| 30470P17R | RIKEN cDNA 4930470P17 g | 67637 | MGI:1914887 | y | n | 6.46 | y |
| Coro2a | coronin, actin binding protei | 107684 | MGI:1345966 | y | n | 6.52 | y |
| Slc8a1 | solute carrier family 8 (sodiu | 20541 | MGI:107956 | y | n | 6.52 | y |
| Ttc29 | tetratricopeptide repeat dom | 73301 | MGI:1920551 | y | n | 6.52 | y |
| Kcna10 | potassium voltage-gated cha | 242151 | MGI:3037820 | y | n | 6.52 | y |
| Iqub | IQ motif and ubiquitin domai | 214704 | MGI:3041159 | y | n | 6.52 | y |
| Rbfox3 | RNA binding protein, fox-1 h | 52897 | MGI:106368 | y | n | 6.64 | y |
| Gm10432 | predicted gene 10432 | 0003871 | MGI:3642011 | y | n | 6.64 | y |
| Mgat5b | mannoside acetylglucosami | 268510 | MGI:3606200 | y | n | 6.75 | y |
| Laptm5 | lysosomal-associated protei | 16792 | MGI:108046 | y | n | 6.81 | y |
| Ak1 | adenylate kinase 1 | 11636 | MGI:87977 | y | n | 6.86 | y |
| Fam78b | family with sequence similar | 226610 | MGI:2443050 | y | n | 6.86 | y |
| Kcnh7 | potassium voltage-gated cha | 170738 | MGI:2159566 | y | n | 6.86 | y |
| Insc | INSC spindle orientation ada | 233752 | MGI:1917942 | y | n | 6.95 | y |
| Faim2 | Fas apoptotic inhibitory mole | 72393 | MGI:1919643 | y | n | 7.00 | y |
| Ankfn1 | ankyrin-repeat and fibronect | 382543 | MGI:2686021 | y | n | 7.09 | y |
| Rph3a | rabphilin 3A | 19894 | MGI:102788 | y | n | 7.09 | y |
| Ush2a | usherin | 22283 | MGI:1341292 | y | n | 7.13 | y |
| Srrm4 | serine/arginine repetitive ma | 68955 | MGI:1916205 | y | n | 7.17 | y |
| Cxcl14 | C-X-C motif chemokine ligar | 57266 | MGI:1888514 | y | n | 7.21 | y |
| Mogat1 | monoacylglycerol O-acyltran | 68393 | MGI:1915643 | y | n | 7.32 | y |
| Crmp1 | collapsin response mediator | 12933 | MGI:107793 | y | n | 7.39 | y |
| St8sia3 | ST8 alpha-N-acetyl-neurami | 20451 | MGI:106019 | y | n | 7.52 | y |
| Grp | gastrin releasing peptide | 225642 | MGI:95833 | y | n | 7.55 | y |
| Mfng | MFNG O-fucosylpeptide 3-b | 17305 | MGI:1095404 | y | n | 7.70 | y |
| Chrna9 | cholinergic receptor, nicotini | 231252 | MGI:1202403 | y | n | 7.93 | y |
| Myo3a | myosin IIIA | 667663 | MGI:2183924 | y | n | 8.13 | y |
| Bdnf | brain derived neurotrophic fa | 12064 | MGI:88145 | y | n | 8.13 | y |
| Pvalb | parvalbumin | 19293 | MGI:97821 | y | n | 8.38 | y |
| Rasd2 | RASD family, member 2 | 75141 | MGI:1922391 | y | n | 8.52 | y |
| Lhfpl5 | lipoma HMGIC fusion partne | 328789 | MGI:1915382 | y | n | 8.55 | y |
| Dync1i1 | dynein cytoplasmic 1 interm | 13426 | MGI:107743 | y | n | -6.70 | n |
| Dok5 | docking protein 5 | 76829 | MGI:1924079 | y | n | -6.09 | n |
| Trp53i11 | transformation related protei | 277414 | MGI:2670995 | y | n | -5.81 | n |
| Itga9 | integrin alpha 9 | 104099 | MGI:104756 | y | n | -5.58 | n |
| Dctd | dCMP deaminase | 320685 | MGI:2444529 | y | n | -5.58 | n |
| Sorl1 | sortilin-related receptor, LDL | 20660 | MGI:1202296 | y | n | -5.32 | n |
| Gins2 | GINS complex subunit 2 | 272551 | MGI:1921019 | y | n | -5.17 | n |
| Mmd2 | monocyte to macrophage di | 75104 | MGI:1922354 | y | n | -5.17 | n |
| Wdr59 | WD repeat domain 59 | 319481 | MGI:2442115 | y | n | -5.17 | n |
| Ark2c | arkadia (RNF111) C-termina | 225743 | MGI:2444521 | y | n | -5.17 | n |
| Prtg | protogenin | 235472 | MGI:2444710 | y | n | -5.17 | n |
| Vps13d | vacuolar protein sorting 13D | 230895 | MGI:2448530 | y | n | -5.17 | n |
| Gm266 | predicted gene 266 | 212539 | MGI:2685112 | y | n | -5.17 | n |
| Slc7a5 | solute carrier family 7 (catior | 20539 | MGI:1298205 | y | n | -5.00 | n |
| Bace2 | beta-site APP-cleaving enzy | 56175 | MGI:1860440 | y | n | -5.00 | n |
| Glipr2 | GLI pathogenesis-related 2 | 384009 | MGI:1917770 | y | n | -5.00 | n |
