## Supplementary material for "CASZ1 regulates the rate at which outer hair cells mature and is required for hearing": Table S2

**Table S2. Gene expression differences between organs of Corti of *Casz1$^{fl/fl}$;Tg(Sox10-Cre)* mice versus WT mice at P4.** (1132 genes are listed.)

| Gene symbol | Gene name | Entrez gene ID | MGI gene ID | Feature type* | logFC *Casz1(fl/fl);Tg(Sox10-Cre)* vs WT | FDR-adjusted P (Benjamini-Yekutieli method) | Predominantly IHC expressed in organ of Corti? y/n | Predominantly OHC expressed in organ of Corti? y/n | Predominantly OHC *and* IHC expressed in organ of Corti? y/n | ATOH1 target gene? y/n | Annotated with 'sensory perception of sound' GO term? y/n | Annotated with 'cilium' GO term? y/n |
|---|---|---|---|---|---|---|---|---|---|---|---|---|
| Serpina1c | serine (or cysteine) peptidase | 20702 | MGI:891969 | protein coding | 10.207 | 1.02E-10 | n | y | n | n | n | n |
| Ccnb1ip1 | cyclin B1 interacting protein 1 | 239083 | MGI:2685134 | protein coding | 9.598 | 3.58E-07 | n | n | n | n | n | n |
| Serpina1a | serine (or cysteine) peptidase | 20700 | MGI:891971 | protein coding | 9.263 | 7.93E-06 | n | n | n | n | n | n |
| Vwa5b1 | von Willebrand factor A doma | 75718 | MGI:1922968 | protein coding | 7.235 | 1.05E-11 | n | n | n | n | n | n |
| Myl1 | myosin, light polypeptide 1 | 17901 | MGI:97269 | protein coding | 6.895 | 3.59E-08 | n | n | n | n | n | n |
| Serpina1b | serine (or cysteine) preptidas | 20701 | MGI:891970 | protein coding | 6.253 | 1.93E-07 | n | n | n | n | n | n |
| Abcc2 | ATP-binding cassette, sub-fa | 12780 | MGI:1352447 | protein coding | 5.689 | 5.56E-09 | n | n | n | n | n | n |
| Cyp2j11 | cytochrome P450, family 2, su | 100066 | MGI:2140224 | protein coding | 5.393 | 3.00E-07 | n | n | n | n | n | n |
| Cfap107 | cilia and flagella associated p | 69364 | MGI:1916614 | protein coding | 5.320 | 9.36E-08 | n | n | n | n | n | y |
| Cort | cortistatin | 12854 | MGI:109538 | protein coding | 5.267 | 6.71E-09 | n | n | n | n | n | n |
| Lhb | luteinizing hormone beta | 16866 | MGI:96782 | protein coding | 4.935 | 3.52E-07 | n | n | n | n | n | n |
| AI429214 | expressed sequence AI42921 | 621080 | MGI:2142538 | protein coding | 4.734 | 5.41E-11 | n | n | n | n | n | n |
| Tagap | T cell activation Rho GTPase | 72536 | MGI:3615484 | protein coding | 4.663 | 8.50E-09 | n | n | n | n | n | n |
| Erich5 | glutamate rich 5 | 239368 | MGI:2447772 | protein coding | 4.454 | 1.10E-05 | n | n | n | y | n | n |
| 4932438H23Rik | RIKEN cDNA 4932438H23 ge | 74387 | MGI:1921637 | protein coding | 4.385 | 4.97E-07 | n | n | n | n | n | n |
| Zmynd10 | zinc finger, MYND domain co | 114602 | MGI:2387863 | protein coding | 4.157 | 7.00E-08 | n | n | n | n | n | n |
| Cysltr1 | cysteinyl leukotriene receptor | 58861 | MGI:1926218 | protein coding | 4.062 | 1.34E-09 | n | n | n | n | n | n |
| Gm14461 | predicted gene 14461 | 329436 | MGI:3651589 | protein coding | 3.973 | 1.33E-08 | n | n | n | n | n | n |
| Masp2 | MBL associated serine protea | 17175 | MGI:1330832 | protein coding | 3.870 | 5.15E-08 | n | n | n | n | n | n |
| Iqca1 | IQ motif containing with AAA | 74918 | MGI:1922168 | protein coding | 3.867 | 2.37E-10 | n | n | n | n | n | y |
| Nme9 | NME/NM23 family member 9 | 623534 | MGI:4359686 | protein coding | 3.778 | 1.08E-08 | n | n | n | n | n | n |
| Cramp1 | cramped chromatin regulator | 57354 | MGI:1930190 | protein coding | 3.769 | 4.34E-07 | n | n | n | n | n | n |
| Pou2af1 | POU domain, class 2, associa | 18985 | MGI:105086 | protein coding | 3.648 | 1.07E-05 | n | n | n | y | n | n |
| Trhde | TRH-degrading enzyme | 237553 | MGI:2384311 | protein coding | 3.646 | 1.04E-07 | n | n | n | n | n | n |
| D7Ertd443e | DNA segment, Chr 7, ERATC | 71007 | MGI:1196431 | protein coding | 3.548 | 2.36E-07 | n | n | y | n | n | n |
| Rbm5 | RNA binding motif protein 5 | 83486 | MGI:1933204 | protein coding | 3.537 | 1.85E-14 | n | n | n | n | n | n |
| Ptp4a2 | protein tyrosine phosphatase | 19244 | MGI:1277117 | protein coding | 3.494 | 1.50E-13 | n | n | n | n | n | n |
| Cfap161 | cilia and flagella associated p | 75556 | MGI:1922806 | protein coding | 3.431 | 9.01E-10 | n | n | n | y | n | y |
| Arhgap22 | Rho GTPase activating protei | 239027 | MGI:2443418 | protein coding | 3.412 | 1.20E-06 | n | n | n | n | n | n |
| Trim21 | tripartite motif-containing 21 | 20821 | MGI:106657 | protein coding | 3.391 | 9.57E-05 | n | n | n | n | n | n |
| Zer1 | zyg-11 related, cell cycle regu | 227693 | MGI:2442511 | protein coding | 3.228 | 8.58E-13 | n | n | n | n | n | n |
| Btnl2 | butyrophilin-like 2 | 547431 | MGI:1859549 | protein coding | 3.201 | 9.26E-10 | n | n | n | n | n | n |
| Ccdc113 | coiled-coil domain containing | 244608 | MGI:3606076 | protein coding | 3.198 | 3.45E-07 | n | n | n | n | n | y |
| Prokr2 | prokineticin receptor 2 | 246313 | MGI:2181363 | protein coding | 3.161 | 6.39E-08 | n | n | n | n | n | n |
| Asb1 | ankyrin repeat and SOCS box | 65247 | MGI:1929735 | protein coding | 3.099 | 2.57E-06 | n | n | n | n | n | n |
| Retn | resistin | 57264 | MGI:1888506 | protein coding | 3.053 | 7.93E-06 | n | n | n | n | n | n |
| Insm2 | insulinoma-associated 2 | 56856 | MGI:1930787 | protein coding | 3.040 | 6.53E-06 | n | y** (PMID: 30305733) | n | y | n | n |
| Xpnpep2 | X-prolyl aminopeptidase (ami | 170745 | MGI:2180001 | protein coding | 3.031 | 1.98E-05 | n | n | n | n | n | n |
| Ocm | oncomodulin | 18261 | MGI:97401 | protein coding | 3.006 | 2.61E-10 | n | y | n | n | y** (PMID: 26843644) | n |
| Zfp868 | zinc finger protein 868 | 234362 | MGI:2142546 | protein coding | 2.996 | 1.85E-09 | n | n | n | n | n | n |
| Xcr1 | chemokine (C motif) receptor | 23832 | MGI:1346338 | protein coding | 2.996 | 8.97E-07 | n | n | n | n | n | n |
| Dnajc5b | DnaJ heat shock protein fami | 66326 | MGI:1913576 | protein coding | 2.985 | 8.73E-08 | n | n | n | n | n | n |
| Nlgn1 | neuroligin 1 | 192167 | MGI:2179435 | protein coding | 2.919 | 3.98E-09 | n | n | n | n | n | n |
| Meiob | meiosis specific with OB dom | 75178 | MGI:1922428 | protein coding | 2.917 | 2.29E-05 | n | n | n | n | n | n |
| Wdr95 | WD40 repeat domain 95 | 381693 | MGI:1923042 | protein coding | 2.911 | 3.75E-06 | y | n | n | n | n | n |
| Pianp | PILR alpha associated neural | 319352 | MGI:2441908 | protein coding | 2.895 | 1.25E-08 | n | n | n | n | n | n |
| Mal2 | mal, T cell differentiation prot | 105853 | MGI:2146021 | protein coding | 2.868 | 3.85E-13 | n | n | n | n | n | n |
| Col6a6 | collagen, type VI, alpha 6 | 245026 | MGI:2444259 | protein coding | 2.838 | 1.18E-07 | n | n | n | n | n | n |
| Grik3 | glutamate receptor, ionotropic | 14807 | MGI:95816 | protein coding | 2.833 | 7.37E-07 | n | n | n | y | n | n |
| Supt16 | SPT16, facilitates chromatin r | 114741 | MGI:1890948 | protein coding | 2.782 | 1.85E-14 | n | n | n | n | n | n |
| Lgals3 | lectin, galactose binding, solu | 16854 | MGI:96778 | protein coding | 2.742 | 7.82E-09 | n | n | n | n | n | n |
| Samd11 | sterile alpha motif domain cor | 231004 | MGI:2446220 | protein coding | 2.715 | 4.05E-11 | n | n | n | n | n | n |
| Lpar2 | lysophosphatidic acid recepto | 53978 | MGI:1858422 | protein coding | 2.684 | 1.33E-09 | n | n | n | n | n | n |
| Cfap95 | cilia and flagella associated p | 67483 | MGI:1914733 | protein coding | 2.654 | 1.65E-05 | n | n | n | n | n | y |
| Tmem130 | transmembrane protein 130 | 243339 | MGI:3607706 | protein coding | 2.562 | 9.71E-08 | n | n | n | n | n | n |
| Cep15 | centrosomal protein 15 | 218734 | MGI:1917937 | protein coding | 2.544 | 2.14E-12 | n | n | n | n | n | y |
| Cpne9 | copine family member IX | 211232 | MGI:2443052 | protein coding | 2.534 | 1.53E-10 | n | n | y | n | n | n |
| Hgf | hepatocyte growth factor | 15234 | MGI:96079 | protein coding | 2.504 | 1.97E-07 | n | n | n | y | y** (PMID: 19576567, 32152201) | n |
| Nsg2 | neuron specific gene family m | 18197 | MGI:1202070 | protein coding | 2.503 | 2.48E-05 | n | y | n | y | n | n |
| Dcdc2a | doublecortin domain containir | 195208 | MGI:2652818 | protein coding | 2.436 | 9.36E-08 | n | n | n | n | y | y |

*Only protein coding genes are listed.

**Manually updated based on indicated PMID.

| | | | | | | | | | | | | |
|---|---|---|---|---|---|---|---|---|---|---|---|---|
| Cfap45 | cilia and flagella associated p | 71870 | MGI:1919120 | protein coding | 2.417 | 2.82E-09 | n | n | n | n | n | y |
| Nppa | natriuretic peptide type A | 230899 | MGI:97367 | protein coding | 2.357 | 2.94E-06 | n | y | n | n | n | n |
| Fbp1 | fructose bisphosphatase 1 | 14121 | MGI:95492 | protein coding | 2.353 | 9.38E-06 | n | n | n | n | n | n |
| Tekt4 | tektin 4 | 71840 | MGI:1919090 | protein coding | 2.325 | 0.000514195 | n | n | n | n | n | y |
| Mapk10 | mitogen-activated protein kina | 26414 | MGI:1346863 | protein coding | 2.317 | 4.10E-10 | n | n | n | n | n | n |
| Myh13 | myosin, heavy polypeptide 13 | 544791 | MGI:1339967 | protein coding | 2.266 | 3.70E-07 | n | n | n | n | n | n |
| Cpeb4 | cytoplasmic polyadenylation e | 67579 | MGI:1914829 | protein coding | 2.252 | 0.000154963 | n | n | n | n | n | n |
| Exd1 | exonuclease 3'-5' domain cor | 241624 | MGI:3045306 | protein coding | 2.240 | 4.49E-06 | n | n | n | y | n | n |
| Slc39a2 | solute carrier family 39 (zinc t | 214922 | MGI:2684326 | protein coding | 2.212 | 1.31E-07 | n | y | n | y | n | n |
| Nod2 | nucleotide-binding oligomeriz | 257632 | MGI:2429397 | protein coding | 2.209 | 1.82E-07 | n | n | n | n | n | n |
| Irak4 | interleukin-1 receptor-associa | 266632 | MGI:2182474 | protein coding | 2.201 | 5.37E-07 | n | n | n | n | n | n |
| Khdrbs2 | KH domain containing, RNA b | 170771 | MGI:2159649 | protein coding | 2.200 | 6.50E-09 | n | n | y | n | n | n |
| Tmem87a | transmembrane protein 87A | 211499 | MGI:2441844 | protein coding | 2.193 | 4.05E-11 | n | n | n | y | n | n |
| Mroh7 | maestro heat-like repeat fami | 381538 | MGI:2685873 | protein coding | 2.177 | 6.33E-06 | n | n | n | n | n | n |
| Ryr3 | ryanodine receptor 3 | 20192 | MGI:99684 | protein coding | 2.153 | 0.000367223 | n | n | n | n | n | n |
| Pla2g4e | phospholipase A2, group IVE | 329502 | MGI:1919144 | protein coding | 2.146 | 7.55E-11 | n | n | n | n | n | n |
| Gfra2 | glial cell line derived neurotro | 14586 | MGI:1195462 | protein coding | 2.140 | 1.03E-11 | n | n | n | y | n | n |
| Il25 | interleukin 25 | 140806 | MGI:2155888 | protein coding | 2.111 | 9.86E-06 | n | n | y | y | n | n |
| Rd3 | retinal degeneration 3 | 74023 | MGI:1921273 | protein coding | 2.071 | 3.81E-05 | n | n | n | n | n | y |
| Tnfrsf10b | tumor necrosis factor recepto | 21933 | MGI:1341090 | protein coding | 2.053 | 4.70E-11 | n | n | n | n | n | n |
| Nxnl1 | nucleoredoxin-like 1 | 234404 | MGI:1924446 | protein coding | 2.041 | 1.18E-05 | n | n | n | n | n | y |
| Rasgrf1 | RAS protein-specific guanine | 19417 | MGI:99694 | protein coding | 1.981 | 1.40E-05 | n | n | n | n | n | n |
| Gpr151 | G protein-coupled receptor 15 | 240239 | MGI:2441887 | protein coding | 1.927 | 6.56E-06 | n | n | n | y | n | n |
| Shcbp1l | Shc SH2-domain binding prot | 71836 | MGI:1919086 | protein coding | 1.921 | 0.00076909 | n | n | n | n | n | n |
| Plbd1 | phospholipase B domain cont | 66857 | MGI:1914107 | protein coding | 1.879 | 2.68E-05 | n | n | n | n | n | n |
| Kcnj4 | potassium inwardly-rectifying | 16520 | MGI:104743 | protein coding | 1.859 | 0.00018791 | n | n | n | y | n | n |
| Ppp2r2c | protein phosphatase 2, regula | 269643 | MGI:2442660 | protein coding | 1.815 | 1.31E-07 | n | n | n | n | n | n |
| Upk1a | uroplakin 1A | 109637 | MGI:98911 | protein coding | 1.813 | 0.000227957 | n | n | n | n | n | n |
| Cfap52 | cilia and flagella associated p | 71860 | MGI:1919110 | protein coding | 1.811 | 5.56E-09 | n | n | y | n | n | y |
| Erlec1 | endoplasmic reticulum lectin | 66753 | MGI:1914003 | protein coding | 1.793 | 0.000481898 | n | n | n | n | n | n |
| Cap1 | cyclase associated actin cyto | 12331 | MGI:88262 | protein coding | 1.784 | 1.77E-12 | n | n | n | y | n | n |
| Rph3a | rabphilin 3A | 19894 | MGI:102788 | protein coding | 1.777 | 1.41E-07 | n | y | n | y | n | n |
| Spag8 | sperm associated antigen 8 | 433700 | MGI:3056295 | protein coding | 1.765 | 3.24E-09 | n | n | n | n | n | y |
| Phyhip | phytanoyl-CoA hydroxylase ir | 105653 | MGI:1860417 | protein coding | 1.720 | 5.37E-06 | n | n | n | n | n | n |
| Sema3e | sema domain, immunoglobuli | 20349 | MGI:1340034 | protein coding | 1.703 | 4.30E-05 | n | n | n | n | n | n |
| Galnt9 | polypeptide N-acetylgalactosa | 231605 | MGI:2677965 | protein coding | 1.677 | 4.20E-06 | n | n | y | y | n | n |
| Cyp2j12 | cytochrome P450, family 2, su | 242546 | MGI:3717097 | protein coding | 1.672 | 2.58E-07 | n | y | n | n | n | n |
| Ptpn22 | protein tyrosine phosphatase, | 19260 | MGI:107170 | protein coding | 1.668 | 3.31E-08 | n | n | n | n | n | n |
| Tmem150c | transmembrane protein 150C | 231503 | MGI:3041258 | protein coding | 1.667 | 5.38E-08 | n | n | n | n | n | n |
| Emp1 | epithelial membrane protein 1 | 13730 | MGI:107941 | protein coding | 1.665 | 9.49E-10 | n | n | n | y | n | n |
| Ppp1r14d | protein phosphatase 1, regula | 72112 | MGI:1919362 | protein coding | 1.664 | 0.000131243 | n | y | n | n | n | n |
| Vipr2 | vasoactive intestinal peptide r | 22355 | MGI:107166 | protein coding | 1.660 | 4.56E-07 | n | n | n | n | n | n |
| Coro2a | coronin, actin binding protein | 107684 | MGI:1345966 | protein coding | 1.655 | 8.73E-08 | n | y | n | y | n | n |
| Chst4 | carbohydrate sulfotransferase | 26887 | MGI:1349479 | protein coding | 1.648 | 5.86E-08 | n | n | n | n | n | n |
| Cacna1h | calcium channel, voltage-dep | 58226 | MGI:1928842 | protein coding | 1.646 | 1.92E-07 | n | n | n | n | n | n |
| Ogdhl | oxoglutarate dehydrogenase- | 239017 | MGI:3616088 | protein coding | 1.626 | 1.29E-09 | n | n | n | n | n | n |
| Rgs7 | regulator of G protein signalin | 24012 | MGI:1346089 | protein coding | 1.620 | 9.76E-06 | n | n | n | n | n | n |
| Cacng2 | calcium channel, voltage-dep | 12300 | MGI:1316660 | protein coding | 1.613 | 3.70E-07 | n | y | n | n | n | n |
| Dnah7a | dynein, axonemal, heavy cha | 627872 | MGI:2685838 | protein coding | 1.612 | 0.000126008 | n | n | n | n | n | y |
| Nol4 | nucleolar protein 4 | 319211 | MGI:2441684 | protein coding | 1.606 | 7.52E-05 | n | n | n | n | n | n |
| Aknad1 | AKNA domain containing 1 | 329738 | MGI:3584453 | protein coding | 1.599 | 7.11E-05 | n | n | n | y | n | n |
| Ntng2 | netrin G2 | 171171 | MGI:2159341 | protein coding | 1.598 | 3.55E-06 | n | n | n | y | n | n |
| Brinp1 | bone morphogenic protein/ret | 56710 | MGI:1928478 | protein coding | 1.597 | 3.09E-09 | n | n | n | n | n | n |
| Ptx4 | pentraxin 4 | 68509 | MGI:1915759 | protein coding | 1.584 | 1.47E-05 | n | n | n | n | n | n |
| Dynlt1b | dynein light chain Tctex-type | 21648 | MGI:98643 | protein coding | 1.569 | 2.89E-05 | n | n | n | n | n | n |
| Grk1 | G protein-coupled receptor ki | 24013 | MGI:1345146 | protein coding | 1.544 | 2.17E-05 | n | y | n | n | n | y |
| Ano2 | anoctamin 2 | 243634 | MGI:2387214 | protein coding | 1.543 | 1.21E-06 | n | n | n | n | n | y |
| Il3ra | interleukin 3 receptor, alpha c | 16188 | MGI:96553 | protein coding | 1.537 | 4.32E-06 | n | n | n | n | n | n |
| R3hdml | R3H domain containing-like | 100043899 | MGI:3650937 | protein coding | 1.535 | 6.39E-08 | n | n | y | y | n | n |
| Ocel1 | occludin/ELL domain containi | 77090 | MGI:1924340 | protein coding | 1.526 | 1.63E-06 | n | n | n | y | n | n |
| Mfap1a | microfibrillar-associated prote | 67532 | MGI:1914782 | protein coding | 1.518 | 1.93E-09 | n | n | n | n | n | n |
| Cfap206 | cilia and flagella associated p | 69329 | MGI:1916579 | protein coding | 1.517 | 7.22E-09 | n | n | y | n | n | y |
| Nrip2 | nuclear receptor interacting p | 60345 | MGI:1891884 | protein coding | 1.507 | 7.24E-08 | n | n | n | n | n | n |
| Chst9 | carbohydrate sulfotransferase | 71367 | MGI:1918617 | protein coding | 1.497 | 6.02E-05 | n | n | n | n | n | n |
| Ccl28 | C-C motif chemokine ligand 2 | 56838 | MGI:1861731 | protein coding | 1.493 | 0.000362603 | n | n | n | n | n | n |
| Car7 | carbonic anhydrase 7 | 12354 | MGI:103100 | protein coding | 1.484 | 1.29E-08 | n | n | y | n | n | n |
| E230025N22Rik | Riken cDNA E230025N22 ge | 240216 | MGI:3687212 | protein coding | 1.475 | 5.03E-08 | y | n | n | n | n | n |

| | | | | | | | | | | | | |
|---|---|---|---|---|---|---|---|---|---|---|---|---|
| Dock5 | dedicator of cytokinesis 5 | 68813 | MGI:2652871 | protein coding | 1.468 | 3.31E-08 | n | n | n | n | n | n |
| Cysltr2 | cysteinyl leukotriene receptor | 70086 | MGI:1917336 | protein coding | 1.468 | 1.37E-05 | n | n | n | n | n | n |
| Gprc5a | G protein-coupled receptor, fa | 232431 | MGI:1891250 | protein coding | 1.466 | 4.90E-06 | n | n | n | n | n | n |
| Abcc8 | ATP-binding cassette, sub-fa | 20927 | MGI:1352629 | protein coding | 1.462 | 5.87E-09 | n | n | n | n | n | n |
| Zkscan16 | zinc finger with KRAB and SC | 100041581 | MGI:3510405 | protein coding | 1.458 | 0.000246065 | n | n | n | y | n | n |
| Pcsk1 | proprotein convertase subtilis | 18548 | MGI:97511 | protein coding | 1.432 | 1.97E-06 | n | n | n | n | n | n |
| Zfp963 | zinc finger protein 963 | 620419 | MGI:4867078 | protein coding | 1.430 | 5.65E-08 | n | n | n | y | n | n |
| Edem1 | ER degradation enhancer, ma | 192193 | MGI:2180139 | protein coding | 1.425 | 9.01E-10 | n | n | n | n | n | n |
| 6430571L13Rik | RIKEN cDNA 6430571L13 ge | 235599 | MGI:2445137 | protein coding | 1.424 | 9.52E-05 | n | n | n | n | n | n |
| Smim18 | small integral membrane prot | 72632 | MGI:1919882 | protein coding | 1.424 | 1.81E-06 | y | n | n | n | n | n |
| Kcnk10 | potassium channel, subfamily | 72258 | MGI:1919508 | protein coding | 1.416 | 0.000415168 | n | n | n | n | n | n |
| Setd6 | SET domain containing 6 | 66083 | MGI:1913333 | protein coding | 1.409 | 1.30E-07 | n | n | n | n | n | n |
| Zfp933 | zinc finger protein 933 | 242747 | MGI:1922865 | protein coding | 1.407 | 2.00E-05 | n | n | n | n | n | n |
| Tslp | thymic stromal lymphopoietin | 53603 | MGI:1855696 | protein coding | 1.401 | 1.20E-06 | n | n | n | n | n | n |
| Eno1b | enolase 1B, retrotransposed | 433182 | MGI:3648653 | protein coding | 1.397 | 7.11E-11 | n | n | n | n | n | n |
| 2410004P03Rik | RIKEN cDNA 2410004P03 ge | 73667 | MGI:1920917 | protein coding | 1.387 | 4.13E-06 | n | n | n | n | n | n |
| Cdkl1 | cyclin dependent kinase like 1 | 71091 | MGI:1918341 | protein coding | 1.384 | 2.94E-05 | n | n | y | n | n | y |
| Sync | syncoilin | 68828 | MGI:1916078 | protein coding | 1.382 | 2.25E-05 | n | n | n | n | n | n |
| Rem2 | rad and gem related GTP bin | 140743 | MGI:2155260 | protein coding | 1.382 | 1.62E-08 | n | n | n | n | n | n |
| Prss56 | serine protease 56 | 69453 | MGI:1916703 | protein coding | 1.381 | 3.08E-07 | n | n | n | n | n | n |
| AI593442 | expressed sequence AI59344 | 330941 | MGI:2143099 | protein coding | 1.370 | 1.26E-07 | n | y | n | n | n | n |
| Gxylt2 | glucoside xylosyltransferase 2 | 232313 | MGI:2682940 | protein coding | 1.368 | 0.000117498 | n | n | n | n | n | n |
| Pak5 | p21 (RAC1) activated kinase | 241656 | MGI:1920334 | protein coding | 1.367 | 0.000774177 | n | n | n | y | n | n |
| Wdfy1 | WD repeat and FYVE domain | 69368 | MGI:1916618 | protein coding | 1.363 | 7.48E-05 | n | n | n | n | n | n |
| Medag | mesenteric estrogen depende | 70717 | MGI:1917967 | protein coding | 1.357 | 0.000109571 | n | n | n | n | n | n |
| Ppip5k1 | diphosphoinositol pentakisph | 327655 | MGI:2443281 | protein coding | 1.355 | 7.22E-09 | n | y | n | n | n | n |
| Tm6sf2 | transmembrane 6 superfamily | 107770 | MGI:1933210 | protein coding | 1.353 | 0.000425409 | n | n | n | n | n | n |
| F2rl1 | F2R like trypsin receptor 1 | 14063 | MGI:101910 | protein coding | 1.337 | 6.33E-06 | n | n | n | n | n | n |
| Ralyl | RALY RNA binding protein-lik | 76897 | MGI:1924147 | protein coding | 1.336 | 2.27E-05 | n | n | n | n | n | n |
| Dynlt5 | dynein light chain Tctex-type | 67344 | MGI:1914594 | protein coding | 1.334 | 0.000439307 | n | n | y | n | n | n |
| Ppp1r13l | protein phosphatase 1, regula | 333654 | MGI:3525053 | protein coding | 1.319 | 0.000301998 | n | n | n | n | n | n |
| Ccdc121rt1 | coiled-coil domain containing | 403180 | MGI:2685601 | protein coding | 1.315 | 0.000535223 | n | n | n | n | n | n |
| Ptgs2 | prostaglandin-endoperoxide s | 19225 | MGI:97798 | protein coding | 1.297 | 0.000771212 | n | n | n | n | n | n |
| Ptgir | prostaglandin I receptor (IP) | 19222 | MGI:99535 | protein coding | 1.296 | 4.05E-07 | n | y | n | n | n | n |
| Ppwd1 | peptidylprolyl isomerase dom | 238831 | MGI:2443069 | protein coding | 1.296 | 2.76E-07 | n | n | n | n | n | n |
| Cd59a | CD59a antigen | 12509 | MGI:109177 | protein coding | 1.296 | 7.22E-09 | n | n | n | n | n | n |
| Hif3a | hypoxia inducible factor 3, alp | 53417 | MGI:1859778 | protein coding | 1.291 | 0.000205291 | n | n | n | y | n | n |
| Acss2 | acyl-CoA synthetase short-ch | 60525 | MGI:1890410 | protein coding | 1.288 | 1.05E-09 | n | n | y | n | n | n |
| Kcnf1 | potassium voltage-gated char | 382571 | MGI:2687399 | protein coding | 1.282 | 3.30E-05 | n | n | n | y | n | y |
| Cabcoco1 | ciliary associated calcium bin | 73287 | MGI:1920537 | protein coding | 1.281 | 5.47E-09 | n | n | n | n | n | y |
| Efcab14 | EF-hand calcium binding dom | 230648 | MGI:2442397 | protein coding | 1.281 | 1.82E-10 | n | n | n | n | n | n |
| Mro | maestro | 71263 | MGI:2152817 | protein coding | 1.269 | 0.000310131 | n | n | n | y | n | n |
| Hemk1 | HemK methyltransferase fam | 69536 | MGI:1916786 | protein coding | 1.260 | 1.28E-08 | n | n | n | n | n | n |
| Oacyl | O-acyltransferase like | 319888 | MGI:2442915 | protein coding | 1.259 | 0.000105999 | n | n | n | y | n | n |
| Hddc3 | HD domain containing 3 | 68695 | MGI:1915945 | protein coding | 1.257 | 4.89E-06 | n | n | n | n | n | n |
| Armc2 | armadillo repeat containing 2 | 213402 | MGI:1916449 | protein coding | 1.256 | 0.000163137 | n | n | n | n | n | n |
| Laptm5 | lysosomal-associated protein | 16792 | MGI:108046 | protein coding | 1.255 | 1.16E-08 | n | y | n | y | n | n |
| Slc18a1 | solute carrier family 18 (vesic | 110877 | MGI:106684 | protein coding | 1.254 | 1.82E-05 | n | n | n | n | n | n |
| Gabrb3 | GABRB3, gamma-aminobuty | 14402 | MGI:95621 | protein coding | 1.243 | 3.06E-07 | n | n | y | n | n | n |
| Dhrs3 | dehydrogenase/reductase 3 | 20148 | MGI:1315215 | protein coding | 1.242 | 2.76E-07 | n | n | n | n | n | n |
| Fndc1 | fibronectin type III domain cor | 68655 | MGI:1915905 | protein coding | 1.241 | 7.03E-06 | n | n | n | n | n | n |
| Pcp4l1 | Purkinje cell protein 4-like 1 | 66425 | MGI:1913675 | protein coding | 1.225 | 1.24E-08 | n | n | n | y | n | n |
| Sting1 | stimulator of interferon respor | 72512 | MGI:1919762 | protein coding | 1.221 | 4.05E-06 | n | n | y | y | n | n |
| Tmem191 | transmembrane protein 191 | 224019 | MGI:107238 | protein coding | 1.221 | 3.30E-09 | n | n | y | n | n | n |
| Unc80 | unc-80, NALCN activator | 329178 | MGI:2652882 | protein coding | 1.221 | 4.52E-05 | n | n | n | n | n | n |
| Ofcc1 | orofacial cleft 1 candidate 1 | 218165 | MGI:2658851 | protein coding | 1.220 | 2.57E-06 | n | n | n | n | n | n |
| Lrrc34 | leucine rich repeat containing | 71827 | MGI:1919077 | protein coding | 1.213 | 0.000559102 | n | n | n | n | n | n |
| Donson | downstream neighbor of SON | 60364 | MGI:1890621 | protein coding | 1.209 | 6.21E-08 | n | y | n | n | n | n |
| Fn3k | fructosamine 3 kinase | 63828 | MGI:1926834 | protein coding | 1.206 | 0.000261705 | n | n | n | n | n | n |
| Pnma8b | PNMA family member 8B | 434128 | MGI:3645856 | protein coding | 1.205 | 1.18E-06 | n | n | n | n | n | n |
| Trim17 | tripartite motif-containing 17 | 56631 | MGI:1861440 | protein coding | 1.204 | 0.000208985 | n | n | n | n | n | n |
| Stk33 | serine/threonine kinase 33 | 117229 | MGI:2152419 | protein coding | 1.200 | 0.000162764 | n | n | n | y | n | n |
| Gm6934 | predicted gene 6934 | 628919 | MGI:3648115 | protein coding | 1.168 | 0.000245659 | n | n | n | n | n | n |
| Dok3 | docking protein 3 | 27261 | MGI:1351490 | protein coding | 1.160 | 0.000201706 | n | n | n | n | n | n |
| Saxo4 | stabilizer of axonemal microtu | 67752 | MGI:1915002 | protein coding | 1.159 | 1.51E-06 | y | n | n | n | n | y |
| Hrob | homologous recombination fa | 217216 | MGI:2387601 | protein coding | 1.158 | 1.26E-07 | n | n | y | n | n | n |
| Gm20939 | predicted gene, 20939 | 100044193 | MGI:5434295 | protein coding | 1.156 | 0.000323549 | n | n | n | n | n | n |

| | | | | | | | | | | | | |
|---|---|---|---|---|---|---|---|---|---|---|---|---|
| Parvb | parvin, beta | 170736 | MGI:2153063 | protein coding | 1.153 | 1.63E-05 | n | n | n | y | n | n |
| Chrng | cholinergic receptor, nicotinic | 11449 | MGI:87895 | protein coding | 1.153 | 4.21E-05 | y | n | n | n | n | n |
| Coa4 | cytochrome c oxidase asseml | 68185 | MGI:1915435 | protein coding | 1.150 | 0.00076909 | n | n | n | n | n | n |
| Tnfrsf18 | tumor necrosis factor recepto | 21936 | MGI:894675 | protein coding | 1.148 | 6.11E-05 | n | n | n | y | n | n |
| Nnt | nicotinamide nucleotide trans | 18115 | MGI:109279 | protein coding | 1.146 | 2.26E-08 | n | n | n | y | n | n |
| Pitpnm1 | phosphatidylinositol transfer p | 18739 | MGI:1197524 | protein coding | 1.142 | 3.09E-09 | y | n | n | n | n | n |
| Kcnh7 | potassium voltage-gated char | 170738 | MGI:2159566 | protein coding | 1.139 | 1.98E-05 | n | n | y | y | n | n |
| Oxa1l | oxidase assembly 1-like | 69089 | MGI:1916339 | protein coding | 1.133 | 1.01E-09 | n | n | n | n | n | n |
| Tmx4 | thioredoxin-related transmem | 52837 | MGI:106558 | protein coding | 1.126 | 6.50E-09 | n | n | n | n | n | n |
| Pim1 | proviral integration site 1 | 18712 | MGI:97584 | protein coding | 1.118 | 2.11E-05 | n | n | n | y | n | n |
| D630023F18Rik | RIKEN cDNA D630023F18 ge | 98303 | MGI:2138198 | protein coding | 1.112 | 6.33E-06 | n | n | n | n | n | n |
| Tspan2 | tetraspanin 2 | 70747 | MGI:1917997 | protein coding | 1.109 | 2.49E-07 | n | n | n | n | n | n |
| D930020B18Rik | RIKEN cDNA D930020B18 ge | 216393 | MGI:2442001 | protein coding | 1.108 | 8.37E-05 | n | n | n | n | n | n |
| Tmem108 | transmembrane protein 108 | 81907 | MGI:1932411 | protein coding | 1.107 | 1.95E-05 | n | n | n | n | n | n |
| 1700088E04Rik | RIKEN cDNA 1700088E04 ge | 27660 | MGI:1920774 | protein coding | 1.102 | 4.07E-05 | n | n | y | n | n | n |
| Ermardl2 | ER membrane associated RN | 100041574 | MGI:3583895 | protein coding | 1.097 | 0.000119327 | n | n | n | n | n | n |
| Crebl2 | cAMP responsive element bir | 232430 | MGI:1889385 | protein coding | 1.093 | 1.17E-06 | n | n | n | y | n | n |
| Dnali1 | dynein, axonemal, light interm | 75563 | MGI:1922813 | protein coding | 1.089 | 0.000123609 | n | n | n | n | n | y |
| Syt14 | synaptotagmin XIV | 329324 | MGI:2444490 | protein coding | 1.088 | 0.000114034 | n | n | y | y | n | n |
| Cd163 | CD163 antigen | 93671 | MGI:2135946 | protein coding | 1.084 | 0.000505711 | n | n | n | n | n | n |
| Qrfpr | pyroglutamylated RFamide pe | 229214 | MGI:2677633 | protein coding | 1.080 | 4.31E-05 | n | n | n | n | n | y |
| Zswim3 | zinc finger SWIM-type contair | 67538 | MGI:1914788 | protein coding | 1.071 | 0.000197819 | n | n | n | n | n | n |
| Smim5 | small integral membrane prot | 66528 | MGI:1913778 | protein coding | 1.064 | 6.26E-07 | n | n | y | n | n | n |
| Cfap91 | cilia and flagella associated p | 320214 | MGI:2443598 | protein coding | 1.059 | 1.57E-05 | n | n | n | n | n | y |
| Cdh13 | cadherin 13 | 12554 | MGI:99551 | protein coding | 1.058 | 1.44E-05 | n | n | n | n | n | n |
| Chrnb2 | cholinergic receptor nicotinic l | 11444 | MGI:87891 | protein coding | 1.058 | 3.62E-07 | n | n | n | n | n** (PMID: 36305825) | n |
| Zfp128 | zinc finger protein 128 | 243833 | MGI:2389445 | protein coding | 1.052 | 0.000672919 | n | n | n | n | n | n |
| Gm14403 | predicted gene 14403 | 433520 | MGI:3649813 | protein coding | 1.051 | 0.00018791 | n | n | n | n | n | n |
| Lhfpl7 | LHFPL tetraspan subfamily m | 333048 | MGI:2685700 | protein coding | 1.049 | 1.69E-05 | n | n | n | n | n | n |
| Cfap65 | cilia and flagella associated p | 241116 | MGI:2444274 | protein coding | 1.047 | 5.36E-06 | n | n | n | n | n | y |
| Capsl | calcyphosine-like | 75568 | MGI:1922818 | protein coding | 1.045 | 4.34E-06 | n | n | y | n | n | n |
| Bmp3 | bone morphogenetic protein 3 | 110075 | MGI:88179 | protein coding | 1.040 | 0.00044637 | n | n | n | n | n | n |
| Ak5 | adenylate kinase 5 | 229949 | MGI:2677491 | protein coding | 1.040 | 0.000728796 | n | n | n | n | n | n |
| Ehhadh | enoyl-Coenzyme A, hydratase | 74147 | MGI:1277964 | protein coding | 1.039 | 1.18E-07 | n | n | n | n | n | n |
| Zfp992 | zinc finger protein 992 | 433791 | MGI:3700963 | protein coding | 1.038 | 6.25E-05 | n | n | n | n | n | n |
| Drc7 | dynein regulatory complex su | 330830 | MGI:2685616 | protein coding | 1.033 | 0.00032074 | n | n | n | n | n | y |
| Hebp1 | heme binding protein 1 | 15199 | MGI:1333880 | protein coding | 1.027 | 2.69E-07 | n | n | n | n | n | n |
| Ccdc96 | coiled-coil domain containing | 66717 | MGI:1913967 | protein coding | 1.025 | 0.000259122 | n | n | n | n | n | y |
| Padi2 | peptidyl arginine deiminase, t | 18600 | MGI:1338892 | protein coding | 1.020 | 1.61E-08 | n | n | n | n | n | n |
| Cdkn2d | cyclin dependent kinase inhib | 12581 | MGI:105387 | protein coding | 1.018 | 1.93E-09 | n | n | y | n | y | n |
| Fgf15 | fibroblast growth factor 15 | 14170 | MGI:1096383 | protein coding | 1.018 | 0.000261705 | n | n | n | n | n | n |
| Trmt1l | tRNA methyltransferase 1 like | 98685 | MGI:1916185 | protein coding | 1.015 | 0.000586655 | n | n | n | n | n | n |
| Jag1 | jagged 1 | 16449 | MGI:1095416 | protein coding | 1.013 | 0.000131999 | n | n | n | y | y** (PMID: 11259677) | n |
| Slc37a1 | solute carrier family 37 (glyce | 224674 | MGI:2446181 | protein coding | 1.011 | 1.75E-07 | n | n | n | y | n | n |
| Elovl2 | ELOVL fatty acid elongase 2 | 54326 | MGI:1858960 | protein coding | 1.005 | 9.33E-07 | n | n | n | n | n | n |
| Fscn2 | fascin actin-bundling protein 2 | 238021 | MGI:2443337 | protein coding | 1.004 | 1.90E-08 | y | n | n | n | y** (PMID: 20660251) | n |
| Inka2 | inka box actin regulator 2 | 109050 | MGI:1923497 | protein coding | 1.004 | 0.000154963 | n | n | n | y | n | n |
| Pm20d1 | peptidase M20 domain contai | 212933 | MGI:2442939 | protein coding | 0.998 | 0.000379443 | n | n | n | n | n | n |
| Celsr1 | cadherin, EGF LAG seven-pa | 12614 | MGI:1100883 | protein coding | 0.996 | 1.12E-06 | n | n | n | n | n | n |
| Zfp691 | zinc finger protein 691 | 195522 | MGI:3041163 | protein coding | 0.994 | 1.06E-06 | n | n | n | n | n | n |
| Degs2 | delta 4-desaturase, sphingolip | 70059 | MGI:1917309 | protein coding | 0.992 | 0.000162339 | n | n | n | n | n | n |
| Atf7ip | activating transcription factor | 54343 | MGI:1858965 | protein coding | 0.991 | 0.000113531 | n | n | y | y | n | n |
| Crhr1 | corticotropin releasing hormo | 12921 | MGI:88498 | protein coding | 0.990 | 1.35E-06 | n | n | n | n | n | n |
| Myo3a | myosin IIIA | 667663 | MGI:2183924 | protein coding | 0.987 | 1.17E-06 | n | n | y** (PMID: 12032315) | y | y | n |
| Fkrp | fukutin related protein | 243853 | MGI:2447586 | protein coding | 0.986 | 1.60E-05 | n | n | n | n | n | n |
| Traf3ip1 | TRAF3 interacting protein 1 | 74019 | MGI:1921269 | protein coding | 0.986 | 1.77E-05 | n | n | n | n | n | y |
| Tspan11 | tetraspanin 11 | 68498 | MGI:1915748 | protein coding | 0.984 | 0.000315351 | n | n | n | y | n | n |
| Mau2 | MAU2 sister chromatid cohes | 74549 | MGI:1921799 | protein coding | 0.980 | 2.86E-05 | n | n | n | n | n | n |
| Cdh18 | cadherin 18 | 320865 | MGI:1344366 | protein coding | 0.978 | 8.11E-05 | n | n | n | y | n | n |
| Dok7 | docking protein 7 | 231134 | MGI:3584043 | protein coding | 0.976 | 3.23E-05 | n | n | n | n | n | n |
| Ubiad1 | UbiA prenyltransferase doma | 71707 | MGI:1918957 | protein coding | 0.968 | 2.12E-08 | n | n | n | n | n | n |
| Cntnap3 | contactin associated protein-l | 238680 | MGI:3588199 | protein coding | 0.967 | 0.000844909 | n | n | n | n | n | n |
| Zfp174 | zinc finger protein 174 | 385674 | MGI:2686600 | protein coding | 0.967 | 0.000158125 | n | n | n | n | n | n |
| Elfn1 | leucine rich repeat and fibron | 243312 | MGI:2442479 | protein coding | 0.966 | 4.67E-06 | n | y | n | y | n | n |
| Oprk1 | opioid receptor, kappa 1 | 18387 | MGI:97439 | protein coding | 0.959 | 1.24E-06 | n | n | n | n | n | n |
| Rsph1 | radial spoke head 1 homolog | 22092 | MGI:1194909 | protein coding | 0.956 | 5.80E-07 | n | n | y | n | n | y |
| Klhdc1 | kelch domain containing 1 | 271005 | MGI:2672853 | protein coding | 0.953 | 0.000399764 | n | n | n | n | n | n |

| | | | | | | | | | | | | |
|---|---|---|---|---|---|---|---|---|---|---|---|---|
| Disp2 | dispatched RND transporter f | 214240 | MGI:2388733 | protein coding | 0.951 | 0.000836046 | n | n | n | n | n | n |
| Rab2b | RAB2B, member RAS oncog | 76338 | MGI:1923588 | protein coding | 0.950 | 1.30E-07 | n | n | n | n | n | n |
| Dpp10 | dipeptidylpeptidase 10 | 269109 | MGI:2442409 | protein coding | 0.949 | 1.43E-06 | n | n | n | y | n | n |
| Stac | src homology three (SH3) and | 20840 | MGI:1201400 | protein coding | 0.945 | 0.0001922 | n | n | n | n | n | n |
| Ugt2a3 | UDP glucuronosyltransferase | 72094 | MGI:1919344 | protein coding | 0.945 | 9.86E-06 | n | n | n | n | n | n |
| Ccdc85a | coiled-coil domain containing | 216613 | MGI:2445069 | protein coding | 0.939 | 1.36E-05 | n | n | n | n | n | n |
| Slc16a3 | solute carrier family 16 (monc | 80879 | MGI:1933438 | protein coding | 0.934 | 0.000103599 | n | n | n | n | n | n |
| Efna5 | ephrin A5 | 13640 | MGI:107444 | protein coding | 0.932 | 5.45E-06 | n | y | n | y | n | n |
| Cngb1 | cyclic nucleotide gated chann | 333329 | MGI:2664102 | protein coding | 0.930 | 8.97E-05 | n | n | n | n | n | y |
| Crtam | cytotoxic and regulatory T cel | 54698 | MGI:1859822 | protein coding | 0.929 | 1.21E-05 | n | n | n | n | n | n |
| Hes5 | hes family bHLH transcription | 15208 | MGI:104876 | protein coding | 0.925 | 0.000733926 | n | n | n | n | n | n |
| Ebf2 | early B cell factor 2 | 13592 | MGI:894332 | protein coding | 0.922 | 3.04E-07 | n | n | n | y | n | n |
| Nos1ap | nitric oxide synthase 1 (neuro | 70729 | MGI:1917979 | protein coding | 0.920 | 7.89E-06 | n | y | n | n | n | n |
| Trim36 | tripartite motif-containing 36 | 28105 | MGI:106264 | protein coding | 0.917 | 0.000425409 | y | n | n | n | n | n |
| Slc25a25 | solute carrier family 25 (mitoc | 227731 | MGI:1915913 | protein coding | 0.916 | 1.38E-07 | n | n | n | y | n | n |
| Zfp719 | zinc finger protein 719 | 210105 | MGI:2444708 | protein coding | 0.913 | 3.48E-06 | n | n | n | n | n | n |
| Myo1h | myosin 1H | 231646 | MGI:1914674 | protein coding | 0.905 | 6.21E-06 | n | y | n | n | n | n |
| Iqcc | IQ motif containing C | 230767 | MGI:2446212 | protein coding | 0.902 | 7.99E-06 | n | n | n | n | n | n |
| Rlig1 | RNA 5'-phosphate and 3'-OH | 68281 | MGI:1921197 | protein coding | 0.898 | 1.58E-05 | n | n | y | n | n | n |
| Bicdl2 | BICD family like cargo adaptc | 212733 | MGI:2388267 | protein coding | 0.898 | 0.000856265 | n | n | n | n | n | n |
| Dmtn | dematin actin binding protein | 13829 | MGI:99670 | protein coding | 0.897 | 1.57E-07 | n | n | n | n | n | n |
| Acp3 | acid phosphatase 3 | 56318 | MGI:1928480 | protein coding | 0.889 | 6.58E-05 | n | n | n | n | n | n |
| Emc9 | ER membrane protein comple | 85308 | MGI:1934682 | protein coding | 0.887 | 0.000155825 | n | n | n | n | n | n |
| Cacng5 | calcium channel, voltage-dep | 140723 | MGI:2157946 | protein coding | 0.887 | 2.60E-05 | n | y | n | y | n | n |
| Cfap43 | cilia and flagella associated p | 100048534 | MGI:1289258 | protein coding | 0.885 | 0.000487331 | n | n | n | n | n | y |
| Pdgfrl | platelet-derived growth factor | 68797 | MGI:1916047 | protein coding | 0.879 | 8.44E-07 | n | n | n | n | n | n |
| Gpr179 | G protein-coupled receptor 17 | 217143 | MGI:2443409 | protein coding | 0.879 | 0.000124389 | n | n | n | n | n | n |
| Timeless | timeless circadian clock 1 | 21853 | MGI:1321393 | protein coding | 0.875 | 4.86E-05 | n | n | n | n | n | n |
| Zfp760 | zinc finger protein 760 | 240034 | MGI:2679257 | protein coding | 0.874 | 1.36E-05 | n | n | n | n | n | n |
| Hsph1 | heat shock 105kDa/110kDa p | 15505 | MGI:105053 | protein coding | 0.872 | 3.59E-07 | n | n | y | n | n | n |
| Cimap1b | ciliary microtubule associated | 70113 | MGI:1917363 | protein coding | 0.872 | 0.000111872 | y | n | n | n | n | y |
| Ide | insulin degrading enzyme | 15925 | MGI:96412 | protein coding | 0.871 | 5.03E-08 | n | n | n | n | n | n |
| Asap3 | ArfGAP with SH3 domain, anl | 230837 | MGI:2684986 | protein coding | 0.871 | 1.11E-07 | n | n | n | n | n | n |
| Drc1 | dynein regulatory complex su | 381738 | MGI:2685906 | protein coding | 0.871 | 0.000631194 | n | n | y | n | n | y |
| Pcsk9 | proprotein convertase subtilis | 100102 | MGI:2140260 | protein coding | 0.870 | 1.54E-05 | n | n | y | n | n | n |
| Katnal2 | katanin p60 subunit A-like 2 | 71206 | MGI:1924234 | protein coding | 0.868 | 0.000190991 | n | n | n | n | n | n |
| Foxj1 | forkhead box J1 | 15223 | MGI:1347474 | protein coding | 0.864 | 0.000236499 | n | n | y | y | n | n |
| Fbxo6 | F-box protein 6 | 50762 | MGI:1354743 | protein coding | 0.863 | 1.29E-08 | n | n | n | n | n | n |
| Pgbd1 | piggyBac transposable eleme | 319207 | MGI:2441675 | protein coding | 0.858 | 0.000637384 | n | n | n | n | n | n |
| C1qtnf12 | C1q and tumor necrosis facto | 67389 | MGI:1914639 | protein coding | 0.852 | 2.14E-08 | n | n | n | n | n | n |
| Pcdhac2 | protocadherin alpha subfamily | 353237 | MGI:1891443 | protein coding | 0.852 | 1.13E-05 | n | n | n | n | n | n |
| Dido1 | death inducer-obliterator 1 | 23856 | MGI:1344352 | protein coding | 0.851 | 3.14E-06 | n | n | n | n | n | n |
| Dlg2 | discs large MAGUK scaffold p | 23859 | MGI:1344351 | protein coding | 0.849 | 3.62E-06 | n | n | n | y | n | n |
| Osbpl6 | oxysterol binding protein-like | 99031 | MGI:2139014 | protein coding | 0.849 | 1.04E-07 | n | n | n | n | n | n |
| Cdk18 | cyclin dependent kinase 18 | 18557 | MGI:97518 | protein coding | 0.843 | 1.30E-05 | n | n | n | n | n | n |
| Zcchc12 | zinc finger, CCHC domain co | 72693 | MGI:1919943 | protein coding | 0.840 | 4.56E-07 | n | n | n | n | n | n |
| Myo15a | myosin XVA | 17910 | MGI:1261811 | protein coding | 0.840 | 4.40E-06 | n | n | y | n | y | n |
| Shisal1 | shisa like 1 | 72301 | MGI:1919551 | protein coding | 0.839 | 4.13E-06 | n | n | n | y | n | n |
| Mogat1 | monoacylglycerol O-acyltrans | 68393 | MGI:1915643 | protein coding | 0.838 | 2.71E-05 | n | n | y | y | n | n |
| Spred3 | sprouty-related EVH1 domain | 101809 | MGI:2142186 | protein coding | 0.837 | 0.000716683 | n | n | n | n | n | n |
| Lmbrd2 | LMBR1 domain containing 2 | 320506 | MGI:2444173 | protein coding | 0.833 | 0.000139893 | n | n | n | n | n | n |
| Zmat1 | zinc finger, matrin type 1 | 215693 | MGI:2442284 | protein coding | 0.831 | 0.000475152 | n | n | n | n | n | n |
| Tram2 | translocating chain-associatin | 170829 | MGI:1924817 | protein coding | 0.828 | 0.000785437 | n | n | n | n | n | n |
| Dzank1 | double zinc ribbon and ankyri | 241688 | MGI:2139080 | protein coding | 0.824 | 3.04E-05 | n | n | n | n | n | n |
| Eif3l | eukaryotic translation initiatio | 223691 | MGI:2386251 | protein coding | 0.814 | 4.59E-08 | n | n | n | n | n | n |
| Gtpbp3 | GTP binding protein 3 | 70359 | MGI:1917609 | protein coding | 0.812 | 3.81E-05 | n | n | n | n | n | n |
| Rab37 | RAB37, member RAS oncoge | 58222 | MGI:1929945 | protein coding | 0.812 | 0.000307362 | n | n | n | y | n | n |
| Abcc10 | ATP-binding cassette, sub-fa | 224814 | MGI:2386976 | protein coding | 0.812 | 1.39E-05 | n | n | n | n | n | n |
| Nphp4 | nephronophthisis 4 (juvenile) | 260305 | MGI:2384210 | protein coding | 0.810 | 0.000176768 | n | n | n | y | n | y |
| Heatr5b | HEAT repeat containing 5B | 320473 | MGI:2444098 | protein coding | 0.801 | 0.000976689 | n | n | n | n | n | n |
| Cfap299 | cilia and flagella associated p | 75784 | MGI:1916571 | protein coding | 0.794 | 0.000848434 | n | n | n | n | n | n |
| Ptprq | protein tyrosine phosphatase | 237523 | MGI:1096349 | protein coding | 0.792 | 0.000803483 | n | n | y | n | y** (PMID: 14534255) | n |
| Zscan29 | zinc finger SCAN domains 29 | 99334 | MGI:2139317 | protein coding | 0.787 | 2.76E-07 | n | n | n | n | n | n |
| Dbndd1 | dysbindin domain containing | 72185 | MGI:1919435 | protein coding | 0.787 | 4.90E-06 | n | y | n | n | n | n |
| Acad12 | acyl-Coenzyme A dehydroge | 338350 | MGI:2443320 | protein coding | 0.786 | 0.00060666 | n | n | n | n | n | n |
| Smap2 | small ArfGAP 2 | 69780 | MGI:1917030 | protein coding | 0.775 | 4.23E-08 | n | n | y | n | n | n |
| Slc9a2 | solute carrier family 9 (sodiun | 226999 | MGI:105075 | protein coding | 0.775 | 1.36E-05 | n | n | n | y | n | n |

| | | | | | | | | | | | | |
|---|---|---|---|---|---|---|---|---|---|---|---|---|
| Ubxn10 | UBX domain protein 10 | 212190 | MGI:2443123 | protein coding | 0.771 | 0.000971013 | n | n | n | y | n | y |
| Lmod1 | leiomodin 1 (smooth muscle) | 93689 | MGI:2135671 | protein coding | 0.769 | 9.07E-05 | n | n | n | y | n | n |
| Ss18l1 | SS18, nBAF chromatin remoc | 269397 | MGI:2444061 | protein coding | 0.765 | 2.18E-06 | n | n | n | n | n | n |
| 1700003E16Rik | RIKEN cDNA 1700003E16 ge | 71837 | MGI:1919087 | protein coding | 0.765 | 0.00076909 | n | n | n | n | n | n |
| Zfp180 | zinc finger protein 180 | 210135 | MGI:1923701 | protein coding | 0.765 | 5.25E-06 | n | n | n | n | n | n |
| Zfp747l1 | zinc finger protein 747 like 1 | 78921 | MGI:1926171 | protein coding | 0.764 | 0.000960485 | n | n | n | n | n | n |
| Prmt8 | protein arginine N-methyltran: | 381813 | MGI:3043083 | protein coding | 0.764 | 0.000983416 | n | n | n | n | n | n |
| Plch2 | phospholipase C, eta 2 | 269615 | MGI:2443078 | protein coding | 0.762 | 2.92E-05 | n | n | y | y | n | n |
| Fbxo27 | F-box protein 27 | 233040 | MGI:2685007 | protein coding | 0.760 | 9.86E-06 | n | n | n | y | n | n |
| Rhov | ras homolog family member V | 228543 | MGI:2444227 | protein coding | 0.758 | 0.000577442 | n | n | n | n | n | n |
| Acss3 | acyl-CoA synthetase short-ch | 380660 | MGI:2685720 | protein coding | 0.753 | 0.000154238 | n | n | n | n | n | n |
| Calb1 | calbindin 1 | 12307 | MGI:88248 | protein coding | 0.751 | 2.96E-07 | n | n | y | n | n | n |
| Spock2 | sparc/osteonectin, cwcv and l | 94214 | MGI:1891351 | protein coding | 0.746 | 5.53E-06 | n | n | y | y | n | n |
| Dytn | dystrotelin | 241073 | MGI:2685061 | protein coding | 0.746 | 3.79E-06 | n | y | n | n | n | n |
| Dnaja4 | DnaJ heat shock protein fami | 58233 | MGI:1927638 | protein coding | 0.740 | 1.48E-06 | n | n | y | n | n | n |
| Pih1d2 | PIH1 domain containing 2 | 72614 | MGI:1919864 | protein coding | 0.738 | 0.000113531 | y | n | n | n | n | n |
| Nipal3 | NIPA-like domain containing : | 74552 | MGI:1921802 | protein coding | 0.737 | 2.83E-05 | n | n | y | n | n | n |
| Pigg | phosphatidylinositol glycan ar | 433931 | MGI:3576484 | protein coding | 0.735 | 6.82E-05 | n | n | n | n | n | n |
| Kcnj11 | potassium inwardly rectifying | 16514 | MGI:107501 | protein coding | 0.733 | 0.000114162 | n | n | n | n | n | n |
| Adcy8 | adenylate cyclase 8 | 11514 | MGI:1341110 | protein coding | 0.731 | 0.000557841 | n | n | n | y | n | n |
| Mettl16 | methyltransferase 16, N6-met | 67493 | MGI:1914743 | protein coding | 0.730 | 4.17E-05 | n | n | n | n | n | n |
| Sgpp2 | sphingosine-1-phosphate phc | 433323 | MGI:3589109 | protein coding | 0.730 | 5.36E-06 | n | n | n | y | n | n |
| Fibin | fin bud initiation factor homolc | 67606 | MGI:1914856 | protein coding | 0.729 | 7.02E-05 | n | n | n | n | n | n |
| Gpr152 | G protein-coupled receptor 1! | 269053 | MGI:2685519 | protein coding | 0.728 | 7.51E-05 | n | n | n | n | n | n |
| Capn15 | calpain 15 | 50817 | MGI:1355075 | protein coding | 0.726 | 0.000836046 | n | n | n | n | n | n |
| Cdkl4 | cyclin dependent kinase like 4 | 381113 | MGI:3587025 | protein coding | 0.715 | 4.56E-07 | n | n | y | n | n | n |
| B3gnt4 | UDP-GlcNAc:betaGal beta-1, | 231727 | MGI:2680208 | protein coding | 0.713 | 0.000116742 | n | n | y | n | n | n |
| Nudt5 | nudix hydrolase 5 | 53893 | MGI:1858232 | protein coding | 0.713 | 3.69E-05 | n | n | n | n | n | n |
| Hdhd5 | haloacid dehalogenase like hy | 214932 | MGI:2136976 | protein coding | 0.713 | 6.33E-06 | n | n | n | n | n | n |
| Tubgcp4 | tubulin, gamma complex com | 51885 | MGI:1196293 | protein coding | 0.713 | 8.33E-06 | n | n | n | n | n | n |
| Mlh3 | mutL homolog 3 | 217716 | MGI:1353455 | protein coding | 0.712 | 3.03E-05 | n | n | n | n | n | n |
| Cyb5d2 | cytochrome b5 domain contai | 192986 | MGI:2684848 | protein coding | 0.712 | 0.000139658 | n | n | n | n | n | n |
| Gfi1 | growth factor independent 1 t | 14581 | MGI:103170 | protein coding | 0.710 | 3.78E-07 | n | n | y | y | y** (PMID: 12441305) | n |
| Snai3 | snail family zinc finger 3 | 30927 | MGI:1353563 | protein coding | 0.706 | 0.000278487 | n | n | n | y | n | n |
| Aass | aminoadipate-semialdehyde s | 30956 | MGI:1353573 | protein coding | 0.704 | 0.000660496 | n | n | n | n | n | n |
| Sesn2 | sestrin 2 | 230784 | MGI:2651874 | protein coding | 0.704 | 2.53E-06 | n | n | n | n | n | n |
| Aqp4 | aquaporin 4 | 11829 | MGI:107387 | protein coding | 0.703 | 1.59E-05 | n | n | n | n | y | n |
| Dffa | DNA fragmentation factor, alp | 13347 | MGI:1196227 | protein coding | 0.702 | 0.000928837 | n | n | n | n | n | n |
| Zfp866 | zinc finger protein 866 | 330788 | MGI:3584369 | protein coding | 0.698 | 0.000151377 | n | n | n | n | n | n |
| Ak1 | adenylate kinase 1 | 11636 | MGI:87977 | protein coding | 0.697 | 9.86E-06 | n | n | y | y | n | y |
| Fam89a | family with sequence similarit | 69627 | MGI:1916877 | protein coding | 0.697 | 0.000340806 | n | n | n | n | n | n |
| Uhmk1 | U2AF homology motif (UHM) | 16589 | MGI:1341908 | protein coding | 0.696 | 0.000607188 | n | n | y | n | n | n |
| Slc25a42 | solute carrier family 25, meml | 73095 | MGI:1920345 | protein coding | 0.696 | 0.000673666 | n | n | n | n | n | n |
| P4ha2 | procollagen-proline, 2-oxoglut | 18452 | MGI:894286 | protein coding | 0.696 | 5.41E-08 | n | n | n | n | n | n |
| Kncn | kinocilin | 654462 | MGI:3614952 | protein coding | 0.696 | 1.14E-05 | n | n | y | n | n | y |
| Pierce2 | piercer of microtubule wall 2 | 546143 | MGI:3648770 | protein coding | 0.693 | 0.0001922 | n | n | y | n | n | y |
| Ppara | peroxisome proliferator activa | 19013 | MGI:104740 | protein coding | 0.693 | 0.000408468 | n | n | n | n | n | n |
| Cygb | cytoglobin | 114886 | MGI:2149481 | protein coding | 0.691 | 0.000970555 | n | n | n | n | n | n |
| Sv2b | synaptic vesicle glycoprotein | 64176 | MGI:1927338 | protein coding | 0.690 | 3.05E-05 | n | n | n | n | n | n |
| Veph1 | ventricular zone expressed PI | 72789 | MGI:1920039 | protein coding | 0.689 | 1.49E-06 | n | y | n | y | n | n |
| Mex3b | mex3 RNA binding family mer | 108797 | MGI:1918252 | protein coding | 0.686 | 0.000735533 | n | n | n | y | n | n |
| Syt13 | synaptotagmin XIII | 80976 | MGI:1933945 | protein coding | 0.686 | 1.95E-05 | n | n | y | y | n | n |
| Micall1 | microtubule associated mono | 27008 | MGI:105870 | protein coding | 0.682 | 2.94E-06 | n | n | n | n | n | n |
| Adgrv1 | adhesion G protein-coupled re | 110789 | MGI:1274784 | protein coding | 0.681 | 0.000110178 | n | n | y | y | y | y |
| Cercam | cerebral endothelial cell adhe | 99151 | MGI:2139134 | protein coding | 0.679 | 3.94E-07 | n | n | n | n | n | n |
| Zfyve16 | zinc finger, FYVE domain cor | 218441 | MGI:2145181 | protein coding | 0.677 | 0.000107051 | n | n | n | n | n | n |
| Cnksr2 | connector enhancer of kinase | 245684 | MGI:2661175 | protein coding | 0.675 | 0.000351654 | n | n | n | n | n | n |
| Dpysl5 | dihydropyrimidinase-like 5 | 65254 | MGI:1929772 | protein coding | 0.675 | 0.000505711 | n | n | n | y | n | n |
| Aifm1 | apoptosis-inducing factor, mit | 26926 | MGI:1349419 | protein coding | 0.673 | 6.09E-08 | n | n | n | n | n | n |
| Nebl | nebulette | 74103 | MGI:1921353 | protein coding | 0.667 | 1.23E-05 | n | n | n | y | n | n |
| Tmem51 | transmembrane protein 51 | 214359 | MGI:2384874 | protein coding | 0.666 | 1.69E-06 | n | n | n | n | n | n |
| Hspa2 | heat shock protein 2 | 15512 | MGI:96243 | protein coding | 0.666 | 3.61E-06 | n | n | n | y | n | n |
| Pccb | propionyl Coenzyme A carbo: | 66904 | MGI:1914154 | protein coding | 0.665 | 1.75E-07 | n | n | n | y | n | n |
| Evc2 | EvC ciliary complex subunit 2 | 68525 | MGI:1915775 | protein coding | 0.665 | 6.46E-06 | n | n | y | y | n | y |
| Chn2 | chimerin 2 | 69993 | MGI:1917243 | protein coding | 0.663 | 0.000253038 | n | n | n | n | n | n |
| Prrt3 | proline-rich transmembrane p | 210673 | MGI:2444810 | protein coding | 0.662 | 0.000308484 | n | n | n | y | n | n |
| Pnpla3 | patatin-like phospholipase do | 116939 | MGI:2151796 | protein coding | 0.662 | 0.000630585 | y | n | n | n | n | n |

| | | | | | | | | | | | | |
|---|---|---|---|---|---|---|---|---|---|---|---|---|
| Sgtb | small glutamine-rich tetratrico | 218544 | MGI:2444615 | protein coding | 0.661 | 0.000636483 | n | n | n | n | n | n |
| Prkg2 | protein kinase, cGMP-depenc | 19092 | MGI:108173 | protein coding | 0.659 | 6.46E-05 | n | n | n | n | n | n |
| Ap3m2 | adaptor-related protein compl | 64933 | MGI:1929214 | protein coding | 0.659 | 1.72E-07 | n | n | y | y | n | n |
| Zfp983 | zinc finger protein 983 | 73229 | MGI:1920479 | protein coding | 0.654 | 0.000805895 | n | n | n | n | n | n |
| Stk35 | serine/threonine kinase 35 | 67333 | MGI:1914583 | protein coding | 0.653 | 0.000388666 | n | n | n | n | n | n |
| Amacr | alpha-methylacyl-CoA racema | 17117 | MGI:1098273 | protein coding | 0.652 | 0.000208936 | n | n | n | n | n | n |
| Polr2f | polymerase (RNA) II (DNA di | 69833 | MGI:1349393 | protein coding | 0.650 | 9.15E-06 | n | n | n | n | n | n |
| Spef2 | sperm flagellar 2 | 320277 | MGI:2443727 | protein coding | 0.647 | 0.000637521 | n | n | n | n | n | y |
| Cers6 | ceramide synthase 6 | 241447 | MGI:2442564 | protein coding | 0.641 | 0.000487148 | n | n | n | n | n | n |
| Raph1 | Ras association (RalGDS/AF- | 77300 | MGI:1924550 | protein coding | 0.639 | 1.16E-05 | n | n | n | y | n | n |
| Ppp2r5b | protein phosphatase 2, regula | 225849 | MGI:2388480 | protein coding | 0.638 | 8.70E-05 | n | n | y | n | n | n |
| Atoh1 | atonal bHLH transcription fact | 11921 | MGI:104654 | protein coding | 0.638 | 0.000889297 | n | n | y | y | y** (PMID: 10364557) | n |
| Gm1322 | predicted gene 1322 | 383709 | MGI:2686168 | protein coding | 0.637 | 0.000259226 | n | n | n | n | n | n |
| Spag6l | sperm associated antigen 6-li | 50525 | MGI:1354388 | protein coding | 0.633 | 0.000228071 | n | n | y | n | n | y |
| Ablim1 | actin-binding LIM protein 1 | 226251 | MGI:1194500 | protein coding | 0.626 | 1.77E-05 | n | n | n | y | n | n |
| Ccdc68 | coiled-coil domain containing | 381175 | MGI:3612676 | protein coding | 0.624 | 1.36E-05 | n | n | n | y | n | y |
| Tekt1 | tektin 1 | 21689 | MGI:1333819 | protein coding | 0.622 | 7.14E-07 | n | n | n | y | n | y |
| Tsen15 | tRNA splicing endonuclease s | 66637 | MGI:1913887 | protein coding | 0.621 | 0.000666466 | n | n | n | n | n | n |
| Pik3cd | phosphatidylinositol-4,5-bisph | 18707 | MGI:1098211 | protein coding | 0.615 | 0.000639689 | n | n | n | n | n | n |
| 1600014C10Rik | RIKEN cDNA 1600014C10 ge | 72244 | MGI:1919494 | protein coding | 0.615 | 0.000779987 | n | n | n | y | n | n |
| B3galt2 | UDP-Gal:betaGlcNAc beta 1, | 26878 | MGI:1349461 | protein coding | 0.613 | 0.000705579 | n | n | n | y | n | n |
| Abcd2 | ATP-binding cassette, sub-fa | 26874 | MGI:1349467 | protein coding | 0.613 | 2.94E-05 | n | n | n | y | n | n |
| Gemin4 | gem nuclear organelle associ | 276919 | MGI:2449313 | protein coding | 0.612 | 0.000925473 | n | n | n | n | n | n |
| Fanca | Fanconi anemia, complement | 14087 | MGI:1341823 | protein coding | 0.609 | 4.23E-05 | n | y | n | n | n | n |
| Pjvk | pejvakin | 381375 | MGI:2685847 | protein coding | 0.608 | 0.000785437 | n | n | n | n | y | y |
| Apobec1 | apolipoprotein B mRNA editin | 11810 | MGI:103298 | protein coding | 0.608 | 0.000604096 | n | n | n | n | n | n |
| Dzip3 | DAZ interacting protein 3, zin | 224170 | MGI:1917433 | protein coding | 0.604 | 0.0005203 | n | n | n | n | n | n |
| Mapkbp1 | mitogen-activated protein kina | 26390 | MGI:1347004 | protein coding | 0.604 | 2.09E-05 | n | y | n | n | n | n |
| Gm7694 | predicted gene 7694 | 665574 | MGI:3649135 | protein coding | 0.603 | 0.000463599 | n | n | n | n | n | n |
| Parp11 | poly (ADP-ribose) polymerase | 101187 | MGI:2141505 | protein coding | 0.602 | 0.000416544 | n | n | n | n | n | n |
| Gpx2 | glutathione peroxidase 2 | 14776 | MGI:106609 | protein coding | 0.601 | 1.54E-05 | n | n | n | y | n | n |
| Tmtc4 | transmembrane and tetratrico | 70551 | MGI:1921050 | protein coding | 0.597 | 2.90E-06 | n | y | n | y | y | n |
| Far1 | fatty acyl CoA reductase 1 | 67420 | MGI:1914670 | protein coding | 0.596 | 0.00024129 | n | n | n | n | n | n |
| Cacnb4 | calcium channel, voltage-dep | 12298 | MGI:103301 | protein coding | 0.596 | 0.000832284 | n | n | n | n | n | n |
| Zfp672 | zinc finger protein 672 | 319475 | MGI:2442105 | protein coding | 0.592 | 0.000360073 | n | n | n | n | n | n |
| Mfng | MFNG O-fucosylpeptide 3-be | 17305 | MGI:1095404 | protein coding | 0.592 | 5.99E-07 | n | n | y | y | n | n |
| Atg16l2 | autophagy related 16 like 2 | 73683 | MGI:1920933 | protein coding | 0.590 | 0.000551603 | n | n | n | y | n | n |
| Aph1c | aph1 homolog C, gamma sec | 68318 | MGI:1915568 | protein coding | 0.590 | 0.000110892 | n | n | n | n | n | n |
| Pigv | phosphatidylinositol glycan ar | 230801 | MGI:2442480 | protein coding | 0.589 | 9.98E-06 | n | n | n | n | n | n |
| Sfxn2 | sideroflexin 2 | 94279 | MGI:2137678 | protein coding | 0.589 | 0.000848434 | n | n | n | n | n | n |
| Arhgef18 | Rho/Rac guanine nucleotide | 102098 | MGI:2142567 | protein coding | 0.588 | 0.000446777 | n | n | n | n | n | n |
| Ubash3b | ubiquitin associated and SH3 | 72828 | MGI:1920078 | protein coding | 0.585 | 6.87E-06 | n | n | n | n | n | n |
| Myo18a | myosin XVIIIA | 360013 | MGI:2667185 | protein coding | 0.585 | 6.21E-06 | n | n | y | y | n | n |
| Ptprg | protein tyrosine phosphatase | 19270 | MGI:97814 | protein coding | 0.585 | 9.29E-05 | n | n | n | y | n | n |
| Eif3j2 | eukaryotic translation initiatio | 100042807 | MGI:3704486 | protein coding | 0.583 | 0.000399708 | n | n | n | n | n | n |
| Apba3 | amyloid beta precursor protei | 57267 | MGI:1888527 | protein coding | 0.582 | 0.000214896 | n | n | n | n | n | n |
| Tjp3 | tight junction protein 3 | 27375 | MGI:1351650 | protein coding | 0.581 | 0.000141622 | n | n | n | n | n | n |
| Marchf8 | membrane associated ring-Cl | 71779 | MGI:1919029 | protein coding | 0.580 | 6.77E-06 | n | n | n | y | n | n |
| Rdh12 | retinol dehydrogenase 12 | 77974 | MGI:1925224 | protein coding | 0.580 | 6.28E-06 | n | n | y | n | n | n |
| Kdm5c | lysine demethylase 5C | 20591 | MGI:99781 | protein coding | 0.579 | 5.14E-06 | n | n | n | n | n | n |
| Fndc7 | fibronectin type III domain cor | 320181 | MGI:2443535 | protein coding | 0.578 | 1.10E-05 | n | n | n | n | n | n |
| Sall2 | spalt like transcription factor 2 | 50524 | MGI:1354373 | protein coding | 0.578 | 1.23E-06 | n | n | n | n | n | n |
| Rtn4ip1 | reticulon 4 interacting protein | 170728 | MGI:2178759 | protein coding | 0.577 | 4.99E-05 | n | n | n | n | n | n |
| Odad2 | outer dynein arm docking con | 74934 | MGI:1922184 | protein coding | 0.576 | 0.000705585 | y | n | n | n | n | y |
| S100pbp | S100P binding protein | 74648 | MGI:1921898 | protein coding | 0.575 | 0.000724984 | n | n | n | n | n | n |
| Ivns1abp | influenza virus NS1A binding | 117198 | MGI:2152389 | protein coding | 0.574 | 1.81E-05 | n | n | n | y | n | n |
| Magee1 | MAGE family member E1 | 107528 | MGI:2148149 | protein coding | 0.573 | 9.90E-06 | n | n | n | n | n | n |
| Mmaa | methylmalonic aciduria (coba | 109136 | MGI:1923805 | protein coding | 0.572 | 0.000126689 | n | n | n | n | n | n |
| Acad11 | acyl-Coenzyme A dehydroge | 102632 | MGI:2143169 | protein coding | 0.571 | 4.46E-05 | n | y | n | n | n | n |
| Lrrc61 | leucine rich repeat containing | 243371 | MGI:2652848 | protein coding | 0.570 | 3.04E-05 | n | n | n | n | n | n |
| Nim1k | NIM1 serine/threonine protein | 245269 | MGI:2442399 | protein coding | 0.568 | 0.000449066 | n | y | n | y | n | n |
| Nck1 | non-catalytic region of tyrosin | 17973 | MGI:109601 | protein coding | 0.566 | 0.000205349 | n | n | n | n | n | n |
| Xirp2 | xin actin-binding repeat conta | 241431 | MGI:2685198 | protein coding | 0.562 | 4.99E-05 | y | n | n | n | y** (PMID: 25772365, 25653358) | n |
| Rb1 | RB transcriptional corepresso | 19645 | MGI:97874 | protein coding | 0.558 | 2.60E-05 | n | n | n | y | n | n |
| Snapc1 | small nuclear RNA activating | 75627 | MGI:1922877 | protein coding | 0.558 | 0.00010225 | n | n | n | n | n | n |
| Twf2 | twinfilin actin binding protein 2 | 23999 | MGI:1346078 | protein coding | 0.556 | 5.34E-06 | n | n | y | n | n | n |
| Pex10 | peroxisomal biogenesis factor | 668173 | MGI:2684988 | protein coding | 0.552 | 0.000309585 | n | n | n | n | n | n |

| | | | | | | | | | | | | |
|---|---|---|---|---|---|---|---|---|---|---|---|---|
| Fndc3a | fibronectin type III domain cor | 319448 | MGI:1196463 | protein coding | 0.552 | 1.19E-05 | n | n | n | n | n | n |
| Dusp14 | dual specificity phosphatase 1 | 56405 | MGI:1927168 | protein coding | 0.551 | 3.37E-05 | n | n | y | y | n | n |
| Ascc2 | activating signal cointegrator | 75452 | MGI:1922702 | protein coding | 0.551 | 0.000235779 | n | n | n | n | n | n |
| Slc35b4 | solute carrier family 35, meml | 58246 | MGI:1931249 | protein coding | 0.549 | 0.000764309 | n | n | n | y | n | n |
| Fut8 | fucosyltransferase 8 | 53618 | MGI:1858901 | protein coding | 0.547 | 3.02E-06 | n | n | n | n | n | n |
| Ecel1 | endothelin converting enzyme | 13599 | MGI:1343461 | protein coding | 0.547 | 0.000189155 | n | n | n | y | n | n |
| Car12 | carbonic anhydrase 12 | 76459 | MGI:1923709 | protein coding | 0.544 | 0.000791453 | n | y | n | n | n | n |
| Atp2c1 | ATPase, Ca++-sequestering | 235574 | MGI:1889008 | protein coding | 0.543 | 3.00E-07 | n | n | n | y | n | n |
| Fadd | Fas associated via death dom | 14082 | MGI:109324 | protein coding | 0.543 | 0.000166832 | n | n | n | n | n | n |
| Gpr155 | G protein-coupled receptor 15 | 68526 | MGI:1915776 | protein coding | 0.542 | 0.000703177 | n | n | y | n | n | n |
| Cog6 | component of oligomeric golg | 67542 | MGI:1914792 | protein coding | 0.540 | 0.000235336 | n | n | n | n | n | n |
| Lnpk | lunapark, ER junction formatio | 69605 | MGI:1918115 | protein coding | 0.539 | 0.000231536 | n | n | n | n | n | n |
| Map3k12 | mitogen-activated protein kina | 26404 | MGI:1346881 | protein coding | 0.538 | 0.000428722 | n | n | n | n | n | n |
| Proser3 | proline and serine rich 3 | 333193 | MGI:2681861 | protein coding | 0.537 | 0.000643881 | n | n | n | n | n | n |
| Phf8 | PHD finger protein 8 | 320595 | MGI:2444341 | protein coding | 0.536 | 0.000471516 | n | n | n | n | n | n |
| Cnot3 | CCR4-NOT transcription com | 232791 | MGI:2385261 | protein coding | 0.534 | 0.000719821 | n | n | n | y | n | n |
| Vars2 | valyl-tRNA synthetase 2, mito | 68915 | MGI:1916165 | protein coding | 0.534 | 9.58E-05 | n | n | n | n | n | n |
| Ing5 | inhibitor of growth family, mer | 66262 | MGI:1922816 | protein coding | 0.532 | 0.000274799 | n | n | n | n | n | n |
| Slc39a3 | solute carrier family 39 (zinc t | 106947 | MGI:2147269 | protein coding | 0.532 | 3.96E-05 | n | n | y | n | n | n |
| Kndc1 | kinase non-catalytic C-lobe do | 76484 | MGI:1923734 | protein coding | 0.531 | 0.000236087 | n | y | n | n | n | n |
| Tyk2 | tyrosine kinase 2 | 54721 | MGI:1929470 | protein coding | 0.530 | 0.000502137 | n | n | n | n | n | n |
| Tmem107 | transmembrane protein 107 | 66910 | MGI:1914160 | protein coding | 0.530 | 0.000498 | n | n | y | n | n | y |
| Atg4c | autophagy related 4C, cystein | 242557 | MGI:2651854 | protein coding | 0.528 | 0.000565443 | n | n | n | y | n | n |
| Gfm2 | G elongation factor, mitochon | 320806 | MGI:2444783 | protein coding | 0.526 | 1.39E-05 | n | n | n | n | n | n |
| Btaf1 | B-TFIID TATA-box binding pr | 107182 | MGI:2147538 | protein coding | 0.525 | 0.000463599 | n | n | n | n | n | n |
| Aplp1 | amyloid beta precursor like pr | 11803 | MGI:88046 | protein coding | 0.524 | 2.69E-07 | n | n | n | n | n | n |
| Snap23 | synaptosomal-associated pro | 20619 | MGI:109356 | protein coding | 0.524 | 0.000177378 | n | n | n | n | n | n |
| Hspa4l | heat shock protein 4 like | 18415 | MGI:107422 | protein coding | 0.524 | 3.05E-05 | n | n | y | y | n | n |
| Fhit | fragile histidine triad gene | 14198 | MGI:1277947 | protein coding | 0.522 | 5.77E-05 | n | n | y | y | n | n |
| Marveld3 | MARVEL (membrane-associa | 73608 | MGI:1920858 | protein coding | 0.522 | 0.00060529 | n | n | n | y | n | n |
| Armcx1 | armadillo repeat containing, X | 78248 | MGI:1925498 | protein coding | 0.521 | 0.000136239 | n | n | n | n | n | n |
| Cap2 | cyclase associated actin cyto | 67252 | MGI:1914502 | protein coding | 0.515 | 7.02E-05 | n | n | n | y | n | n |
| Cdkl2 | cyclin dependent kinase like 2 | 53886 | MGI:1858227 | protein coding | 0.513 | 6.34E-05 | n | n | y | n | n | n |
| Tdrd3 | tudor domain containing 3 | 219249 | MGI:2444023 | protein coding | 0.512 | 0.000643881 | n | n | n | y | n | n |
| Slc4a2 | solute carrier family 4 (anion e | 20535 | MGI:109351 | protein coding | 0.512 | 9.03E-06 | n | n | n | n | n | n |
| Rexo4 | REX4, 3'-5' exonuclease | 227656 | MGI:2684957 | protein coding | 0.511 | 0.000157927 | n | n | n | n | n | n |
| Rab3b | RAB3B, member RAS oncoge | 69908 | MGI:1917158 | protein coding | 0.510 | 6.29E-05 | n | n | n | y | n | n |
| Trabd | TraB domain containing | 67976 | MGI:1915226 | protein coding | 0.509 | 0.000709185 | n | n | n | n | n | n |
| Abitram | actin binding transcription mo | 230234 | MGI:2677850 | protein coding | 0.508 | 0.000212673 | n | n | n | n | n | n |
| Tjap1 | tight junction associated prote | 74094 | MGI:1921344 | protein coding | 0.507 | 8.24E-06 | n | n | y | y | n | n |
| Pgm2l1 | phosphoglucomutase 2-like 1 | 70974 | MGI:1918224 | protein coding | 0.506 | 6.02E-05 | n | n | y | y | n | n |
| Mcoln3 | mucolipin 3 | 171166 | MGI:1890500 | protein coding | 0.506 | 2.75E-05 | n | n | n | n | y | n |
| Map2 | microtubule-associated protei | 17756 | MGI:97175 | protein coding | 0.505 | 0.000226324 | n | y | n | y | n | n |
| Me2 | malic enzyme 2, NAD(+)-depe | 107029 | MGI:2147351 | protein coding | 0.504 | 7.52E-05 | n | n | n | n | n | n |
| Frmd3 | FERM domain containing 3 | 242506 | MGI:2442466 | protein coding | 0.502 | 1.65E-05 | n | n | n | y | n | n |
| Ank3 | ankyrin 3, epithelial | 11735 | MGI:88026 | protein coding | 0.502 | 0.000394773 | n | n | y | y | n | n |
| Ift140 | intraflagellar transport 140 | 106633 | MGI:2146906 | protein coding | 0.502 | 4.21E-05 | n | n | n | n | n | y |
| Zfp324 | zinc finger protein 324 | 243834 | MGI:2444641 | protein coding | 0.501 | 0.000978867 | n | n | n | n | n | n |
| Plk2 | polo like kinase 2 | 20620 | MGI:1099790 | protein coding | 0.500 | 0.00040715 | n | n | n | n | n | n |
| Abhd4 | abhydrolase domain containir | 105501 | MGI:1915938 | protein coding | 0.499 | 3.00E-06 | n | n | n | n | n | n |
| Ttpal | tocopherol (alpha) transfer pro | 76080 | MGI:1923330 | protein coding | 0.498 | 0.000277178 | n | n | n | n | n | n |
| Hid1 | HID1 domain containing | 217310 | MGI:2445087 | protein coding | 0.496 | 1.84E-05 | n | n | n | y | n | n |
| Fkbp4 | FK506 binding protein 4 | 14228 | MGI:95543 | protein coding | 0.494 | 4.43E-06 | n | n | n | n | n | n |
| Mlf1 | myeloid leukemia factor 1 | 17349 | MGI:1341819 | protein coding | 0.493 | 0.0001922 | n | n | y | n | n | y |
| Mis12 | MIS12 kinetochore complex c | 67139 | MGI:1914389 | protein coding | 0.492 | 0.000970555 | n | n | n | n | n | n |
| Fbxo42 | F-box protein 42 | 213499 | MGI:1924992 | protein coding | 0.489 | 0.000144272 | n | n | n | n | n | n |
| Atg4d | autophagy related 4D, cystein | 235040 | MGI:2444308 | protein coding | 0.489 | 0.000135568 | n | n | y | n | n | n |
| Mid1ip1 | Mid1 interacting protein 1 (ga | 68041 | MGI:1915291 | protein coding | 0.488 | 0.000139658 | n | n | n | n | n | n |
| Slc9a9 | solute carrier family 9 (sodiun | 331004 | MGI:2679732 | protein coding | 0.486 | 0.00088973 | n | n | n | y | n | n |
| Slc2a12 | solute carrier family 2 (facilita | 353169 | MGI:3052471 | protein coding | 0.486 | 0.00084698 | n | n | n | n | n | n |
| Arv1 | ARV1 homolog, fatty acid hor | 68865 | MGI:1916115 | protein coding | 0.483 | 0.000676269 | n | n | n | y | n | n |
| Ckmt1 | creatine kinase, mitochondria | 12716 | MGI:99441 | protein coding | 0.483 | 0.000102818 | n | n | y | n | y** (PMID: 14977190) | n |
| Vcl | vinculin | 22330 | MGI:98927 | protein coding | 0.481 | 6.10E-05 | n | n | n | n | n | n |
| Rnpep | arginyl aminopeptidase (amin | 215615 | MGI:2384902 | protein coding | 0.479 | 0.000267028 | n | n | n | n | n | n |
| Soat1 | sterol O-acyltransferase 1 | 20652 | MGI:104665 | protein coding | 0.476 | 0.000392094 | n | n | n | n | n | n |
| Tmem131l | transmembrane 131 like | 229473 | MGI:2443399 | protein coding | 0.476 | 0.000465657 | n | n | n | y | n | n |
| Vti1a | vesicle transport through inter | 53611 | MGI:1855699 | protein coding | 0.475 | 0.000336102 | n | n | n | y | n | n |

| | | | | | | | | | | | | |
|---|---|---|---|---|---|---|---|---|---|---|---|---|
| Tmcc2 | transmembrane and coiled-cc | 68875 | MGI:1916125 | protein coding | 0.475 | 3.89E-05 | n | n | y | y | n | n |
| Clic5 | chloride intracellular channel | 224796 | MGI:1917912 | protein coding | 0.473 | 0.000419881 | n | n | y | n | y | n |
| Cfap69 | cilia and flagella associated p | 207686 | MGI:2443778 | protein coding | 0.473 | 0.000637521 | n | n | y | n | n | y |
| Insc | INSC spindle orientation adap | 233752 | MGI:1917942 | protein coding | 0.472 | 0.000161382 | n | n | y | y | n | n |
| Parm1 | prostate androgen-regulated r | 231440 | MGI:2443349 | protein coding | 0.472 | 0.000250058 | n | n | n | n | n | n |
| Rffl | ring finger and FYVE like dom | 67338 | MGI:1914588 | protein coding | 0.471 | 0.000236499 | n | n | n | y | n | n |
| Trappc13 | trafficking protein particle com | 66975 | MGI:1914225 | protein coding | 0.471 | 9.68E-05 | n | n | n | n | n | n |
| Creld1 | cysteine-rich with EGF-like do | 171508 | MGI:2152539 | protein coding | 0.470 | 9.15E-05 | n | n | n | n | n | n |
| Gorasp1 | golgi reassembly stacking pro | 74498 | MGI:1921748 | protein coding | 0.469 | 0.000946603 | n | n | n | n | n | n |
| Rtl6 | retrotransposon Gag like 6 | 223732 | MGI:2675858 | protein coding | 0.467 | 0.000338163 | n | n | n | y | n | n |
| Agpat4 | 1-acylglycerol-3-phosphate O | 68262 | MGI:1915512 | protein coding | 0.467 | 0.000243657 | n | n | n | y | n | n |
| B3gnt7 | UDP-GlcNAc:betaGal beta-1, | 227327 | MGI:2384394 | protein coding | 0.467 | 5.56E-05 | n | n | n | n | n | n |
| Acp6 | acid phosphatase 6, lysophos | 66659 | MGI:1931010 | protein coding | 0.466 | 0.000362676 | n | n | n | n | n | n |
| Poli | polymerase (DNA directed), io | 26447 | MGI:1347081 | protein coding | 0.465 | 0.000166319 | n | n | n | n | n | n |
| Tango2 | transport and golgi organizatio | 27883 | MGI:101825 | protein coding | 0.464 | 0.000162545 | n | n | n | n | n | n |
| Tmem117 | transmembrane protein 117 | 320709 | MGI:2444580 | protein coding | 0.464 | 5.20E-05 | n | n | n | y | n | n |
| Tfcp2 | transcription factor CP2 | 21422 | MGI:98509 | protein coding | 0.462 | 0.000441847 | n | n | n | n | n | n |
| Sugp2 | SURP and G patch domain co | 234373 | MGI:2678085 | protein coding | 0.462 | 0.000470676 | n | y | n | y | n | n |
| Agpat3 | 1-acylglycerol-3-phosphate O | 28169 | MGI:1336186 | protein coding | 0.461 | 0.0005257 | n | n | n | n | n | n |
| Dhx29 | DExH-box helicase 29 | 218629 | MGI:2145374 | protein coding | 0.461 | 0.000161312 | n | n | n | n | n | n |
| Map10 | microtubule-associated protei | 74393 | MGI:1921643 | protein coding | 0.460 | 0.000944557 | n | n | n | n | n | n |
| Slc22a5 | solute carrier family 22 (organ | 20520 | MGI:1329012 | protein coding | 0.460 | 7.82E-05 | n | n | n | n | n | n |
| Slc27a4 | solute carrier family 27 (fatty a | 26569 | MGI:1347347 | protein coding | 0.458 | 0.000153247 | n | n | n | n | n | n |
| Rspry1 | ring finger and SPRY domain | 67610 | MGI:1914860 | protein coding | 0.457 | 0.000955152 | n | n | n | n | n | n |
| Adgrg2 | adhesion G protein-coupled re | 237175 | MGI:2446854 | protein coding | 0.456 | 0.000933452 | n | n | n | n | n | n |
| Ahcyl2 | S-adenosylhomocysteine hyd | 74340 | MGI:1921590 | protein coding | 0.456 | 6.67E-05 | n | n | n | n | n | n |
| Usp20 | ubiquitin specific peptidase 20 | 74270 | MGI:1921520 | protein coding | 0.455 | 0.000209844 | n | n | y | y | n | n |
| Eif4e3 | eukaryotic translation initiation | 66892 | MGI:1914142 | protein coding | 0.455 | 0.00012618 | n | n | n | n | n | n |
| Ccar2 | cell cycle activator and apopto | 219158 | MGI:2444228 | protein coding | 0.454 | 2.88E-05 | n | n | n | n | n | n |
| Trpm4 | transient receptor potential ca | 68667 | MGI:1915917 | protein coding | 0.453 | 0.000224745 | n | n | n | n | n | n |
| Rap1gds1 | RAP1, GTP-GDP dissociation | 229877 | MGI:2385189 | protein coding | 0.453 | 0.000604745 | n | n | n | n | n | n |
| Ptpn3 | protein tyrosine phosphatase, | 545622 | MGI:105307 | protein coding | 0.453 | 0.00017024 | n | n | y | n | n | n |
| Snx1 | sorting nexin 1 | 56440 | MGI:1928395 | protein coding | 0.453 | 1.10E-05 | n | n | n | n | n | n |
| Ncald | neurocalcin delta | 52589 | MGI:1196326 | protein coding | 0.452 | 0.000277737 | n | n | n | y | n | n |
| Tceal3 | transcription elongation factor | 594844 | MGI:1913354 | protein coding | 0.452 | 0.000494769 | n | n | n | n | n | n |
| Sgsm1 | small G protein signaling mod | 52850 | MGI:107320 | protein coding | 0.451 | 0.000237097 | n | n | n | n | n | n |
| Cpt2 | carnitine palmitoyltransferase | 12896 | MGI:109176 | protein coding | 0.451 | 0.000255804 | n | n | n | n | n | n |
| Ercc6 | excision repair cross-complen | 319955 | MGI:1100494 | protein coding | 0.450 | 0.000971013 | n | y | n | y | y** (PMID: 25762674) | n |
| Mier2 | MIER family member 2 | 70427 | MGI:1917677 | protein coding | 0.450 | 0.000232653 | n | n | n | n | n | n |
| Pick1 | protein interacting with C kina | 18693 | MGI:894645 | protein coding | 0.448 | 7.75E-06 | n | n | n | y | n | n |
| Slitrk6 | SLIT and NTRK-like family, m | 239250 | MGI:2443198 | protein coding | 0.447 | 0.000191487 | n | n | n | n | y | n |
| Itsn2 | intersectin 2 | 20403 | MGI:1338049 | protein coding | 0.447 | 0.000778934 | n | n | n | n | n | n |
| Txlna | taxilin alpha | 109658 | MGI:105968 | protein coding | 0.446 | 4.15E-05 | n | n | n | n | n | n |
| Cyb561a3 | cytochrome b561 family, mem | 225912 | MGI:2686925 | protein coding | 0.446 | 0.000229096 | n | n | n | n | n | n |
| Rab4a | RAB4A, member RAS oncoge | 19341 | MGI:105069 | protein coding | 0.445 | 0.000162339 | n | n | n | n | n | n |
| Strip2 | striatin interacting protein 2 | 320609 | MGI:2444363 | protein coding | 0.445 | 0.000236087 | n | y | n | n | n | n |
| Arvcf | armadillo repeat gene deleted | 11877 | MGI:109620 | protein coding | 0.445 | 4.18E-05 | n | n | n | n | n | n |
| Prss36 | serine protease 36 | 77613 | MGI:1924863 | protein coding | 0.444 | 0.000111239 | n | n | n | n | n | n |
| Usp33 | ubiquitin specific peptidase 33 | 170822 | MGI:2159711 | protein coding | 0.444 | 6.69E-05 | n | n | n | y | n | n |
| Mettl17 | methyltransferase like 17 | 52535 | MGI:1098577 | protein coding | 0.443 | 0.000114071 | n | n | n | n | n | n |
| Tdrkh | tudor and KH domain contain | 72634 | MGI:1919884 | protein coding | 0.441 | 0.000525479 | n | n | n | n | n | n |
| Shkbp1 | Sh3kbp1 binding protein 1 | 192192 | MGI:2385803 | protein coding | 0.441 | 0.000189922 | n | n | n | n | n | n |
| Atp9a | ATPase, class II, type 9A | 11981 | MGI:1330826 | protein coding | 0.438 | 5.15E-06 | n | n | n | n | n | n |
| Prkcz | protein kinase C, zeta | 18762 | MGI:97602 | protein coding | 0.438 | 0.00087057 | n | n | n | n | n | n |
| Atosa | atos homolog A | 235493 | MGI:2387648 | protein coding | 0.436 | 0.000346078 | n | y | n | y | n | n |
| Tomt | transmembrane O-methyltran | 791260 | MGI:3769724 | protein coding | 0.435 | 0.000670323 | n | n | y | n | y | n |
| Dclk1 | doublecortin-like kinase 1 | 13175 | MGI:1330861 | protein coding | 0.434 | 0.000185058 | n | n | n | n | n | n |
| Zfp346 | zinc finger protein 346 | 26919 | MGI:1349417 | protein coding | 0.432 | 0.000925473 | n | n | n | n | n | n |
| Abcf3 | ATP-binding cassette, sub-far | 27406 | MGI:1351656 | protein coding | 0.430 | 0.000258754 | n | n | n | n | n | n |
| Pde6d | phosphodiesterase 6D, cGMP | 18582 | MGI:1270843 | protein coding | 0.429 | 3.01E-05 | n | n | n | n | n | n |
| Sec14l1 | SEC14-like lipid binding 1 | 74136 | MGI:1921386 | protein coding | 0.428 | 0.000596047 | n | n | n | y | n | n |
| Cstf3 | cleavage stimulation factor, 3 | 228410 | MGI:1351825 | protein coding | 0.428 | 0.000719821 | n | n | n | n | n | n |
| Map9 | microtubule-associated protei | 213582 | MGI:2442208 | protein coding | 0.428 | 0.000317066 | n | n | y | n | n | n |
| Bicd2 | BICD cargo adaptor 2 | 76895 | MGI:1924145 | protein coding | 0.425 | 0.000564741 | n | n | n | n | n | n |
| Sel1l3 | sel-1 suppressor of lin-12-like | 231238 | MGI:1916941 | protein coding | 0.425 | 0.000214151 | n | n | n | n | n | n |
| Tmem30b | transmembrane protein 30B | 238257 | MGI:2442082 | protein coding | 0.424 | 0.00025105 | n | n | n | n | n | n |
| Cfap418 | cilia and flagella associated p | 67157 | MGI:1914407 | protein coding | 0.423 | 5.52E-05 | n | n | n | n | n | y |

| | | | | | | | | | | | | |
|---|---|---|---|---|---|---|---|---|---|---|---|---|
| Ric8b | RIC8 guanine nucleotide excl | 237422 | MGI:2682307 | protein coding | 0.422 | 0.000645205 | n | n | n | y | n | n |
| Creld2 | cysteine-rich with EGF-like do | 76737 | MGI:1923987 | protein coding | 0.421 | 0.000175604 | n | n | n | n | n | n |
| Arhgef10l | Rho guanine nucleotide excha | 72754 | MGI:1920004 | protein coding | 0.421 | 0.00012535 | n | n | n | y | n | n |
| Kcnj13 | potassium inwardly-rectifying | 100040591 | MGI:3781032 | protein coding | 0.419 | 0.000261875 | n | n | y | n | n | n |
| C9orf72 | C9orf72, member of C9orf72- | 73205 | MGI:1920455 | protein coding | 0.418 | 1.84E-05 | n | n | n | y | n | n |
| Pomgnt2 | protein O-linked mannose bet | 215494 | MGI:2143424 | protein coding | 0.418 | 0.000124389 | n | n | n | y | n | n |
| Ate1 | arginyltransferase 1 | 11907 | MGI:1333870 | protein coding | 0.418 | 0.000447824 | n | n | y | y | n | n |
| Tmem184b | transmembrane protein 184b | 223693 | MGI:2445179 | protein coding | 0.418 | 0.000608279 | n | n | n | y | n | n |
| Vwa5a | von Willebrand factor A doma | 67776 | MGI:1915026 | protein coding | 0.418 | 0.000479547 | n | n | n | n | n | n |
| Ugp2 | UDP-glucose pyrophosphoryl | 216558 | MGI:2183447 | protein coding | 0.415 | 2.96E-06 | n | n | n | n | n | n |
| Nek4 | NIMA (never in mitosis gene a | 23955 | MGI:1344404 | protein coding | 0.414 | 0.000517424 | n | n | n | n | n | y |
| Uaca | uveal autoantigen with coiled- | 72565 | MGI:1919815 | protein coding | 0.413 | 0.000716683 | n | n | y | n | n | n |
| Anxa5 | annexin A5 | 11747 | MGI:106008 | protein coding | 0.413 | 0.000383621 | n | n | n | n | n | n |
| Sacm1l | SAC1 suppressor of actin mu | 83493 | MGI:1933169 | protein coding | 0.412 | 0.000155212 | n | n | y | n | n | n |
| Sugp1 | SURP and G patch domain co | 70616 | MGI:1917866 | protein coding | 0.411 | 4.30E-05 | n | n | n | n | n | n |
| Fhip2b | FHF complex subunit HOOK | 239170 | MGI:3036290 | protein coding | 0.408 | 0.000417769 | n | n | n | n | n | n |
| Irf3 | interferon regulatory factor 3 | 54131 | MGI:1859179 | protein coding | 0.407 | 0.000348975 | n | n | n | n | n | n |
| Lhx3 | LIM homeobox protein 3 | 16871 | MGI:102673 | protein coding | 0.404 | 0.000557081 | n | n | y | n | n | n |
| Cds2 | CDP-diacylglycerol synthase | 110911 | MGI:1332236 | protein coding | 0.403 | 0.000228772 | n | n | y | n | n | n |
| Sorbs2 | sorbin and SH3 domain conta | 234214 | MGI:1924574 | protein coding | 0.402 | 0.000148155 | n | n | n | y | n | n |
| Golm1 | golgi membrane protein 1 | 105348 | MGI:1917329 | protein coding | 0.399 | 0.000310131 | n | n | y | y | n | n |
| G3bp1 | G3BP stress granule assemb | 27041 | MGI:1351465 | protein coding | 0.399 | 0.000508922 | n | n | n | n | n | n |
| Pxylp1 | 2-phosphoxylose phosphatas | 235534 | MGI:2442444 | protein coding | 0.397 | 0.000153247 | n | n | n | n | n | n |
| Tex264 | testis expressed gene 264 | 21767 | MGI:1096570 | protein coding | 0.396 | 0.000122402 | n | n | n | n | n | n |
| Zfp612 | zinc finger protein 612 | 234725 | MGI:2443465 | protein coding | 0.396 | 0.000177378 | n | n | y | n | n | n |
| Mfsd14b | major facilitator superfamily d | 66631 | MGI:1913881 | protein coding | 0.395 | 0.000311688 | n | n | n | n | n | n |
| Srd5a1 | steroid 5 alpha-reductase 1 | 78925 | MGI:98400 | protein coding | 0.395 | 9.04E-05 | n | n | y | y | n | n |
| Hgs | HGF-regulated tyrosine kinas | 15239 | MGI:104681 | protein coding | 0.395 | 0.000494769 | n | n | y | n | n | n |
| Muc15 | mucin 15 | 269328 | MGI:2442110 | protein coding | 0.394 | 7.90E-05 | n | n | n | n | n | n |
| Ciapin1 | cytokine induced apoptosis in | 109006 | MGI:1922083 | protein coding | 0.393 | 0.000136239 | n | n | n | n | n | n |
| Nek1 | NIMA (never in mitosis gene a | 18004 | MGI:97303 | protein coding | 0.393 | 0.000759763 | n | n | n | n | n | n |
| Lrrc20 | leucine rich repeat containing | 216011 | MGI:2387182 | protein coding | 0.391 | 0.000602891 | n | n | n | y | n | n |
| Ccnd1 | cyclin D1 | 12443 | MGI:88313 | protein coding | 0.391 | 0.000238775 | n | n | n | n | n | n |
| Snap91 | synaptosomal-associated pro | 20616 | MGI:109132 | protein coding | 0.390 | 2.15E-05 | n | n | n | n | n | n |
| Tmem184c | transmembrane protein 184C | 234463 | MGI:2384562 | protein coding | 0.390 | 5.32E-05 | n | n | n | n | n | n |
| Ufsp2 | UFM1-specific peptidase 2 | 192169 | MGI:1913679 | protein coding | 0.388 | 0.000245659 | n | n | n | n | n | n |
| Ift70b | intraflagellar transport 70B | 72421 | MGI:1919671 | protein coding | 0.388 | 0.000306045 | n | n | n | n | n | y |
| Ppa1 | pyrophosphatase (inorganic) | 67895 | MGI:97831 | protein coding | 0.388 | 4.84E-05 | n | n | n | n | n | n |
| Usp46 | ubiquitin specific peptidase 46 | 69727 | MGI:1916977 | protein coding | 0.387 | 0.000725083 | n | n | y | y | n | n |
| Rab36 | RAB36, member RAS oncoge | 76877 | MGI:1924127 | protein coding | 0.387 | 5.95E-05 | n | n | y | y | n | n |
| Misp | mitotic spindle positioning | 78906 | MGI:1926156 | protein coding | 0.386 | 0.000515786 | n | n | n | n | n | n |
| Zfp512 | zinc finger protein 512 | 269639 | MGI:1917345 | protein coding | 0.383 | 0.000331954 | n | n | n | n | n | n |
| Tars2 | threonyl-tRNA synthetase 2, r | 71807 | MGI:1919057 | protein coding | 0.382 | 0.000415168 | n | n | n | y | n | n |
| Gys1 | glycogen synthase 1, muscle | 14936 | MGI:101805 | protein coding | 0.382 | 0.000308484 | n | n | n | n | n | n |
| Fam222b | family with sequence similarit | 216971 | MGI:2384939 | protein coding | 0.381 | 0.0001307 | n | n | n | n | n | n |
| Gart | phosphoribosylglycinamide fo | 14450 | MGI:95654 | protein coding | 0.381 | 0.000365855 | n | n | n | n | n | n |
| Smpx | small muscle protein, X-linked | 66106 | MGI:1913356 | protein coding | 0.380 | 0.000207342 | n | n | y | y | y** (PMID: 34722533) | n |
| Ttll12 | tubulin tyrosine ligase-like fan | 223723 | MGI:3039573 | protein coding | 0.378 | 0.00035479 | n | n | n | n | n | n |
| Ncbp2 | nuclear cap binding protein su | 68092 | MGI:1915342 | protein coding | 0.378 | 0.000236087 | n | n | n | n | n | n |
| Flot1 | flotillin 1 | 14251 | MGI:1100500 | protein coding | 0.375 | 5.89E-05 | n | n | n | n | n | n |
| Scmh1 | sex comb on midleg homolog | 29871 | MGI:1352762 | protein coding | 0.375 | 0.000596339 | n | n | n | n | n | n |
| Stx2 | syntaxin 2 | 13852 | MGI:108059 | protein coding | 0.375 | 0.000501224 | n | n | n | n | n | n |
| Ttc19 | tetratricopeptide repeat doma | 72795 | MGI:1920045 | protein coding | 0.374 | 0.000233407 | n | n | y | y | n | n |
| Ccdc91 | coiled-coil domain containing | 67015 | MGI:1914265 | protein coding | 0.373 | 0.000634646 | n | n | n | n | n | n |
| Miga2 | mitoguardin 2 | 108958 | MGI:1922035 | protein coding | 0.373 | 0.000475152 | n | n | y | n | n | n |
| Wscd2 | WSC domain containing 2 | 320916 | MGI:2445030 | protein coding | 0.372 | 6.31E-05 | n | n | n | n | n | n |
| Dxo | decapping exoribonuclease | 112403 | MGI:1890444 | protein coding | 0.372 | 0.0006791 | n | n | n | n | n | n |
| Mocos | molybdenum cofactor sulfura: | 68591 | MGI:1915841 | protein coding | 0.370 | 0.000724395 | n | n | n | n | n | n |
| Oxct1 | 3-oxoacid CoA transferase 1 | 67041 | MGI:1914291 | protein coding | 0.370 | 0.000437435 | n | n | n | n | n | n |
| Ro60 | Ro60, Y RNA binding protein | 20822 | MGI:106652 | protein coding | 0.366 | 0.000226396 | n | n | n | n | n | n |
| Ccn4 | cellular communication netwo | 22402 | MGI:1197008 | protein coding | 0.365 | 0.000291115 | n | n | n | n | n | n |
| Tcaf1 | TRPM8 channel-associated fa | 77574 | MGI:1914665 | protein coding | 0.365 | 0.000517424 | n | n | n | n | n | n |
| Mtg2 | mitochondrial ribosome assoc | 52856 | MGI:106565 | protein coding | 0.364 | 0.000932068 | n | n | n | y | n | n |
| Tmem255b | transmembrane protein 255B | 272465 | MGI:2685533 | protein coding | 0.363 | 0.00016505 | n | n | y | y | n | n |
| Oxr1 | oxidation resistance 1 | 170719 | MGI:2179326 | protein coding | 0.363 | 0.000432072 | n | n | n | n | n | n |
| Ap2a2 | adaptor-related protein compl | 11772 | MGI:101920 | protein coding | 0.362 | 5.09E-05 | n | n | n | n | n | n |
| Stk16 | serine/threonine kinase 16 | 20872 | MGI:1313271 | protein coding | 0.362 | 0.000679935 | n | n | n | n | n | n |

| | | | | | | | | | | | | |
|---|---|---|---|---|---|---|---|---|---|---|---|---|
| Syvn1 | synovial apoptosis inhibitor 1, | 74126 | MGI:1921376 | protein coding | 0.358 | 0.000262692 | n | n | n | y | n | n |
| Gpd2 | glycerol phosphate dehydrog | 14571 | MGI:99778 | protein coding | 0.358 | 0.000196359 | n | n | n | n | n | n |
| Nprl2 | NPR2 like, GATOR1 complex | 56032 | MGI:1914482 | protein coding | 0.357 | 0.000637086 | n | n | y | n | n | n |
| Lima1 | LIM domain and actin binding | 65970 | MGI:1920992 | protein coding | 0.356 | 0.000601212 | n | n | n | y | n | n |
| Gnpat | glyceronephosphate O-acyltra | 14712 | MGI:1343460 | protein coding | 0.355 | 0.000208746 | n | n | n | n | n | n |
| Atp6v1b2 | ATPase, H+ transporting, lys | 11966 | MGI:109618 | protein coding | 0.355 | 0.000207561 | n | n | n | n | n | n |
| Cacnb3 | calcium channel, voltage-dep | 12297 | MGI:103307 | protein coding | 0.355 | 2.46E-05 | n | n | n | y | n | n |
| Rnf5 | ring finger protein 5 | 54197 | MGI:1860076 | protein coding | 0.355 | 0.000120292 | n | n | n | n | n | n |
| Taok2 | TAO kinase 2 | 381921 | MGI:1915919 | protein coding | 0.349 | 0.000502308 | n | n | n | n | n | n |
| Hpcal1 | hippocalcin-like 1 | 53602 | MGI:1855689 | protein coding | 0.348 | 0.000171489 | n | n | n | y | n | n |
| Atp6v0e | ATPase, H+ transporting, lys | 11974 | MGI:1328318 | protein coding | 0.347 | 0.000782777 | n | n | n | y | n | n |
| Gabarapl1 | GABA type A receptor associ | 57436 | MGI:1914980 | protein coding | 0.346 | 0.000374456 | n | n | n | n | n | n |
| Ppp5c | protein phosphatase 5, cataly | 19060 | MGI:102666 | protein coding | 0.345 | 7.52E-05 | n | n | n | n | n | n |
| Slco3a1 | solute carrier organic anion tr | 108116 | MGI:1351867 | protein coding | 0.344 | 0.000140331 | n | n | n | y | n | n |
| Rcan1 | regulator of calcineurin 1 | 54720 | MGI:1890564 | protein coding | 0.342 | 0.000925473 | n | n | n | n | n | n |
| 2810004N23Rik | RIKEN cDNA 2810004N23 ge | 66523 | MGI:1913773 | protein coding | 0.341 | 0.000415168 | n | n | n | n | n | n |
| Ppme1 | protein phosphatase methyles | 72590 | MGI:1919840 | protein coding | 0.339 | 0.000218532 | n | n | n | n | n | n |
| Yipf3 | Yip1 domain family, member | 28064 | MGI:106280 | protein coding | 0.337 | 0.000955152 | n | n | n | n | n | n |
| Dbn1 | drebrin 1 | 56320 | MGI:1931838 | protein coding | 0.336 | 0.000459513 | n | n | n | n | n | n |
| Rnf6 | ring finger protein (C3H2C3 ty | 74132 | MGI:1921382 | protein coding | 0.335 | 0.000767488 | n | n | n | n | n | n |
| Fuca1 | fucosidase, alpha-L- 1, tissue | 71665 | MGI:95593 | protein coding | 0.335 | 0.000111259 | n | n | n | n | n | n |
| Lrrtm1 | leucine rich repeat transmeml | 74342 | MGI:2389173 | protein coding | 0.332 | 0.000108888 | n | n | n | n | n | n |
| Nrdc | nardilysin convertase | 230598 | MGI:1201386 | protein coding | 0.332 | 0.000449066 | n | n | y | n | n | n |
| Iars1 | isoleucyl-tRNA synthetase 1 | 105148 | MGI:2145219 | protein coding | 0.331 | 0.000970036 | n | n | n | n | n | n |
| Zranb2 | zinc finger, RAN-binding dom | 53861 | MGI:1858211 | protein coding | 0.330 | 0.0006791 | n | n | n | n | n | n |
| Bphl | biphenyl hydrolase like | 68021 | MGI:1915271 | protein coding | 0.329 | 0.000521532 | n | n | n | n | n | n |
| Kifap3 | kinesin-associated protein 3 | 16579 | MGI:107566 | protein coding | 0.329 | 0.000971013 | y | n | n | n | n | y |
| Trim13 | tripartite motif-containing 13 | 66597 | MGI:1913847 | protein coding | 0.324 | 0.000975788 | n | n | n | n | n | n |
| Osbpl11 | oxysterol binding protein-like | 106326 | MGI:2146553 | protein coding | 0.322 | 0.000636483 | n | n | y | n | n | n |
| Dcaf12 | DDB1 and CUL4 associated f | 68970 | MGI:1916220 | protein coding | 0.322 | 0.000832284 | n | n | n | n | n | n |
| Hdac5 | histone deacetylase 5 | 15184 | MGI:1333784 | protein coding | 0.321 | 0.000502308 | n | n | y | n | n | n |
| Cct4 | chaperonin containing TCP1 s | 12464 | MGI:104689 | protein coding | 0.321 | 0.000108025 | n | n | n | n | n | y |
| Prep | prolyl endopeptidase | 19072 | MGI:1270863 | protein coding | 0.318 | 0.000192253 | n | n | n | n | n | n |
| Ddr1 | discoidin domain receptor fan | 12305 | MGI:99216 | protein coding | 0.313 | 0.0005203 | n | n | n | y | n | n |
| Nagk | N-acetylglucosamine kinase | 56174 | MGI:1860418 | protein coding | 0.313 | 0.000851021 | n | n | n | n | n | n |
| Yars1 | tyrosyl-tRNA synthetase 1 | 107271 | MGI:2147627 | protein coding | 0.312 | 0.000171489 | n | n | y | n | n | n |
| Spire2 | spire type actin nucleation fac | 234857 | MGI:2446256 | protein coding | 0.308 | 0.000752339 | n | n | n | n | n | n |
| Eya1 | EYA transcriptional coactivato | 14048 | MGI:109344 | protein coding | 0.306 | 0.00082903 | n | n | n | y | y** (PMID: 18678597) | n |
| Sae1 | SUMO1 activating enzyme su | 56459 | MGI:1929264 | protein coding | 0.305 | 0.000135568 | n | n | n | y | n | n |
| Camk2b | calcium/calmodulin-depender | 12323 | MGI:88257 | protein coding | 0.305 | 0.000227957 | n | n | y | n | n | n |
| Kmt5c | lysine methyltransferase 5C | 232811 | MGI:2385262 | protein coding | 0.302 | 0.000724984 | n | n | n | n | n | n |
| Faah | fatty acid amide hydrolase | 14073 | MGI:109609 | protein coding | 0.296 | 0.00044637 | n | n | n | y | n | n |
| Ugdh | UDP-glucose dehydrogenase | 22235 | MGI:1306785 | protein coding | 0.296 | 0.000475284 | n | n | n | n | n | n |
| Fbxo3 | F-box protein 3 | 57443 | MGI:1929084 | protein coding | 0.292 | 0.00035479 | n | n | n | n | n | n |
| Fads2 | fatty acid desaturase 2 | 56473 | MGI:1930079 | protein coding | 0.291 | 0.000505711 | n | n | n | n | n | n |
| Tmem30a | transmembrane protein 30A | 69981 | MGI:106402 | protein coding | 0.289 | 0.000900618 | n | n | n | n | n | n |
| Fkbp8 | FK506 binding protein 8 | 14232 | MGI:1341070 | protein coding | 0.288 | 0.000745635 | n | n | n | n | n | n |
| Camta1 | calmodulin binding transcripti | 100072 | MGI:2140230 | protein coding | 0.285 | 0.000347975 | n | n | n | y | n | n |
| Lztr1 | leucine-zipper-like transcriptic | 66863 | MGI:1914113 | protein coding | 0.280 | 0.000607188 | n | n | n | n | n | n |
| Trpv4 | transient receptor potential ca | 63873 | MGI:1926945 | protein coding | 0.279 | 0.000955152 | n | n | n | n | n | y |
| Afg3l1 | AFG3-like AAA ATPase 1 | 114896 | MGI:1928277 | protein coding | 0.275 | 0.000857537 | n | n | n | n | n | n |
| Parp2 | poly (ADP-ribose) polymerase | 11546 | MGI:1341112 | protein coding | 0.274 | 0.000557081 | n | n | n | n | n | n |
| Bnip3l | BCL2/adenovirus E1B interac | 12177 | MGI:1332659 | protein coding | -0.273 | 0.000604096 | n | n | n | y | n | n |
| Mfsd10 | major facilitator superfamily d | 68294 | MGI:1915544 | protein coding | -0.274 | 0.000724984 | n | n | n | n | n | n |
| Ncam1 | neural cell adhesion molecule | 17967 | MGI:97281 | protein coding | -0.279 | 0.000719821 | n | n | n | n | n | n |
| 2410002F23Rik | RIKEN cDNA 2410002F23 ge | 668661 | MGI:1914226 | protein coding | -0.282 | 0.000425409 | n | n | n | n | n | n |
| Polr2e | polymerase (RNA) II (DNA dir | 66420 | MGI:1913670 | protein coding | -0.283 | 0.000818241 | n | n | n | n | n | n |
| Arl2bp | ADP-ribosylation factor-like 2 | 107566 | MGI:1349429 | protein coding | -0.285 | 0.00076909 | n | n | n | n | n | y |
| Bccip | BRCA2 and CDKN1A interact | 66165 | MGI:1913415 | protein coding | -0.288 | 0.000925473 | n | n | n | n | n | n |
| Rps6ka1 | ribosomal protein S6 kinase p | 20111 | MGI:104558 | protein coding | -0.295 | 0.000684005 | n | n | n | y | n | n |
| Ddx39a | DEAD box helicase 39a | 68278 | MGI:1915528 | protein coding | -0.295 | 0.000976572 | n | n | n | n | n | n |
| Rhbdf1 | rhomboid 5 homolog 1 | 13650 | MGI:104328 | protein coding | -0.299 | 0.000309585 | n | n | n | n | n | n |
| Eif3i | eukaryotic translation initiatior | 54709 | MGI:1860763 | protein coding | -0.299 | 0.000805895 | n | n | n | n | n | n |
| Cnbp | cellular nucleic acid binding p | 12785 | MGI:88431 | protein coding | -0.300 | 0.000649533 | n | n | n | n | n | n |
| Ppie | peptidylprolyl isomerase E (cy | 56031 | MGI:1917118 | protein coding | -0.306 | 0.00015055 | n | n | n | n | n | n |
| Strap | serine/threonine kinase recep | 20901 | MGI:1329037 | protein coding | -0.309 | 0.000325942 | n | n | n | n | n | n |
| Calml4 | calmodulin-like 4 | 75600 | MGI:1922850 | protein coding | -0.313 | 0.000343608 | n | n | n | n | n | n |

| | | | | | | | | | | | | |
|---|---|---|---|---|---|---|---|---|---|---|---|---|
| Cep57 | centrosomal protein 57 | 74360 | MGI:1915551 | protein coding | -0.315 | 0.000666015 | n | n | n | y | n | n |
| Thoc3 | THO complex 3 | 73666 | MGI:1920916 | protein coding | -0.320 | 0.000680269 | n | n | n | n | n | n |
| Ptbp1 | polypyrimidine tract binding p | 19205 | MGI:97791 | protein coding | -0.321 | 0.000306116 | n | n | y | y | n | n |
| Ecd | ecdysoneless cell cycle regul | 70601 | MGI:1917851 | protein coding | -0.322 | 0.000236326 | n | n | n | n | n | n |
| Dazap2 | DAZ associated protein 2 | 23994 | MGI:1344344 | protein coding | -0.322 | 0.000502308 | n | n | n | n | n | n |
| Scara3 | scavenger receptor class A, n | 219151 | MGI:2444418 | protein coding | -0.327 | 0.000786799 | n | n | n | n | n | n |
| Dstn | destrin | 56431 | MGI:1929270 | protein coding | -0.327 | 0.000600601 | n | n | n | n | n | n |
| Lancl1 | LanC (bacterial lantibiotic syn | 14768 | MGI:1336997 | protein coding | -0.333 | 0.000439307 | n | n | n | n | n | n |
| Cpne8 | copine VIII | 66871 | MGI:1914121 | protein coding | -0.333 | 0.000934048 | n | n | n | n | n | n |
| Apoe | apolipoprotein E | 11816 | MGI:88057 | protein coding | -0.335 | 0.000779202 | n | n | n | n | n | n |
| Rps3a1 | ribosomal protein S3A1 | 20091 | MGI:1202063 | protein coding | -0.335 | 0.000526722 | n | n | n | n | n | n |
| Rpl7 | ribosomal protein L7 | 19989 | MGI:98073 | protein coding | -0.338 | 0.000516912 | n | n | n | n | n | n |
| Timp2 | tissue inhibitor of metalloprote | 21858 | MGI:98753 | protein coding | -0.338 | 5.11E-05 | n | n | n | n | n | n |
| Ccz1 | CCZ1 vacuolar protein traffick | 231874 | MGI:2141070 | protein coding | -0.341 | 0.000107631 | n | n | n | y | n | n |
| Lhfpl5 | lipoma HMGIC fusion partner | 328789 | MGI:1915382 | protein coding | -0.343 | 0.000805895 | n | n | y | y | y | n |
| 1110038F14Rik | RIKEN cDNA 1110038F14 ge | 117171 | MGI:2152337 | protein coding | -0.343 | 0.000721338 | n | n | n | y | n | n |
| Eif2s2 | eukaryotic translation initiatio | 67204 | MGI:1914454 | protein coding | -0.343 | 0.000580797 | n | n | n | n | n | n |
| Timm44 | translocase of inner mitochon | 21856 | MGI:1343262 | protein coding | -0.345 | 0.000250349 | n | n | n | n | n | n |
| Cox4i1 | cytochrome c oxidase subuni | 12857 | MGI:88473 | protein coding | -0.347 | 0.000392086 | n | n | n | n | n | n |
| Mmp14 | matrix metallopeptidase 14 (n | 17387 | MGI:101900 | protein coding | -0.348 | 3.82E-05 | n | n | n | n | n | n |
| Mfge8 | milk fat globule EGF and facto | 17304 | MGI:102768 | protein coding | -0.351 | 0.000346078 | n | n | n | y | n | n |
| Tpm4 | tropomyosin 4 | 326618 | MGI:2449202 | protein coding | -0.352 | 0.000314352 | n | n | n | y | n | n |
| Smim10l1 | small integral membrane prot | 381820 | MGI:1914379 | protein coding | -0.354 | 0.000249993 | n | n | n | n | n | n |
| Tubb4a | tubulin, beta 4A class IVA | 22153 | MGI:107848 | protein coding | -0.354 | 0.000738513 | n | n | n | n | n | y |
| Ctsf | cathepsin F | 56464 | MGI:1861434 | protein coding | -0.359 | 0.000107051 | n | n | y | y | n | n |
| Colgalt1 | collagen beta(1-O)galactosylt | 234407 | MGI:1924348 | protein coding | -0.361 | 0.000126073 | n | n | n | n | n | n |
| Myl9 | myosin, light polypeptide 9, re | 98932 | MGI:2138915 | protein coding | -0.362 | 3.37E-05 | n | n | n | n | n | n |
| Lpcat2 | lysophosphatidylcholine acylt | 270084 | MGI:3606214 | protein coding | -0.362 | 0.00016505 | n | n | n | n | n | n |
| Acot2 | acyl-CoA thioesterase 2 | 171210 | MGI:2159605 | protein coding | -0.365 | 0.000818069 | n | n | n | n | n | n |
| Habp4 | hyaluronic acid binding protei | 56541 | MGI:1891713 | protein coding | -0.370 | 0.000475152 | n | n | n | n | n | n |
| Hmgb1 | high mobility group box 1 | 15289 | MGI:96113 | protein coding | -0.371 | 0.000172158 | n | n | n | n | n | n |
| Ramp3 | receptor (calcitonin) activity m | 56089 | MGI:1860292 | protein coding | -0.376 | 0.000955152 | n | n | n | n | n | n |
| Nt5c2 | 5'-nucleotidase, cytosolic II | 76952 | MGI:2178563 | protein coding | -0.377 | 0.000306486 | n | n | n | n | n | n |
| Use1 | unconventional SNARE in the | 67023 | MGI:1914273 | protein coding | -0.377 | 0.000303617 | n | n | n | n | n | n |
| Ctdsp2 | CTD small phosphatase 2 | 52468 | MGI:1098748 | protein coding | -0.377 | 5.77E-05 | n | n | n | n | n | n |
| Selenop | selenoprotein P | 20363 | MGI:894288 | protein coding | -0.378 | 4.21E-05 | n | n | n | n | n | n |
| Coq7 | demethyl-Q 7 | 12850 | MGI:107207 | protein coding | -0.378 | 0.000643881 | n | n | n | n | n | n |
| Cbfb | core binding factor beta | 12400 | MGI:99851 | protein coding | -0.380 | 0.000275445 | n | n | n | y | n | n |
| Pear1 | platelet endothelial aggregatio | 73182 | MGI:1920432 | protein coding | -0.384 | 0.00063702 | n | n | n | n | n | n |
| Serpine2 | serine (or cysteine) peptidase | 20720 | MGI:101780 | protein coding | -0.388 | 0.000149776 | n | n | n | n | n | n |
| Nhp2 | NHP2 ribonucleoprotein | 52530 | MGI:1098547 | protein coding | -0.388 | 0.00012668 | n | n | n | n | n | n |
| Cep43 | centrosomal protein 43 | 75296 | MGI:1922546 | protein coding | -0.389 | 0.000559147 | n | n | n | n | n | n |
| Tmem179 | transmembrane protein 179 | 104885 | MGI:2144891 | protein coding | -0.390 | 0.000544428 | n | n | n | n | n | n |
| Esd | esterase D/formylglutathione | 13885 | MGI:95421 | protein coding | -0.392 | 0.000213359 | n | n | n | y | n | n |
| Nphp1 | nephronophthisis 1 (juvenile) | 53885 | MGI:1858233 | protein coding | -0.392 | 0.000314158 | n | n | y | n | n | y |
| Egfl7 | EGF-like domain 7 | 353156 | MGI:2449923 | protein coding | -0.393 | 0.000933452 | n | n | y | n | n | n |
| Selenoh | selenoprotein H | 72657 | MGI:1919907 | protein coding | -0.395 | 0.000192253 | n | n | n | n | n | n |
| Tspan15 | tetraspanin 15 | 70423 | MGI:1917673 | protein coding | -0.396 | 0.00023563 | n | n | n | n | n | n |
| Tomm22 | translocase of outer mitochon | 223696 | MGI:2450248 | protein coding | -0.401 | 0.000360196 | n | n | n | n | n | n |
| Maip1 | matrix AAA peptidase interact | 68115 | MGI:1915365 | protein coding | -0.402 | 0.000756559 | n | n | n | n | n | n |
| Pnrc2 | proline-rich nuclear receptor c | 52830 | MGI:106512 | protein coding | -0.402 | 1.02E-05 | n | n | n | n | n | n |
| Clic4 | chloride intracellular channel | 29876 | MGI:1352754 | protein coding | -0.403 | 2.91E-05 | n | n | n | y | n | n |
| Ybx1 | Y box protein 1 | 22608 | MGI:99146 | protein coding | -0.404 | 0.000162339 | n | n | n | n | n | n |
| Zfp131 | zinc finger protein 131 | 72465 | MGI:1919715 | protein coding | -0.405 | 0.000727166 | n | n | n | n | n | n |
| Ppp1r15a | protein phosphatase 1, regula | 17872 | MGI:1927072 | protein coding | -0.410 | 0.000841905 | n | n | n | n | n | n |
| S100b | S100 protein, beta polypeptid | 20203 | MGI:98217 | protein coding | -0.411 | 0.000499686 | n | n | n | n | n | n |
| Rpl13a | ribosomal protein L13A | 22121 | MGI:1351455 | protein coding | -0.411 | 0.000903517 | n | n | n | n | n | n |
| Cdv3 | carnitine deficiency-associate | 321022 | MGI:2448759 | protein coding | -0.412 | 5.16E-05 | n | n | n | n | n | n |
| Cope | coatomer protein complex, su | 59042 | MGI:1891702 | protein coding | -0.417 | 0.000102818 | n | n | n | n | n | n |
| Eef2k | eukaryotic elongation factor-2 | 13631 | MGI:1195261 | protein coding | -0.418 | 7.99E-05 | n | n | n | n | n | n |
| Etv4 | ets variant 4 | 18612 | MGI:99423 | protein coding | -0.423 | 0.000303719 | n | n | n | n | n | n |
| Nap1l1 | nucleosome assembly protein | 53605 | MGI:1855693 | protein coding | -0.425 | 3.30E-05 | n | n | n | n | n | n |
| Dchs1 | dachsous cadherin related 1 | 233651 | MGI:2685011 | protein coding | -0.426 | 0.000938057 | n | n | n | n | n | n |
| Arpc1b | actin related protein 2/3 comp | 11867 | MGI:1343142 | protein coding | -0.427 | 0.000228817 | n | n | n | n | n | n |
| Pam | peptidylglycine alpha-amidati | 18484 | MGI:97475 | protein coding | -0.427 | 0.00017726 | n | n | n | n | n | n |
| Irx5 | Iroquois homeobox 5 | 54352 | MGI:1859086 | protein coding | -0.427 | 0.000385082 | n | n | n | n | n | n |
| Emcn | endomucin | 59308 | MGI:1891716 | protein coding | -0.428 | 0.000799739 | n | n | n | y | n | n |

| | | | | | | | | | | | | |
|---|---|---|---|---|---|---|---|---|---|---|---|---|
| Ace | angiotensin I converting enzy | 11421 | MGI:87874 | protein coding | -0.430 | 0.000719821 | n | n | n | n | n | y |
| Mterf4 | mitochondrial transcription ter | 69821 | MGI:1918355 | protein coding | -0.436 | 0.000522519 | n | n | n | n | n | n |
| Gstm7 | glutathione S-transferase, mu | 68312 | MGI:1915562 | protein coding | -0.439 | 0.000144272 | n | n | n | n | n | n |
| Tmem25 | transmembrane protein 25 | 71687 | MGI:1918937 | protein coding | -0.439 | 0.000577442 | n | n | n | n | n | n |
| Ptn | pleiotrophin | 19242 | MGI:97804 | protein coding | -0.441 | 2.94E-05 | n | n | n | y | n | n |
| Ppp1r16b | protein phosphatase 1, regula | 228852 | MGI:2151841 | protein coding | -0.443 | 0.000345757 | n | n | n | n | n | n |
| Lhfpl6 | LHFPL tetraspan subfamily m | 108927 | MGI:1920048 | protein coding | -0.444 | 0.000779486 | n | n | n | n | n | n |
| Mthfsl | 5, 10-methenyltetrahydrofolat | 100039707 | MGI:3780550 | protein coding | -0.445 | 0.000971224 | n | n | n | n | n | n |
| Srsf10 | serine and arginine-rich splici | 14105 | MGI:1333805 | protein coding | -0.446 | 0.00073827 | n | n | n | n | n | n |
| Gpr85 | G protein-coupled receptor 85 | 64450 | MGI:1927851 | protein coding | -0.447 | 0.000942351 | n | n | n | n | n | n |
| Svbp | small vasohibin binding protei | 69216 | MGI:1916466 | protein coding | -0.448 | 9.39E-05 | n | n | y | n | n | n |
| Cox11 | cytochrome c oxidase assemb | 69802 | MGI:1917052 | protein coding | -0.449 | 0.000148567 | n | n | n | n | n | n |
| Btf3l4 | basic transcription factor 3-lik | 70533 | MGI:1915312 | protein coding | -0.449 | 2.32E-05 | n | n | n | n | n | n |
| Osgep | O-sialoglycoprotein endopept | 66246 | MGI:1913496 | protein coding | -0.450 | 0.000200212 | n | n | n | n | n | n |
| Fam234b | family with sequence similarit | 74525 | MGI:1921775 | protein coding | -0.451 | 1.44E-05 | n | n | y | n | n | n |
| Sec61a2 | SEC61 translocon subunit alp | 57743 | MGI:1931071 | protein coding | -0.451 | 0.000139658 | n | n | y | n | n | n |
| Nmi | N-myc (and STAT) interactor | 64685 | MGI:1928368 | protein coding | -0.453 | 0.000865883 | n | n | n | n | n | n |
| Pdcd5 | programmed cell death 5 | 56330 | MGI:1913538 | protein coding | -0.453 | 0.000296098 | n | n | n | n | n | n |
| Gpatch4 | G patch domain containing 4 | 66614 | MGI:1913864 | protein coding | -0.455 | 0.000677957 | n | n | n | n | n | n |
| Ttyh2 | tweety family member 2 | 117160 | MGI:2157091 | protein coding | -0.461 | 0.00024897 | n | n | n | n | n | n |
| Atg14 | autophagy related 14 | 100504663 | MGI:1261775 | protein coding | -0.465 | 5.77E-05 | n | n | n | n | n | y |
| Arsb | arylsulfatase B | 11881 | MGI:88075 | protein coding | -0.465 | 0.000928304 | n | n | n | n | n | n |
| Pgpep1 | pyroglutamyl-peptidase I | 66522 | MGI:1913772 | protein coding | -0.469 | 0.000103367 | n | n | n | n | n | n |
| 1700025G04Rik | RIKEN cDNA 1700025G04 ge | 69399 | MGI:1916649 | protein coding | -0.470 | 0.000247118 | n | n | n | y | n | n |
| Ifitm2 | interferon induced transmemb | 80876 | MGI:1933382 | protein coding | -0.471 | 0.000531078 | n | n | n | n | n | n |
| Selenbp1 | selenium binding protein 1 | 20341 | MGI:96825 | protein coding | -0.471 | 0.000976708 | n | n | n | n | n | n |
| Ppp1r1a | protein phosphatase 1, regula | 58200 | MGI:1889595 | protein coding | -0.473 | 6.08E-05 | n | n | n | n | n | n |
| Rnf214 | ring finger protein 214 | 235315 | MGI:2444451 | protein coding | -0.477 | 0.000166874 | n | n | n | n | n | n |
| Rpl18a | ribosomal protein L18A | 76808 | MGI:1924058 | protein coding | -0.480 | 0.000803392 | n | n | n | n | n | n |
| Meis3 | Meis homeobox 3 | 17537 | MGI:108519 | protein coding | -0.480 | 0.000138462 | n | n | n | n | n | n |
| Erbin | Erbb2 interacting protein | 59079 | MGI:1890169 | protein coding | -0.481 | 0.00097668 | n | n | n | y | n | n |
| Cp | ceruloplasmin | 12870 | MGI:88476 | protein coding | -0.481 | 0.000109102 | n | n | n | n | n | n |
| Ggact | gamma-glutamylamine cyclot | 223267 | MGI:2385008 | protein coding | -0.484 | 0.000726877 | n | n | n | n | n | n |
| Wee1 | WEE 1 homolog 1 (S. pombe | 22390 | MGI:103075 | protein coding | -0.486 | 0.000461825 | n | n | n | y | n | n |
| Eral1 | Era like 12S mitochondrial rR | 57837 | MGI:1889295 | protein coding | -0.489 | 0.000211842 | n | n | n | n | n | n |
| Pink1 | PTEN induced putative kinase | 68943 | MGI:1916193 | protein coding | -0.489 | 1.58E-05 | n | n | n | n | n | n |
| Fyn | Fyn proto-oncogene | 14360 | MGI:95602 | protein coding | -0.489 | 0.000167697 | n | n | n | n | n | n |
| Rpl32 | ribosomal protein L32 | 19951 | MGI:98038 | protein coding | -0.490 | 0.000422229 | n | n | n | n | n | n |
| Lbr | lamin B receptor | 98386 | MGI:2138281 | protein coding | -0.492 | 0.000112667 | n | n | n | y | n | n |
| Tmem192 | transmembrane protein 192 | 73067 | MGI:1920317 | protein coding | -0.492 | 0.000291607 | n | n | n | n | n | n |
| Tmem234 | transmembrane protein 234 | 76799 | MGI:1924049 | protein coding | -0.496 | 4.94E-06 | n | n | n | y | n | n |
| Cd164l2 | CD164 sialomucin-like 2 | 69655 | MGI:1916905 | protein coding | -0.498 | 1.45E-05 | n | n | n | n | n | n |
| Syt6 | synaptotagmin VI | 54524 | MGI:1859544 | protein coding | -0.505 | 0.000229652 | n | n | n | n | n | n |
| Tacc3 | transforming, acidic coiled-co | 21335 | MGI:1341163 | protein coding | -0.505 | 0.000767667 | n | n | n | y | n | n |
| Pmf1 | polyamine-modulated factor 1 | 67037 | MGI:1914287 | protein coding | -0.506 | 0.000314398 | n | n | n | n | n | n |
| Mycl | v-myc avian myelocytomatosi | 16918 | MGI:96799 | protein coding | -0.507 | 2.90E-05 | n | n | n | y | n | n |
| Pou3f4 | POU domain, class 3, transcr | 18994 | MGI:101894 | protein coding | -0.511 | 0.000240401 | n | n | n | n | y | n |
| Cdk8 | cyclin dependent kinase 8 | 264064 | MGI:1196224 | protein coding | -0.511 | 0.00024587 | n | n | n | y | n | n |
| Aebp1 | AE binding protein 1 | 11568 | MGI:1197012 | protein coding | -0.512 | 5.04E-05 | n | n | n | n | n | n |
| Rcan3 | regulator of calcineurin 3 | 53902 | MGI:1858220 | protein coding | -0.517 | 0.000114071 | n | n | n | n | n | n |
| Cnrip1 | cannabinoid receptor interacti | 380686 | MGI:1917505 | protein coding | -0.520 | 0.000355395 | n | n | n | n | n | n |
| Grxcr1 | glutaredoxin, cysteine rich 1 | 433899 | MGI:3577767 | protein coding | -0.521 | 0.000213385 | n | n | y | y | y | y |
| Atp1a2 | ATPase, Na+/K+ transporting | 98660 | MGI:88106 | protein coding | -0.522 | 1.38E-05 | n | n | n | n | n | n |
| Bod1 | biorientation of chromosomes | 69556 | MGI:1916806 | protein coding | -0.528 | 0.000736153 | n | n | n | n | n | n |
| Pcna | proliferating cell nuclear antig | 18538 | MGI:97503 | protein coding | -0.532 | 0.000798956 | n | n | n | n | n | n |
| Hlf | hepatic leukemia factor | 217082 | MGI:96108 | protein coding | -0.535 | 0.000147582 | n | n | n | n | n | n |
| Pla2g7 | phospholipase A2, group VII ( | 27226 | MGI:1351327 | protein coding | -0.536 | 2.29E-06 | n | n | n | n | n | n |
| Tspan17 | tetraspanin 17 | 74257 | MGI:1921507 | protein coding | -0.539 | 1.82E-05 | n | n | n | n | n | n |
| Eno1 | enolase 1, alpha non-neuron | 13806 | MGI:95393 | protein coding | -0.540 | 1.39E-05 | n | n | n | n | n | n |
| Oga | O-GlcNAcase | 76055 | MGI:1932139 | protein coding | -0.541 | 0.000832284 | n | n | n | n | n | n |
| Kif2a | kinesin family member 2A | 16563 | MGI:108390 | protein coding | -0.543 | 0.000162339 | n | n | n | n | n | y |
| Bub1b | BUB1B, mitotic checkpoint se | 12236 | MGI:1333889 | protein coding | -0.544 | 0.000502308 | n | n | n | n | n | n |
| Dpysl3 | dihydropyrimidinase-like 3 | 22240 | MGI:1349762 | protein coding | -0.544 | 7.82E-05 | n | n | y | y | n | n |
| Ephx4 | epoxide hydrolase 4 | 384214 | MGI:2686228 | protein coding | -0.545 | 0.00046171 | n | n | n | n | n | n |
| Csrp1 | cysteine and glycine-rich prot | 13007 | MGI:88549 | protein coding | -0.549 | 0.000482778 | n | n | n | y | n | n |
| Trim45 | tripartite motif-containing 45 | 229644 | MGI:1918187 | protein coding | -0.550 | 7.52E-05 | n | n | y | n | n | n |
| Itpr3 | inositol 1,4,5-triphosphate rec | 16440 | MGI:96624 | protein coding | -0.551 | 0.000116742 | n | n | n | y | n | n |

| | | | | | | | | | | | | |
|---|---|---|---|---|---|---|---|---|---|---|---|---|
| Phlda3 | pleckstrin homology like doma | 27280 | MGI:1351485 | protein coding | -0.552 | 0.000906689 | n | n | n | n | n | n |
| Ndufaf6 | NADH:ubiquinone oxidoreduc | 76947 | MGI:1924197 | protein coding | -0.552 | 0.000275432 | n | n | y | y | n | n |
| Cdkn1c | cyclin dependent kinase inhib | 12577 | MGI:104564 | protein coding | -0.553 | 0.000744451 | n | n | n | n | n | n |
| Calb2 | calbindin 2 | 12308 | MGI:101914 | protein coding | -0.558 | 3.87E-07 | n | n | y | y | n | n |
| Ube2j2 | ubiquitin-conjugating enzyme | 140499 | MGI:2153608 | protein coding | -0.559 | 2.46E-05 | n | n | n | n | n | n |
| Trmt6 | tRNA methyltransferase 6 | 66926 | MGI:1914176 | protein coding | -0.560 | 1.92E-05 | n | n | n | n | n | n |
| Tbk1 | TANK-binding kinase 1 | 56480 | MGI:1929658 | protein coding | -0.562 | 0.00035406 | n | n | n | n | n | n |
| Lsm4 | LSM4 homolog, U6 small nuc | 50783 | MGI:1354692 | protein coding | -0.567 | 4.45E-06 | n | n | n | n | n | n |
| Inafm1 | InaF motif containing 1 | 66300 | MGI:1913550 | protein coding | -0.568 | 5.50E-05 | n | n | n | n | n | n |
| Timp3 | tissue inhibitor of metalloprote | 21859 | MGI:98754 | protein coding | -0.569 | 8.34E-05 | n | n | n | y | n | n |
| Flt1 | FMS-like tyrosine kinase 1 | 14254 | MGI:95558 | protein coding | -0.569 | 0.000359142 | n | n | n | n | n | n |
| Pdhb | pyruvate dehydrogenase (lipo | 68263 | MGI:1915513 | protein coding | -0.569 | 6.60E-06 | n | n | n | n | n | n |
| Rbpms | RNA binding protein gene wit | 19663 | MGI:1334446 | protein coding | -0.572 | 0.000344064 | n | n | n | n | n | n |
| Snrpb2 | U2 small nuclear ribonucleop | 20639 | MGI:104805 | protein coding | -0.573 | 1.51E-05 | n | n | n | y | n | n |
| Morf4l1 | mortality factor 4 like 1 | 21761 | MGI:1096551 | protein coding | -0.576 | 0.000176768 | n | n | n | n | n | n |
| Ppm1m | protein phosphatase 1M | 67905 | MGI:1915155 | protein coding | -0.577 | 0.000117342 | n | n | n | n | n | n |
| Rpl36 | ribosomal protein L36 | 54217 | MGI:1860603 | protein coding | -0.578 | 0.000570584 | n | n | n | n | n | n |
| Pttg1 | pituitary tumor-transforming g | 30939 | MGI:1353578 | protein coding | -0.580 | 0.000778934 | n | n | n | n | n | n |
| Efemp1 | epidermal growth factor-conta | 216616 | MGI:1339998 | protein coding | -0.588 | 5.63E-05 | n | n | n | n | n | n |
| Gypc | glycophorin C | 71683 | MGI:1098566 | protein coding | -0.591 | 0.000514195 | n | n | n | n | n | n |
| Atp8a2 | ATPase, aminophospholipid t | 50769 | MGI:1354710 | protein coding | -0.592 | 0.000673666 | n | n | n | n | y** (PMID: 24413176) | n |
| Fam210b | family with sequence similarit | 67017 | MGI:1914267 | protein coding | -0.593 | 2.69E-05 | n | n | n | n | n | n |
| Polg2 | polymerase (DNA directed), g | 50776 | MGI:1354947 | protein coding | -0.595 | 0.000353981 | n | n | n | n | n | n |
| Cnep1r1 | CTD nuclear envelope phosp | 382030 | MGI:1921981 | protein coding | -0.599 | 0.000181516 | n | n | n | n | n | n |
| Dgat1 | diacylglycerol O-acyltransfera | 13350 | MGI:1333825 | protein coding | -0.601 | 3.63E-06 | n | n | n | n | n | n |
| Col6a2 | collagen, type VI, alpha 2 | 12834 | MGI:88460 | protein coding | -0.605 | 0.000240567 | n | n | n | n | n | n |
| Tmem216 | transmembrane protein 216 | 68642 | MGI:1920020 | protein coding | -0.607 | 1.39E-05 | y | n | n | n | n | y |
| Ajuba | ajuba LIM protein | 16475 | MGI:1341886 | protein coding | -0.609 | 0.000162868 | n | n | n | n | n | n |
| Acp1 | acid phosphatase 1, soluble | 11431 | MGI:87881 | protein coding | -0.610 | 0.000461705 | n | n | n | n | n | n |
| Plaat3 | phospholipase A and acyltran | 225845 | MGI:2179715 | protein coding | -0.610 | 1.06E-05 | n | n | n | y | n | n |
| Arrdc4 | arrestin domain containing 4 | 66412 | MGI:1913662 | protein coding | -0.611 | 9.30E-06 | n | n | n | y | n | n |
| B2m | beta-2 microglobulin | 12010 | MGI:88127 | protein coding | -0.612 | 0.000254308 | n | n | n | n | n | n |
| Notch2 | notch 2 | 18129 | MGI:97364 | protein coding | -0.615 | 0.000113531 | n | n | n | n | n | y |
| Bgn | biglycan | 12111 | MGI:88158 | protein coding | -0.621 | 0.000484644 | n | n | n | n | n | n |
| Smtn | smoothelin | 29856 | MGI:1354727 | protein coding | -0.622 | 0.000144272 | n | n | n | n | n | n |
| Cfap20 | cilia and flagella associated p | 14894 | MGI:107428 | protein coding | -0.629 | 1.14E-05 | n | n | n | n | n | y |
| Lats2 | large tumor suppressor 2 | 50523 | MGI:1354386 | protein coding | -0.629 | 0.000577129 | n | n | n | n | n | n |
| Col5a2 | collagen, type V, alpha 2 | 12832 | MGI:88458 | protein coding | -0.631 | 2.64E-05 | n | n | n | n | n | n |
| Ppm1j | protein phosphatase 1J | 71887 | MGI:1919137 | protein coding | -0.634 | 0.000258248 | y | n | n | n | n | n |
| Slc38a3 | solute carrier family 38, meml | 76257 | MGI:1923507 | protein coding | -0.635 | 1.16E-05 | n | n | n | n | n | n |
| Pcolce | procollagen C-endopeptidase | 18542 | MGI:105099 | protein coding | -0.636 | 0.000300982 | n | n | n | n | n | n |
| Rtraf | RNA transcription, translation | 68045 | MGI:1915295 | protein coding | -0.639 | 2.28E-06 | n | n | n | y | n | n |
| Tesc | tescalcin | 57816 | MGI:1930803 | protein coding | -0.643 | 1.03E-05 | n | n | n | y | n | n |
| Endod1 | endonuclease domain contair | 71946 | MGI:1919196 | protein coding | -0.648 | 1.23E-05 | n | n | y | n | n | n |
| Atp6v0c | ATPase, H+ transporting, lysc | 11984 | MGI:88116 | protein coding | -0.649 | 0.000624366 | n | n | n | n | n | n |
| Urah | urate (5-hydroxyiso-) hydrolas | 76974 | MGI:1916142 | protein coding | -0.651 | 2.23E-05 | n | n | n | n | n | n |
| Aqp1 | aquaporin 1 | 11826 | MGI:103201 | protein coding | -0.652 | 0.000272028 | n | n | n | n | n | n |
| Frg2f1 | FSHD region gene 2 family m | 433752 | MGI:3035485 | protein coding | -0.654 | 6.46E-05 | n | n | n | n | n | n |
| Axin1 | axin 1 | 12005 | MGI:1096327 | protein coding | -0.656 | 0.000368589 | n | n | n | n | y | n |
| Sfn | stratifin | 55948 | MGI:1891831 | protein coding | -0.657 | 0.00024897 | n | n | n | n | n | n |
| Cdk15 | cyclin dependent kinase 15 | 271697 | MGI:3583944 | protein coding | -0.661 | 0.000673666 | n | n | y | y | n | n |
| Cd83 | CD83 antigen | 12522 | MGI:1328316 | protein coding | -0.663 | 0.000102797 | n | n | n | n | n | n |
| Rhno1 | RAD9-HUS1-RAD1 interactin | 72440 | MGI:1915315 | protein coding | -0.666 | 6.24E-08 | n | n | n | n | n | n |
| Nkd1 | naked cuticle 1 | 93960 | MGI:2135954 | protein coding | -0.668 | 1.52E-05 | n | n | n | y | n | n |
| Otor | otoraplin | 57329 | MGI:1888678 | protein coding | -0.669 | 9.88E-05 | n | n | n | n | n | n |
| Wipf3 | WAS/WASL interacting protei | 330319 | MGI:3044681 | protein coding | -0.673 | 0.000840784 | n | n | n | n | n | n |
| Igfbp2 | insulin-like growth factor bindi | 16008 | MGI:96437 | protein coding | -0.677 | 6.77E-05 | n | n | n | n | n | n |
| Dnm3 | dynamin 3 | 103967 | MGI:1341299 | protein coding | -0.678 | 7.99E-05 | n | y | n | n | n | n |
| Fam149a | family with sequence similarit | 212326 | MGI:2387177 | protein coding | -0.679 | 6.50E-06 | n | n | n | n | n | n |
| St8sia3 | ST8 alpha-N-acetyl-neuramin | 20451 | MGI:106019 | protein coding | -0.681 | 2.54E-06 | n | n | y | y | n | n |
| Fkbp7 | FK506 binding protein 7 | 14231 | MGI:1336879 | protein coding | -0.681 | 0.000264043 | n | n | n | n | n | n |
| Cd248 | CD248 antigen, endosialin | 70445 | MGI:1917695 | protein coding | -0.682 | 6.06E-05 | n | n | n | y | n | n |
| Olfm1 | olfactomedin 1 | 56177 | MGI:1860437 | protein coding | -0.695 | 1.71E-05 | n | n | y | y | n | n |
| Zfp365 | zinc finger protein 365 | 216049 | MGI:2143676 | protein coding | -0.697 | 4.45E-06 | n | n | n | n | n | n |
| Hnrnpdl | heterogeneous nuclear ribonu | 50926 | MGI:1355299 | protein coding | -0.698 | 1.86E-05 | n | n | n | n | n | n |
| Nrsn1 | neurensin 1 | 22360 | MGI:894662 | protein coding | -0.699 | 1.41E-05 | y | n | n | n | n | n |
| Peli2 | pellino 2 | 93834 | MGI:1891445 | protein coding | -0.702 | 5.79E-07 | n | n | n | y | n | n |

| | | | | | | | | | | | | |
|---|---|---|---|---|---|---|---|---|---|---|---|---|
| Ifitm3 | interferon induced transmemb | 66141 | MGI:1913391 | protein coding | -0.703 | 2.09E-05 | n | n | n | n | n | n |
| Trh | thyrotropin releasing hormone | 22044 | MGI:98823 | protein coding | -0.708 | 3.80E-05 | n | n | n | y | n | n |
| Syce2 | synaptonemal complex centra | 71846 | MGI:1919096 | protein coding | -0.720 | 3.79E-06 | n | n | n | n | n | n |
| Rab3c | RAB3C, member RAS oncog | 67295 | MGI:1914545 | protein coding | -0.722 | 1.80E-05 | n | n | n | n | n | n |
| Synj2 | synaptojanin 2 | 20975 | MGI:1201671 | protein coding | -0.725 | 1.45E-05 | y | n | n | n | y** (PMID: 21423608, 33100973) | n |
| Spon2 | spondin 2, extracellular matrix | 100689 | MGI:1923724 | protein coding | -0.725 | 2.05E-05 | n | n | n | n | n | n |
| Tex30 | testis expressed 30 | 75623 | MGI:1922873 | protein coding | -0.725 | 0.000139658 | n | n | n | n | n | n |
| Mycbpap | MYCBP associated protein | 104601 | MGI:2388726 | protein coding | -0.729 | 6.24E-07 | n | y | n | n | n | n |
| Tgfbi | transforming growth factor, be | 21810 | MGI:99959 | protein coding | -0.730 | 9.10E-05 | n | n | n | n | n | n |
| Atp6v1c2 | ATPase, H+ transporting, lyso | 68775 | MGI:1916025 | protein coding | -0.733 | 0.000848434 | n | y | n | n | n | n |
| Ttll7 | tubulin tyrosine ligase-like fan | 70892 | MGI:1918142 | protein coding | -0.733 | 0.00025105 | n | y | n | y | n | y |
| Kcp | kielin/chordin-like protein | 333088 | MGI:2141640 | protein coding | -0.735 | 5.15E-05 | n | n | y | n | n | n |
| Tbx18 | T-box18 | 76365 | MGI:1923615 | protein coding | -0.738 | 4.31E-05 | n | n | n | n | y | n |
| Apod | apolipoprotein D | 11815 | MGI:88056 | protein coding | -0.740 | 0.000441035 | n | n | n | n | n | n |
| Rad54l2 | RAD54 like 2 (S. cerevisiae) | 81000 | MGI:1933196 | protein coding | -0.741 | 0.000125812 | n | n | n | n | n | n |
| Tuba1b | tubulin, alpha 1B | 22143 | MGI:107804 | protein coding | -0.742 | 0.000318522 | n | n | n | y | n | n |
| Lat2 | linker for activation of T cells f | 56743 | MGI:1926479 | protein coding | -0.744 | 0.000415615 | n | n | n | n | n | n |
| Pcdhb9 | protocadherin beta 9 | 93880 | MGI:2136744 | protein coding | -0.744 | 0.00076909 | n | n | n | n | n | n |
| Calm2 | calmodulin 2 | 12314 | MGI:103250 | protein coding | -0.747 | 2.23E-05 | n | n | n | n | n | n |
| Bok | BCL2-related ovarian killer | 51800 | MGI:1858494 | protein coding | -0.754 | 0.000208936 | n | n | n | n | n | n |
| Haus4 | HAUS augmin-like complex, s | 219072 | MGI:1261794 | protein coding | -0.756 | 1.23E-05 | n | n | n | n | n | n |
| Tspan4 | tetraspanin 4 | 64540 | MGI:1928097 | protein coding | -0.757 | 3.00E-05 | n | n | n | n | n | n |
| Rnps1 | RNA binding protein with serii | 19826 | MGI:97960 | protein coding | -0.758 | 1.67E-06 | n | n | n | n | n | n |
| Sbsn | suprabasin | 282619 | MGI:2446326 | protein coding | -0.759 | 0.000705585 | n | n | n | n | n | n |
| Slc22a18 | solute carrier family 22 (organ | 18400 | MGI:1336884 | protein coding | -0.763 | 0.000250473 | n | n | n | n | n | n |
| Hmga2 | high mobility group AT-hook 2 | 15364 | MGI:101761 | protein coding | -0.776 | 0.000848434 | n | n | n | n | n | n |
| Col26a1 | collagen, type XXVI, alpha 1 | 140709 | MGI:2155345 | protein coding | -0.777 | 2.17E-05 | n | n | n | y | n | n |
| Snrnp25 | small nuclear ribonucleoprote | 78372 | MGI:1925622 | protein coding | -0.780 | 3.82E-06 | n | n | y | n | n | n |
| Pnp | purine-nucleoside phosphoryl | 18950 | MGI:97365 | protein coding | -0.786 | 1.12E-05 | n | n | n | n | n | n |
| Tfb2m | transcription factor B2, mitoch | 15278 | MGI:107937 | protein coding | -0.794 | 0.00060529 | n | n | n | n | n | n |
| Hdac1 | histone deacetylase 1 | 433759 | MGI:108086 | protein coding | -0.801 | 2.96E-06 | n | n | n | n | n | n |
| Kctd6 | potassium channel tetrameris | 71393 | MGI:1918643 | protein coding | -0.808 | 0.00012412 | n | n | n | n | n | n |
| Tcf7l2 | transcription factor 7 like 2, T | 21416 | MGI:1202879 | protein coding | -0.808 | 0.000955152 | n | n | n | n | n | n |
| Phgdh | 3-phosphoglycerate dehydrog | 236539 | MGI:1355330 | protein coding | -0.811 | 4.91E-09 | n | n | n | n | n | n |
| Fhod3 | formin homology 2 domain co | 225288 | MGI:1925847 | protein coding | -0.827 | 2.24E-05 | n | n | y | n | n | n |
| Tpm2 | tropomyosin 2, beta | 22004 | MGI:98810 | protein coding | -0.829 | 7.57E-06 | n | n | n | n | n | n |
| Frmpd1 | FERM and PDZ domain conta | 666060 | MGI:2446274 | protein coding | -0.843 | 5.62E-06 | n | n | n | y | n | n |
| Znrf3 | zinc and ring finger 3 | 407821 | MGI:3039616 | protein coding | -0.844 | 2.16E-07 | n | n | n | n | n | n |
| Fez1 | fasciculation and elongation p | 235180 | MGI:2670976 | protein coding | -0.855 | 1.36E-08 | n | n | y | y | n | n |
| Efcc1 | EF hand and coiled-coil doma | 58229 | MGI:3611451 | protein coding | -0.857 | 2.63E-05 | n | n | n | n | n | n |
| Abca6 | ATP-binding cassette, sub-fai | 76184 | MGI:1923434 | protein coding | -0.857 | 0.000731922 | n | n | n | n | n | n |
| Ucp3 | uncoupling protein 3 (mitocho | 22229 | MGI:1099787 | protein coding | -0.864 | 0.000236087 | n | y | n | n | n | n |
| Kcnn4 | potassium intermediate/small | 16534 | MGI:1277957 | protein coding | -0.864 | 0.000925473 | n | n | n | n | n | n |
| Paqr7 | progestin and adipoQ recepto | 71904 | MGI:1919154 | protein coding | -0.878 | 0.000207342 | n | n | n | n | n | n |
| Pvr | poliovirus receptor | 52118 | MGI:107741 | protein coding | -0.879 | 0.000559102 | n | n | n | n | n | n |
| Casq1 | calsequestrin 1 | 12372 | MGI:1309468 | protein coding | -0.879 | 1.58E-05 | n | n | n | n | n | n |
| Rtp4 | receptor transporter protein 4 | 67775 | MGI:1915025 | protein coding | -0.884 | 6.72E-06 | n | n | n | n | n | n |
| Reep6 | receptor accessory protein 6 | 70335 | MGI:1917585 | protein coding | -0.894 | 7.24E-08 | n | n | n | n | n | n |
| Cbr3 | carbonyl reductase 3 | 109857 | MGI:1309992 | protein coding | -0.896 | 0.00020919 | n | n | n | n | n | n |
| Cdkn1b | cyclin dependent kinase inhib | 12576 | MGI:104565 | protein coding | -0.896 | 1.48E-06 | n | n | n | n | y | n |
| Bid | BH3 interacting domain death | 12122 | MGI:108093 | protein coding | -0.898 | 2.16E-07 | n | n | n | n | n | n |
| Lacc1 | laccase domain containing 1 | 210808 | MGI:2445077 | protein coding | -0.905 | 6.21E-06 | n | n | n | n | n | n |
| Zfp991 | zinc finger protein 991 | 666532 | MGI:3701604 | protein coding | -0.907 | 0.000185425 | n | n | n | n | n | n |
| A930009A15Rik | RIKEN cDNA A930009A15 ge | 77798 | MGI:1925048 | protein coding | -0.908 | 4.43E-06 | y | n | n | y | n | n |
| Scara5 | scavenger receptor class A, n | 71145 | MGI:1918395 | protein coding | -0.913 | 2.74E-05 | n | n | n | n | n | n |
| Sgip1 | SH3-domain GRB2-like (endo | 73094 | MGI:1920344 | protein coding | -0.914 | 4.40E-06 | n | n | n | n | n | n |
| Klhl15 | kelch-like 15 | 236904 | MGI:1923400 | protein coding | -0.917 | 0.000164023 | n | n | n | n | n | n |
| Pvalb | parvalbumin | 19293 | MGI:97821 | protein coding | -0.921 | 6.64E-08 | n | n | y | y | n | n |
| Tagap1 | T cell activation GTPase activ | 380608 | MGI:1919786 | protein coding | -0.924 | 0.000369362 | n | n | n | n | n | n |
| Parp3 | poly (ADP-ribose) polymerase | 235587 | MGI:1891258 | protein coding | -0.938 | 3.86E-05 | n | n | n | n | n | n |
| Otof | otoferlin | 83762 | MGI:1891247 | protein coding | -0.938 | 6.18E-06 | y | n | n | n | y | n |
| Zfp383 | zinc finger protein 383 | 73729 | MGI:1920979 | protein coding | -0.942 | 5.86E-05 | n | n | n | n | n | n |
| Sos2 | SOS Ras/Rho guanine nucleo | 20663 | MGI:98355 | protein coding | -0.947 | 0.00025026 | n | n | n | n | n | n |
| Agtrap | angiotensin II, type I receptor- | 11610 | MGI:1339977 | protein coding | -0.947 | 4.09E-09 | n | n | n | y | n | n |
| Ccdc122 | coiled-coil domain containing | 108811 | MGI:1918358 | protein coding | -0.952 | 4.51E-06 | n | n | n | n | n | n |
| Zbtb8a | zinc finger and BTB domain c | 73680 | MGI:1920930 | protein coding | -0.954 | 9.57E-05 | n | n | n | n | n | n |
| Ncan | neurocan | 13004 | MGI:104694 | protein coding | -0.957 | 1.24E-06 | n | n | n | n | n | n |

| | | | | | | | | | | | | |
|---|---|---|---|---|---|---|---|---|---|---|---|---|
| Gas2l3 | growth arrest-specific 2 like 3 | 237436 | MGI:1918780 | protein coding | -0.959 | 1.78E-07 | n | n | n | n | n | n |
| Ncoa4 | nuclear receptor coactivator 4 | 27057 | MGI:1350932 | protein coding | -0.977 | 2.62E-06 | n | n | n | n | n | n |
| Wdhd1 | WD repeat and HMG-box DN | 218973 | MGI:2443514 | protein coding | -0.977 | 0.000902981 | n | n | n | n | n | n |
| Pnma2 | paraneoplastic antigen MA2 | 239157 | MGI:2444129 | protein coding | -0.979 | 7.33E-05 | n | n | y | y | n | n |
| Hmgb2 | high mobility group box 2 | 97165 | MGI:96157 | protein coding | -0.990 | 0.000172489 | n | n | n | n | n | n |
| Xkr8 | X-linked Kx blood group relat | 381560 | MGI:2685877 | protein coding | -0.993 | 1.47E-05 | n | n | n | n | n | n |
| Bst2 | bone marrow stromal cell anti | 69550 | MGI:1916800 | protein coding | -0.996 | 1.51E-05 | n | n | n | n | n | n |
| Tmem178 | transmembrane protein 178 | 68027 | MGI:1915277 | protein coding | -0.996 | 0.000333809 | n | n | n | n | n | n |
| Hnrnpa0 | heterogeneous nuclear ribonu | 77134 | MGI:1924384 | protein coding | -1.003 | 0.000435205 | n | n | n | n | n | n |
| Gm15217 | predicted gene 15217 | 100041724 | MGI:3705233 | protein coding | -1.003 | 0.000751472 | n | n | n | n | n | n |
| Spmip1 | sperm microtubule inner prote | 668210 | MGI:3644212 | protein coding | -1.006 | 0.000109997 | n | n | n | n | n | n |
| Trmt9b | tRNA methyltransferase 9B | 319582 | MGI:2442328 | protein coding | -1.015 | 0.000731922 | n | n | n | y | n | n |
| Sh2d4a | SH2 domain containing 4A | 72281 | MGI:1919531 | protein coding | -1.028 | 4.67E-06 | n | n | y | y | n | n |
| Tlcd3b | TLC domain containing 3B | 68952 | MGI:1916202 | protein coding | -1.033 | 5.40E-07 | n | n | y | n | n | n |
| Chgb | chromogranin B | 12653 | MGI:88395 | protein coding | -1.042 | 1.14E-07 | n | n | y | y | n | n |
| Zic2 | zinc finger protein of the cerel | 22772 | MGI:106679 | protein coding | -1.047 | 0.000446777 | n | n | n | n | n | n |
| Abhd14b | abhydrolase domain containir | 76491 | MGI:1923741 | protein coding | -1.056 | 4.94E-09 | n | n | n | n | n | n |
| Matn1 | matrilin 1, cartilage matrix pro | 17180 | MGI:106591 | protein coding | -1.064 | 0.000235336 | n | n | n | n | n | n |
| Pak6 | p21 (RAC1) activated kinase | 214230 | MGI:2679420 | protein coding | -1.073 | 0.000162545 | n | n | n | n | n | n |
| Slc4a11 | solute carrier family 4, sodiur | 269356 | MGI:2138987 | protein coding | -1.080 | 4.63E-05 | n | n | n | n | n | n |
| Mmrn2 | multimerin 2 | 105450 | MGI:2385618 | protein coding | -1.081 | 0.000453006 | n | n | n | n | n | n |
| Apobec2 | apolipoprotein B mRNA editin | 11811 | MGI:1343178 | protein coding | -1.082 | 0.000939185 | n | n | n | n | n | n |
| A530016L24Rik | RIKEN cDNA A530016L24 ge | 319942 | MGI:2443020 | protein coding | -1.093 | 7.12E-05 | y | n | n | y | n | n |
| Dnah8 | dynein, axonemal, heavy cha | 13417 | MGI:107714 | protein coding | -1.096 | 3.02E-06 | n | n | n | n | n | y |
| Zfp950 | zinc finger protein 950 | 414758 | MGI:2652824 | protein coding | -1.097 | 9.86E-06 | n | n | n | n | n | n |
| Insig1 | insulin induced gene 1 | 231070 | MGI:1916289 | protein coding | -1.103 | 0.000160417 | n | n | n | n | n | n |
| Rpgrip1 | retinitis pigmentosa GTPase r | 77945 | MGI:1932134 | protein coding | -1.112 | 0.000427669 | n | n | n | n | n | y |
| Crls1 | cardiolipin synthase 1 | 66586 | MGI:1913836 | protein coding | -1.117 | 0.000303012 | n | n | n | n | n | n |
| Rnaset2b | ribonuclease T2B | 68195 | MGI:3702087 | protein coding | -1.118 | 0.000273977 | n | n | n | n | n | n |
| Eif3j1 | eukaryotic translation initiatio | 78655 | MGI:1925905 | protein coding | -1.121 | 0.000226104 | n | n | n | n | n | n |
| Nf2 | neurofibromin 2 | 18016 | MGI:97307 | protein coding | -1.125 | 3.98E-09 | n | n | y | n | n | n |
| Dhtkd1 | dehydrogenase E1 and transl | 209692 | MGI:2445096 | protein coding | -1.125 | 6.29E-05 | n | n | n | n | n | n |
| Lig1 | ligase I, DNA, ATP-dependen | 16881 | MGI:101789 | protein coding | -1.133 | 7.17E-08 | y | n | n | n | n | n |
| Necab2 | N-terminal EF-hand calcium b | 117148 | MGI:2152211 | protein coding | -1.140 | 8.87E-07 | y | n | n | y | n | n |
| Gpr62 | G protein-coupled receptor 62 | 436090 | MGI:3525078 | protein coding | -1.193 | 0.000976708 | n | n | n | n | n | n |
| Rps2 | ribosomal protein S2 | 16898 | MGI:105110 | protein coding | -1.201 | 8.75E-05 | n | n | n | n | n | n |
| Gpr176 | G protein-coupled receptor 17 | 381413 | MGI:2685858 | protein coding | -1.202 | 0.000756559 | n | n | n | n | n | n |
| Prom2 | prominin 2 | 192212 | MGI:2138997 | protein coding | -1.217 | 7.17E-08 | n | n | n | n | n | y |
| Rpl3 | ribosomal protein L3 | 27367 | MGI:1351605 | protein coding | -1.237 | 9.33E-07 | n | n | n | n | n | n |
| Gm14305 | predicted gene 14305 | 100043387 | MGI:3709632 | protein coding | -1.250 | 0.000392635 | n | n | n | n | n | n |
| Oxsm | 3-oxoacyl-ACP synthase, mit | 71147 | MGI:1918397 | protein coding | -1.255 | 0.000505407 | n | n | n | n | n | n |
| Casz1 | castor zinc finger 1 | 69743 | MGI:1196251 | protein coding | -1.267 | 5.91E-05 | n | n | y | y | n | n |
| Psmb5 | proteasome (prosome, macro | 19173 | MGI:1194513 | protein coding | -1.268 | 3.29E-09 | n | n | n | n | n | n |
| Stmn1 | stathmin 1 | 16765 | MGI:96739 | protein coding | -1.275 | 5.36E-06 | n | n | n | n | n | n |
| Cmtm5 | CKLF-like MARVEL transmer | 67272 | MGI:2447164 | protein coding | -1.276 | 7.24E-08 | n | n | n | n | n | n |
| Gfod1 | glucose-fructose oxidoreduct | 328232 | MGI:2145304 | protein coding | -1.290 | 1.82E-05 | n | y | n | y | n | n |
| Miip | migration and invasion inhibit | 28010 | MGI:106506 | protein coding | -1.295 | 1.34E-09 | n | n | n | n | n | n |
| Chit1 | chitinase 1 | 71884 | MGI:1919134 | protein coding | -1.302 | 0.000236314 | n | n | n | y | n | n |
| Unc5a | unc-5 netrin receptor A | 107448 | MGI:894682 | protein coding | -1.304 | 0.000105229 | n | n | n | n | n | n |
| Jmjd7 | jumonji domain containing 7 | 433466 | MGI:3845785 | protein coding | -1.309 | 8.63E-09 | n | n | n | n | n | n |
| Bcas1 | brain enriched myelin associa | 76960 | MGI:1924210 | protein coding | -1.319 | 1.51E-06 | n | n | n | y | n | n |
| Htr3a | 5-hydroxytryptamine (seroton | 15561 | MGI:96282 | protein coding | -1.325 | 4.34E-07 | n | y | n | n | n | n |
| Grp | gastrin releasing peptide | 225642 | MGI:95833 | protein coding | -1.328 | 1.31E-07 | n | y | n | y | n | n |
| Plaat1 | phospholipase A and acyltran | 27281 | MGI:1351473 | protein coding | -1.364 | 4.56E-07 | n | n | n | y | n | n |
| Ppp1r14a | protein phosphatase 1, regula | 68458 | MGI:1931139 | protein coding | -1.397 | 7.27E-06 | n | n | y | y | n | n |
| Mocs1 | molybdenum cofactor synthes | 56738 | MGI:1928904 | protein coding | -1.410 | 8.73E-08 | n | n | n | n | n | n |
| Fgl1 | fibrinogen-like protein 1 | 234199 | MGI:102795 | protein coding | -1.414 | 0.00026818 | n | n | n | n | n | n |
| Kalrn | kalirin, RhoGEF kinase | 545156 | MGI:2685385 | protein coding | -1.422 | 1.73E-05 | n | y | n | y | n | n |
| Art5 | ADP-ribosyltransferase 5 | 11875 | MGI:107948 | protein coding | -1.423 | 1.49E-05 | n | n | y | n | n | n |
| Ssc5d | scavenger receptor cysteine r | 269855 | MGI:3606211 | protein coding | -1.426 | 0.000182715 | n | n | n | n | n | n |
| Cel | carboxyl ester lipase | 12613 | MGI:88374 | protein coding | -1.442 | 2.47E-05 | n | n | n | n | n | n |
| Gm7932 | predicted gene 7932 | 666105 | MGI:3643446 | protein coding | -1.452 | 0.000250473 | n | n | n | n | n | n |
| Gnmt | glycine N-methyltransferase | 14711 | MGI:1202304 | protein coding | -1.455 | 1.90E-06 | n | n | y | n | n | n |
| Tuba8 | tubulin, alpha 8 | 53857 | MGI:1858225 | protein coding | -1.476 | 1.95E-05 | y | n | n | n | n | n |
| Cplane2 | ciliogenesis and planar polari | 76166 | MGI:1923416 | protein coding | -1.504 | 1.50E-05 | n | n | n | n | n | y |
| Inafm2 | InaF motif containing 2 | 100043272 | MGI:1915354 | protein coding | -1.516 | 7.99E-05 | n | n | y | n | n | n |
| 4833420G17Rik | RIKEN cDNA 4833420G17 ge | 67392 | MGI:1914642 | protein coding | -1.516 | 1.82E-07 | n | n | n | n | n | n |

| | | | | | | | | | | | | |
|---|---|---|---|---|---|---|---|---|---|---|---|---|
| Bend6 | BEN domain containing 6 | 320705 | MGI:2444572 | protein coding | -1.526 | 4.60E-07 | n | n | n | n | n | n |
| Rin1 | Ras and Rab interactor 1 | 225870 | MGI:2385695 | protein coding | -1.534 | 4.01E-06 | n | n | n | n | n | n |
| Ackr4 | atypical chemokine receptor 4 | 252837 | MGI:2181676 | protein coding | -1.542 | 9.28E-09 | n | n | n | n | n | n |
| Slc13a4 | solute carrier family 13 (sodiu | 243755 | MGI:2442367 | protein coding | -1.546 | 0.000211651 | n | n | n | n | n | n |
| Scn4a | sodium channel, voltage-gate | 110880 | MGI:98250 | protein coding | -1.566 | 0.000258248 | n | n | n | y | n | n |
| Sh3rf2 | SH3 domain containing ring fi | 269016 | MGI:2444628 | protein coding | -1.566 | 1.43E-06 | n | n | n | n | n | n |
| Disp3 | dispatched RND transporter f | 242748 | MGI:2444403 | protein coding | -1.582 | 0.000306411 | n | n | n | n | n | n |
| Slc45a3 | solute carrier family 45, meml | 212980 | MGI:1922082 | protein coding | -1.592 | 3.82E-06 | n | y | n | n | n | n |
| Cldn19 | claudin 19 | 242653 | MGI:3033992 | protein coding | -1.609 | 0.00097844 | n | n | n | n | n | n |
| Cimip3 | ciliary microtubule inner prote | 75462 | MGI:1922712 | protein coding | -1.637 | 6.59E-05 | n | n | n | n | n | n |
| Ets1 | E26 avian leukemia oncogen | 23871 | MGI:95455 | protein coding | -1.673 | 0.00038047 | n | n | n | n | n | n |
| Il11 | interleukin 11 | 16156 | MGI:107613 | protein coding | -1.696 | 7.83E-06 | n | n | n | n | n | n |
| Pkhd1l1 | polycystic kidney and hepatic | 192190 | MGI:2183153 | protein coding | -1.701 | 6.50E-09 | n | y | n | n | y | n |
| A2ml1 | alpha-2-macroglobulin like 1 | 232400 | MGI:3039594 | protein coding | -1.707 | 8.53E-05 | n | n | n | n | n | n |
| Tub | tubby bipartite transcription fa | 22141 | MGI:2651573 | protein coding | -1.708 | 2.16E-07 | n | n | n | n | y | y |
| Zfp386 | zinc finger protein 386 (Krupp | 56220 | MGI:1930708 | protein coding | -1.776 | 7.54E-12 | n | n | n | n | n | n |
| Slc1a6 | solute carrier family 1 (high af | 20513 | MGI:1096331 | protein coding | -1.778 | 1.72E-07 | n | n | n | n | n | n |
| Grid1 | glutamate receptor, ionotropic | 14803 | MGI:95812 | protein coding | -1.811 | 3.69E-08 | n | n | n | y | n | n |
| Ang4 | angiogenin, ribonuclease A fa | 219033 | MGI:2656551 | protein coding | -1.813 | 0.000228782 | n | n | n | n | n | n |
| Mpz | myelin protein zero | 17528 | MGI:103177 | protein coding | -1.829 | 2.75E-05 | n | n | n | n | n | n |
| Zfp46 | zinc finger protein 46 | 22704 | MGI:99192 | protein coding | -1.844 | 1.54E-07 | n | n | n | n | n | n |
| Mamdc2 | MAM domain containing 2 | 71738 | MGI:1918988 | protein coding | -1.852 | 3.92E-07 | y | n | n | n | n | n |
| Rprm | reprimo, TP53 dependent G2 | 67874 | MGI:1915124 | protein coding | -1.853 | 1.50E-13 | n | n | y | y | n | n |
| Alox5 | arachidonate 5-lipoxygenase | 11689 | MGI:87999 | protein coding | -1.884 | 1.23E-05 | n | n | n | y | n | n |
| Il20rb | interleukin 20 receptor beta | 213208 | MGI:2143266 | protein coding | -1.938 | 0.000132661 | n | n | n | n | n | n |
| Serpina3g | serine (or cysteine) peptidase | 20715 | MGI:105046 | protein coding | -1.944 | 0.000208936 | n | n | n | n | n | n |
| Cyp4f18 | cytochrome P450, family 4, su | 72054 | MGI:1919304 | protein coding | -1.971 | 7.99E-05 | n | n | n | n | n | n |
| Slc1a7 | solute carrier family 1 (glutam | 242607 | MGI:2444087 | protein coding | -1.986 | 0.000168829 | n | n | n | y | n | n |
| Styxl2 | serine/threonine/tyrosine inte | 240892 | MGI:2685055 | protein coding | -2.010 | 9.64E-10 | n | n | y | n | n | n |
| Gnb1 | guanine nucleotide binding pr | 14688 | MGI:95781 | protein coding | -2.024 | 1.05E-11 | n | n | n | n | n | y |
| Pdpn | podoplanin | 14726 | MGI:103098 | protein coding | -2.055 | 1.05E-11 | n | n | n | n | n | n |
| Chrna1 | cholinergic receptor nicotinic a | 11435 | MGI:87885 | protein coding | -2.083 | 2.21E-11 | y | n | n | y | n | n |
| Slc6a20a | solute carrier family 6 (neurot | 102680 | MGI:2143217 | protein coding | -2.124 | 0.000218701 | n | n | n | n | n | n |
| Slc34a3 | solute carrier family 34 (sodiu | 142681 | MGI:2159410 | protein coding | -2.140 | 9.26E-08 | y | n | n | n | n | n |
| Rab6b | RAB6B, member RAS oncog | 270192 | MGI:107283 | protein coding | -2.157 | 3.14E-10 | n | n | n | y | n | n |
| Gm14295 | predicted gene 14295 | 100039123 | MGI:3709624 | protein coding | -2.222 | 3.38E-05 | n | n | n | n | n | n |
| Tekt5 | tektin 5 | 70426 | MGI:1917676 | protein coding | -2.243 | 9.36E-06 | n | n | n | n | n | y |
| Ndufs5 | NADH:ubiquinone oxidoreduc | 595136 | MGI:1890889 | protein coding | -2.255 | 1.28E-08 | n | n | n | n | n | n |
| Kif21b | kinesin family member 21B | 16565 | MGI:109234 | protein coding | -2.352 | 6.06E-09 | n | n | y | n | n | n |
| Nuggc | nuclear GTPase, germinal ce | 100503545 | MGI:2685446 | protein coding | -2.353 | 2.54E-06 | n | n | n | n | n | n |
| Hmgn2 | high mobility group nucleosor | 15331 | MGI:96136 | protein coding | -2.372 | 8.40E-07 | n | n | n | n | n | n |
| Samd8 | sterile alpha motif domain cor | 67630 | MGI:1914880 | protein coding | -2.376 | 2.04E-05 | n | n | n | y | n | n |
| Hpca | hippocalcin | 15444 | MGI:1336200 | protein coding | -2.429 | 2.04E-07 | n | n | n | n | n | n |
| Htr1a | 5-hydroxytryptamine (seroton | 15550 | MGI:96273 | protein coding | -2.537 | 0.000108162 | n | n | n | n | n | n |
| Atp2a3 | ATPase, Ca++ transporting, u | 53313 | MGI:1194503 | protein coding | -2.576 | 1.75E-12 | y | n | n | n | n | n |
| Gpr12 | G-protein coupled receptor 12 | 14738 | MGI:101909 | protein coding | -2.758 | 2.92E-08 | n | n | n | n | n | n |
| Tac4 | tachykinin 4 | 93670 | MGI:1931130 | protein coding | -3.016 | 1.04E-06 | n | y | n | n | n | n |
| Zfp979 | zinc finger protein 979 | 112422 | MGI:2148252 | protein coding | -3.073 | 1.75E-05 | n | n | n | n | n | n |
| Rhbdl2 | rhomboid like 2 | 230726 | MGI:3608413 | protein coding | -3.471 | 3.68E-08 | n | n | n | n | n | n |
| Tnfrsf8 | tumor necrosis factor recepto | 21941 | MGI:99908 | protein coding | -4.086 | 7.67E-07 | n | n | n | n | n | n |
| Ghsr | growth hormone secretagogu | 208188 | MGI:2441906 | protein coding | -4.594 | 3.43E-06 | n | n | n | y | n | n |
| Rpl29 | ribosomal protein L29 | 19944 | MGI:99687 | protein coding | -4.969 | 4.32E-05 | n | n | n | n | n | n |
| Emx1 | empty spiracles homeobox 1 | 13796 | MGI:95387 | protein coding | -5.321 | 3.30E-09 | n | n | n | n | n | n |
| Rbl2 | RB transcriptional corepresso | 19651 | MGI:105085 | protein coding | 0.479 | 0.001000805 | n | n | n | y | n | n |
| Ube2ql1 | ubiquitin-conjugating enzyme | 76980 | MGI:1924230 | protein coding | -0.634 | 0.001001086 | n | n | y | n | n | n |
| Creb3l4 | cAMP responsive element bir | 78284 | MGI:1916603 | protein coding | 0.498 | 0.001001086 | n | n | n | n | n | n |
| Dalrd3 | DALR anticodon binding dom | 67789 | MGI:1915039 | protein coding | 0.329 | 0.001001752 | n | n | y | n | n | n |
| Dmc1 | DNA meiotic recombinase 1 | 13404 | MGI:105393 | protein coding | 1.112 | 0.001002932 | n | n | n | n | n | n |
| Pld5 | phospholipase D family meml | 319455 | MGI:2442056 | protein coding | 0.488 | 0.001002932 | n | n | n | n | n | n |
| Rab11fip5 | RAB11 family interacting prot | 52055 | MGI:1098586 | protein coding | -0.513 | 0.001003904 | n | n | n | n | n | n |
| G3bp2 | G3BP stress granule assemb | 23881 | MGI:2442040 | protein coding | 0.302 | 0.00100548 | n | n | n | n | n | n |
| Xylb | xylulokinase homolog (H. influ | 102448 | MGI:2142985 | protein coding | 0.707 | 0.001007915 | n | n | n | n | n | n |
| Pfkl | phosphofructokinase, liver, B- | 18641 | MGI:97547 | protein coding | 0.319 | 0.001016241 | n | n | n | n | n | n |
| Depdc5 | DEP domain containing 5 | 277854 | MGI:2141101 | protein coding | 0.556 | 0.00102466 | n | n | n | n | n | n |
| Htra4 | HtrA serine peptidase 4 | 330723 | MGI:3036260 | protein coding | -0.665 | 0.00102466 | n | n | n | y | n | n |
| Wdr47 | WD repeat domain 47 | 99512 | MGI:2139593 | protein coding | 0.456 | 0.001028818 | n | n | n | y | n | n |
| Anxa1 | annexin A1 | 16952 | MGI:96819 | protein coding | -0.344 | 0.001038354 | n | n | n | n | n | y |
