## Supplementary material for "CASZ1 regulates the rate at which outer hair cells mature and is required for hearing": Table S3

**Table S3. Groups of genes expressed predominantly in OHCs, IHCs, or OHCs and IHCs during first postnatal week.**

| Gene symbol | Gene name | Entrez gene ID | MGI gene ID | logFC *Casz1(fl/fl);Tg(Sox10-Cre)* vs WT | FDR-adjusted P (Benjamini-Yekutieli method) | Expressed predominantly in OHCs in organ of Corti? y/n | Expressed predominantly in IHCs in organ of Corti? y/n | Expressed predominantly in OHCs *and* IHCs (vs non-HCs) in organ of Corti? y/n | Feature type | |
|---|---|---|---|---|---|---|---|---|---|---|
| Tac4 | tachykinin 4 | 93670 | MGI:1931130 | -3.016 | 1.03768E-06 | y | n | n | protein coding gene | *not detected (ND) |
| Pkhd1l1 | polycystic kidney and hepatic disease 1-lil | 192190 | MGI:2183153 | -1.701 | 6.49542E-09 | y | n | n | protein coding gene | **Predomninantly OHC expressed based on PMID: 30305733 |
| Slc45a3 | solute carrier family 45, member 3 | 212980 | MGI:1922082 | -1.592 | 3.82187E-06 | y | n | n | protein coding gene | |
| Kalrn | kalirin, RhoGEF kinase | 545156 | MGI:2685385 | -1.422 | 1.72862E-05 | y | n | n | protein coding gene | |
| Grp | gastrin releasing peptide | 225642 | MGI:95833 | -1.328 | 1.30536E-07 | y | n | n | protein coding gene | |
| Htr3a | 5-hydroxytryptamine (serotonin) receptor : | 15561 | MGI:96282 | -1.325 | 4.33575E-07 | y | n | n | protein coding gene | |
| Gfod1 | glucose-fructose oxidoreductase domain c | 328232 | MGI:2145304 | -1.290 | 1.8202E-05 | y | n | n | protein coding gene | |
| Ucp3 | uncoupling protein 3 (mitochondrial, proto | 22229 | MGI:1099787 | -0.864 | 0.000236087 | y | n | n | protein coding gene | |
| Ttll7 | tubulin tyrosine ligase-like family, member | 70892 | MGI:1918142 | -0.733 | 0.00025105 | y | n | n | protein coding gene | |
| Atp6v1c2 | ATPase, H+ transporting, lysosomal V1 su | 68775 | MGI:1916025 | -0.733 | 0.000848434 | y | n | n | protein coding gene | |
| Mycbpap | MYCBP associated protein | 104601 | MGI:2388726 | -0.729 | 6.24396E-07 | y | n | n | protein coding gene | |
| Dnm3 | dynamin 3 | 103967 | MGI:1341299 | -0.678 | 7.99203E-05 | y | n | n | protein coding gene | |
| Fam214a | atos homolog A | 235493 | MGI:2387648 | 0.436 | 0.000346078 | y | n | n | protein coding gene | |
| Strip2 | striatin interacting protein 2 | 320609 | MGI:2444363 | 0.445 | 0.000236087 | y | n | n | protein coding gene | |
| Ercc6 | excision repair cross-complementing rode | 319955 | MGI:1100494 | 0.450 | 0.000971013 | y | n | n | protein coding gene | |
| Sugp2 | SURP and G patch domain containing 2 | 234373 | MGI:2678085 | 0.462 | 0.000470676 | y | n | n | protein coding gene | |
| Map2 | microtubule-associated protein 2 | 17756 | MGI:97175 | 0.505 | 0.000226324 | y | n | n | protein coding gene | |
| Kndc1 | kinase non-catalytic C-lobe domain (KIND | 76484 | MGI:1923734 | 0.531 | 0.000236087 | y | n | n | protein coding gene | |
| Car12 | carbonic anhydrase 12 | 76459 | MGI:1923709 | 0.544 | 0.000791453 | y | n | n | protein coding gene | |
| Nim1k | NIM1 serine/threonine protein kinase | 245269 | MGI:2442399 | 0.568 | 0.000449066 | y | n | n | protein coding gene | |
| Acad11 | acyl-Coenzyme A dehydrogenase family, | 102632 | MGI:2143169 | 0.571 | 4.46477E-05 | y | n | n | protein coding gene | |
| Tmtc4 | transmembrane and tetratricopeptide repe | 70551 | MGI:1921050 | 0.597 | 2.89877E-06 | y | n | n | protein coding gene | |
| Mapkbp1 | mitogen-activated protein kinase binding p | 26390 | MGI:1347004 | 0.604 | 2.09179E-05 | y | n | n | protein coding gene | |
| Fanca | Fanconi anemia, complementation group . | 14087 | MGI:1341823 | 0.609 | 4.22751E-05 | y | n | n | protein coding gene | |
| Veph1 | ventricular zone expressed PH domain-co | 72789 | MGI:1920039 | 0.689 | 1.48851E-06 | y | n | n | protein coding gene | |
| Dytn | dystrotelin | 241073 | MGI:2685061 | 0.746 | 3.78915E-06 | y | n | n | protein coding gene | |
| Dbndd1 | dysbindin domain containing 1 | 72185 | MGI:1919435 | 0.787 | 4.90257E-06 | y | n | n | protein coding gene | |
| Cacng5 | calcium channel, voltage-dependent, gam | 140723 | MGI:2157946 | 0.887 | 2.59539E-05 | y | n | n | protein coding gene | |
| Myo1h | myosin 1H | 231646 | MGI:1914674 | 0.905 | 6.21486E-06 | y | n | n | protein coding gene | |
| Nos1ap | nitric oxide synthase 1 (neuronal) adaptor | 70729 | MGI:1917979 | 0.920 | 7.8874E-06 | y | n | n | protein coding gene | |
| Efna5 | ephrin A5 | 13640 | MGI:107444 | 0.932 | 5.45231E-06 | y | n | n | protein coding gene | |
| Elfn1 | leucine rich repeat and fibronectin type III, | 243312 | MGI:2442479 | 0.966 | 4.66767E-06 | y | n | n | protein coding gene | |
| Donson | downstream neighbor of SON | 60364 | MGI:1890621 | 1.209 | 6.21214E-08 | y | n | n | protein coding gene | |
| Laptm5 | lysosomal-associated protein transmembr | 16792 | MGI:108046 | 1.255 | 1.16041E-08 | y | n | n | protein coding gene | |
| Ptgir | prostaglandin I receptor (IP) | 19222 | MGI:99535 | 1.296 | 4.05111E-07 | y | n | n | protein coding gene | |
| Ppip5k1 | diphosphoinositol pentakisphosphate kina | 327655 | MGI:2443281 | 1.355 | 7.22073E-09 | y | n | n | protein coding gene | |
| AI593442 | expressed sequence AI593442 | 330941 | MGI:2143099 | 1.370 | 1.25813E-07 | y | n | n | protein coding gene | |
| Grk1 | G protein-coupled receptor kinase 1 | 24013 | MGI:1345146 | 1.544 | 2.17026E-05 | y | n | n | protein coding gene | |
| Cacng2 | calcium channel, voltage-dependent, gam | 12300 | MGI:1316660 | 1.613 | 3.7042E-07 | y | n | n | protein coding gene | |
| Coro2a | coronin, actin binding protein 2A | 107684 | MGI:1345966 | 1.655 | 8.72675E-08 | y | n | n | protein coding gene | |
| Ppp1r14d | protein phosphatase 1, regulatory inhibitor | 72112 | MGI:1919362 | 1.664 | 0.000131243 | y | n | n | protein coding gene | |
| Cyp2j12 | cytochrome P450, family 2, subfamily j, po | 242546 | MGI:3717097 | 1.672 | 2.5808E-07 | y | n | n | protein coding gene | |
| Rph3a | rabphilin 3A | 19894 | MGI:102788 | 1.777 | 1.40713E-07 | y | n | n | protein coding gene | |
| Slc39a2 | solute carrier family 39 (zinc transporter), | 214922 | MGI:2684326 | 2.212 | 1.31012E-07 | y | n | n | protein coding gene | |
| Nppa | natriuretic peptide type A | 230899 | MGI:97367 | 2.357 | 2.93947E-06 | y | n | n | protein coding gene | |
| Nsg2 | neuron specific gene family member 2 | 18197 | MGI:1202070 | 2.503 | 2.47702E-05 | y | n | n | protein coding gene | |
| Ocm | oncomodulin | 18261 | MGI:97401 | 3.006 | 2.6102E-10 | y | n | n | protein coding gene | |
| Insm2 ** | insulinoma-associated 2 ** | 56856 ** | MGI:1930787 ** | 3.039** | 6.52568274E-06** | y** | n** | n** | protein coding gene** | |
| Serpina1c | serine (or cysteine) peptidase inhibitor, cla | 20702 | MGI:891969 | 10.207 | 1.01817E-10 | y | n | n | protein coding gene | |
| Calca | calcitonin/calcitonin-related polypeptide, a | 12310 | MGI:2151253 | -0.780 | 0.002421952 | y | n | n | protein coding gene | |
| Six2 | sine oculis-related homeobox 2 | 20472 | MGI:102778 | -0.778 | 0.082085564 | y | n | n | protein coding gene | |
| Tmem145 | transmembrane protein 145 | 330485 | MGI:3607779 | -0.660 | 0.010044403 | y | n | n | protein coding gene | |
| Pdlim1 | PDZ and LIM domain 1 (elfin) | 54132 | MGI:1860611 | -0.631 | 0.019458036 | y | n | n | protein coding gene | |
| Gigyf1 | GRB10 interacting GYF protein 1 | 57330 | MGI:1888677 | -0.503 | 0.001294985 | y | n | n | protein coding gene | |
| Rnf19b | ring finger protein 19B | 75234 | MGI:1922484 | -0.493 | 0.006277903 | y | n | n | protein coding gene | |
| Adamts13 | ADAM metallopeptidase with thrombospor | 279028 | MGI:2685556 | -0.486 | 0.011189162 | y | n | n | protein coding gene | |
| Cep63 | centrosomal protein 63 | 28135 | MGI:2158560 | -0.469 | 0.003264211 | y | n | n | protein coding gene | |
| Jakmip1 | janus kinase and microtubule interacting p | 76071 | MGI:1923321 | -0.427 | 0.032485856 | y | n | n | protein coding gene | |

| | | | | | | | | | |
|---|---|---|---|---|---|---|---|---|---|
| Stx1a | syntaxin 1A (brain) | 20907 | MGI:109355 | -0.425 | 0.001753823 | y | n | n | protein coding gene |
| Ifit1 | interferon-induced protein with tetratricope | 15957 | MGI:99450 | -0.392 | 0.002287566 | y | n | n | protein coding gene |
| Vwc2 | von Willebrand factor C domain containing | 319922 | MGI:2442987 | -0.355 | 1 | y | n | n | protein coding gene |
| Tmem38a | transmembrane protein 38A | 74166 | MGI:1921416 | -0.343 | 0.007828064 | y | n | n | protein coding gene |
| Baiap2l2 | BAI1-associated protein 2-like 2 | 207495 | MGI:2652819 | -0.340 | 0.301438431 | y | n | n | protein coding gene |
| Dusp8 | dual specificity phosphatase 8 | 18218 | MGI:106626 | -0.329 | 0.026453454 | y | n | n | protein coding gene |
| Tprn | taperin | 97031 | MGI:2139535 | -0.304 | 0.067451503 | y | n | n | protein coding gene |
| Shroom3 | shroom family member 3 | 27428 | MGI:1351655 | -0.304 | 0.540957303 | y | n | n | protein coding gene |
| Tlk1 | tousled-like kinase 1 | 228012 | MGI:2441683 | -0.300 | 0.005791364 | y | n | n | protein coding gene |
| Plk4 | polo like kinase 4 | 20873 | MGI:101783 | -0.281 | 0.346425611 | y | n | n | protein coding gene |
| 4930558C23Rik | cortexin domain containing 2 | 67654 | MGI:1914904 | -0.272 | 0.336058788 | y | n | n | protein coding gene |
| Coro2b | coronin, actin binding protein, 2B | 235431 | MGI:2444283 | -0.235 | 0.045860454 | y | n | n | protein coding gene |
| Rapgef1 | Rap guanine nucleotide exchange factor ( | 107746 | MGI:104580 | -0.216 | 0.017881104 | y | n | n | protein coding gene |
| Rbfox3 | RNA binding protein, fox-1 homolog (C. el | 52897 | MGI:106368 | -0.216 | 0.13711716 | y | n | n | protein coding gene |
| Mslnl | mesothelin-like | 328783 | MGI:3607710 | -0.192 | 1 | y | n | n | protein coding gene |
| Pald1 | phosphatase domain containing, paladin 1 | 27355 | MGI:1351623 | -0.182 | 0.050613127 | y | n | n | protein coding gene |
| Bcl9 | B cell CLL/lymphoma 9 | 77578 | MGI:1924828 | -0.172 | 1 | y | n | n | protein coding gene |
| Kif13a | kinesin family member 13A | 16553 | MGI:1098264 | -0.170 | 1 | y | n | n | protein coding gene |
| Cds1 | CDP-diacylglycerol synthase 1 | 74596 | MGI:1921846 | -0.146 | 0.89676792 | y | n | n | protein coding gene |
| Phldb1 | pleckstrin homology like domain, family B, | 102693 | MGI:2143230 | -0.143 | 0.764924851 | y | n | n | protein coding gene |
| Arhgap1 | Rho GTPase activating protein 1 | 228359 | MGI:2445003 | -0.140 | 0.201298927 | y | n | n | protein coding gene |
| Osbpl9 | oxysterol binding protein-like 9 | 100273 | MGI:1923784 | -0.128 | 0.314349046 | y | n | n | protein coding gene |
| Lasp1 | LIM and SH3 protein 1 | 16796 | MGI:109656 | -0.119 | 0.687697833 | y | n | n | protein coding gene |
| Pgf | placental growth factor | 18654 | MGI:105095 | -0.113 | 1 | y | n | n | protein coding gene |
| Camsap3 | calmodulin regulated spectrin-associated | 69697 | MGI:1916947 | -0.113 | 1 | y | n | n | protein coding gene |
| Tmem218 | transmembrane protein 218 | 66279 | MGI:1913529 | -0.109 | 1 | y | n | n | protein coding gene |
| Sh2d4b | SH2 domain containing 4B | 328381 | MGI:1925182 | -0.106 | 1 | y | n | n | protein coding gene |
| Zfp618 | zinc finger protein 618 | 72701 | MGI:1919950 | -0.102 | 1 | y | n | n | protein coding gene |
| Mpc1 | mitochondrial pyruvate carrier 1 | 55951 | MGI:1915240 | -0.101 | 1 | y | n | n | protein coding gene |
| Tppp | tubulin polymerization promoting protein | 72948 | MGI:1920198 | -0.097 | 1 | y | n | n | protein coding gene |
| Tmem91 | transmembrane protein 91 | 320208 | MGI:2443589 | -0.094 | 1 | y | n | n | protein coding gene |
| Trp53inp1 | transformation related protein 53 inducible | 60599 | MGI:1926609 | -0.092 | 1 | y | n | n | protein coding gene |
| Palld | palladin, cytoskeletal associated protein | 72333 | MGI:1919583 | -0.091 | 1 | y | n | n | protein coding gene |
| Prkcd | protein kinase C, delta | 18753 | MGI:97598 | -0.070 | 1 | y | n | n | protein coding gene |
| Zfp423 | zinc finger protein 423 | 94187 | MGI:1891217 | -0.067 | 1 | y | n | n | protein coding gene |
| Gpcpd1 | glycerophosphocholine phosphodiesteras | 74182 | MGI:104898 | -0.067 | 1 | y | n | n | protein coding gene |
| Kit | KIT proto-oncogene receptor tyrosine kina | 16590 | MGI:96677 | -0.060 | 1 | y | n | n | protein coding gene |
| Klhl25 | kelch-like 25 | 207952 | MGI:2668031 | -0.057 | 1 | y | n | n | protein coding gene |
| Pard3b | par-3 family cell polarity regulator beta | 72823 | MGI:1919301 | -0.054 | 1 | y | n | n | protein coding gene |
| Aifm3 | apoptosis-inducing factor, mitochondrion-a | 72168 | MGI:1919418 | -0.052 | 1 | y | n | n | protein coding gene |
| Strc | stereocilin | 140476 | MGI:2153816 | -0.047 | 1 | y | n | n | protein coding gene |
| Gcnt2 | glucosaminyl (N-acetyl) transferase 2 (I bl | 14538 | MGI:1100870 | -0.046 | 1 | y | n | n | protein coding gene |
| Tmem59l | transmembrane protein 59-like | 67937 | MGI:1915187 | -0.038 | 1 | y | n | n | protein coding gene |
| Wrap73 | WD repeat containing, antisense to Trp73 | 59002 | MGI:1891749 | -0.037 | 1 | y | n | n | protein coding gene |
| Tmem120a | transmembrane protein 120A | 215210 | MGI:2686991 | -0.034 | 1 | y | n | n | protein coding gene |
| Disc1 | disrupted in schizophrenia 1 | 244667 | MGI:2447658 | -0.027 | 1 | y | n | n | protein coding gene |
| Sstr2 | somatostatin receptor 2 | 20606 | MGI:98328 | -0.019 | 1 | y | n | n | protein coding gene |
| 0610030E20Rik | RIKEN cDNA 0610030E20 gene | 68364 | MGI:1915614 | -0.011 | 1 | y | n | n | protein coding gene |
| Cbwd1 | Zn regulated GTPase metalloprotein activ | 226043 | MGI:2385089 | -0.009 | 1 | y | n | n | protein coding gene |
| Gpr27 | G protein-coupled receptor 27 | 14761 | MGI:1202299 | -0.007 | 1 | y | n | n | protein coding gene |
| Ablim3 | actin binding LIM protein family, member 3 | 319713 | MGI:2442582 | -0.004 | 1 | y | n | n | protein coding gene |
| Bcor | BCL6 interacting corepressor | 71458 | MGI:1918708 | -0.003 | 1 | y | n | n | protein coding gene |
| Mknk1 | MAP kinase-interacting serine/threonine k | 17346 | MGI:894316 | 0.016 | 1 | y | n | n | protein coding gene |
| Ano10 | anoctamin 10 | 102566 | MGI:2143103 | 0.023 | 1 | y | n | n | protein coding gene |
| Gan | giant axonal neuropathy | 209239 | MGI:1890619 | 0.034 | 1 | y | n | n | protein coding gene |
| Tmem163 | transmembrane protein 163 | 72160 | MGI:1919410 | 0.037 | 1 | y | n | n | protein coding gene |
| Hdac11 | histone deacetylase 11 | 232232 | MGI:2385252 | 0.038 | 1 | y | n | n | protein coding gene |
| Tmem229a | transmembrane protein 229A | 319832 | MGI:2442812 | 0.043 | 1 | y | n | n | protein coding gene |
| Sox6 | SRY (sex determining region Y)-box 6 | 20679 | MGI:98368 | 0.051 | 1 | y | n | n | protein coding gene |
| Cdr2 | cerebellar degeneration-related 2 | 12585 | MGI:1100885 | 0.051 | 1 | y | n | n | protein coding gene |
| Slc43a1 | solute carrier family 43, member 1 | 72401 | MGI:1931352 | 0.054 | 1 | y | n | n | protein coding gene |
| Herc2 | HECT and RLD domain containing E3 ubi | 15204 | MGI:103234 | 0.055 | 1 | y | n | n | protein coding gene |
| Agl | amylo-1,6-glucosidase, 4-alpha-glucanotra | 77559 | MGI:1924809 | 0.059 | 1 | y | n | n | protein coding gene |
| Cep44 | centrosomal protein 44 | 382010 | MGI:3525111 | 0.064 | 1 | y | n | n | protein coding gene |

| | | | | | | | | | |
|---|---|---|---|---|---|---|---|---|---|
| Mboat7 | membrane bound O-acyltransferase doma | 77582 | MGI:1924832 | 0.072 | 1 | y | n | n | protein coding gene |
| Atp2b2 | ATPase, Ca++ transporting, plasma mem | 11941 | MGI:105368 | 0.078 | 1 | y | n | n | protein coding gene |
| Mef2a | myocyte enhancer factor 2A | 17258 | MGI:99532 | 0.093 | 1 | y | n | n | protein coding gene |
| Fggy | FGGY carbohydrate kinase domain conta | 75578 | MGI:1922828 | 0.107 | 1 | y | n | n | protein coding gene |
| Ptcd1 | pentatricopeptide repeat domain 1 | 71799 | MGI:1919049 | 0.115 | 1 | y | n | n | protein coding gene |
| Gnao1 | guanine nucleotide binding protein, alpha | 14681 | MGI:95775 | 0.117 | 1 | y | n | n | protein coding gene |
| Mmp23 | matrix metallopeptidase 23 | 26561 | MGI:1347361 | 0.119 | 1 | y | n | n | protein coding gene |
| Sh3pxd2a | SH3 and PX domains 2A | 14218 | MGI:1298393 | 0.121 | 1 | y | n | n | protein coding gene |
| Cog1 | component of oligomeric golgi complex 1 | 16834 | MGI:1333873 | 0.123 | 0.568760285 | y | n | n | protein coding gene |
| Tmprss7 | transmembrane serine protease 7 | 208171 | MGI:2686594 | 0.136 | 1 | y | n | n | protein coding gene |
| Atp8a1 | ATPase phospholipid transporting 8A1 | 11980 | MGI:1330848 | 0.136 | 0.851466615 | y | n | n | protein coding gene |
| Synpo | synaptopodin | 104027 | MGI:1099446 | 0.140 | 1 | y | n | n | protein coding gene |
| Adarb1 | adenosine deaminase, RNA-specific, B1 | 110532 | MGI:891999 | 0.143 | 1 | y | n | n | protein coding gene |
| Drosha | drosha, ribonuclease type III | 14000 | MGI:1261425 | 0.146 | 0.875762463 | y | n | n | protein coding gene |
| Mical3 | microtubule associated monooxygenase, | 194401 | MGI:2442733 | 0.149 | 1 | y | n | n | protein coding gene |
| Aox2 | aldehyde oxidase 2 | 213043 | MGI:3529596 | 0.162 | 1 | y | n | n | protein coding gene |
| Serinc3 | serine incorporator 3 | 26943 | MGI:1349457 | 0.164 | 1 | y | n | n | protein coding gene |
| Hsd17b7 | hydroxysteroid (17-beta) dehydrogenase | 15490 | MGI:1330808 | 0.170 | 0.401094651 | y | n | n | protein coding gene |
| Zfp503 | zinc finger protein 503 | 218820 | MGI:1353644 | 0.184 | 1 | y | n | n | protein coding gene |
| Bcat2 | branched chain aminotransferase 2, mitoc | 12036 | MGI:1276534 | 0.186 | 0.19656288 | y | n | n | protein coding gene |
| Hlcs | holocarboxylase synthetase (biotin- [propr | 110948 | MGI:894646 | 0.189 | 0.173107391 | y | n | n | protein coding gene |
| Map4k2 | mitogen-activated protein kinase kinase ki | 26412 | MGI:1346883 | 0.195 | 0.237057863 | y | n | n | protein coding gene |
| Ampd3 | adenosine monophosphate deaminase 3 | 11717 | MGI:1096344 | 0.195 | 0.926068085 | y | n | n | protein coding gene |
| Grk3 | G protein-coupled receptor kinase 3 | 320129 | MGI:87941 | 0.203 | 0.052925698 | y | n | n | protein coding gene |
| Bdp1 | B double prime 1, subunit of RNA polymer | 544971 | MGI:1347077 | 0.218 | 0.440633195 | y | n | n | protein coding gene |
| Gse1 | genetic suppressor element 1, coiled-coil | 382034 | MGI:1098275 | 0.225 | 0.536303672 | y | n | n | protein coding gene |
| Tecpr1 | tectonin beta-propeller repeat containing 1 | 70381 | MGI:1917631 | 0.226 | 0.004899373 | y | n | n | protein coding gene |
| Mfsd4a | major facilitator superfamily domain conta | 213006 | MGI:2442786 | 0.261 | 1 | y | n | n | protein coding gene |
| Igsf21 | immunoglobulin superfamily, member 21 | 230868 | MGI:2681842 | 0.261 | 1 | y | n | n | protein coding gene |
| Naa16 | N(alpha)-acetyltransferase 16, NatA auxili | 66897 | MGI:1914147 | 0.267 | 0.057944601 | y | n | n | protein coding gene |
| Ikzf2 | IKAROS family zinc finger 2 | 22779 | MGI:1342541 | 0.269 | 0.94804301 | y | n | n | protein coding gene |
| Acsl1 | acyl-CoA synthetase long-chain family me | 14081 | MGI:102797 | 0.275 | 0.035260579 | y | n | n | protein coding gene |
| Ypel1 | yippee like 1 | 106369 | MGI:1913303 | 0.279 | 0.012138929 | y | n | n | protein coding gene |
| Fbxl20 | F-box and leucine-rich repeat protein 20 | 72194 | MGI:1919444 | 0.293 | 0.001476111 | y | n | n | protein coding gene |
| Pfkfb4 | 6-phosphofructo-2-kinase/fructose-2,6-bip | 270198 | MGI:2687284 | 0.309 | 0.005141427 | y | n | n | protein coding gene |
| Septin4 | septin 4 | 18952 | MGI:1270156 | 0.323 | 0.02255227 | y | n | n | protein coding gene |
| Fer1l6 | fer-1 like family member 6 | 631797 | MGI:3645398 | 0.337 | 0.056156032 | y | n | n | protein coding gene |
| Leng1 | leukocyte receptor cluster (LRC) member | 69757 | MGI:1917007 | 0.338 | 0.024862413 | y | n | n | protein coding gene |
| Cep78 | centrosomal protein 78 | 208518 | MGI:1924386 | 0.339 | 0.079164367 | y | n | n | protein coding gene |
| Eif2ak4 | eukaryotic translation initiation factor 2 alp | 27103 | MGI:1353427 | 0.365 | 0.004369472 | y | n | n | protein coding gene |
| Sipa1l3 | signal-induced proliferation-associated 1 l | 74206 | MGI:1921456 | 0.367 | 0.225382057 | y | n | n | protein coding gene |
| Bach2 | BTB and CNC homology, basic leucine zip | 12014 | MGI:894679 | 0.390 | 0.055076096 | y | n | n | protein coding gene |
| Ccdc92 | coiled-coil domain containing 92 | 215707 | MGI:106485 | 0.394 | 0.002297433 | y | n | n | protein coding gene |
| Lrrk1 | leucine-rich repeat kinase 1 | 233328 | MGI:2142227 | 0.405 | 0.00191099 | y | n | n | protein coding gene |
| Pkn3 | protein kinase N3 | 263803 | MGI:2388285 | 0.406 | 0.214596787 | y | n | n | protein coding gene |
| Dock4 | dedicator of cytokinesis 4 | 238130 | MGI:1918006 | 0.409 | 0.040008638 | y | n | n | protein coding gene |
| Tmtc1 | transmembrane and tetratricopeptide repe | 387314 | MGI:3039590 | 0.458 | 0.002256286 | y | n | n | protein coding gene |
| Plekhg6 | pleckstrin homology domain containing, fa | 213522 | MGI:2682298 | 0.459 | 0.00265095 | y | n | n | protein coding gene |
| Bcl11b | B cell leukemia/lymphoma 11B | 58208 | MGI:1929913 | 0.493 | 0.12521077 | y | n | n | protein coding gene |
| Fgf21 | fibroblast growth factor 21 | 56636 | MGI:1861377 | 0.574 | 0.155114208 | y | n | n | protein coding gene |
| P2rx3 | purinergic receptor P2X, ligand-gated ion | 228139 | MGI:1097160 | 0.595 | 0.377894956 | y | n | n | protein coding gene |
| Ush2a | usherin | 22283 | MGI:1341292 | 0.672 | 0.010014608 | y | n | n | protein coding gene |
| Dll1 | delta like canonical Notch ligand 1 | 13388 | MGI:104659 | 0.772 | 0.002232668 | y | n | n | protein coding gene |
| Prcd | photoreceptor disc component | 100038570 | MGI:3649529 | 0.861 | 0.008157974 | y | n | n | protein coding gene |
| Slc47a2 | solute carrier family 47, member 2 | 380701 | MGI:3588190 | 0.871 | 0.020772433 | y | n | n | protein coding gene |
| Cdyl2 | chromodomain protein, Y chromosome-lik | 75796 | MGI:1923046 | 0.964 | 0.015065784 | y | n | n | protein coding gene |
| Slc26a5 | solute carrier family 26, member 5 | 80979 | MGI:1933154 | 1.022 | 0.005192689 | y | n | n | protein coding gene |
| Spns3 | SPNS lysolipid transporter 3, sphingosine | 77577 | MGI:1924827 | 1.043 | 0.026801677 | y | n | n | protein coding gene |
| Insm1 | insulinoma-associated 1 | 53626 | MGI:1859980 | ND* | ND* | y | n | n | protein coding gene |
| Neurod6 | neurogenic differentiation 6 | 11922 | MGI:106593 | ND* | ND* | y | n | n | protein coding gene |
| Pcdh8 | protocadherin 8 | 18530 | MGI:1306800 | ND* | ND* | y | n | n | protein coding gene |
| Dll3 | delta like canonical Notch ligand 3 | 13389 | MGI:1096877 | ND* | ND* | y | n | n | protein coding gene |
| Gpha2 | glycoprotein hormone alpha 2 | 170458 | MGI:2156541 | ND* | ND* | y | n | n | protein coding gene |
| Pax2 | paired box 2 | 18504 | MGI:97486 | ND* | ND* | y | n | n | protein coding gene |

| | | | | | | | | | |
|---|---|---|---|---|---|---|---|---|---|
| Rtn4r | reticulon 4 receptor | 65079 | MGI:2136886 | ND* | ND* | y | n | n | protein coding gene |
| Them7 | thioesterase superfamily member 7 | 74088 | MGI:1921338 | ND* | ND* | y | n | n | protein coding gene |
| Hspa1b | heat shock protein 1B | 15511 | MGI:99517 | ND* | ND* | y | n | n | protein coding gene |
| Ryr1 | ryanodine receptor 1, skeletal muscle | 20190 | MGI:99659 | ND* | ND* | y | n | n | protein coding gene |
| Atp2a3 | ATPase, Ca++ transporting, ubiquitous | 53313 | MGI:1194503 | -2.576 | 1.75073E-12 | n | y | n | protein coding gene |
| Slc34a3 | solute carrier family 34 (sodium phosphate | 142681 | MGI:2159410 | -2.140 | 9.25547E-08 | n | y | n | protein coding gene |
| Chrna1 | cholinergic receptor nicotinic alpha 1 subu | 11435 | MGI:87885 | -2.083 | 2.21234E-11 | n | y | n | protein coding gene |
| Mamdc2 | MAM domain containing 2 | 71738 | MGI:1918988 | -1.852 | 3.91724E-07 | n | y | n | protein coding gene |
| Tuba8 | tubulin, alpha 8 | 53857 | MGI:1858225 | -1.476 | 1.95187E-05 | n | y | n | protein coding gene |
| Necab2 | N-terminal EF-hand calcium binding prote | 117148 | MGI:2152211 | -1.140 | 8.87034E-07 | n | y | n | protein coding gene |
| Lig1 | ligase I, DNA, ATP-dependent | 16881 | MGI:101789 | -1.133 | 7.16722E-08 | n | y | n | protein coding gene |
| A530016L24Rik | RIKEN cDNA A530016L24 gene | 319942 | MGI:2443020 | -1.093 | 7.1186E-05 | n | y | n | protein coding gene |
| Otof | otoferlin | 83762 | MGI:1891247 | -0.938 | 6.18154E-06 | n | y | n | protein coding gene |
| A930009A15Rik | RIKEN cDNA A930009A15 gene | 77798 | MGI:1925048 | -0.908 | 4.42872E-06 | n | y | n | protein coding gene |
| Synj2 | synaptojanin 2 | 20975 | MGI:1201671 | -0.725 | 1.45495E-05 | n | y | n | protein coding gene |
| Nrsn1 | neurensin 1 | 22360 | MGI:894662 | -0.699 | 1.41178E-05 | n | y | n | protein coding gene |
| Ppm1j | protein phosphatase 1J | 71887 | MGI:1919137 | -0.634 | 0.000258248 | n | y | n | protein coding gene |
| Tmem216 | transmembrane protein 216 | 68642 | MGI:1920020 | -0.607 | 1.38825E-05 | n | y | n | protein coding gene |
| Kifap3 | kinesin-associated protein 3 | 16579 | MGI:107566 | 0.329 | 0.000971013 | n | y | n | protein coding gene |
| Xirp2 | xin actin-binding repeat containing 2 | 241431 | MGI:2685198 | 0.562 | 4.99276E-05 | n | y | n | protein coding gene |
| Odad2 | outer dynein arm docking complex subuni | 74934 | MGI:1922184 | 0.576 | 0.000705585 | n | y | n | protein coding gene |
| Pnpla3 | patatin-like phospholipase domain contain | 116939 | MGI:2151796 | 0.662 | 0.000630585 | n | y | n | protein coding gene |
| Pih1d2 | PIH1 domain containing 2 | 72614 | MGI:1919864 | 0.738 | 0.000113531 | n | y | n | protein coding gene |
| Cimap1b | ciliary microtubule associated protein 1B | 70113 | MGI:1917363 | 0.872 | 0.000111872 | n | y | n | protein coding gene |
| Trim36 | tripartite motif-containing 36 | 28105 | MGI:106264 | 0.917 | 0.000425409 | n | y | n | protein coding gene |
| Fscn2 | fascin actin-bundling protein 2 | 238021 | MGI:2443337 | 1.004 | 1.90407E-08 | n | y | n | protein coding gene |
| Pitpnm1 | phosphatidylinositol transfer protein, mem | 18739 | MGI:1197524 | 1.142 | 3.08514E-09 | n | y | n | protein coding gene |
| Chrng | cholinergic receptor, nicotinic, gamma pol | 11449 | MGI:87895 | 1.153 | 4.21003E-05 | n | y | n | protein coding gene |
| Saxo4 | stabilizer of axonemal microtubules 4 | 67752 | MGI:1915002 | 1.159 | 1.51385E-06 | n | y | n | protein coding gene |
| Smim18 | small integral membrane protein 18 | 72632 | MGI:1919882 | 1.424 | 1.8052E-06 | n | y | n | protein coding gene |
| E230025N22Rik | Riken cDNA E230025N22 gene | 240216 | MGI:3687212 | 1.475 | 5.03046E-08 | n | y | n | protein coding gene |
| Wdr95 | WD40 repeat domain 95 | 381693 | MGI:1923042 | 2.911 | 3.75414E-06 | n | y | n | protein coding gene |
| Fcrlb | Fc receptor-like B | 435653 | MGI:3576487 | -0.497 | 0.001099825 | n | y | n | protein coding gene |
| Slc17a8 | solute carrier family 17 (sodium-depender | 216227 | MGI:3039629 | 1.006 | 0.001441647 | n | y | n | protein coding gene |
| Cdk14 | cyclin dependent kinase 14 | 18647 | MGI:894318 | -0.371 | 0.001691711 | n | y | n | protein coding gene |
| Smim1 | small integral membrane protein 1 | 68859 | MGI:1916109 | -0.363 | 0.002138013 | n | y | n | protein coding gene |
| Pgam2 | phosphoglycerate mutase 2 | 56012 | MGI:1933118 | -0.574 | 0.002281303 | n | y | n | protein coding gene |
| Dnm1 | dynamin 1 | 13429 | MGI:107384 | -0.449 | 0.002490555 | n | y | n | protein coding gene |
| Ttc28 | tetratricopeptide repeat domain 28 | 209683 | MGI:2140873 | -0.459 | 0.002831615 | n | y | n | protein coding gene |
| Sult4a1 | sulfotransferase family 4A, member 1 | 29859 | MGI:1888971 | 0.408 | 0.003273093 | n | y | n | protein coding gene |
| Dgkg | diacylglycerol kinase, gamma | 110197 | MGI:105060 | 0.490 | 0.005587271 | n | y | n | protein coding gene |
| Nefl | neurofilament, light polypeptide | 18039 | MGI:97313 | 0.548 | 0.008208396 | n | y | n | protein coding gene |
| B4galt2 | UDP-Gal:betaGlcNAc beta 1,4- galactosyl | 53418 | MGI:1858493 | -0.257 | 0.009335808 | n | y | n | protein coding gene |
| Asphd2 | aspartate beta-hydroxylase domain contai | 72898 | MGI:1920148 | 0.535 | 0.009871938 | n | y | n | protein coding gene |
| Immp2l | IMP2 inner mitochondrial membrane pepti | 93757 | MGI:2135611 | -0.430 | 0.009904114 | n | y | n | protein coding gene |
| Plcxd2 | phosphatidylinositol-specific phospholipas | 433022 | MGI:3647874 | 1.033 | 0.011414785 | n | y | n | protein coding gene |
| Tbc1d16 | TBC1 domain family, member 16 | 207592 | MGI:2652878 | -0.347 | 0.011968115 | n | y | n | protein coding gene |
| Stk32c | serine/threonine kinase 32C | 57740 | MGI:2385336 | -1.092 | 0.016601522 | n | y | n | protein coding gene |
| Slc1a5 | solute carrier family 1 (neutral amino acid | 20514 | MGI:105305 | -0.535 | 0.020247645 | n | y | n | protein coding gene |
| Pcnx4 | pecanex homolog 4 | 67708 | MGI:1914958 | 0.271 | 0.021559099 | n | y | n | protein coding gene |
| Rnf224 | ring finger protein 224 | 329360 | MGI:2685603 | 0.616 | 0.022106636 | n | y | n | protein coding gene |
| Cys1 | cystin 1 | 12879 | MGI:2177632 | -0.767 | 0.022802266 | n | y | n | protein coding gene |
| Nmnat3 | nicotinamide nucleotide adenylyltransfera: | 74080 | MGI:1921330 | 0.358 | 0.028591867 | n | y | n | protein coding gene |
| Rwdd3 | RWD domain containing 3 | 66568 | MGI:1920420 | -0.247 | 0.037516002 | n | y | n | protein coding gene |
| Myl4 | myosin, light polypeptide 4 | 17896 | MGI:97267 | 0.407 | 0.042942676 | n | y | n | protein coding gene |
| Grwd1 | glutamate-rich WD repeat containing 1 | 101612 | MGI:2141989 | -0.191 | 0.056166003 | n | y | n | protein coding gene |
| Kcnh8 | potassium voltage-gated channel, subfam | 211468 | MGI:2445160 | 0.266 | 0.057235404 | n | y | n | protein coding gene |
| Enkur | enkurin, TRPC channel interacting protein | 71233 | MGI:1918483 | 0.344 | 0.066888352 | n | y | n | protein coding gene |
| Spmip6 | sperm microtubule inner protein 6 | 73721 | MGI:1920971 | 0.545 | 0.073032411 | n | y | n | protein coding gene |
| 1190005I06Rik | RIKEN cDNA 1190005I06 gene | 68918 | MGI:1916168 | -0.321 | 0.084243815 | n | y | n | protein coding gene |
| Kcnab2 | potassium voltage-gated channel, shaker- | 16498 | MGI:109239 | 0.281 | 0.098724007 | n | y | n | protein coding gene |
| Kif5c | kinesin family member 5C | 16574 | MGI:1098269 | -0.233 | 0.113500393 | n | y | n | protein coding gene |
| Efl1 | elongation factor like GTPase 1 | 101592 | MGI:2141969 | 0.198 | 0.12521077 | n | y | n | protein coding gene |
| Dpysl2 | dihydropyrimidinase-like 2 | 12934 | MGI:1349763 | -0.218 | 0.177829126 | n | y | n | protein coding gene |

| | | | | | | | | | |
|---|---|---|---|---|---|---|---|---|---|
| Liat1 | ligand of ATE1 | 74230 | MGI:1921480 | 0.331 | 0.186888879 | n | y | n | protein coding gene |
| Tbx2 | T-box 2 | 21385 | MGI:98494 | -0.463 | 0.209233539 | n | y | n | protein coding gene |
| Zfp276 | zinc finger protein (C2H2 type) 276 | 57247 | MGI:1888495 | 0.174 | 0.227428517 | n | y | n | protein coding gene |
| Nup205 | nucleoporin 205 | 70699 | MGI:2141625 | 0.153 | 0.227533548 | n | y | n | protein coding gene |
| Pex5l | peroxisomal biogenesis factor 5-like | 58869 | MGI:1916672 | 0.526 | 0.233979837 | n | y | n | protein coding gene |
| Cracdl | capping protein inhibiting regulator of actir | 72097 | MGI:1919347 | -0.418 | 0.250820416 | n | y | n | protein coding gene |
| Nfasc | neurofascin | 269116 | MGI:104753 | 0.243 | 0.253686174 | n | y | n | protein coding gene |
| Intu | inturned planar cell polarity protein | 380614 | MGI:2443752 | 0.296 | 0.325102688 | n | y | n | protein coding gene |
| Fgf8 | fibroblast growth factor 8 | 14179 | MGI:99604 | -0.241 | 0.335487325 | n | y | n | protein coding gene |
| Nalf1 | NALCN channel auxiliary factor 1 | 270028 | MGI:2142765 | -0.578 | 0.350867422 | n | y | n | protein coding gene |
| Cacna2d2 | calcium channel, voltage-dependent, alph | 56808 | MGI:1929813 | -0.276 | 0.441791553 | n | y | n | protein coding gene |
| Mapk12 | mitogen-activated protein kinase 12 | 29857 | MGI:1353438 | -0.258 | 0.45991977 | n | y | n | protein coding gene |
| Ube2q1 | ubiquitin-conjugating enzyme E2Q family | 70093 | MGI:1917343 | 0.440 | 0.468705165 | n | y | n | protein coding gene |
| Kctd1 | potassium channel tetramerisation domair | 106931 | MGI:1918269 | -0.236 | 0.493088568 | n | y | n | protein coding gene |
| Syt2 | synaptotagmin II | 20980 | MGI:99666 | -0.405 | 0.533365655 | n | y | n | protein coding gene |
| Dusp19 | dual specificity phosphatase 19 | 68082 | MGI:1915332 | 0.179 | 0.637121456 | n | y | n | protein coding gene |
| Scx | scleraxis scleraxis bHLH transcription fact | 20289 | MGI:102934 | -0.222 | 0.658164978 | n | y | n | protein coding gene |
| Plin5 | perilipin 5 | 66968 | MGI:1914218 | -0.197 | 0.815325182 | n | y | n | protein coding gene |
| Tln2 | talin 2 | 70549 | MGI:1917799 | -0.166 | 0.819031561 | n | y | n | protein coding gene |
| Odad1 | outer dynein arm docking complex subuni | 211535 | MGI:2446120 | 0.172 | 0.939087704 | n | y | n | protein coding gene |
| Fxyd2 | FXYD domain-containing ion transport reg | 11936 | MGI:1195260 | -0.366 | 1 | n | y | n | protein coding gene |
| Cab39 | calcium binding protein 39 | 12283 | MGI:107438 | 0.065 | 1 | n | y | n | protein coding gene |
| Eno2 | enolase 2, gamma neuronal | 13807 | MGI:95394 | -0.072 | 1 | n | y | n | protein coding gene |
| Vegfd | vascular endothelial growth factor D | 14205 | MGI:108037 | 0.026 | 1 | n | y | n | protein coding gene |
| Polg | polymerase (DNA directed), gamma | 18975 | MGI:1196389 | 0.016 | 1 | n | y | n | protein coding gene |
| Trp53bp1 | transformation related protein 53 binding p | 27223 | MGI:1351320 | 0.000 | 1 | n | y | n | protein coding gene |
| Cabp2 | calcium binding protein 2 | 29866 | MGI:1352749 | 0.106 | 1 | n | y | n | protein coding gene |
| Fbxw4 | F-box and WD-40 domain protein 4 | 30838 | MGI:1354698 | -0.079 | 1 | n | y | n | protein coding gene |
| Septin1 | septin 1 | 54204 | MGI:1858916 | 0.111 | 1 | n | y | n | protein coding gene |
| Kcnip3 | Kv channel interacting protein 3, calsenilir | 56461 | MGI:1929258 | 0.132 | 1 | n | y | n | protein coding gene |
| Dnajb13 | DnaJ heat shock protein family (Hsp40) m | 69387 | MGI:1916637 | 0.211 | 1 | n | y | n | protein coding gene |
| Draxin | dorsal inhibitory axon guidance protein | 70433 | MGI:1917683 | 0.092 | 1 | n | y | n | protein coding gene |
| Usp38 | ubiquitin specific peptidase 38 | 74841 | MGI:1922091 | -0.008 | 1 | n | y | n | protein coding gene |
| Rilpl1 | Rab interacting lysosomal protein-like 1 | 75695 | MGI:1922945 | -0.037 | 1 | n | y | n | protein coding gene |
| Tanc2 | tetratricopeptide repeat, ankyrin repeat an | 77097 | MGI:2444121 | 0.149 | 1 | n | y | n | protein coding gene |
| Heg1 | heart development protein with EGF-like c | 77446 | MGI:1924696 | -0.002 | 1 | n | y | n | protein coding gene |
| Asb8 | ankyrin repeat and SOCS box-containing | 78541 | MGI:1925791 | -0.048 | 1 | n | y | n | protein coding gene |
| Cpsf1 | cleavage and polyadenylation specific fac | 94230 | MGI:2679722 | 0.043 | 1 | n | y | n | protein coding gene |
| Fbxl14 | F-box and leucine-rich repeat protein 14 | 101358 | MGI:2141676 | -0.158 | 1 | n | y | n | protein coding gene |
| Adi1 | acireductone dioxygenase 1 | 104923 | MGI:2144929 | -0.027 | 1 | n | y | n | protein coding gene |
| Pacs1 | phosphofurin acidic cluster sorting protein | 107975 | MGI:1277113 | -0.024 | 1 | n | y | n | protein coding gene |
| Rbm15b | RNA binding motif protein 15B | 109095 | MGI:1923598 | -0.283 | 1 | n | y | n | protein coding gene |
| Crip3 | cysteine-rich protein 3 | 114570 | MGI:2152434 | -0.214 | 1 | n | y | n | protein coding gene |
| Bcas3 | BCAS3 microtubule associated cell migrat | 192197 | MGI:2385848 | -0.112 | 1 | n | y | n | protein coding gene |
| Nmrk1 | nicotinamide riboside kinase 1 | 225994 | MGI:2147434 | 0.035 | 1 | n | y | n | protein coding gene |
| Sestd1 | SEC14 and spectrin domains 1 | 228071 | MGI:1916262 | -0.182 | 1 | n | y | n | protein coding gene |
| Scn3b | sodium channel, voltage-gated, type III, be | 235281 | MGI:1918882 | -0.099 | 1 | n | y | n | protein coding gene |
| Brip1 | BRCA1 interacting protein C-terminal helic | 237911 | MGI:2442836 | -0.120 | 1 | n | y | n | protein coding gene |
| Lonrf1 | LON peptidase N-terminal domain and rin | 244421 | MGI:3609241 | -0.210 | 1 | n | y | n | protein coding gene |
| Spata2 | spermatogenesis associated 2 | 263876 | MGI:2146885 | -0.076 | 1 | n | y | n | protein coding gene |
| Hs3st6 | heparan sulfate (glucosamine) 3-O-sulfotr | 328779 | MGI:3580487 | -0.007 | 1 | n | y | n | protein coding gene |
| Abi2 | abl interactor 2 | 329165 | MGI:106913 | 0.110 | 1 | n | y | n | protein coding gene |
| Tafa3 | TAFA chemokine like family member 3 | 329731 | MGI:3046463 | -0.071 | 1 | n | y | n | protein coding gene |
| Cacfd1 | calcium channel flower domain containing | 381356 | MGI:1924317 | 0.075 | 1 | n | y | n | protein coding gene |
| AB124611 | cDNA sequence AB124611 | 382062 | MGI:3043001 | -0.239 | 1 | n | y | n | protein coding gene |
| Zyg11b | zyg-ll family member B, cell cycle regulatc | 414872 | MGI:2685277 | 0.001 | 1 | n | y | n | protein coding gene |
| Toporsl | topoisomerase I binding, arginine/serine-r | 68274 | MGI:1915524 | ND* | ND* | n | y | n | protein coding gene |
| Dydc2 | DPY30 domain containing 2 | 71200 | MGI:1918450 | ND* | ND* | n | y | n | protein coding gene |
| Usp50 | ubiquitin specific peptidase 50 | 75083 | MGI:1922333 | ND* | ND* | n | y | n | protein coding gene |
| Cabp7 | calcium binding protein 7 | 192650 | MGI:2183437 | ND* | ND* | n | y | n | protein coding gene |
| Kif28 | kinesin family member 28 | 383592 | MGI:2686151 | ND* | ND* | n | y | n | protein coding gene |
| Kif21b | kinesin family member 21B | 16565 | MGI:109234 | -2.352 | 6.06E-09 | n | n | y | protein coding gene |
| Styxl2 | serine/threonine/tyrosine interacting like 2 | 240892 | MGI:2685055 | -2.010 | 9.64E-10 | n | n | y | protein coding gene |
| Rprm | reprimo, TP53 dependent G2 arrest media | 67874 | MGI:1915124 | -1.853 | 1.50E-13 | n | n | y | protein coding gene |

| | | | | | | | | | |
|---|---|---|---|---|---|---|---|---|---|
| Inafm2 | InaF motif containing 2 | 100043272 | MGI:1915354 | -1.516 | 7.99E-05 | n | n | y | protein coding gene |
| Gnmt | glycine N-methyltransferase | 14711 | MGI:1202304 | -1.455 | 1.90E-06 | n | n | y | protein coding gene |
| Art5 | ADP-ribosyltransferase 5 | 11875 | MGI:107948 | -1.423 | 1.49E-05 | n | n | y | protein coding gene |
| Ppp1r14a | protein phosphatase 1, regulatory inhibitor | 68458 | MGI:1931139 | -1.397 | 7.27E-06 | n | n | y | protein coding gene |
| Casz1 | castor zinc finger 1 | 69743 | MGI:1196251 | -1.267 | 5.91E-05 | n | n | y | protein coding gene |
| Nf2 | neurofibromin 2 | 18016 | MGI:97307 | -1.125 | 3.98E-09 | n | n | y | protein coding gene |
| Chgb | chromogranin B | 12653 | MGI:88395 | -1.042 | 1.14E-07 | n | n | y | protein coding gene |
| Tlcd3b | TLC domain containing 3B | 68952 | MGI:1916202 | -1.033 | 5.40E-07 | n | n | y | protein coding gene |
| Sh2d4a | SH2 domain containing 4A | 72281 | MGI:1919531 | -1.028 | 4.67E-06 | n | n | y | protein coding gene |
| Pnma2 | paraneoplastic antigen MA2 | 239157 | MGI:2444129 | -0.979 | 7.33E-05 | n | n | y | protein coding gene |
| Pvalb | parvalbumin | 19293 | MGI:97821 | -0.921 | 6.64E-08 | n | n | y | protein coding gene |
| Fez1 | fasciculation and elongation protein zeta 1 | 235180 | MGI:2670976 | -0.855 | 1.36E-08 | n | n | y | protein coding gene |
| Fhod3 | formin homology 2 domain containing 3 | 225288 | MGI:1925847 | -0.827 | 2.24E-05 | n | n | y | protein coding gene |
| Snrnp25 | small nuclear ribonucleoprotein 25 (U11/U | 78372 | MGI:1925622 | -0.780 | 3.82E-06 | n | n | y | protein coding gene |
| Kcp | kielin/chordin-like protein | 333088 | MGI:2141640 | -0.735 | 5.15E-05 | n | n | y | protein coding gene |
| Olfm1 | olfactomedin 1 | 56177 | MGI:1860437 | -0.695 | 1.71E-05 | n | n | y | protein coding gene |
| St8sia3 | ST8 alpha-N-acetyl-neuraminide alpha-2,8 | 20451 | MGI:106019 | -0.681 | 2.54E-06 | n | n | y | protein coding gene |
| Cdk15 | cyclin dependent kinase 15 | 271697 | MGI:3583944 | -0.661 | 0.000673666 | n | n | y | protein coding gene |
| Endod1 | endonuclease domain containing 1 | 71946 | MGI:1919196 | -0.648 | 1.23E-05 | n | n | y | protein coding gene |
| Calb2 | calbindin 2 | 12308 | MGI:101914 | -0.558 | 3.87E-07 | n | n | y | protein coding gene |
| Ndufaf6 | NADH:ubiquinone oxidoreductase comple | 76947 | MGI:1924197 | -0.552 | 0.000275432 | n | n | y | protein coding gene |
| Trim45 | tripartite motif-containing 45 | 229644 | MGI:1918187 | -0.550 | 7.52E-05 | n | n | y | protein coding gene |
| Dpysl3 | dihydropyrimidinase-like 3 | 22240 | MGI:1349762 | -0.544 | 7.82E-05 | n | n | y | protein coding gene |
| Grxcr1 | glutaredoxin, cysteine rich 1 | 433899 | MGI:3577767 | -0.521 | 0.000213385 | n | n | y | protein coding gene |
| Sec61a2 | SEC61 translocon subunit alpha 2 | 57743 | MGI:1931071 | -0.451 | 0.000139658 | n | n | y | protein coding gene |
| Fam234b | family with sequence similarity 234, memb | 74525 | MGI:1921775 | -0.451 | 1.44E-05 | n | n | y | protein coding gene |
| Svbp | small vasohibin binding protein | 69216 | MGI:1916466 | -0.448 | 9.39E-05 | n | n | y | protein coding gene |
| Egfl7 | EGF-like domain 7 | 353156 | MGI:2449923 | -0.393 | 0.000933452 | n | n | y | protein coding gene |
| Nphp1 | nephronophthisis 1 (juvenile) homolog (hu | 53885 | MGI:1858233 | -0.392 | 0.000314158 | n | n | y | protein coding gene |
| Ctsf | cathepsin F | 56464 | MGI:1861434 | -0.359 | 0.000107051 | n | n | y | protein coding gene |
| Lhfpl5 | lipoma HMGIC fusion partner-like 5 | 328789 | MGI:1915382 | -0.343 | 0.000805895 | n | n | y | protein coding gene |
| Ptbp1 | polypyrimidine tract binding protein 1 | 19205 | MGI:97791 | -0.321 | 0.000306116 | n | n | y | protein coding gene |
| Camk2b | calcium/calmodulin-dependent protein kina | 12323 | MGI:88257 | 0.305 | 0.000227957 | n | n | y | protein coding gene |
| Yars1 | tyrosyl-tRNA synthetase 1 | 107271 | MGI:2147627 | 0.312 | 0.000171489 | n | n | y | protein coding gene |
| Hdac5 | histone deacetylase 5 | 15184 | MGI:1333784 | 0.321 | 0.000502308 | n | n | y | protein coding gene |
| Osbpl11 | oxysterol binding protein-like 11 | 106326 | MGI:2146553 | 0.322 | 0.000636483 | n | n | y | protein coding gene |
| Nrdc | nardilysin convertase | 230598 | MGI:1201386 | 0.332 | 0.000449066 | n | n | y | protein coding gene |
| Nprl2 | NPR2 like, GATOR1 complex subunit | 56032 | MGI:1914482 | 0.357 | 0.000637086 | n | n | y | protein coding gene |
| Tmem255b | transmembrane protein 255B | 272465 | MGI:2685533 | 0.363 | 0.00016505 | n | n | y | protein coding gene |
| Miga2 | mitoguardin 2 | 108958 | MGI:1922035 | 0.373 | 0.000475152 | n | n | y | protein coding gene |
| Ttc19 | tetratricopeptide repeat domain 19 | 72795 | MGI:1920045 | 0.374 | 0.000233407 | n | n | y | protein coding gene |
| Smpx | small muscle protein, X-linked | 66106 | MGI:1913356 | 0.380 | 0.000207342 | n | n | y | protein coding gene |
| Rab36 | RAB36, member RAS oncogene family | 76877 | MGI:1924127 | 0.387 | 5.95E-05 | n | n | y | protein coding gene |
| Usp46 | ubiquitin specific peptidase 46 | 69727 | MGI:1916977 | 0.387 | 0.000725083 | n | n | y | protein coding gene |
| Hgs | HGF-regulated tyrosine kinase substrate | 15239 | MGI:104681 | 0.395 | 0.000494769 | n | n | y | protein coding gene |
| Srd5a1 | steroid 5 alpha-reductase 1 | 78925 | MGI:98400 | 0.395 | 9.04E-05 | n | n | y | protein coding gene |
| Zfp612 | zinc finger protein 612 | 234725 | MGI:2443465 | 0.396 | 0.000177378 | n | n | y | protein coding gene |
| Golm1 | golgi membrane protein 1 | 105348 | MGI:1917329 | 0.399 | 0.000310131 | n | n | y | protein coding gene |
| Cds2 | CDP-diacylglycerol synthase 2 | 110911 | MGI:1332236 | 0.403 | 0.000228772 | n | n | y | protein coding gene |
| Lhx3 | LIM homeobox protein 3 | 16871 | MGI:102673 | 0.404 | 0.000557081 | n | n | y | protein coding gene |
| Sacm1l | SAC1 suppressor of actin mutations 1-like | 83493 | MGI:1933169 | 0.412 | 0.000155212 | n | n | y | protein coding gene |
| Uaca | uveal autoantigen with coiled-coil domains | 72565 | MGI:1919815 | 0.413 | 0.000716683 | n | n | y | protein coding gene |
| Ate1 | arginyltransferase 1 | 11907 | MGI:1333870 | 0.418 | 0.000447824 | n | n | y | protein coding gene |
| Kcnj13 | potassium inwardly-rectifying channel, sub | 100040591 | MGI:3781032 | 0.419 | 0.000261875 | n | n | y | protein coding gene |
| Map9 | microtubule-associated protein 9 | 213582 | MGI:2442208 | 0.428 | 0.000317066 | n | n | y | protein coding gene |
| Tomt | transmembrane O-methyltransferase | 791260 | MGI:3769724 | 0.435 | 0.000670323 | n | n | y | protein coding gene |
| Ptpn3 | protein tyrosine phosphatase, non-recepto | 545622 | MGI:105307 | 0.453 | 0.00017024 | n | n | y | protein coding gene |
| Usp20 | ubiquitin specific peptidase 20 | 74270 | MGI:1921520 | 0.455 | 0.000209844 | n | n | y | protein coding gene |
| Insc | INSC spindle orientation adaptor protein | 233752 | MGI:1917942 | 0.472 | 0.000161382 | n | n | y | protein coding gene |
| Cfap69 | cilia and flagella associated protein 69 | 207686 | MGI:2443778 | 0.473 | 0.000637521 | n | n | y | protein coding gene |
| Clic5 | chloride intracellular channel 5 | 224796 | MGI:1917912 | 0.473 | 0.000419881 | n | n | y | protein coding gene |
| Tmcc2 | transmembrane and coiled-coil domains 2 | 68875 | MGI:1916125 | 0.475 | 3.89E-05 | n | n | y | protein coding gene |
| Ckmt1 | creatine kinase, mitochondrial 1, ubiquitou | 12716 | MGI:99441 | 0.483 | 0.000102818 | n | n | y | protein coding gene |
| Atg4d | autophagy related 4D, cysteine peptidase | 235040 | MGI:2444308 | 0.489 | 0.000135568 | n | n | y | protein coding gene |

| | | | | | | | | | |
|---|---|---|---|---|---|---|---|---|---|
| Mlf1 | myeloid leukemia factor 1 | 17349 | MGI:1341819 | 0.493 | 0.0001922 | n | n | y | protein coding gene |
| Ank3 | ankyrin 3, epithelial | 11735 | MGI:88026 | 0.502 | 0.000394773 | n | n | y | protein coding gene |
| Pgm2l1 | phosphoglucomutase 2-like 1 | 70974 | MGI:1918224 | 0.506 | 6.02E-05 | n | n | y | protein coding gene |
| Tjap1 | tight junction associated protein 1 | 74094 | MGI:1921344 | 0.507 | 8.24E-06 | n | n | y | protein coding gene |
| Cdkl2 | cyclin dependent kinase like 2 | 53886 | MGI:1858227 | 0.513 | 6.34E-05 | n | n | y | protein coding gene |
| Fhit | fragile histidine triad gene | 14198 | MGI:1277947 | 0.522 | 5.77E-05 | n | n | y | protein coding gene |
| Hspa4l | heat shock protein 4 like | 18415 | MGI:107422 | 0.524 | 3.05E-05 | n | n | y | protein coding gene |
| Tmem107 | transmembrane protein 107 | 66910 | MGI:1914160 | 0.530 | 0.000498 | n | n | y | protein coding gene |
| Slc39a3 | solute carrier family 39 (zinc transporter), | 106947 | MGI:2147269 | 0.532 | 3.96E-05 | n | n | y | protein coding gene |
| Gpr155 | G protein-coupled receptor 155 | 68526 | MGI:1915776 | 0.542 | 0.000703177 | n | n | y | protein coding gene |
| Dusp14 | dual specificity phosphatase 14 | 56405 | MGI:1927168 | 0.551 | 3.37E-05 | n | n | y | protein coding gene |
| Twf2 | twinfilin actin binding protein 2 | 23999 | MGI:1346078 | 0.556 | 5.34E-06 | n | n | y | protein coding gene |
| Rdh12 | retinol dehydrogenase 12 | 77974 | MGI:1925224 | 0.580 | 6.28E-06 | n | n | y | protein coding gene |
| Myo18a | myosin XVIIIA | 360013 | MGI:2667185 | 0.585 | 6.21E-06 | n | n | y | protein coding gene |
| Mfng | MFNG O-fucosylpeptide 3-beta-N-acetylg | 17305 | MGI:1095404 | 0.592 | 5.99E-07 | n | n | y | protein coding gene |
| Spag6l | sperm associated antigen 6-like | 50525 | MGI:1354388 | 0.633 | 0.000228071 | n | n | y | protein coding gene |
| Atoh1 | atonal bHLH transcription factor 1 | 11921 | MGI:104654 | 0.638 | 0.000889297 | n | n | y | protein coding gene |
| Ppp2r5b | protein phosphatase 2, regulatory subunit | 225849 | MGI:2388480 | 0.638 | 8.70E-05 | n | n | y | protein coding gene |
| Ap3m2 | adaptor-related protein complex 3, mu 2 s | 64933 | MGI:1929214 | 0.659 | 1.72E-07 | n | n | y | protein coding gene |
| Evc2 | EvC ciliary complex subunit 2 | 68525 | MGI:1915775 | 0.665 | 6.46E-06 | n | n | y | protein coding gene |
| Adgrv1 | adhesion G protein-coupled receptor V1 | 110789 | MGI:1274784 | 0.681 | 0.000110178 | n | n | y | protein coding gene |
| Syt13 | synaptotagmin XIII | 80976 | MGI:1933945 | 0.686 | 1.95E-05 | n | n | y | protein coding gene |
| Pierce2 | piercer of microtubule wall 2 | 546143 | MGI:3648770 | 0.693 | 0.0001922 | n | n | y | protein coding gene |
| Kncn | kinocilin | 654462 | MGI:3614952 | 0.696 | 1.14E-05 | n | n | y | protein coding gene |
| Uhmk1 | U2AF homology motif (UHM) kinase 1 | 16589 | MGI:1341908 | 0.696 | 0.000607188 | n | n | y | protein coding gene |
| Ak1 | adenylate kinase 1 | 11636 | MGI:87977 | 0.697 | 9.86E-06 | n | n | y | protein coding gene |
| Gfi1 | growth factor independent 1 transcription | 14581 | MGI:103170 | 0.710 | 3.78E-07 | n | n | y | protein coding gene |
| B3gnt4 | UDP-GlcNAc:betaGal beta-1,3-N-acetylgl | 231727 | MGI:2680208 | 0.713 | 0.000116742 | n | n | y | protein coding gene |
| Cdkl4 | cyclin dependent kinase like 4 | 381113 | MGI:3587025 | 0.715 | 4.56E-07 | n | n | y | protein coding gene |
| Nipal3 | NIPA-like domain containing 3 | 74552 | MGI:1921802 | 0.737 | 2.83E-05 | n | n | y | protein coding gene |
| Dnaja4 | DnaJ heat shock protein family (Hsp40) m | 58233 | MGI:1927638 | 0.740 | 1.48E-06 | n | n | y | protein coding gene |
| Spock2 | sparc/osteonectin, cwcv and kazal-like do | 94214 | MGI:1891351 | 0.746 | 5.53E-06 | n | n | y | protein coding gene |
| Calb1 | calbindin 1 | 12307 | MGI:88248 | 0.751 | 2.96E-07 | n | n | y | protein coding gene |
| Plch2 | phospholipase C, eta 2 | 269615 | MGI:2443078 | 0.762 | 2.92E-05 | n | n | y | protein coding gene |
| Smap2 | small ArfGAP 2 | 69780 | MGI:1917030 | 0.775 | 4.23E-08 | n | n | y | protein coding gene |
| Ptprq | protein tyrosine phosphatase receptor typ | 237523 | MGI:1096349 | 0.792 | 0.000803483 | n | n | y | protein coding gene |
| Mogat1 | monoacylglycerol O-acyltransferase 1 | 68393 | MGI:1915643 | 0.838 | 2.71E-05 | n | n | y | protein coding gene |
| Myo15a | myosin XVA | 17910 | MGI:1261811 | 0.840 | 4.40E-06 | n | n | y | protein coding gene |
| Foxj1 | forkhead box J1 | 15223 | MGI:1347474 | 0.864 | 0.000236499 | n | n | y | protein coding gene |
| Pcsk9 | proprotein convertase subtilisin/kexin type | 100102 | MGI:2140260 | 0.870 | 1.54E-05 | n | n | y | protein coding gene |
| Drc1 | dynein regulatory complex subunit 1 | 381738 | MGI:2685906 | 0.871 | 0.000631194 | n | n | y | protein coding gene |
| Hsph1 | heat shock 105kDa/110kDa protein 1 | 15505 | MGI:105053 | 0.872 | 3.59E-07 | n | n | y | protein coding gene |
| Rlig1 | RNA 5'-phosphate and 3'-OH ligase 1 | 68281 | MGI:1921197 | 0.898 | 1.58E-05 | n | n | y | protein coding gene |
| Rsph1 | radial spoke head 1 homolog (Chlamydom | 22092 | MGI:1194909 | 0.956 | 5.80E-07 | n | n | y | protein coding gene |
| Myo3a | myosin IIIA | 667663 | MGI:2183924 | 0.987 | 1.17E-06 | n | n | y | protein coding gene |
| Atf7ip | activating transcription factor 7 interacting | 54343 | MGI:1858965 | 0.991 | 0.000113531 | n | n | y | protein coding gene |
| Cdkn2d | cyclin dependent kinase inhibitor 2D | 12581 | MGI:105387 | 1.018 | 1.93E-09 | n | n | y | protein coding gene |
| Capsl | calcyphosine-like | 75568 | MGI:1922818 | 1.045 | 4.34E-06 | n | n | y | protein coding gene |
| Smim5 | small integral membrane protein 5 | 66528 | MGI:1913778 | 1.064 | 6.26E-07 | n | n | y | protein coding gene |
| Syt14 | synaptotagmin XIV | 329324 | MGI:2444490 | 1.088 | 0.000114034 | n | n | y | protein coding gene |
| 1700088E04Rik | RIKEN cDNA 1700088E04 gene | 27660 | MGI:1920774 | 1.102 | 4.07E-05 | n | n | y | protein coding gene |
| Kcnh7 | potassium voltage-gated channel, subfam | 170738 | MGI:2159566 | 1.139 | 1.98E-05 | n | n | y | protein coding gene |
| Hrob | homologous recombination factor with OB | 217216 | MGI:2387601 | 1.158 | 1.26E-07 | n | n | y | protein coding gene |
| Tmem191 | transmembrane protein 191 | 224019 | MGI:107238 | 1.221 | 3.30E-09 | n | n | y | protein coding gene |
| Sting1 | stimulator of interferon response cGAMP i | 72512 | MGI:1919762 | 1.221 | 4.05E-06 | n | n | y | protein coding gene |
| Gabrb3 | GABRB3, gamma-aminobutyric acid type | 14402 | MGI:95621 | 1.243 | 3.06E-07 | n | n | y | protein coding gene |
| Acss2 | acyl-CoA synthetase short-chain family m | 60525 | MGI:1890410 | 1.288 | 1.05E-09 | n | n | y | protein coding gene |
| Dynlt5 | dynein light chain Tctex-type 5 | 67344 | MGI:1914594 | 1.334 | 0.000439307 | n | n | y | protein coding gene |
| Cdkl1 | cyclin dependent kinase like 1 | 71091 | MGI:1918341 | 1.384 | 2.94E-05 | n | n | y | protein coding gene |
| Car7 | carbonic anhydrase 7 | 12354 | MGI:103100 | 1.484 | 1.29E-08 | n | n | y | protein coding gene |
| Cfap206 | cilia and flagella associated protein 206 | 69329 | MGI:1916579 | 1.517 | 7.22E-09 | n | n | y | protein coding gene |
| R3hdml | R3H domain containing-like | 100043899 | MGI:3650937 | 1.535 | 6.39E-08 | n | n | y | protein coding gene |
| Galnt9 | polypeptide N-acetylgalactosaminyltransfe | 231605 | MGI:2677965 | 1.677 | 4.20E-06 | n | n | y | protein coding gene |
| Cfap52 | cilia and flagella associated protein 52 | 71860 | MGI:1919110 | 1.811 | 5.56E-09 | n | n | y | protein coding gene |

| | | | | | | | | | |
|---|---|---|---|---|---|---|---|---|---|
| Il25 | interleukin 25 | 140806 | MGI:2155888 | 2.111 | 9.86E-06 | n | n | y | protein coding gene |
| Khdrbs2 | KH domain containing, RNA binding, signa | 170771 | MGI:2159649 | 2.200 | 6.50E-09 | n | n | y | protein coding gene |
| Cpne9 | copine family member IX | 211232 | MGI:2443052 | 2.534 | 1.53E-10 | n | n | y | protein coding gene |
| D7Ertd443e | DNA segment, Chr 7, ERATO Doi 443, ex | 71007 | MGI:1196431 | 3.548 | 2.36E-07 | n | n | y | protein coding gene |
| Ube2ql1 | ubiquitin-conjugating enzyme E2Q family- | 76980 | MGI:1924230 | -0.634 | 0.001001086 | n | n | y | protein coding gene |
| Dalrd3 | DALR anticodon binding domain containin | 67789 | MGI:1915039 | 0.329 | 0.001001752 | n | n | y | protein coding gene |
| Arhgef28 | Rho guanine nucleotide exchange factor 2 | 110596 | MGI:1346016 | 0.468 | 0.001090417 | n | n | y | protein coding gene |
| Crlf1 | cytokine receptor-like factor 1 | 12931 | MGI:1340030 | -0.859 | 0.001133057 | n | n | y | protein coding gene |
| Unc45a | unc-45 myosin chaperone A | 101869 | MGI:2142246 | 0.269 | 0.001168619 | n | n | y | protein coding gene |
| Ldb3 | LIM domain binding 3 | 24131 | MGI:1344412 | 0.702 | 0.001249297 | n | n | y | protein coding gene |
| Slc14a1 | solute carrier family 14 (urea transporter), | 108052 | MGI:1351654 | -0.750 | 0.001249297 | n | n | y | protein coding gene |
| Cenpb | centromere protein B | 12616 | MGI:88376 | -0.541 | 0.001280559 | n | n | y | protein coding gene |
| Parp6 | poly (ADP-ribose) polymerase family, mer | 67287 | MGI:1914537 | 0.273 | 0.001289683 | n | n | y | protein coding gene |
| Kcnb1 | potassium voltage gated channel, Shab-re | 16500 | MGI:96666 | 0.559 | 0.001352826 | n | n | y | protein coding gene |
| Dynlrb2 | dynein light chain roadblock-type 2 | 75465 | MGI:1922715 | 0.613 | 0.001359643 | n | n | y | protein coding gene |
| Galt | galactose-1-phosphate uridyl transferase | 14430 | MGI:95638 | -0.291 | 0.001509125 | n | n | y | protein coding gene |
| Cfap119 | cilia and flagella associated protein 119 | 233899 | MGI:2685012 | 0.371 | 0.001537281 | n | n | y | protein coding gene |
| Dtna | dystrobrevin alpha | 13527 | MGI:106039 | 0.392 | 0.0015601 | n | n | y | protein coding gene |
| Lhfpl4 | lipoma HMGIC fusion partner-like protein 4 | 269788 | MGI:3057108 | 0.383 | 0.001619246 | n | n | y | protein coding gene |
| Dapk3 | death-associated protein kinase 3 | 13144 | MGI:1203520 | 0.467 | 0.001644791 | n | n | y | protein coding gene |
| Tmie | transmembrane inner ear | 20776 | MGI:2159400 | 0.409 | 0.001702759 | n | n | y | protein coding gene |
| Clrn1 | clarin 1 | 229320 | MGI:2388124 | 0.533 | 0.001737091 | n | n | y | protein coding gene |
| Rhpn1 | rhophilin, Rho GTPase binding protein 1 | 14787 | MGI:1098783 | -0.298 | 0.0017764 | n | n | y | protein coding gene |
| Zfp949 | zinc finger protein 949 | 71640 | MGI:1918890 | 0.615 | 0.001805749 | n | n | y | protein coding gene |
| Cdc42bpa | CDC42 binding protein kinase alpha | 226751 | MGI:2441841 | -1.491 | 0.001888697 | n | n | y | protein coding gene |
| Srebf2 | sterol regulatory element binding factor 2 | 20788 | MGI:107585 | 0.389 | 0.001936665 | n | n | y | protein coding gene |
| Scg3 | secretogranin III | 20255 | MGI:103032 | 0.373 | 0.001949302 | n | n | y | protein coding gene |
| Chac2 | ChaC, cation transport regulator 2 | 68044 | MGI:1915294 | 0.859 | 0.002105842 | n | n | y | protein coding gene |
| Nt5dc3 | 5'-nucleotidase domain containing 3 | 103466 | MGI:3513266 | 0.395 | 0.00220312 | n | n | y | protein coding gene |
| Msra | methionine sulfoxide reductase A | 110265 | MGI:106916 | -0.565 | 0.002208819 | n | n | y | protein coding gene |
| Reps1 | RalBP1 associated Eps domain containing | 19707 | MGI:1196373 | -0.380 | 0.00231608 | n | n | y | protein coding gene |
| Pacsin1 | protein kinase C and casein kinase substr | 23969 | MGI:1345181 | 0.362 | 0.002338702 | n | n | y | protein coding gene |
| Efr3a | EFR3 homolog A | 76740 | MGI:1923990 | 0.315 | 0.002602964 | n | n | y | protein coding gene |
| Rasd2 | RASD family, member 2 | 75141 | MGI:1922391 | -0.258 | 0.002661938 | n | n | y | protein coding gene |
| Cep19 | centrosomal protein 19 | 66994 | MGI:1914244 | 0.273 | 0.002781033 | n | n | y | protein coding gene |
| Vps45 | vacuolar protein sorting 45 | 22365 | MGI:891965 | 0.426 | 0.002782287 | n | n | y | protein coding gene |
| Aftph | aftiphilin | 216549 | MGI:1923012 | 0.262 | 0.002868764 | n | n | y | protein coding gene |
| Dctn1 | dynactin 1 | 13191 | MGI:107745 | 0.456 | 0.002922716 | n | n | y | protein coding gene |
| Clip4 | CAP-GLY domain containing linker protein | 78785 | MGI:1919100 | 0.378 | 0.003057423 | n | n | y | protein coding gene |
| Pkig | protein kinase inhibitor, gamma | 18769 | MGI:1343086 | -0.245 | 0.003071236 | n | n | y | protein coding gene |
| Lmod3 | leiomodin 3 (fetal) | 320502 | MGI:2444169 | -0.299 | 0.003351021 | n | n | y | protein coding gene |
| Whrn | whirlin | 73750 | MGI:2682003 | -0.361 | 0.003488215 | n | n | y | protein coding gene |
| Nectin1 | nectin cell adhesion molecule 1 | 58235 | MGI:1926483 | 0.415 | 0.003501536 | n | n | y | protein coding gene |
| Nhlh1 | nescient helix loop helix 1 | 18071 | MGI:98481 | 1.162 | 0.003729208 | n | n | y | protein coding gene |
| Rundc3a | RUN domain containing 3A | 51799 | MGI:1858752 | 0.285 | 0.003796384 | n | n | y | protein coding gene |
| Nefm | neurofilament, medium polypeptide | 18040 | MGI:97314 | 0.636 | 0.003806179 | n | n | y | protein coding gene |
| Rtn2 | reticulon 2 (Z-band associated protein) | 20167 | MGI:107612 | -0.361 | 0.003933109 | n | n | y | protein coding gene |
| Usp19 | ubiquitin specific peptidase 19 | 71472 | MGI:1918722 | 0.288 | 0.004119503 | n | n | y | protein coding gene |
| Gipc3 | GIPC PDZ domain containing family, mem | 209047 | MGI:2387006 | 0.628 | 0.004216138 | n | n | y | protein coding gene |
| Blzf1 | basic leucine zipper nuclear factor 1 | 66352 | MGI:1201607 | -0.252 | 0.004280601 | n | n | y | protein coding gene |
| Scaf8 | SR-related CTD-associated factor 8 | 106583 | MGI:1925212 | -0.332 | 0.004299117 | n | n | y | protein coding gene |
| Sympk | symplekin | 68188 | MGI:1915438 | 0.356 | 0.004330505 | n | n | y | protein coding gene |
| Ttc21a | tetratricopeptide repeat domain 21A | 74052 | MGI:1921302 | 0.607 | 0.004448554 | n | n | y | protein coding gene |
| Gas8 | growth arrest specific 8 | 104346 | MGI:1202386 | 0.303 | 0.004550714 | n | n | y | protein coding gene |
| Fktn | fukutin | 246179 | MGI:2179507 | 0.307 | 0.004670656 | n | n | y | protein coding gene |
| Dock9 | dedicator of cytokinesis 9 | 105445 | MGI:106321 | -0.360 | 0.004675587 | n | n | y | protein coding gene |
| Rab3a | RAB3A, member RAS oncogene family | 19339 | MGI:97843 | 0.249 | 0.004733786 | n | n | y | protein coding gene |
| Psmc3ip | proteasome (prosome, macropain) 26S su | 19183 | MGI:1098610 | 1.079 | 0.004774222 | n | n | y | protein coding gene |
| Slc37a4 | solute carrier family 37 (glucose-6-phosph | 14385 | MGI:1316650 | 0.304 | 0.004775903 | n | n | y | protein coding gene |
| Upf2 | UPF2 regulator of nonsense transcripts ho | 326622 | MGI:2449307 | 0.426 | 0.005058891 | n | n | y | protein coding gene |
| Slc52a3 | solute carrier protein family 52, member 3 | 69698 | MGI:1916948 | 0.610 | 0.005115438 | n | n | y | protein coding gene |
| Eif4enif1 | eukaryotic translation initiation factor 4E n | 74203 | MGI:1921453 | -0.302 | 0.005131508 | n | n | y | protein coding gene |
| Pigyl | phosphatidylinositol glycan anchor biosyn | 66268 | MGI:1913518 | -0.308 | 0.005162979 | n | n | y | protein coding gene |
| Agtpbp1 | ATP/GTP binding protein 1 | 67269 | MGI:2159437 | 0.438 | 0.00521368 | n | n | y | protein coding gene |

| | | | | | | | | | |
|---|---|---|---|---|---|---|---|---|---|
| Usp7 | ubiquitin specific peptidase 7 | 252870 | MGI:2182061 | -0.366 | 0.005356929 | n | n | y | protein coding gene |
| Dnai2 | dynein axonemal intermediate chain 2 | 432611 | MGI:2685574 | 0.384 | 0.005403524 | n | n | y | protein coding gene |
| Polr3gl | polymerase (RNA) III (DNA directed) poly | 69870 | MGI:1917120 | -0.282 | 0.005451431 | n | n | y | protein coding gene |
| Rab11fip1 | RAB11 family interacting protein 1 (class I | 75767 | MGI:1923017 | 0.330 | 0.005507449 | n | n | y | protein coding gene |
| Rcbtb2 | regulator of chromosome condensation (R | 105670 | MGI:1917200 | 0.255 | 0.005567566 | n | n | y | protein coding gene |
| Gpr156 | G protein-coupled receptor 156 | 239845 | MGI:2653880 | 0.409 | 0.005724316 | n | n | y | protein coding gene |
| Rgs12 | regulator of G-protein signaling 12 | 71729 | MGI:1918979 | 0.270 | 0.005731406 | n | n | y | protein coding gene |
| Ccdc65 | coiled-coil domain containing 65 | 105833 | MGI:2146001 | 0.492 | 0.005942632 | n | n | y | protein coding gene |
| Phtf2 | putative homeodomain transcription factor | 68770 | MGI:1916020 | 0.343 | 0.005943115 | n | n | y | protein coding gene |
| Mindy1 | MINDY lysine 48 deubiquitinase 1 | 75007 | MGI:1922257 | 0.240 | 0.006225796 | n | n | y | protein coding gene |
| Hsd17b14 | hydroxysteroid (17-beta) dehydrogenase | 66065 | MGI:1913315 | 0.335 | 0.007074599 | n | n | y | protein coding gene |
| Wdr33 | WD repeat domain 33 | 74320 | MGI:1921570 | 0.233 | 0.00714272 | n | n | y | protein coding gene |
| Bpnt1 | 3'(2'), 5'-bisphosphate nucleotidase 1 | 23827 | MGI:1338800 | 0.286 | 0.007831832 | n | n | y | protein coding gene |
| Chka | choline kinase alpha | 12660 | MGI:107760 | -0.374 | 0.007959702 | n | n | y | protein coding gene |
| Tmem184a | transmembrane protein 184a | 231832 | MGI:2385897 | 0.266 | 0.008059332 | n | n | y | protein coding gene |
| Lnx2 | ligand of numb-protein X 2 | 140887 | MGI:2155959 | 0.301 | 0.008289125 | n | n | y | protein coding gene |
| E130308A19Rik | RIKEN cDNA E130308A19 gene | 230259 | MGI:2442164 | -0.277 | 0.009106328 | n | n | y | protein coding gene |
| Ngrn | neugrin, neurite outgrowth associated | 83485 | MGI:1933212 | 0.260 | 0.009594833 | n | n | y | protein coding gene |
| Mkrn2os | makorin, ring finger protein 2, opposite str | 70291 | MGI:1917541 | 0.339 | 0.010094894 | n | n | y | protein coding gene |
| Mtf1 | metal response element binding transcript | 17764 | MGI:101786 | 0.265 | 0.011001311 | n | n | y | protein coding gene |
| Cdk5rap2 | CDK5 regulatory subunit associated prote | 214444 | MGI:2384875 | 0.210 | 0.011018251 | n | n | y | protein coding gene |
| Tmem183a | transmembrane protein 183A | 57439 | MGI:1914729 | 0.198 | 0.01106509 | n | n | y | protein coding gene |
| Ipo13 | importin 13 | 230673 | MGI:2385205 | 0.238 | 0.0110912 | n | n | y | protein coding gene |
| Ica1 | islet cell autoantigen 1 | 15893 | MGI:96391 | -0.254 | 0.011492466 | n | n | y | protein coding gene |
| Kcnmb2 | potassium large conductance calcium-acti | 72413 | MGI:1919663 | 0.327 | 0.011531649 | n | n | y | protein coding gene |
| Erich3 | glutamate rich 3 | 209601 | MGI:1919095 | 0.494 | 0.011763066 | n | n | y | protein coding gene |
| Cfap126 | cilia and flagella associated protein 126 | 75472 | MGI:1922722 | 0.576 | 0.011942321 | n | n | y | protein coding gene |
| Serpinb6a | serine (or cysteine) peptidase inhibitor, cla | 20719 | MGI:103123 | 0.265 | 0.01225794 | n | n | y | protein coding gene |
| Sik3 | SIK family kinase 3 | 70661 | MGI:2446296 | -0.406 | 0.012291805 | n | n | y | protein coding gene |
| Zfp865 | zinc finger protein 865 | 319748 | MGI:2442656 | -0.445 | 0.012503856 | n | n | y | protein coding gene |
| Nup153 | nucleoporin 153 | 218210 | MGI:2385621 | 0.323 | 0.012853278 | n | n | y | protein coding gene |
| Mthfd2 | methylenetetrahydrofolate dehydrogenase | 17768 | MGI:1338850 | 0.278 | 0.013259432 | n | n | y | protein coding gene |
| Rabl2 | RAB, member RAS oncogene family-like 2 | 68708 | MGI:1915958 | -0.461 | 0.013962551 | n | n | y | protein coding gene |
| Atp5mk | ATP synthase membrane subunit k | 66477 | MGI:1891435 | -0.759 | 0.015086358 | n | n | y | protein coding gene |
| Dhx38 | DEAH-box helicase 38 | 64340 | MGI:1927617 | -0.201 | 0.01709175 | n | n | y | protein coding gene |
| Pard6a | par-6 family cell polarity regulator alpha | 56513 | MGI:1927223 | 0.357 | 0.017218741 | n | n | y | protein coding gene |
| Rogdi | rogdi homolog | 66049 | MGI:1913299 | -0.208 | 0.017218741 | n | n | y | protein coding gene |
| Noa1 | nitric oxide associated 1 | 56412 | MGI:1914306 | 0.416 | 0.017717142 | n | n | y | protein coding gene |
| Rph3al | rabphilin 3A-like (without C2 domains) | 380714 | MGI:1923492 | 0.256 | 0.017950116 | n | n | y | protein coding gene |
| Ppfia3 | protein tyrosine phosphatase, receptor typ | 76787 | MGI:1924037 | 0.378 | 0.01929301 | n | n | y | protein coding gene |
| Atmin | ATM interactor | 234776 | MGI:2682328 | -0.249 | 0.020331953 | n | n | y | protein coding gene |
| Bola2 | bolA family member 2 | 66162 | MGI:1913412 | -0.511 | 0.02048481 | n | n | y | protein coding gene |
| Tmco3 | transmembrane and coiled-coil domains 3 | 234076 | MGI:2444946 | 0.290 | 0.020641549 | n | n | y | protein coding gene |
| Acd | adrenocortical dysplasia | 497652 | MGI:87873 | -0.239 | 0.021809333 | n | n | y | protein coding gene |
| Zdhhc16 | zinc finger, DHHC domain containing 16 | 74168 | MGI:1921418 | -0.177 | 0.021870968 | n | n | y | protein coding gene |
| Gm11837 | predicted gene 11837 | 100038514 | MGI:3702175 | -0.810 | 0.022050203 | n | n | y | protein coding gene |
| Morn1 | MORN repeat containing 1 | 76866 | MGI:1924116 | -0.255 | 0.023017834 | n | n | y | protein coding gene |
| Mthfr | methylenetetrahydrofolate reductase | 17769 | MGI:106639 | 0.333 | 0.02421524 | n | n | y | protein coding gene |
| Slc25a19 | solute carrier family 25 (mitochondrial thia | 67283 | MGI:1914533 | 0.289 | 0.02485877 | n | n | y | protein coding gene |
| Extl2 | exostosin-like glycosyltransferase 2 | 58193 | MGI:1889574 | -0.191 | 0.025207117 | n | n | y | protein coding gene |
| Homer1 | homer scaffolding protein 1 | 26556 | MGI:1347345 | 0.392 | 0.025895949 | n | n | y | protein coding gene |
| Zbtb18 | zinc finger and BTB domain containing 18 | 30928 | MGI:1353609 | -0.765 | 0.025976055 | n | n | y | protein coding gene |
| Drc3 | dynein regulatory complex subunit 3 | 74665 | MGI:1921915 | 0.325 | 0.026061316 | n | n | y | protein coding gene |
| Jup | junction plakoglobin | 16480 | MGI:96650 | 0.208 | 0.026624088 | n | n | y | protein coding gene |
| Peak1 | pseudopodium-enriched atypical kinase 1 | 244895 | MGI:2442366 | 0.538 | 0.028172829 | n | n | y | protein coding gene |
| Cplx1 | complexin 1 | 12889 | MGI:104727 | -0.650 | 0.028754908 | n | n | y | protein coding gene |
| Prkx | protein kinase, X-linked | 19108 | MGI:1309999 | 0.247 | 0.029122634 | n | n | y | protein coding gene |
| Cimap3 | ciliary microtubule associated protein 3 | 100503311 | MGI:1923670 | -0.437 | 0.029349647 | n | n | y | protein coding gene |
| Six4 | sine oculis-related homeobox 4 | 20474 | MGI:106034 | 0.236 | 0.030389584 | n | n | y | protein coding gene |
| Cyld | CYLD lysine 63 deubiquitinase | 74256 | MGI:1921506 | 0.203 | 0.030667497 | n | n | y | protein coding gene |
| Slk | STE20-like kinase | 20874 | MGI:103241 | -0.404 | 0.032626741 | n | n | y | protein coding gene |
| Abr | active BCR-related gene | 109934 | MGI:107771 | 0.285 | 0.032872883 | n | n | y | protein coding gene |
| Cand1 | cullin associated and neddylation disasso | 71902 | MGI:1261820 | 0.193 | 0.033769442 | n | n | y | protein coding gene |
| Nbeal1 | neurobeachin like 1 | 269198 | MGI:2444343 | 0.591 | 0.034153166 | n | n | y | protein coding gene |

| | | | | | | | | | |
|---|---|---|---|---|---|---|---|---|---|
| Iqcg | IQ motif containing G | 69707 | MGI:1916957 | 0.380 | 0.03457872 | n | n | y | protein coding gene |
| Stmn3 | stathmin-like 3 | 20262 | MGI:1277137 | -0.621 | 0.034879295 | n | n | y | protein coding gene |
| Scn1b | sodium channel, voltage-gated, type I, bet | 20266 | MGI:98247 | 0.636 | 0.035503712 | n | n | y | protein coding gene |
| Cmip | c-Maf inducing protein | 74440 | MGI:1921690 | -0.381 | 0.035553649 | n | n | y | protein coding gene |
| Nceh1 | neutral cholesterol ester hydrolase 1 | 320024 | MGI:2443191 | 0.199 | 0.035741963 | n | n | y | protein coding gene |
| Tprkb | Tp53rk binding protein | 69786 | MGI:1917036 | -0.192 | 0.036196541 | n | n | y | protein coding gene |
| Mon2 | MON2 homolog, regulator of endosome tc | 67074 | MGI:1914324 | 0.684 | 0.038435538 | n | n | y | protein coding gene |
| Fcor | Foxo1 corepressor | 100503924 | MGI:1915484 | 0.847 | 0.04061253 | n | n | y | protein coding gene |
| Wdr11 | WD repeat domain 11 | 207425 | MGI:1920230 | 0.273 | 0.040668817 | n | n | y | protein coding gene |
| Rnf157 | ring finger protein 157 | 217340 | MGI:2442484 | -0.462 | 0.041357799 | n | n | y | protein coding gene |
| Ift20 | intraflagellar transport 20 | 55978 | MGI:1915585 | -0.199 | 0.041738325 | n | n | y | protein coding gene |
| Gtpbp1 | GTP binding protein 1 | 14904 | MGI:109443 | -0.244 | 0.043823949 | n | n | y | protein coding gene |
| Pcdh15 | protocadherin 15 | 11994 | MGI:1891428 | 0.384 | 0.045198998 | n | n | y | protein coding gene |
| Scg5 | secretogranin V | 20394 | MGI:98289 | -0.208 | 0.046027242 | n | n | y | protein coding gene |
| Myo6 | myosin VI | 17920 | MGI:104785 | 0.253 | 0.046031843 | n | n | y | protein coding gene |
| Parp1 | poly (ADP-ribose) polymerase family, mer | 11545 | MGI:1340806 | 0.181 | 0.048452001 | n | n | y | protein coding gene |
| Lrtm2 | leucine-rich repeats and transmembrane c | 211187 | MGI:2141485 | 0.285 | 0.048912683 | n | n | y | protein coding gene |
| Wdsub1 | WD repeat, SAM and U-box domain conta | 72137 | MGI:1919387 | 0.222 | 0.049781881 | n | n | y | protein coding gene |
| C2cd5 | C2 calcium-dependent domain containing | 74741 | MGI:1921991 | 0.371 | 0.049851504 | n | n | y | protein coding gene |
| Abca5 | ATP-binding cassette, sub-family A memb | 217265 | MGI:2386607 | 0.380 | 0.050318629 | n | n | y | protein coding gene |
| Elmod1 | ELMO/CED-12 domain containing 1 | 270162 | MGI:3583900 | -0.527 | 0.050371324 | n | n | y | protein coding gene |
| Rbfox2 | RNA binding protein, fox-1 homolog (C. el | 93686 | MGI:1933973 | -0.238 | 0.050506741 | n | n | y | protein coding gene |
| Ica1l | islet cell autoantigen 1-like | 70375 | MGI:1917625 | 0.235 | 0.051156257 | n | n | y | protein coding gene |
| Ppp2r1b | protein phosphatase 2, regulatory subunit | 73699 | MGI:1920949 | -0.172 | 0.051955551 | n | n | y | protein coding gene |
| Entrep3 | endosomal transmembrane epsin interact | 68521 | MGI:1915771 | 0.251 | 0.053093965 | n | n | y | protein coding gene |
| Megf8 | multiple EGF-like-domains 8 | 269878 | MGI:2446294 | 0.221 | 0.053293005 | n | n | y | protein coding gene |
| Pdhx | pyruvate dehydrogenase complex, compo | 27402 | MGI:1351627 | -0.200 | 0.05466547 | n | n | y | protein coding gene |
| Ift27 | intraflagellar transport 27 | 67042 | MGI:1914292 | -0.184 | 0.056096326 | n | n | y | protein coding gene |
| Tet3 | tet methylcytosine dioxygenase 3 | 194388 | MGI:2446229 | -0.208 | 0.056264053 | n | n | y | protein coding gene |
| Fcho1 | FCH domain only 1 | 74015 | MGI:1921265 | 0.255 | 0.057285191 | n | n | y | protein coding gene |
| Fam98c | family with sequence similarity 98, membe | 73833 | MGI:1921083 | -0.272 | 0.057762352 | n | n | y | protein coding gene |
| Syt7 | synaptotagmin VII | 54525 | MGI:1859545 | -0.488 | 0.05942571 | n | n | y | protein coding gene |
| Ankrd42 | ankyrin repeat domain 42 | 73845 | MGI:1921095 | 0.291 | 0.061011663 | n | n | y | protein coding gene |
| Ddx6 | DEAD-box helicase 6 | 13209 | MGI:104976 | -0.233 | 0.061905951 | n | n | y | protein coding gene |
| Cdkn2aipnl | CDKN2A interacting protein N-terminal lik | 52626 | MGI:1261797 | 0.155 | 0.063122282 | n | n | y | protein coding gene |
| Golga1 | golgin A1 | 76899 | MGI:1924149 | 0.275 | 0.063338565 | n | n | y | protein coding gene |
| Prkca | protein kinase C, alpha | 18750 | MGI:97595 | 0.306 | 0.063646223 | n | n | y | protein coding gene |
| Ankrd24 | ankyrin repeat domain 24 | 70615 | MGI:1890394 | 0.192 | 0.066907281 | n | n | y | protein coding gene |
| Spata7 | spermatogenesis associated 7 | 104871 | MGI:2144877 | -0.255 | 0.067901426 | n | n | y | protein coding gene |
| AU022252 | expressed sequence AU022252 | 230696 | MGI:2140466 | 0.262 | 0.068219154 | n | n | y | protein coding gene |
| Prr14 | proline rich 14 | 233895 | MGI:2384565 | -0.181 | 0.069299074 | n | n | y | protein coding gene |
| Tcp11 | t-complex protein 11 | 21463 | MGI:98544 | 0.479 | 0.069389199 | n | n | y | protein coding gene |
| Atp5me | ATP synthase membrane subunit e | 11958 | MGI:106636 | -0.365 | 0.069947952 | n | n | y | protein coding gene |
| Hps4 | HPS4, biogenesis of lysosomal organelles | 192232 | MGI:2177742 | 0.259 | 0.070362135 | n | n | y | protein coding gene |
| Mipep | mitochondrial intermediate peptidase | 70478 | MGI:1917728 | 0.220 | 0.072182803 | n | n | y | protein coding gene |
| Pknox2 | Pbx/knotted 1 homeobox 2 | 208076 | MGI:2445415 | -0.231 | 0.073650031 | n | n | y | protein coding gene |
| Ppp1r9b | protein phosphatase 1, regulatory subunit | 217124 | MGI:2387581 | -0.492 | 0.073907983 | n | n | y | protein coding gene |
| Tmf1 | TATA element modulatory factor 1 | 232286 | MGI:2684999 | 0.237 | 0.074019366 | n | n | y | protein coding gene |
| Thsd7b | thrombospondin, type I, domain containing | 210417 | MGI:2443925 | 0.339 | 0.074254086 | n | n | y | protein coding gene |
| Otos | otospiralin | 260301 | MGI:2672814 | -1.718 | 0.074284084 | n | n | y | protein coding gene |
| Pierce1 | piercer of microtubule wall 1 | 69327 | MGI:1916577 | 0.501 | 0.076304747 | n | n | y | protein coding gene |
| Zmynd12 | zinc finger, MYND domain containing 12 | 332934 | MGI:2140259 | -0.568 | 0.076670483 | n | n | y | protein coding gene |
| Satb1 | special AT-rich sequence binding protein | 20230 | MGI:105084 | 0.272 | 0.077124215 | n | n | y | protein coding gene |
| Nxnl2 | nucleoredoxin-like 2 | 75124 | MGI:1922374 | -0.551 | 0.078093834 | n | n | y | protein coding gene |
| Card10 | caspase recruitment domain family, memb | 105844 | MGI:2146012 | -0.239 | 0.078893578 | n | n | y | protein coding gene |
| Ormdl3 | ORM1-like 3 (S. cerevisiae) | 66612 | MGI:1913862 | 0.156 | 0.079304864 | n | n | y | protein coding gene |
| Fbxo9 | f-box protein 9 | 71538 | MGI:1918788 | 0.188 | 0.080330667 | n | n | y | protein coding gene |
| Rab11fip4 | RAB11 family interacting protein 4 (class I | 268451 | MGI:2442920 | 0.397 | 0.080568711 | n | n | y | protein coding gene |
| Morn5 | MORN repeat containing 5 | 75495 | MGI:1922745 | 0.459 | 0.081891005 | n | n | y | protein coding gene |
| Snap25 | synaptosomal-associated protein 25 | 20614 | MGI:98331 | -0.319 | 0.083969355 | n | n | y | protein coding gene |
| Ybx3 | Y box protein 3 | 56449 | MGI:2137670 | -0.576 | 0.084052988 | n | n | y | protein coding gene |
| Syne4 | spectrin repeat containing, nuclear envelo | 233066 | MGI:2141950 | 0.145 | 0.088727494 | n | n | y | protein coding gene |
| Sox12 | SRY (sex determining region Y)-box 12 | 20667 | MGI:98360 | -0.340 | 0.090840656 | n | n | y | protein coding gene |
| Abca3 | ATP-binding cassette, sub-family A memb | 27410 | MGI:1351617 | 0.216 | 0.09289952 | n | n | y | protein coding gene |

| | | | | | | | | | |
|---|---|---|---|---|---|---|---|---|---|
| Dll4 | delta like canonical Notch ligand 4 | 54485 | MGI:1859388 | 0.298 | 0.093713449 | n | n | y | protein coding gene |
| Mfsd1 | major facilitator superfamily domain conta | 66868 | MGI:1914118 | 0.169 | 0.099505521 | n | n | y | protein coding gene |
| Trim62 | tripartite motif-containing 62 | 67525 | MGI:1914775 | 0.263 | 0.101788503 | n | n | y | protein coding gene |
| Otud3 | OTU domain containing 3 | 73162 | MGI:1920412 | 0.220 | 0.10231107 | n | n | y | protein coding gene |
| Dlst | dihydrolipoamide S-succinyltransferase | 78920 | MGI:1926170 | -0.207 | 0.102613886 | n | n | y | protein coding gene |
| Thoc2l | THO complex subunit 2-like | 100042165 | MGI:3040669 | -0.279 | 0.103440944 | n | n | y | protein coding gene |
| Man1a2 | mannosidase, alpha, class 1A, member 2 | 17156 | MGI:104676 | 0.184 | 0.104556218 | n | n | y | protein coding gene |
| Meak7 | MTOR associated protein, eak-7 homolog | 74347 | MGI:1921597 | 0.199 | 0.105917777 | n | n | y | protein coding gene |
| Prkaa2 | protein kinase, AMP-activated, alpha 2 ca | 108079 | MGI:1336173 | 0.253 | 0.107518826 | n | n | y | protein coding gene |
| Smim26 | small integral membrane protein 26 | 228715 | MGI:2685407 | -0.412 | 0.108664975 | n | n | y | protein coding gene |
| Pacrg | PARK2 co-regulated | 69310 | MGI:1916560 | 0.231 | 0.110678652 | n | n | y | protein coding gene |
| Mib2 | mindbomb E3 ubiquitin protein ligase 2 | 76580 | MGI:2679684 | 0.279 | 0.111855505 | n | n | y | protein coding gene |
| Mtpn | myotrophin | 14489 | MGI:99445 | 0.147 | 0.114364923 | n | n | y | protein coding gene |
| Nsf | N-ethylmaleimide sensitive fusion protein | 18195 | MGI:104560 | 0.209 | 0.115584476 | n | n | y | protein coding gene |
| Mri1 | methylthioribose-1-phosphate isomerase | 67873 | MGI:1915123 | -0.270 | 0.116531698 | n | n | y | protein coding gene |
| Mllt6 | myeloid/lymphoid or mixed-lineage leuken | 246198 | MGI:1935145 | 0.720 | 0.117699449 | n | n | y | protein coding gene |
| Kcnh2 | potassium voltage-gated channel, subfam | 16511 | MGI:1341722 | 0.340 | 0.122320647 | n | n | y | protein coding gene |
| Arhgef7 | Rho guanine nucleotide exchange factor | 54126 | MGI:1860493 | 0.198 | 0.12823311 | n | n | y | protein coding gene |
| Mtmr11 | myotubularin related protein 11 | 194126 | MGI:2652817 | -0.143 | 0.12927345 | n | n | y | protein coding gene |
| Mprip | myosin phosphatase Rho interacting prote | 26936 | MGI:1349438 | -0.196 | 0.130218357 | n | n | y | protein coding gene |
| Dctn3 | dynactin 3 | 53598 | MGI:1859251 | -0.161 | 0.132159197 | n | n | y | protein coding gene |
| Frmd4b | FERM domain containing 4B | 232288 | MGI:2141794 | 0.196 | 0.138070007 | n | n | y | protein coding gene |
| Fam217b | family with sequence similarity 217, memb | 71532 | MGI:1918782 | -0.322 | 0.140366292 | n | n | y | protein coding gene |
| Usp25 | ubiquitin specific peptidase 25 | 30940 | MGI:1353655 | 0.737 | 0.14287518 | n | n | y | protein coding gene |
| Ift22 | intraflagellar transport 22 | 67286 | MGI:1914536 | 0.178 | 0.143086595 | n | n | y | protein coding gene |
| Rpl5 | ribosomal protein L5 | 100503670 | MGI:102854 | -0.251 | 0.145251075 | n | n | y | protein coding gene |
| Wdr24 | WD repeat domain 24 | 268933 | MGI:2446285 | 0.153 | 0.148094389 | n | n | y | protein coding gene |
| Mgat5b | mannoside acetylglucosaminyltransferase | 268510 | MGI:3606200 | 0.384 | 0.15036656 | n | n | y | protein coding gene |
| Rims3 | regulating synaptic membrane exocytosis | 242662 | MGI:2443331 | 0.407 | 0.150533746 | n | n | y | protein coding gene |
| Thap3 | THAP domain containing, apoptosis assoc | 69876 | MGI:1917126 | -0.233 | 0.150695937 | n | n | y | protein coding gene |
| Hs3st3b1 | heparan sulfate (glucosamine) 3-O-sulfotr | 54710 | MGI:1333853 | 0.316 | 0.151644803 | n | n | y | protein coding gene |
| Comtd1 | catechol-O-methyltransferase domain con | 69156 | MGI:1916406 | -0.360 | 0.154546581 | n | n | y | protein coding gene |
| Ndufa5 | NADH:ubiquinone oxidoreductase subunit | 68202 | MGI:1915452 | -0.481 | 0.155153452 | n | n | y | protein coding gene |
| Cgn | cingulin | 70737 | MGI:1927237 | -0.153 | 0.155778978 | n | n | y | protein coding gene |
| Rsbn1 | rosbin, round spermatid basic protein 1 | 229675 | MGI:2444993 | 0.257 | 0.156767763 | n | n | y | protein coding gene |
| Neurl1b | neuralized E3 ubiquitin protein ligase 1B | 240055 | MGI:3643092 | -0.416 | 0.162614789 | n | n | y | protein coding gene |
| Obscn | obscurin, cytoskeletal calmodulin and titin | 380698 | MGI:2681862 | -0.431 | 0.165166671 | n | n | y | protein coding gene |
| Bbs1 | Bardet-Biedl syndrome 1 | 52028 | MGI:1277215 | -0.304 | 0.165845728 | n | n | y | protein coding gene |
| Ccdc40 | coiled-coil domain containing 40 | 207607 | MGI:2443893 | -0.228 | 0.166462144 | n | n | y | protein coding gene |
| Arhgap39 | Rho GTPase activating protein 39 | 223666 | MGI:107858 | 0.357 | 0.166720381 | n | n | y | protein coding gene |
| Fry | FRY microtubule binding protein | 320365 | MGI:2443895 | -0.258 | 0.167964852 | n | n | y | protein coding gene |
| Cfap77 | cilia and flagella associated protein 77 | 329375 | MGI:2685669 | 0.456 | 0.175984456 | n | n | y | protein coding gene |
| Nrxn3 | neurexin III | 18191 | MGI:1096389 | -0.170 | 0.195183977 | n | n | y | protein coding gene |
| Slc6a17 | solute carrier family 6 (neurotransmitter tra | 229706 | MGI:2442535 | 0.442 | 0.19826764 | n | n | y | protein coding gene |
| Madd | MAP-kinase activating death domain | 228355 | MGI:2444672 | 0.201 | 0.201801186 | n | n | y | protein coding gene |
| Rem1 | rad and gem related GTP binding protein | 19700 | MGI:1097696 | 0.233 | 0.206629779 | n | n | y | protein coding gene |
| Nmb | neuromedin B | 68039 | MGI:1915289 | -0.293 | 0.211581271 | n | n | y | protein coding gene |
| Tango6 | transport and golgi organization 6 | 272538 | MGI:2142786 | 0.205 | 0.214547564 | n | n | y | protein coding gene |
| Gpr4 | G protein-coupled receptor 4 | 319197 | MGI:2441992 | 0.222 | 0.220491388 | n | n | y | protein coding gene |
| Ppp1cb | protein phosphatase 1 catalytic subunit be | 19046 | MGI:104871 | -0.119 | 0.226316942 | n | n | y | protein coding gene |
| Osbp | oxysterol binding protein | 76303 | MGI:97447 | -0.336 | 0.226325218 | n | n | y | protein coding gene |
| Cyb561d2 | cytochrome b-561 domain containing 2 | 56368 | MGI:1929280 | 0.156 | 0.227821953 | n | n | y | protein coding gene |
| Mzt2 | mitotic spindle organizing protein 2 | 72083 | MGI:1922845 | -0.231 | 0.235645791 | n | n | y | protein coding gene |
| Ctbp2 | C-terminal binding protein 2 | 13017 | MGI:1201686 | 0.137 | 0.247176729 | n | n | y | protein coding gene |
| Nup210 | nucleoporin 210 | 54563 | MGI:1859555 | 0.224 | 0.248742896 | n | n | y | protein coding gene |
| Usp2 | ubiquitin specific peptidase 2 | 53376 | MGI:1858178 | -0.147 | 0.249316026 | n | n | y | protein coding gene |
| Ptgfrn | prostaglandin F2 receptor negative regula | 19221 | MGI:1277114 | -0.429 | 0.2493371 | n | n | y | protein coding gene |
| Mcf2l | mcf.2 transforming sequence-like | 17207 | MGI:103263 | 0.157 | 0.254812665 | n | n | y | protein coding gene |
| Nin | ninein | 18080 | MGI:105108 | 0.188 | 0.256518874 | n | n | y | protein coding gene |
| Nup214 | nucleoporin 214 | 227720 | MGI:1095411 | 0.255 | 0.260297089 | n | n | y | protein coding gene |
| Wdr35 | WD repeat domain 35 | 74682 | MGI:1921932 | 0.190 | 0.261915841 | n | n | y | protein coding gene |
| Nbas | neuroblastoma amplified sequence | 71169 | MGI:1918419 | -0.165 | 0.268384179 | n | n | y | protein coding gene |
| Usp48 | ubiquitin specific peptidase 48 | 170707 | MGI:2158502 | 0.177 | 0.277873493 | n | n | y | protein coding gene |
| Fam193a | family with sequence homology 193, mem | 231128 | MGI:2447768 | -0.216 | 0.279850108 | n | n | y | protein coding gene |

| | | | | | | | | | |
|---|---|---|---|---|---|---|---|---|---|
| Cep104 | centrosomal protein 104 | 230967 | MGI:2687282 | 0.195 | 0.289810346 | n | n | y | protein coding gene |
| Ubqln1 | ubiquilin 1 | 56085 | MGI:1860276 | 0.132 | 0.29741587 | n | n | y | protein coding gene |
| Tfdp2 | transcription factor Dp 2 | 211586 | MGI:107167 | -0.169 | 0.300061456 | n | n | y | protein coding gene |
| Pcgf6 | polycomb group ring finger 6 | 71041 | MGI:1918291 | -0.195 | 0.301129101 | n | n | y | protein coding gene |
| Nrxn2 | neurexin II | 18190 | MGI:1096362 | -0.515 | 0.308476106 | n | n | y | protein coding gene |
| Bmyc | brain expressed myelocytomatosis oncog | 107771 | MGI:88184 | -0.132 | 0.308708343 | n | n | y | protein coding gene |
| Zfp385a | zinc finger protein 385A | 29813 | MGI:1352495 | 0.240 | 0.31663477 | n | n | y | protein coding gene |
| Ndrg3 | N-myc downstream regulated gene 3 | 29812 | MGI:1352499 | 0.128 | 0.322264503 | n | n | y | protein coding gene |
| Kcna10 | potassium voltage-gated channel, shaker- | 242151 | MGI:3037820 | 0.219 | 0.327732857 | n | n | y | protein coding gene |
| Gng3 | guanine nucleotide binding protein (G prot | 14704 | MGI:102704 | -0.404 | 0.340486569 | n | n | y | protein coding gene |
| Adprhl1 | ADP-ribosylhydrolase like 1 | 234072 | MGI:2442168 | -0.205 | 0.343210259 | n | n | y | protein coding gene |
| Snrk | SNF related kinase | 20623 | MGI:108104 | 0.172 | 0.344797097 | n | n | y | protein coding gene |
| Mark4 | MAP/microtubule affinity regulating kinase | 232944 | MGI:1920955 | -0.302 | 0.347742636 | n | n | y | protein coding gene |
| Mblac2 | metallo-beta-lactamase domain containing | 72852 | MGI:1920102 | 0.256 | 0.350001173 | n | n | y | protein coding gene |
| Inpp5f | inositol polyphosphate-5-phosphatase F | 101490 | MGI:2141867 | 0.192 | 0.353557862 | n | n | y | protein coding gene |
| Ccdc88c | coiled-coil domain containing 88C | 68339 | MGI:1915589 | 0.195 | 0.354557409 | n | n | y | protein coding gene |
| Adck1 | aarF domain containing kinase 1 | 72113 | MGI:1919363 | -0.139 | 0.359348262 | n | n | y | protein coding gene |
| Rxra | retinoid X receptor alpha | 20181 | MGI:98214 | -0.284 | 0.363478282 | n | n | y | protein coding gene |
| Eps8l2 | EPS8-like 2 | 98845 | MGI:2138828 | 0.129 | 0.364950287 | n | n | y | protein coding gene |
| Akap9 | A kinase anchor protein 9 | 100986 | MGI:2178217 | 0.162 | 0.366587768 | n | n | y | protein coding gene |
| Pak3 | p21 (RAC1) activated kinase 3 | 18481 | MGI:1339656 | 0.144 | 0.370833237 | n | n | y | protein coding gene |
| Cdc25b | cell division cycle 25B | 12531 | MGI:99701 | -0.170 | 0.371375162 | n | n | y | protein coding gene |
| Marchf9 | membrane associated ring-CH-type finger | 216438 | MGI:2446144 | -0.589 | 0.372903271 | n | n | y | protein coding gene |
| Coprs | coordinator of PRMT5, differentiation stim | 66423 | MGI:1913673 | 0.211 | 0.382110974 | n | n | y | protein coding gene |
| Dennd5b | DENN domain containing 5B | 320560 | MGI:2444273 | 0.283 | 0.385485636 | n | n | y | protein coding gene |
| Dhx33 | DEAH-box helicase 33 | 216877 | MGI:2445102 | 0.180 | 0.404813102 | n | n | y | protein coding gene |
| Triobp | TRIO and F-actin binding protein | 110253 | MGI:1349410 | -0.128 | 0.404948339 | n | n | y | protein coding gene |
| Cars1 | cysteinyl-tRNA synthetase 1 | 27267 | MGI:1351477 | 0.137 | 0.411749616 | n | n | y | protein coding gene |
| Pcbp3 | poly(rC) binding protein 3 | 59093 | MGI:1890470 | -0.140 | 0.411954211 | n | n | y | protein coding gene |
| Tbc1d7 | TBC1 domain family, member 7 | 67046 | MGI:1914296 | -0.135 | 0.415640529 | n | n | y | protein coding gene |
| Mob2 | MOB kinase activator 2 | 101513 | MGI:1919891 | -0.122 | 0.419758515 | n | n | y | protein coding gene |
| Rbm24 | RNA binding motif protein 24 | 666794 | MGI:3610364 | -0.283 | 0.420751639 | n | n | y | protein coding gene |
| Mob1a | MOB kinase activator 1A | 232157 | MGI:2442631 | 0.184 | 0.429066589 | n | n | y | protein coding gene |
| Rptor | regulatory associated protein of MTOR, cc | 74370 | MGI:1921620 | 0.198 | 0.441908644 | n | n | y | protein coding gene |
| Lsm7 | LSM7 homolog, U6 small nuclear RNA an | 66094 | MGI:1913344 | -0.275 | 0.442018529 | n | n | y | protein coding gene |
| Trim33 | tripartite motif-containing 33 | 94093 | MGI:2137357 | -0.211 | 0.450496581 | n | n | y | protein coding gene |
| 1110032F04Rik | RIKEN cDNA 1110032F04 gene | 68725 | MGI:1915975 | 0.380 | 0.451626338 | n | n | y | protein coding gene |
| Dedd2 | death effector domain-containing DNA bin | 67379 | MGI:1914629 | -0.254 | 0.453147334 | n | n | y | protein coding gene |
| Dscaml1 | DS cell adhesion molecule like 1 | 114873 | MGI:2150309 | 0.195 | 0.458065897 | n | n | y | protein coding gene |
| Rmnd5a | required for meiotic nuclear division 5 hon | 68477 | MGI:1915727 | -0.237 | 0.468040077 | n | n | y | protein coding gene |
| Krit1 | KRIT1, ankyrin repeat containing | 79264 | MGI:1930618 | 0.153 | 0.469126194 | n | n | y | protein coding gene |
| Atp13a1 | ATPase type 13A1 | 170759 | MGI:2180801 | -0.119 | 0.46950763 | n | n | y | protein coding gene |
| Myo7a | myosin VIIA | 17921 | MGI:104510 | 0.275 | 0.46957917 | n | n | y | protein coding gene |
| Col7a1 | collagen, type VII, alpha 1 | 12836 | MGI:88462 | -0.447 | 0.472419608 | n | n | y | protein coding gene |
| Sptan1 | spectrin alpha, non-erythrocytic 1 | 20740 | MGI:98386 | -0.133 | 0.473794971 | n | n | y | protein coding gene |
| Hagh | hydroxyacyl glutathione hydrolase | 14651 | MGI:95745 | 0.142 | 0.48117529 | n | n | y | protein coding gene |
| Thop1 | thimet oligopeptidase 1 | 50492 | MGI:1354165 | 0.138 | 0.483912306 | n | n | y | protein coding gene |
| Asic1 | acid-sensing ion channel 1 | 11419 | MGI:1194915 | 0.129 | 0.492853074 | n | n | y | protein coding gene |
| Slc45a4 | solute carrier family 45, member 4 | 106068 | MGI:2146236 | -0.321 | 0.497234484 | n | n | y | protein coding gene |
| Ubr4 | ubiquitin protein ligase E3 component n-re | 69116 | MGI:1916366 | 0.152 | 0.503266095 | n | n | y | protein coding gene |
| Rc3h2 | ring finger and CCCH-type zinc finger don | 319817 | MGI:2442789 | 0.184 | 0.5043354 | n | n | y | protein coding gene |
| Pak1 | p21 (RAC1) activated kinase 1 | 18479 | MGI:1339975 | 0.122 | 0.51374927 | n | n | y | protein coding gene |
| Mepce | methylphosphate capping enzyme | 231803 | MGI:106477 | -0.224 | 0.518846695 | n | n | y | protein coding gene |
| Hdac4 | histone deacetylase 4 | 208727 | MGI:3036234 | -0.229 | 0.522974998 | n | n | y | protein coding gene |
| Cibar2 | CBY1 interacting BAR domain containing | 436062 | MGI:3588213 | -0.419 | 0.527190303 | n | n | y | protein coding gene |
| Necap1 | NECAP endocytosis associated 1 | 67602 | MGI:1914852 | 0.111 | 0.537949017 | n | n | y | protein coding gene |
| Mlxip | MLX interacting protein | 208104 | MGI:2141183 | -0.157 | 0.540306489 | n | n | y | protein coding gene |
| Ccdc181 | coiled-coil domain containing 181 | 74895 | MGI:1922145 | -0.136 | 0.542238079 | n | n | y | protein coding gene |
| Ahi1 | Abelson helper integration site 1 | 52906 | MGI:87971 | 0.160 | 0.542820366 | n | n | y | protein coding gene |
| Pdzph1 | PDZ and pleckstrin homology domains 1 | 69239 | MGI:1916489 | 0.221 | 0.547271702 | n | n | y | protein coding gene |
| Prepl | prolyl endopeptidase-like | 213760 | MGI:2441932 | 0.127 | 0.557204726 | n | n | y | protein coding gene |
| Gm1043 | predicted gene 1043 | 381634 | MGI:2685889 | 0.275 | 0.562533578 | n | n | y | protein coding gene |
| Strbp | spermatid perinuclear RNA binding protei | 20744 | MGI:104626 | 0.215 | 0.572321969 | n | n | y | protein coding gene |
| Loxhd1 | lipoxygenase homology domains 1 | 240411 | MGI:1914609 | 0.178 | 0.581660949 | n | n | y | protein coding gene |

| | | | | | | | | | |
|---|---|---|---|---|---|---|---|---|---|
| Cimip2c | ciliary microtubule inner protein 2C | 75434 | MGI:1922684 | -0.209 | 0.588374452 | n | n | y | protein coding gene |
| Fam81b | family with sequence similarity 81, membe | 238726 | MGI:2685122 | -0.235 | 0.598863662 | n | n | y | protein coding gene |
| Gdap1l1 | ganglioside-induced differentiation-associ | 228858 | MGI:2385163 | 0.191 | 0.615555398 | n | n | y | protein coding gene |
| Ankrd17 | ankyrin repeat domain 17 | 81702 | MGI:1932101 | -0.165 | 0.618278336 | n | n | y | protein coding gene |
| Pcdh7 | protocadherin 7 | 54216 | MGI:1860487 | 0.162 | 0.623891673 | n | n | y | protein coding gene |
| Katnip | katanin interacting protein | 233865 | MGI:2442760 | 0.235 | 0.62420869 | n | n | y | protein coding gene |
| Runx1t1 | RUNX1 translocation partner 1 | 12395 | MGI:104793 | 0.147 | 0.632101793 | n | n | y | protein coding gene |
| Dync2h1 | dynein cytoplasmic 2 heavy chain 1 | 110350 | MGI:107736 | 0.258 | 0.639480949 | n | n | y | protein coding gene |
| Cacna1d | calcium channel, voltage-dependent, L typ | 12289 | MGI:88293 | 0.144 | 0.640308462 | n | n | y | protein coding gene |
| Ap3b2 | adaptor-related protein complex 3, beta 2 | 11775 | MGI:1100869 | 0.154 | 0.646160167 | n | n | y | protein coding gene |
| Wdr7 | WD repeat domain 7 | 104082 | MGI:1860197 | 0.283 | 0.646725641 | n | n | y | protein coding gene |
| Rnf8 | ring finger protein 8 | 58230 | MGI:1929069 | -0.100 | 0.653495296 | n | n | y | protein coding gene |
| Hectd4 | HECT domain E3 ubiquitin protein ligase 4 | 269700 | MGI:3647820 | -0.125 | 0.654413998 | n | n | y | protein coding gene |
| Gramd4 | GRAM domain containing 4 | 223752 | MGI:2676308 | 0.099 | 0.662120613 | n | n | y | protein coding gene |
| Afdn | afadin, adherens junction formation factor | 17356 | MGI:1314653 | 0.240 | 0.662697419 | n | n | y | protein coding gene |
| Fbrs | fibrosin | 14123 | MGI:104648 | -0.271 | 0.676836296 | n | n | y | protein coding gene |
| Trit1 | tRNA isopentenyltransferase 1 | 66966 | MGI:1914216 | -0.122 | 0.677486494 | n | n | y | protein coding gene |
| Espn | espin | 56226 | MGI:1861630 | 0.169 | 0.680705294 | n | n | y | protein coding gene |
| Cables1 | CDK5 and Abl enzyme substrate 1 | 63955 | MGI:1927065 | -0.263 | 0.690504497 | n | n | y | protein coding gene |
| C2cd2l | C2 calcium-dependent domain containing | 71764 | MGI:1919014 | 0.190 | 0.705023354 | n | n | y | protein coding gene |
| Nfkb1 | nuclear factor of kappa light polypeptide g | 18033 | MGI:97312 | -0.228 | 0.717319596 | n | n | y | protein coding gene |
| Rab3ip | RAB3A interacting protein | 216363 | MGI:105933 | 0.105 | 0.723179073 | n | n | y | protein coding gene |
| Adgrl3 | adhesion G protein-coupled receptor L3 | 319387 | MGI:2441950 | -0.211 | 0.724096266 | n | n | y | protein coding gene |
| Stard10 | StAR related lipid transfer domain contain | 56018 | MGI:1860093 | 0.143 | 0.746550541 | n | n | y | protein coding gene |
| Pitpna | phosphatidylinositol transfer protein, alpha | 18738 | MGI:99887 | 0.091 | 0.747618228 | n | n | y | protein coding gene |
| Spa17 | sperm autoantigenic protein 17 | 20686 | MGI:1333778 | 0.223 | 0.76808558 | n | n | y | protein coding gene |
| Rassf2 | Ras association (RalGDS/AF-6) domain fa | 215653 | MGI:2442060 | -0.181 | 0.801639224 | n | n | y | protein coding gene |
| Stxbp1 | syntaxin binding protein 1 | 20910 | MGI:107363 | -0.138 | 0.828061426 | n | n | y | protein coding gene |
| Fbxo16 | F-box protein 16 | 50759 | MGI:1354706 | 0.140 | 0.851583804 | n | n | y | protein coding gene |
| Wdr31 | WD repeat domain 31 | 71354 | MGI:1918604 | -0.142 | 0.85519053 | n | n | y | protein coding gene |
| Impdh1 | inosine monophosphate dehydrogenase 1 | 23917 | MGI:96567 | -0.129 | 0.859922673 | n | n | y | protein coding gene |
| Tcerg1l | transcription elongation regulator 1-like | 70571 | MGI:1917821 | 0.239 | 0.866507045 | n | n | y | protein coding gene |
| Ift172 | intraflagellar transport 172 | 67661 | MGI:2682064 | 0.135 | 0.868017465 | n | n | y | protein coding gene |
| Mier1 | MEIR1 treanscription regulator | 71148 | MGI:1918398 | -0.120 | 0.86863136 | n | n | y | protein coding gene |
| Prpf18 | pre-mRNA processing factor 18 | 67229 | MGI:1914479 | -0.130 | 0.868752173 | n | n | y | protein coding gene |
| Nt5m | 5',3'-nucleotidase, mitochondrial | 103850 | MGI:1917127 | 0.151 | 0.872647658 | n | n | y | protein coding gene |
| Agap3 | ArfGAP with GTPase domain, ankyrin rep | 213990 | MGI:2183446 | -0.337 | 0.875340469 | n | n | y | protein coding gene |
| Mpdz | multiple PDZ domain crumbs cell polarity c | 17475 | MGI:1343489 | 0.183 | 0.884373095 | n | n | y | protein coding gene |
| Mgat4b | mannoside acetylglucosaminyltransferase | 103534 | MGI:2143974 | -0.229 | 0.891013293 | n | n | y | protein coding gene |
| 4930470P17Rik | RIKEN cDNA 4930470P17 gene | 67637 | MGI:1914887 | 0.479 | 0.897665959 | n | n | y | protein coding gene |
| Dym | dymeclin | 69190 | MGI:1918480 | 0.104 | 0.90575472 | n | n | y | protein coding gene |
| Cdh23 | cadherin related 23 (otocadherin) | 22295 | MGI:1890219 | -0.268 | 0.917388453 | n | n | y | protein coding gene |
| Rpgr | retinitis pigmentosa GTPase regulator | 19893 | MGI:1344037 | 0.160 | 0.926967548 | n | n | y | protein coding gene |
| Slc7a14 | solute carrier family 7 (cationic amino acid | 241919 | MGI:3040688 | -0.179 | 0.945795202 | n | n | y | protein coding gene |
| Ubl3 | ubiquitin-like 3 | 24109 | MGI:1344373 | 0.126 | 0.974144003 | n | n | y | protein coding gene |
| Fam43a | family with sequence similarity 43, membe | 224093 | MGI:2676309 | 0.215 | 0.990327833 | n | n | y | protein coding gene |
| Cdc14a | CDC14 cell division cycle 14A | 229776 | MGI:2442676 | 0.168 | 0.993827622 | n | n | y | protein coding gene |
| Atp6v0a1 | ATPase, H+ transporting, lysosomal V0 su | 11975 | MGI:103286 | 0.153 | 0.993976102 | n | n | y | protein coding gene |
| Ralgds | ral guanine nucleotide dissociation stimula | 19730 | MGI:107485 | -0.103 | 1 | n | n | y | protein coding gene |
| Nell1 | NEL-like 1 | 338352 | MGI:2443902 | -0.206 | 1 | n | n | y | protein coding gene |
| Wdr12 | WD repeat domain 12 | 57750 | MGI:1927241 | 0.124 | 1 | n | n | y | protein coding gene |
| Bace1 | beta-site APP cleaving enzyme 1 | 23821 | MGI:1346542 | 0.186 | 1 | n | n | y | protein coding gene |
| Brf1 | BRF1, RNA polymerase III transcription in | 72308 | MGI:1919558 | -0.090 | 1 | n | n | y | protein coding gene |
| Nhsl3 | NHS like 3 | 97130 | MGI:2140651 | 0.122 | 1 | n | n | y | protein coding gene |
| Agfg2 | ArfGAP with FG repeats 2 | 231801 | MGI:2443267 | 0.147 | 1 | n | n | y | protein coding gene |
| Paqr9 | progestin and adipoQ receptor family mem | 75552 | MGI:1922802 | 0.197 | 1 | n | n | y | protein coding gene |
| Neurl1a | neuralized E3 ubiquitin protein ligase 1A | 18011 | MGI:1334263 | -0.240 | 1 | n | n | y | protein coding gene |
| Utp14a | UTP14A small subunit processome compo | 72554 | MGI:1919804 | 0.124 | 1 | n | n | y | protein coding gene |
| Trnp1 | TMF1-regulated nuclear protein 1 | 69539 | MGI:1916789 | 0.325 | 1 | n | n | y | protein coding gene |
| Reep1 | receptor accessory protein 1 | 52250 | MGI:1098827 | -0.143 | 1 | n | n | y | protein coding gene |
| Btbd9 | BTB domain containing 9 | 224671 | MGI:1916625 | -0.105 | 1 | n | n | y | protein coding gene |
| Tmem41a | transmembrane protein 41a | 66664 | MGI:1913914 | 0.097 | 1 | n | n | y | protein coding gene |
| Ttll3 | tubulin tyrosine ligase-like family, member | 101100 | MGI:2141418 | -0.139 | 1 | n | n | y | protein coding gene |
| Fam193b | family with sequence similarity 193, memb | 212483 | MGI:2385851 | 0.177 | 1 | n | n | y | protein coding gene |

| | | | | | | | | | |
|---|---|---|---|---|---|---|---|---|---|
| Mfsd6 | major facilitator superfamily domain conta | 98682 | MGI:1922925 | 0.106 | 1 | n | n | y | protein coding gene |
| Chrna9 | cholinergic receptor, nicotinic, alpha polyp | 231252 | MGI:1202403 | 0.091 | 1 | n | n | y | protein coding gene |
| Pcgf3 | polycomb group ring finger 3 | 69587 | MGI:1916837 | 0.146 | 1 | n | n | y | protein coding gene |
| Capn7 | calpain 7 | 12339 | MGI:1338030 | 0.090 | 1 | n | n | y | protein coding gene |
| Nme5 | NME/NM23 family member 5 | 75533 | MGI:1922783 | -0.132 | 1 | n | n | y | protein coding gene |
| Trappc5 | trafficking protein particle complex 5 | 66682 | MGI:1913932 | -0.128 | 1 | n | n | y | protein coding gene |
| Synj1 | synaptojanin 1 | 104015 | MGI:1354961 | 0.211 | 1 | n | n | y | protein coding gene |
| Gpsm2 | G-protein signalling modulator 2 (AGS3-lik | 76123 | MGI:1923373 | -0.090 | 1 | n | n | y | protein coding gene |
| Tnks2 | tankyrase, TRF1-interacting ankyrin-relate | 74493 | MGI:1921743 | 0.269 | 1 | n | n | y | protein coding gene |
| Rnf11 | ring finger protein 11 | 29864 | MGI:1352759 | 0.124 | 1 | n | n | y | protein coding gene |
| Adipor2 | adiponectin receptor 2 | 68465 | MGI:93830 | -0.133 | 1 | n | n | y | protein coding gene |
| Pdgfra | platelet derived growth factor receptor, alp | 18595 | MGI:97530 | 0.123 | 1 | n | n | y | protein coding gene |
| Coq10a | coenzyme Q10A | 210582 | MGI:2684847 | 0.207 | 1 | n | n | y | protein coding gene |
| Fem1a | fem 1 homolog a | 14154 | MGI:1335089 | -0.142 | 1 | n | n | y | protein coding gene |
| Wipf2 | WAS/WASL interacting protein family, me | 68524 | MGI:1924462 | 0.134 | 1 | n | n | y | protein coding gene |
| Dot1l | DOT1 like histone lysine methyltransferas | 208266 | MGI:2143886 | -0.280 | 1 | n | n | y | protein coding gene |
| Bdnf | brain derived neurotrophic factor | 12064 | MGI:88145 | 0.104 | 1 | n | n | y | protein coding gene |
| Arhgdig | Rho GDP dissociation inhibitor gamma | 14570 | MGI:108430 | -0.213 | 1 | n | n | y | protein coding gene |
| Ppp1r27 | protein phosphatase 1, regulatory subunit | 68701 | MGI:1915951 | -0.095 | 1 | n | n | y | protein coding gene |
| Phrf1 | PHD and ring finger domains 1 | 101471 | MGI:2141847 | 0.114 | 1 | n | n | y | protein coding gene |
| Prkag2 | protein kinase, AMP-activated, gamma 2 n | 108099 | MGI:1336153 | 0.114 | 1 | n | n | y | protein coding gene |
| Pou4f3 | POU domain, class 4, transcription factor | 18998 | MGI:102523 | 0.112 | 1 | n | n | y | protein coding gene |
| Rgs11 | regulator of G-protein signaling 11 | 50782 | MGI:1354739 | 0.102 | 1 | n | n | y | protein coding gene |
| Misp3 | MISP family member 3 | 70134 | MGI:1917384 | -0.311 | 1 | n | n | y | protein coding gene |
| Cops3 | COP9 signalosome subunit 3 | 26572 | MGI:1349409 | -0.091 | 1 | n | n | y | protein coding gene |
| Chd8 | chromodomain helicase DNA binding prot | 67772 | MGI:1915022 | -0.118 | 1 | n | n | y | protein coding gene |
| Espnl | espin-like | 227357 | MGI:2685402 | -0.161 | 1 | n | n | y | protein coding gene |
| Gtf3c4 | general transcription factor IIIC, polypeptic | 269252 | MGI:2138937 | -0.140 | 1 | n | n | y | protein coding gene |
| Zbtb7a | zinc finger and BTB domain containing 7a | 16969 | MGI:1335091 | -0.128 | 1 | n | n | y | protein coding gene |
| Fhip2a | FHF complex subunit HOOK interacting p | 226252 | MGI:2147545 | 0.134 | 1 | n | n | y | protein coding gene |
| Kcnn2 | potassium intermediate/small conductanc | 140492 | MGI:2153182 | -0.165 | 1 | n | n | y | protein coding gene |
| Lrpprc | leucine-rich PPR-motif containing | 72416 | MGI:1919666 | 0.113 | 1 | n | n | y | protein coding gene |
| Ddit3 | DNA-damage inducible transcript 3 | 13198 | MGI:109247 | -0.162 | 1 | n | n | y | protein coding gene |
| Arid3b | AT-rich interaction domain 3B | 56380 | MGI:1930768 | 0.166 | 1 | n | n | y | protein coding gene |
| Srrm4 | serine/arginine repetitive matrix 4 | 68955 | MGI:1916205 | 0.186 | 1 | n | n | y | protein coding gene |
| Ift81 | intraflagellar transport 81 | 12589 | MGI:1098597 | 0.108 | 1 | n | n | y | protein coding gene |
| Ube3b | ubiquitin protein ligase E3B | 117146 | MGI:1891295 | 0.082 | 1 | n | n | y | protein coding gene |
| Rnf38 | ring finger protein 38 | 73469 | MGI:1920719 | 0.095 | 1 | n | n | y | protein coding gene |
| Ints10 | integrator complex subunit 10 | 70885 | MGI:1918135 | -0.099 | 1 | n | n | y | protein coding gene |
| 2210016L21Rik | RIKEN cDNA 2210016L21 gene | 72357 | MGI:1919607 | -0.089 | 1 | n | n | y | protein coding gene |
| Ankrd37 | ankyrin repeat domain 37 | 654824 | MGI:3603344 | 0.217 | 1 | n | n | y | protein coding gene |
| Ppat | phosphoribosyl pyrophosphate amidotrans | 231327 | MGI:2387203 | -0.108 | 1 | n | n | y | protein coding gene |
| Tsc2 | TSC complex subunit 2 | 22084 | MGI:102548 | -0.152 | 1 | n | n | y | protein coding gene |
| Hyou1 | hypoxia up-regulated 1 | 12282 | MGI:108030 | 0.111 | 1 | n | n | y | protein coding gene |
| Abl2 | ABL proto-oncogene 2, non-receptor tyros | 11352 | MGI:87860 | -0.143 | 1 | n | n | y | protein coding gene |
| Cyb5r1 | cytochrome b5 reductase 1 | 72017 | MGI:1919267 | 0.115 | 1 | n | n | y | protein coding gene |
| Ago2 | argonaute RISC catalytic subunit 2 | 239528 | MGI:2446632 | 0.190 | 1 | n | n | y | protein coding gene |
| Rsph9 | radial spoke head 9 homolog (Chlamydon | 75564 | MGI:1922814 | -0.116 | 1 | n | n | y | protein coding gene |
| Cxcl14 | C-X-C motif chemokine ligand 14 | 57266 | MGI:1888514 | 0.098 | 1 | n | n | y | protein coding gene |
| Dnm1l | dynamin 1-like | 74006 | MGI:1921256 | 0.087 | 1 | n | n | y | protein coding gene |
| Ssbp3 | single-stranded DNA binding protein 3 | 72475 | MGI:1919725 | -0.136 | 1 | n | n | y | protein coding gene |
| Wrap53 | WD repeat containing, antisense to Trp53 | 216853 | MGI:2384933 | -0.106 | 1 | n | n | y | protein coding gene |
| Ppp2r3d | protein phosphatase 2 (formerly 2A), regu | 19054 | MGI:1335093 | -0.191 | 1 | n | n | y | protein coding gene |
| Eml1 | echinoderm microtubule associated protei | 68519 | MGI:1915769 | -0.085 | 1 | n | n | y | protein coding gene |
| Zfp474 | zinc finger protein 474 | 66758 | MGI:1914008 | 0.185 | 1 | n | n | y | protein coding gene |
| Msrb3 | methionine sulfoxide reductase B3 | 320183 | MGI:2443538 | 0.160 | 1 | n | n | y | protein coding gene |
| Rnf149 | ring finger protein 149 | 67702 | MGI:2677438 | 0.126 | 1 | n | n | y | protein coding gene |
| Lmo1 | LIM domain only 1 | 109594 | MGI:102812 | -0.090 | 1 | n | n | y | protein coding gene |
| Epn3 | epsin 3 | 71889 | MGI:1919139 | 0.120 | 1 | n | n | y | protein coding gene |
| Serf1 | small EDRK-rich factor 1 | 20365 | MGI:1337114 | 0.102 | 1 | n | n | y | protein coding gene |
| Aak1 | AP2 associated kinase 1 | 269774 | MGI:1098687 | 0.116 | 1 | n | n | y | protein coding gene |
| Mbd2 | methyl-CpG binding domain protein 2 | 17191 | MGI:1333813 | -0.190 | 1 | n | n | y | protein coding gene |
| Klc2 | kinesin light chain 2 | 16594 | MGI:107953 | -0.072 | 1 | n | n | y | protein coding gene |
| Ehmt2 | euchromatic histone lysine N-methyltransf | 110147 | MGI:2148922 | 0.076 | 1 | n | n | y | protein coding gene |

| | | | | | | | | | |
|---|---|---|---|---|---|---|---|---|---|
| Atn1 | atrophin 1 | 13498 | MGI:104725 | 0.097 | 1 | n | n | y | protein coding gene |
| Selenoi | selenoprotein I | 28042 | MGI:107898 | 0.148 | 1 | n | n | y | protein coding gene |
| Safb2 | scaffold attachment factor B2 | 224902 | MGI:2146808 | 0.115 | 1 | n | n | y | protein coding gene |
| Oscp1 | organic solute carrier partner 1 | 230751 | MGI:1916308 | -0.075 | 1 | n | n | y | protein coding gene |
| Lrp8 | low density lipoprotein receptor-related pr | 16975 | MGI:1340044 | 0.134 | 1 | n | n | y | protein coding gene |
| Dclre1a | DNA cross-link repair 1A | 55947 | MGI:1930042 | 0.179 | 1 | n | n | y | protein coding gene |
| Lrrc73 | leucine rich repeat containing 73 | 224813 | MGI:2684934 | 0.160 | 1 | n | n | y | protein coding gene |
| Mns1 | meiosis-specific nuclear structural protein | 17427 | MGI:107933 | 0.175 | 1 | n | n | y | protein coding gene |
| Zfp667 | zinc finger protein 667 | 384763 | MGI:2442757 | 0.096 | 1 | n | n | y | protein coding gene |
| Slc35c2 | solute carrier family 35, member C2 | 228875 | MGI:2385166 | 0.078 | 1 | n | n | y | protein coding gene |
| Fbxo36 | F-box protein 36 | 66153 | MGI:1289192 | -0.096 | 1 | n | n | y | protein coding gene |
| Arf3 | ADP-ribosylation factor 3 | 11842 | MGI:99432 | -0.082 | 1 | n | n | y | protein coding gene |
| Hook1 | hook microtubule tethering protein 1 | 77963 | MGI:1925213 | 0.124 | 1 | n | n | y | protein coding gene |
| Arnt | aryl hydrocarbon receptor nuclear transloc | 11863 | MGI:88071 | 0.105 | 1 | n | n | y | protein coding gene |
| Larp1 | La ribonucleoprotein 1, translational regula | 73158 | MGI:1890165 | -0.183 | 1 | n | n | y | protein coding gene |
| Polb | polymerase (DNA directed), beta | 18970 | MGI:97740 | 0.112 | 1 | n | n | y | protein coding gene |
| Mreg | melanoregulin | 381269 | MGI:2151839 | -0.074 | 1 | n | n | y | protein coding gene |
| Icmt | isoprenylcysteine carboxyl methyltransfera | 57295 | MGI:1888594 | 0.078 | 1 | n | n | y | protein coding gene |
| Dhx30 | DExH-box helicase 30 | 72831 | MGI:1920081 | 0.091 | 1 | n | n | y | protein coding gene |
| Strn3 | striatin, calmodulin binding protein 3 | 94186 | MGI:2151064 | 0.143 | 1 | n | n | y | protein coding gene |
| Lrrc47 | leucine rich repeat containing 47 | 72946 | MGI:1920196 | -0.141 | 1 | n | n | y | protein coding gene |
| Fbxl16 | F-box and leucine-rich repeat protein 16 | 214931 | MGI:2448488 | -0.149 | 1 | n | n | y | protein coding gene |
| Lrrc10b | leucine rich repeat containing 10B | 278795 | MGI:2685551 | -0.276 | 1 | n | n | y | protein coding gene |
| Stxbp5 | syntaxin binding protein 5 (tomosyn) | 78808 | MGI:1926058 | 0.170 | 1 | n | n | y | protein coding gene |
| Rab11fip3 | RAB11 family interacting protein 3 (class I | 215445 | MGI:2444431 | 0.112 | 1 | n | n | y | protein coding gene |
| Kif3a | kinesin family member 3A | 16568 | MGI:107689 | -0.077 | 1 | n | n | y | protein coding gene |
| Serpine3 | serpin peptidase inhibitor, clade E (nexin, | 319433 | MGI:2442020 | -0.196 | 1 | n | n | y | protein coding gene |
| Bcr | BCR activator of RhoGEF and GTPase | 110279 | MGI:88141 | -0.151 | 1 | n | n | y | protein coding gene |
| Tnrc6b | trinucleotide repeat containing 6b | 213988 | MGI:2443730 | 0.114 | 1 | n | n | y | protein coding gene |
| Josd1 | Josephin domain containing 1 | 74158 | MGI:1921408 | 0.107 | 1 | n | n | y | protein coding gene |
| Sema5b | sema domain, seven thrombospondin rep | 20357 | MGI:107555 | 0.073 | 1 | n | n | y | protein coding gene |
| Rimbp2 | RIMS binding protein 2 | 231760 | MGI:2443235 | 0.180 | 1 | n | n | y | protein coding gene |
| Fam168a | family with sequence similarity 168, memb | 319604 | MGI:2442372 | 0.106 | 1 | n | n | y | protein coding gene |
| Cdc42bpb | CDC42 binding protein kinase beta | 217866 | MGI:2136459 | -0.089 | 1 | n | n | y | protein coding gene |
| Hexd | hexosaminidase D | 238023 | MGI:3605542 | -0.082 | 1 | n | n | y | protein coding gene |
| Sema6b | sema domain, transmembrane domain (TI | 20359 | MGI:1202889 | 0.162 | 1 | n | n | y | protein coding gene |
| Bcl2l1 | BCL2-like 1 | 12048 | MGI:88139 | 0.113 | 1 | n | n | y | protein coding gene |
| Dlg1 | discs large MAGUK scaffold protein 1 | 13383 | MGI:107231 | 0.077 | 1 | n | n | y | protein coding gene |
| Wnk2 | WNK lysine deficient protein kinase 2 | 75607 | MGI:1922857 | -0.086 | 1 | n | n | y | protein coding gene |
| Dync2i1 | dynein 2 intermediate chain 1 | 217935 | MGI:2445085 | 0.085 | 1 | n | n | y | protein coding gene |
| Aacs | acetoacetyl-CoA synthetase | 78894 | MGI:1926144 | 0.079 | 1 | n | n | y | protein coding gene |
| Ranbp10 | RAN binding protein 10 | 74334 | MGI:1921584 | 0.091 | 1 | n | n | y | protein coding gene |
| Pfkp | phosphofructokinase, platelet | 56421 | MGI:1891833 | -0.117 | 1 | n | n | y | protein coding gene |
| Faim2 | Fas apoptotic inhibitory molecule 2 | 72393 | MGI:1919643 | 0.101 | 1 | n | n | y | protein coding gene |
| Ap4m1 | adaptor-related protein complex AP-4, mu | 11781 | MGI:1337063 | 0.146 | 1 | n | n | y | protein coding gene |
| Actr3b | ARP3 actin-related protein 3B | 242894 | MGI:2661120 | 0.089 | 1 | n | n | y | protein coding gene |
| Stxbp2 | syntaxin binding protein 2 | 20911 | MGI:107370 | 0.059 | 1 | n | n | y | protein coding gene |
| Ankrd13b | ankyrin repeat domain 13b | 268445 | MGI:2144501 | -0.216 | 1 | n | n | y | protein coding gene |
| Skor1 | SKI family transcriptional corepressor 1 | 207667 | MGI:2443473 | 0.213 | 1 | n | n | y | protein coding gene |
| Zswim8 | zinc finger SWIM-type containing 8 | 268721 | MGI:1919156 | 0.073 | 1 | n | n | y | protein coding gene |
| Vwa5b2 | von Willebrand factor A domain containing | 328643 | MGI:2681859 | 0.155 | 1 | n | n | y | protein coding gene |
| Acbd7 | acyl-Coenzyme A binding domain contain | 78245 | MGI:1925495 | 0.138 | 1 | n | n | y | protein coding gene |
| Mob3b | MOB kinase activator 3B | 214944 | MGI:2664539 | 0.123 | 1 | n | n | y | protein coding gene |
| D430019H16Rik | RIKEN cDNA D430019H16 gene | 268595 | MGI:2443127 | -0.121 | 1 | n | n | y | protein coding gene |
| Tekt2 | tektin 2 | 24084 | MGI:1346335 | 0.122 | 1 | n | n | y | protein coding gene |
| Armc9 | armadillo repeat containing 9 | 78795 | MGI:1926045 | 0.077 | 1 | n | n | y | protein coding gene |
| Ndufaf3 | NADH:ubiquinone oxidoreductase comple | 66706 | MGI:1913956 | -0.064 | 1 | n | n | y | protein coding gene |
| Scoc | short coiled-coil protein | 56367 | MGI:1927654 | 0.076 | 1 | n | n | y | protein coding gene |
| Lats1 | large tumor suppressor | 16798 | MGI:1333883 | 0.113 | 1 | n | n | y | protein coding gene |
| Palm3 | paralemmin 3 | 74337 | MGI:1921587 | -0.154 | 1 | n | n | y | protein coding gene |
| Spsb4 | splA/ryanodine receptor domain and SOC | 211949 | MGI:2183445 | -0.101 | 1 | n | n | y | protein coding gene |
| Lmln | leishmanolysin-like (metallopeptidase M8 | 239833 | MGI:2444736 | 0.106 | 1 | n | n | y | protein coding gene |
| Dlk2 | delta like non-canonical Notch ligand 2 | 106565 | MGI:2146838 | 0.104 | 1 | n | n | y | protein coding gene |
| Dhrs11 | dehydrogenase/reductase 11 | 192970 | MGI:2652816 | -0.108 | 1 | n | n | y | protein coding gene |

| | | | | | | | | | |
|---|---|---|---|---|---|---|---|---|---|
| Evl | Ena-vasodilator stimulated phosphoprotei | 14026 | MGI:1194884 | 0.060 | 1 | n | n | y | protein coding gene |
| Herc3 | hect domain and RLD 3 | 73998 | MGI:1921248 | -0.070 | 1 | n | n | y | protein coding gene |
| Herc1 | HECT and RLD domain containing E3 ubi | 235439 | MGI:2384589 | 0.109 | 1 | n | n | y | protein coding gene |
| Ubxn11 | UBX domain protein 11 | 67586 | MGI:1914836 | -0.056 | 1 | n | n | y | protein coding gene |
| Bltp3a | bridge-like lipid transfer protein family mer | 224648 | MGI:3041238 | 0.077 | 1 | n | n | y | protein coding gene |
| Cep85l | centrosomal protein 85-like | 100038725 | MGI:3642684 | 0.131 | 1 | n | n | y | protein coding gene |
| Homer2 | homer scaffolding protein 2 | 26557 | MGI:1347354 | 0.077 | 1 | n | n | y | protein coding gene |
| Pip4k2a | phosphatidylinositol-5-phosphate 4-kinase | 18718 | MGI:1298206 | -0.066 | 1 | n | n | y | protein coding gene |
| Atxn7l3 | ataxin 7-like 3 | 217218 | MGI:3036270 | -0.128 | 1 | n | n | y | protein coding gene |
| Nudcd3 | NudC domain containing 3 | 209586 | MGI:2144158 | -0.063 | 1 | n | n | y | protein coding gene |
| Barhl1 | BarH like homeobox 1 | 54422 | MGI:1859288 | 0.094 | 1 | n | n | y | protein coding gene |
| Sars2 | seryl-aminoacyl-tRNA synthetase 2 | 71984 | MGI:1919234 | -0.069 | 1 | n | n | y | protein coding gene |
| Sncg | synuclein, gamma | 20618 | MGI:1298397 | -0.119 | 1 | n | n | y | protein coding gene |
| Lrrc51 | leucine rich repeat containing 51 | 69358 | MGI:1916608 | 0.118 | 1 | n | n | y | protein coding gene |
| Rnf182 | ring finger protein 182 | 328234 | MGI:3045355 | 0.092 | 1 | n | n | y | protein coding gene |
| Polr3c | polymerase (RNA) III (DNA directed) polyp | 74414 | MGI:1921664 | -0.062 | 1 | n | n | y | protein coding gene |
| Csnk1g2 | casein kinase 1, gamma 2 | 103236 | MGI:1920014 | 0.176 | 1 | n | n | y | protein coding gene |
| Hsdl1 | hydroxysteroid dehydrogenase like 1 | 72552 | MGI:1919802 | -0.054 | 1 | n | n | y | protein coding gene |
| Braf | Braf transforming gene | 109880 | MGI:88190 | -0.143 | 1 | n | n | y | protein coding gene |
| Amph | amphiphysin | 218038 | MGI:103574 | 0.071 | 1 | n | n | y | protein coding gene |
| Dmxl2 | Dmx-like 2 | 235380 | MGI:2444630 | 0.116 | 1 | n | n | y | protein coding gene |
| Cwc22 | CWC22 spliceosome-associated protein | 80744 | MGI:2136773 | 0.091 | 1 | n | n | y | protein coding gene |
| Ubr2 | ubiquitin protein ligase E3 component n-re | 224826 | MGI:1861099 | 0.063 | 1 | n | n | y | protein coding gene |
| Pcsk5 | proprotein convertase subtilisin/kexin type | 18552 | MGI:97515 | -0.070 | 1 | n | n | y | protein coding gene |
| Ilf3 | interleukin enhancer binding factor 3 | 16201 | MGI:1339973 | 0.104 | 1 | n | n | y | protein coding gene |
| Slc6a11 | solute carrier family 6 (neurotransmitter tra | 243616 | MGI:95630 | 0.267 | 1 | n | n | y | protein coding gene |
| Rab3gap1 | RAB3 GTPase activating protein subunit 1 | 226407 | MGI:2445001 | -0.122 | 1 | n | n | y | protein coding gene |
| Zfp101 | zinc finger protein 101 | 22643 | MGI:107547 | 0.099 | 1 | n | n | y | protein coding gene |
| Ubn1 | ubinuclein 1 | 170644 | MGI:1891307 | -0.065 | 1 | n | n | y | protein coding gene |
| Nub1 | negative regulator of ubiquitin-like proteins | 53312 | MGI:1889001 | -0.049 | 1 | n | n | y | protein coding gene |
| Osbpl3 | oxysterol binding protein-like 3 | 71720 | MGI:1918970 | -0.058 | 1 | n | n | y | protein coding gene |
| Fam222a | family with sequence similarity 222, memb | 433940 | MGI:3605543 | -0.103 | 1 | n | n | y | protein coding gene |
| Hipk3 | homeodomain interacting protein kinase 3 | 15259 | MGI:1314882 | 0.078 | 1 | n | n | y | protein coding gene |
| Crybb3 | crystallin, beta B3 | 12962 | MGI:102717 | -0.155 | 1 | n | n | y | protein coding gene |
| Dnajc6 | DnaJ heat shock protein family (Hsp40) m | 72685 | MGI:1919935 | 0.089 | 1 | n | n | y | protein coding gene |
| Shroom2 | shroom family member 2 | 110380 | MGI:107194 | 0.072 | 1 | n | n | y | protein coding gene |
| Nup35 | nucleoporin 35 | 69482 | MGI:1916732 | 0.093 | 1 | n | n | y | protein coding gene |
| Gripap1 | GRIP1 associated protein 1 | 54645 | MGI:1859616 | 0.056 | 1 | n | n | y | protein coding gene |
| Rfng | RFNG O-fucosylpeptide 3-beta-N-acetylgl | 19719 | MGI:894275 | 0.053 | 1 | n | n | y | protein coding gene |
| Lrig2 | leucine-rich repeats and immunoglobulin-l | 269473 | MGI:2443718 | 0.053 | 1 | n | n | y | protein coding gene |
| Gigyf2 | GRB10 interacting GYF protein 2 | 227331 | MGI:2138584 | -0.070 | 1 | n | n | y | protein coding gene |
| Gtf2f2 | general transcription factor IIF, polypeptide | 68705 | MGI:1915955 | -0.056 | 1 | n | n | y | protein coding gene |
| Phf10 | PHD finger protein 10 | 72057 | MGI:1919307 | 0.061 | 1 | n | n | y | protein coding gene |
| Nudt14 | nudix hydrolase 14 | 66174 | MGI:1913424 | -0.048 | 1 | n | n | y | protein coding gene |
| Rrp36 | ribosomal RNA processing 36 | 224823 | MGI:2385053 | -0.057 | 1 | n | n | y | protein coding gene |
| Stxbp4 | syntaxin binding protein 4 | 20913 | MGI:1342296 | 0.088 | 1 | n | n | y | protein coding gene |
| Ttc7 | tetratricopeptide repeat domain 7 | 225049 | MGI:1920999 | -0.090 | 1 | n | n | y | protein coding gene |
| Defb25 | defensin beta 25 | 654459 | MGI:3651158 | -0.090 | 1 | n | n | y | protein coding gene |
| Sgsm2 | small G protein signaling modulator 2 | 97761 | MGI:2144695 | 0.047 | 1 | n | n | y | protein coding gene |
| Znhit1 | zinc finger, HIT domain containing 1 | 70103 | MGI:1917353 | -0.055 | 1 | n | n | y | protein coding gene |
| Prkab1 | protein kinase, AMP-activated, beta 1 non | 19079 | MGI:1336167 | -0.041 | 1 | n | n | y | protein coding gene |
| Katnb1 | katanin p80 (WD40-containing) subunit B | 74187 | MGI:1921437 | 0.040 | 1 | n | n | y | protein coding gene |
| Pi4ka | phosphatidylinositol 4-kinase alpha | 224020 | MGI:2448506 | 0.070 | 1 | n | n | y | protein coding gene |
| Pgrmc2 | progesterone receptor membrane compor | 70804 | MGI:1918054 | 0.069 | 1 | n | n | y | protein coding gene |
| Ppp1r16a | protein phosphatase 1, regulatory subunit | 73062 | MGI:1920312 | 0.042 | 1 | n | n | y | protein coding gene |
| Pxdc1 | PX domain containing 1 | 66895 | MGI:1914145 | 0.056 | 1 | n | n | y | protein coding gene |
| Dhps | deoxyhypusine synthase | 330817 | MGI:2683592 | -0.059 | 1 | n | n | y | protein coding gene |
| Ccdc74a | coiled-coil domain containing 74A | 72315 | MGI:1919565 | -0.073 | 1 | n | n | y | protein coding gene |
| Specc1l | sperm antigen with calponin homology and | 74392 | MGI:1921642 | 0.073 | 1 | n | n | y | protein coding gene |
| Kdm4b | lysine (K)-specific demethylase 4B | 193796 | MGI:2442355 | 0.039 | 1 | n | n | y | protein coding gene |
| Slc9a8 | solute carrier family 9 (sodium/hydrogen e | 77031 | MGI:1924281 | -0.060 | 1 | n | n | y | protein coding gene |
| Pdzrn3 | PDZ domain containing RING finger 3 | 55983 | MGI:1933157 | -0.049 | 1 | n | n | y | protein coding gene |
| Slc22a23 | solute carrier family 22, member 23 | 73102 | MGI:1920352 | -0.075 | 1 | n | n | y | protein coding gene |
| Ddx55 | DEAD box helicase 55 | 67848 | MGI:1915098 | -0.049 | 1 | n | n | y | protein coding gene |

| | | | | | | | | | |
|---|---|---|---|---|---|---|---|---|---|
| Fndc5 | fibronectin type III domain containing 5 | 384061 | MGI:1917614 | -0.075 | 1 | n | n | y | protein coding gene |
| Tmem120b | transmembrane protein 120B | 330189 | MGI:3603158 | 0.064 | 1 | n | n | y | protein coding gene |
| Cdk17 | cyclin dependent kinase 17 | 237459 | MGI:97517 | -0.061 | 1 | n | n | y | protein coding gene |
| Zdhhc7 | zinc finger, DHHC domain containing 7 | 102193 | MGI:2142662 | -0.049 | 1 | n | n | y | protein coding gene |
| Cln6 | ceroid-lipofuscinosis, neuronal 6 | 76524 | MGI:2159324 | -0.060 | 1 | n | n | y | protein coding gene |
| Cnnm2 | cyclin M2 | 94219 | MGI:2151054 | 0.076 | 1 | n | n | y | protein coding gene |
| Grxcr2 | glutaredoxin, cysteine rich 2 | 332309 | MGI:2685697 | -0.034 | 1 | n | n | y | protein coding gene |
| Tanc1 | tetratricopeptide repeat, ankyrin repeat an | 66860 | MGI:1914110 | 0.065 | 1 | n | n | y | protein coding gene |
| Cnih2 | cornichon family AMPA receptor auxiliary | 12794 | MGI:1277225 | 0.054 | 1 | n | n | y | protein coding gene |
| Ep400 | E1A binding protein p400 | 75560 | MGI:1276124 | -0.048 | 1 | n | n | y | protein coding gene |
| Acap2 | ArfGAP with coiled-coil, ankyrin repeat an | 78618 | MGI:1925868 | -0.055 | 1 | n | n | y | protein coding gene |
| D5Ertd579e | DNA segment, Chr 5, ERATO Doi 579, ex | 320661 | MGI:1261849 | 0.061 | 1 | n | n | y | protein coding gene |
| Samd14 | sterile alpha motif domain containing 14 | 217125 | MGI:2384945 | -0.046 | 1 | n | n | y | protein coding gene |
| Tmem138 | transmembrane protein 138 | 72982 | MGI:1920232 | -0.042 | 1 | n | n | y | protein coding gene |
| Nfic | nuclear factor I/C | 18029 | MGI:109591 | 0.053 | 1 | n | n | y | protein coding gene |
| Gls | glutaminase | 14660 | MGI:95752 | 0.074 | 1 | n | n | y | protein coding gene |
| Ttc24 | tetratricopeptide repeat domain 24 | 214191 | MGI:2443841 | 0.057 | 1 | n | n | y | protein coding gene |
| Otud4 | OTU domain containing 4 | 73945 | MGI:1098801 | -0.062 | 1 | n | n | y | protein coding gene |
| Meig1 | meiosis expressed gene 1 | 104362 | MGI:1202878 | -0.096 | 1 | n | n | y | protein coding gene |
| Apba1 | amyloid beta precursor protein binding far | 319924 | MGI:1860297 | -0.050 | 1 | n | n | y | protein coding gene |
| Mmp24 | matrix metallopeptidase 24 | 17391 | MGI:1341867 | -0.080 | 1 | n | n | y | protein coding gene |
| Ccl27a | C-C motif chemokine ligand 27A | 20301 | MGI:1343459 | -0.043 | 1 | n | n | y | protein coding gene |
| Pi4k2a | phosphatidylinositol 4-kinase type 2 alpha | 84095 | MGI:1934031 | -0.048 | 1 | n | n | y | protein coding gene |
| Ablim2 | actin-binding LIM protein 2 | 231148 | MGI:2385758 | 0.053 | 1 | n | n | y | protein coding gene |
| Dpcd | deleted in primary ciliary dyskinesia | 226162 | MGI:1924407 | 0.029 | 1 | n | n | y | protein coding gene |
| Ncoa6 | nuclear receptor coactivator 6 | 56406 | MGI:1929915 | 0.097 | 1 | n | n | y | protein coding gene |
| Kcns3 | potassium voltage-gated channel, delayec | 238076 | MGI:1098804 | -0.041 | 1 | n | n | y | protein coding gene |
| Apip | APAF1 interacting protein | 56369 | MGI:1926788 | 0.047 | 1 | n | n | y | protein coding gene |
| Hic2 | hypermethylated in cancer 2 | 58180 | MGI:1929869 | -0.068 | 1 | n | n | y | protein coding gene |
| Pygo1 | pygopus 1 | 72135 | MGI:1919385 | 0.059 | 1 | n | n | y | protein coding gene |
| Rbm26 | RNA binding motif protein 26 | 74213 | MGI:1921463 | 0.062 | 1 | n | n | y | protein coding gene |
| Chrna10 | cholinergic receptor, nicotinic, alpha polyp | 504186 | MGI:3609260 | 0.031 | 1 | n | n | y | protein coding gene |
| Dcun1d2 | defective in cullin neddylation 1 domain cc | 102323 | MGI:2142792 | -0.040 | 1 | n | n | y | protein coding gene |
| Spef1 | sperm flagellar 1 | 70997 | MGI:3513546 | 0.060 | 1 | n | n | y | protein coding gene |
| Rara | retinoic acid receptor, alpha | 19401 | MGI:97856 | 0.036 | 1 | n | n | y | protein coding gene |
| Srrm3 | serine/arginine repetitive matrix 3 | 58212 | MGI:1920309 | -0.086 | 1 | n | n | y | protein coding gene |
| Ubn2 | ubinuclein 2 | 320538 | MGI:2444236 | -0.068 | 1 | n | n | y | protein coding gene |
| Cnot6l | CCR4-NOT transcription complex, subunit | 231464 | MGI:2443154 | -0.055 | 1 | n | n | y | protein coding gene |
| Atxn7 | ataxin 7 | 246103 | MGI:2179277 | 0.056 | 1 | n | n | y | protein coding gene |
| Nherf2 | NHERF family PDZ scaffold protein 2 | 65962 | MGI:1890662 | -0.032 | 1 | n | n | y | protein coding gene |
| Orai2 | ORAI calcium release-activated calcium m | 269717 | MGI:2443195 | 0.032 | 1 | n | n | y | protein coding gene |
| Hira | histone cell cycle regulator | 15260 | MGI:99430 | 0.045 | 1 | n | n | y | protein coding gene |
| Fbxw17 | F-box and WD-40 domain protein 17 | 109082 | MGI:1923584 | 0.033 | 1 | n | n | y | protein coding gene |
| Slc35e4 | solute carrier family 35, member E4 | 103710 | MGI:2144150 | -0.046 | 1 | n | n | y | protein coding gene |
| Nomo1 | nodal modulator 1 | 211548 | MGI:2385850 | 0.029 | 1 | n | n | y | protein coding gene |
| Arl6 | ADP-ribosylation factor-like 6 | 56297 | MGI:1927136 | 0.039 | 1 | n | n | y | protein coding gene |
| Slc38a1 | solute carrier family 38, member 1 | 105727 | MGI:2145895 | 0.033 | 1 | n | n | y | protein coding gene |
| Hacd3 | 3-hydroxyacyl-CoA dehydratase 3 | 57874 | MGI:1889341 | 0.027 | 1 | n | n | y | protein coding gene |
| Coq4 | coenzyme Q4 | 227683 | MGI:1098826 | -0.033 | 1 | n | n | y | protein coding gene |
| Fryl | FRY like transcription coactivator | 72313 | MGI:1919563 | 0.064 | 1 | n | n | y | protein coding gene |
| Tapt1 | transmembrane anterior posterior transfor | 231225 | MGI:2683537 | -0.029 | 1 | n | n | y | protein coding gene |
| Rufy3 | RUN and FYVE domain containing 3 | 52822 | MGI:106484 | -0.039 | 1 | n | n | y | protein coding gene |
| Ppil6 | peptidylprolyl isomerase (cyclophilin)-like ( | 73075 | MGI:1920325 | 0.043 | 1 | n | n | y | protein coding gene |
| Fam120a | family with sequence similarity 120, memb | 218236 | MGI:2446163 | -0.039 | 1 | n | n | y | protein coding gene |
| Atxn1 | ataxin 1 | 20238 | MGI:104783 | -0.039 | 1 | n | n | y | protein coding gene |
| Irx2 | Iroquois homeobox 2 | 16372 | MGI:1197526 | -0.099 | 1 | n | n | y | protein coding gene |
| Ptpn13 | protein tyrosine phosphatase, non-recepto | 19249 | MGI:103293 | 0.050 | 1 | n | n | y | protein coding gene |
| Smim29 | small integral membrane protein 29 | 106672 | MGI:2146839 | 0.044 | 1 | n | n | y | protein coding gene |
| Cnot8 | CCR4-NOT transcription complex, subunit | 69125 | MGI:1916375 | -0.019 | 1 | n | n | y | protein coding gene |
| Wasf1 | WASP family, member 1 | 83767 | MGI:1890563 | -0.027 | 1 | n | n | y | protein coding gene |
| Clasp1 | CLIP associating protein 1 | 76707 | MGI:1923957 | 0.092 | 1 | n | n | y | protein coding gene |
| Cplx2 | complexin 2 | 12890 | MGI:104726 | -0.053 | 1 | n | n | y | protein coding gene |
| Coq9 | coenzyme Q9 | 67914 | MGI:1915164 | 0.031 | 1 | n | n | y | protein coding gene |
| Gns | glucosamine (N-acetyl)-6-sulfatase | 75612 | MGI:1922862 | -0.019 | 1 | n | n | y | protein coding gene |

| | | | | | | | | | |
|---|---|---|---|---|---|---|---|---|---|
| Myo16 | myosin XVI | 244281 | MGI:2685951 | -0.076 | 1 | n | n | y | protein coding gene |
| Taf15 | TATA-box binding protein associated facto | 70439 | MGI:1917689 | -0.038 | 1 | n | n | y | protein coding gene |
| Ankra2 | ankyrin repeat family A member 2 | 68558 | MGI:1915808 | 0.023 | 1 | n | n | y | protein coding gene |
| Ttc39b | tetratricopeptide repeat domain 39B | 69863 | MGI:1917113 | 0.033 | 1 | n | n | y | protein coding gene |
| Uckl1 | uridine-cytidine kinase 1-like 1 | 68556 | MGI:1915806 | 0.027 | 1 | n | n | y | protein coding gene |
| Anxa4 | annexin A4 | 11746 | MGI:88030 | -0.019 | 1 | n | n | y | protein coding gene |
| Arhgap26 | Rho GTPase activating protein 26 | 71302 | MGI:1918552 | 0.058 | 1 | n | n | y | protein coding gene |
| Tex9 | testis expressed gene 9 | 21778 | MGI:1201610 | -0.032 | 1 | n | n | y | protein coding gene |
| Wwc2 | WW, C2 and coiled-coil domain containing | 52357 | MGI:1261872 | 0.023 | 1 | n | n | y | protein coding gene |
| Khdrbs3 | KH domain containing, RNA binding, signa | 13992 | MGI:1313312 | -0.070 | 1 | n | n | y | protein coding gene |
| Ylpm1 | YLP motif containing 1 | 56531 | MGI:1926195 | -0.027 | 1 | n | n | y | protein coding gene |
| Pdcd7 | programmed cell death 7 | 50996 | MGI:1859170 | 0.026 | 1 | n | n | y | protein coding gene |
| Ak8 | adenylate kinase 8 | 68870 | MGI:1916120 | 0.035 | 1 | n | n | y | protein coding gene |
| Cog7 | component of oligomeric golgi complex 7 | 233824 | MGI:2685013 | -0.018 | 1 | n | n | y | protein coding gene |
| Jag2 | jagged 2 | 16450 | MGI:1098270 | 0.042 | 1 | n | n | y | protein coding gene |
| Carf | calcium response factor | 241066 | MGI:2182269 | 0.035 | 1 | n | n | y | protein coding gene |
| Dnajc15 | DnaJ heat shock protein family (Hsp40) m | 66148 | MGI:1913398 | -0.021 | 1 | n | n | y | protein coding gene |
| Tpm1 | tropomyosin 1, alpha | 22003 | MGI:98809 | 0.027 | 1 | n | n | y | protein coding gene |
| Asxl2 | ASXL transcriptional regulator 2 | 75302 | MGI:1922552 | 0.040 | 1 | n | n | y | protein coding gene |
| Mapk6 | mitogen-activated protein kinase 6 | 50772 | MGI:1354946 | -0.018 | 1 | n | n | y | protein coding gene |
| Chga | chromogranin A | 12652 | MGI:88394 | 0.022 | 1 | n | n | y | protein coding gene |
| Mycbp2 | MYC binding protein 2, E3 ubiquitin protei | 105689 | MGI:2179432 | 0.021 | 1 | n | n | y | protein coding gene |
| Epha4 | Eph receptor A4 | 13838 | MGI:98277 | 0.020 | 1 | n | n | y | protein coding gene |
| Pcgf1 | polycomb group ring finger 1 | 69837 | MGI:1917087 | 0.028 | 1 | n | n | y | protein coding gene |
| Synrg | synergin, gamma | 217030 | MGI:1354742 | -0.029 | 1 | n | n | y | protein coding gene |
| Gars1 | glycyl-tRNA synthetase 1 | 353172 | MGI:2449057 | -0.014 | 1 | n | n | y | protein coding gene |
| Haghl | hydroxyacylglutathione hydrolase-like | 68977 | MGI:1919877 | 0.019 | 1 | n | n | y | protein coding gene |
| Lpin2 | lipin 2 | 64898 | MGI:1891341 | 0.017 | 1 | n | n | y | protein coding gene |
| Mapk8ip3 | mitogen-activated protein kinase 8 interac | 30957 | MGI:1353598 | -0.016 | 1 | n | n | y | protein coding gene |
| Dcaf5 | DDB1 and CUL4 associated factor 5 | 320808 | MGI:2444785 | 0.013 | 1 | n | n | y | protein coding gene |
| Ulk2 | unc-51 like kinase 2 | 29869 | MGI:1352758 | 0.018 | 1 | n | n | y | protein coding gene |
| Brsk1 | BR serine/threonine kinase 1 | 381979 | MGI:2685946 | -0.033 | 1 | n | n | y | protein coding gene |
| Ccp110 | centriolar coiled coil protein 110 | 101565 | MGI:2141942 | -0.028 | 1 | n | n | y | protein coding gene |
| Pop5 | processing of precursor 5, ribonuclease P | 117109 | MGI:2151221 | -0.019 | 1 | n | n | y | protein coding gene |
| Ctnnd2 | catenin delta 2 | 18163 | MGI:1195966 | -0.019 | 1 | n | n | y | protein coding gene |
| Camk2g | calcium/calmodulin-dependent protein kina | 12325 | MGI:88259 | -0.013 | 1 | n | n | y | protein coding gene |
| Bcl7a | B cell CLL/lymphoma 7A | 77045 | MGI:1924295 | -0.024 | 1 | n | n | y | protein coding gene |
| Bola1 | bolA family member 1 | 69168 | MGI:1916418 | -0.019 | 1 | n | n | y | protein coding gene |
| Hip1r | huntingtin interacting protein 1 related | 29816 | MGI:1352504 | 0.017 | 1 | n | n | y | protein coding gene |
| Plekho1 | pleckstrin homology domain containing, fa | 67220 | MGI:1914470 | -0.023 | 1 | n | n | y | protein coding gene |
| Trappc9 | trafficking protein particle complex 9 | 76510 | MGI:1923760 | 0.020 | 1 | n | n | y | protein coding gene |
| Plce1 | phospholipase C, epsilon 1 | 74055 | MGI:1921305 | -0.021 | 1 | n | n | y | protein coding gene |
| Tnrc6a | trinucleotide repeat containing 6a | 233833 | MGI:2385292 | 0.024 | 1 | n | n | y | protein coding gene |
| Mtmr3 | myotubularin related protein 3 | 74302 | MGI:1921552 | 0.016 | 1 | n | n | y | protein coding gene |
| Magi3 | membrane associated guanylate kinase, W | 99470 | MGI:1923484 | 0.020 | 1 | n | n | y | protein coding gene |
| Brf2 | BRF2, RNA polymerase III transcription in | 66653 | MGI:1913903 | 0.014 | 1 | n | n | y | protein coding gene |
| Crmp1 | collapsin response mediator protein 1 | 12933 | MGI:107793 | 0.016 | 1 | n | n | y | protein coding gene |
| Cib2 | calcium and integrin binding family membe | 56506 | MGI:1929293 | -0.009 | 1 | n | n | y | protein coding gene |
| Tnc | tenascin C | 21923 | MGI:101922 | -0.026 | 1 | n | n | y | protein coding gene |
| Fank1 | fibronectin type 3 and ankyrin repeat dom | 66930 | MGI:1914180 | -0.020 | 1 | n | n | y | protein coding gene |
| Ogfod2 | 2-oxoglutarate and iron-dependent oxyger | 66627 | MGI:1913877 | 0.010 | 1 | n | n | y | protein coding gene |
| Cetn2 | centrin 2 | 26370 | MGI:1347085 | 0.008 | 1 | n | n | y | protein coding gene |
| Tmc1 | transmembrane channel-like gene family 1 | 13409 | MGI:2151016 | 0.012 | 1 | n | n | y | protein coding gene |
| Pip5k1c | phosphatidylinositol-4-phosphate 5-kinase | 18717 | MGI:1298224 | 0.012 | 1 | n | n | y | protein coding gene |
| Ik | IK cytokine | 24010 | MGI:1345142 | 0.007 | 1 | n | n | y | protein coding gene |
| Bbip1 | BBSome interacting protein 1 | 100503572 | MGI:1913610 | -0.008 | 1 | n | n | y | protein coding gene |
| Hmg20a | high mobility group 20A | 66867 | MGI:1914117 | 0.007 | 1 | n | n | y | protein coding gene |
| Brpf1 | bromodomain and PHD finger containing, | 78783 | MGI:1926033 | -0.009 | 1 | n | n | y | protein coding gene |
| Sptb | spectrin beta, erythrocytic | 20741 | MGI:98387 | 0.010 | 1 | n | n | y | protein coding gene |
| Ift74 | intraflagellar transport 74 | 67694 | MGI:1914944 | -0.005 | 1 | n | n | y | protein coding gene |
| 2510002D24Rik | RIKEN cDNA 2510002D24 gene | 72307 | MGI:1919557 | 0.006 | 1 | n | n | y | protein coding gene |
| Znhit2 | zinc finger, HIT domain containing 2 | 29805 | MGI:1352481 | 0.008 | 1 | n | n | y | protein coding gene |
| Dnal4 | dynein, axonemal, light chain 4 | 54152 | MGI:1859217 | -0.004 | 1 | n | n | y | protein coding gene |
| B230219D22Rik | RIKEN cDNA B230219D22 gene | 78521 | MGI:1925771 | -0.004 | 1 | n | n | y | protein coding gene |

| | | | | | | | | | |
|---|---|---|---|---|---|---|---|---|---|
| Ppfia4 | protein tyrosine phosphatase, receptor typ | 68507 | MGI:1915757 | -0.008 | 1 | n | n | y | protein coding gene |
| Gng8 | guanine nucleotide binding protein (G prot | 14709 | MGI:109163 | 0.013 | 1 | n | n | y | protein coding gene |
| Ubac1 | ubiquitin associated domain containing 1 | 98766 | MGI:1920995 | 0.004 | 1 | n | n | y | protein coding gene |
| Rnf150 | ring finger protein 150 | 330812 | MGI:2443860 | 0.007 | 1 | n | n | y | protein coding gene |
| Frmd6 | FERM domain containing 6 | 319710 | MGI:2442579 | 0.004 | 1 | n | n | y | protein coding gene |
| Ift122 | intraflagellar transport 122 | 81896 | MGI:1932386 | 0.003 | 1 | n | n | y | protein coding gene |
| Hrk | harakiri, BCL2 interacting protein (contains | 12123 | MGI:1201608 | 0.007 | 1 | n | n | y | protein coding gene |
| Purb | purine rich element binding protein B | 19291 | MGI:1338779 | -0.003 | 1 | n | n | y | protein coding gene |
| Hpf1 | histone PARylation factor 1 | 72612 | MGI:1919862 | 0.001 | 1 | n | n | y | protein coding gene |
| Slc8a2 | solute carrier family 8 (sodium/calcium exc | 110891 | MGI:107996 | -0.002 | 1 | n | n | y | protein coding gene |
| Cfap144 | cilia and flagella associated protein 144 | 75429 | MGI:1922679 | ND* | ND* | n | n | y | protein coding gene |
| Pam16l | presequence translocase associated moto | 100042179 | MGI:3704359 | ND* | ND* | n | n | y | protein coding gene |
| Gm32742 | predicted gene, 32742 | 102635385 | MGI:5591901 | ND* | ND* | n | n | y | protein coding gene |
| Gm30191 | predicted gene, 30191 | 115489946 | MGI:5589350 | ND* | ND* | n | n | y | protein coding gene |
| Sco2 | SCO2 cytochrome c oxidase assembly pro | 100126824 | MGI:3818630 | ND* | ND* | n | n | y | protein coding gene |
| Ccer2 | coiled-coil glutamate-rich protein 2 | 100504112 | MGI:3645242 | ND* | ND* | n | n | y | protein coding gene |
| Ccl21a | C-C motif chemokine ligand 21 (serine) | 18829 | MGI:1349183 | ND* | ND* | n | n | y | protein coding gene |
| Tcf15 | transcription factor 15 | 21407 | MGI:104664 | ND* | ND* | n | n | y | protein coding gene |
