## Supplementary material for "CASZ1 regulates the rate at which outer hair cells mature and is required for hearing": Table S4

**Table S4. Genes annotated with 'sensory perception of sound' GO term and differentially expressed in organs of Corti of *Casz1*$^{fl/fl}$*;Tg(Sox10-Cre)* versus WT mice.**

| Gene symbol | Gene name | Entrez gene ID | MGI gene ID | Annotated with 'sensory perception of sound' GO term? y/n | logFC *Casz1(fl/fl);Tg(Sox10-Cre)* vs WT | More than 2-fold difference in expression? y/n | FDR-adjusted P (Benjamini-Yekutieli method) | Previously reported association between gene defect and hearing loss or HC loss. Listed for genes with 2-fold difference in expression between *Casz1(fl/fl);Tg(Sox10-Cre)* vs WT. |
|---|---|---|---|---|---|---|---|---|
| Ocm | oncomodulin | 18261 | MGI:97401 | y* (PMID: 26843644) | 3.006 | y | 2.61E-10 | *Ocm* overexpressor mouse line has not been produced or characterized for hearing loss or HC loss. |
| Hgf | hepatocyte growth factor | 15234 | MGI:96079 | y* (PMID: 19576567, 32152201) | 2.504 | y | 1.97E-07 | Transgenic overexpression of Hgf (MH19-Hgf) causes progressive OHC loss and hearing loss, but it does not cause disorganization of stereocilium bundles (PMID: 19576567). |
| Dcdc2a | doublecortin domain containing 2a | 195208 | MGI:2652818 | y | 2.436 | y | 9.36E-08 | Overexpression of wild-type DCDC2a-GFP in HCs causes elongation of kinocilia (PMID: 25601850). |
| Cdkn2d | cyclin dependent kinase inhibitor 2D | 12581 | MGI:105387 | y | 1.018 | y | 1.93E-09 | *Cdkn2d* overexpressor mouse line has not been produced or characterized for hearing loss or HC loss. |
| Jag1 | jagged 1 | 16449 | MGI:1095416 | y* (PMID: 11259677) | 1.013 | y | 0.000131999 | JAG1-peptide treatment boosts HC formation in cochlear explants (doi: https://doi.org/10.1101/2025.03.02.640998). |
| Fscn2 | fascin actin-bundling protein 2 | 238021 | MGI:2443337 | y* (PMID: 20660251) | 1.004 | y | 1.90E-08 | Overexpression of EGFP-FSCN2 in HCs does not impair auditory function in mice (PMID: 29874122). |
| Pkhd1l1 | polycystic kidney and hepatic disea | 192190 | MGI:2183153 | y | -1.701 | y | 6.50E-09 | Inactivation of *Pkhd1l1* in HCs causes progressive loss of OHC stereocilia starting ~6 weeks after birth, but it does not cause loss of OHCs or disorganization of stereocilium bundles at rootlet level (doi: 10.1101/2024.02.29.582786). |
| Tub | tubby bipartite transcription factor | 22141 | MGI:2651573 | y | -1.708 | y | 2.16E-07 | Inactivation of *Tub* causes loss of cohesion in stereocilium bundles in OHCs, but it does not cause OHC loss or disorganization of stereocilium bundles at rootlet level (PMID: 32358189). |
| Myo3a | myosin IIIA | 667663 | MGI:2183924 | y | 0.987 | n | 1.17E-06 | |
| Myo15a | myosin XVA | 17910 | MGI:1261811 | y | 0.840 | n | 4.40E-06 | |
| Ptprq | protein tyrosine phosphatase recep | 237523 | MGI:1096349 | y* (PMID: 14534255) | 0.792 | n | 0.000803483 | |
| Gfi1 | growth factor independent 1 transcı | 14581 | MGI:103170 | y* (PMID: 12441305) | 0.710 | n | 3.78E-07 | |
| Aqp4 | aquaporin 4 | 11829 | MGI:107387 | y | 0.703 | n | 1.59E-05 | |
| Adgrv1 | adhesion G protein-coupled recepto | 110789 | MGI:1274784 | y | 0.681 | n | 0.000110178 | |
| Atoh1 | atonal bHLH transcription factor 1 | 11921 | MGI:104654 | y* (PMID: 10364557) | 0.638 | n | 0.000889297 | |
| Pjvk | pejvakin | 381375 | MGI:2685847 | y | 0.608 | n | 0.000785437 | |
| Tmtc4 | transmembrane and tetratricopeptic | 70551 | MGI:1921050 | y | 0.597 | n | 2.90E-06 | |
| Xirp2 | xin actin-binding repeat containing : | 241431 | MGI:2685198 | y* (PMID: 25772365, 25653358) | 0.562 | n | 4.99E-05 | |
| Mcoln3 | mucolipin 3 | 171166 | MGI:1890500 | y | 0.506 | n | 2.75E-05 | |
| Ckmt1 | creatine kinase, mitochondrial 1, ub | 12716 | MGI:99441 | y* (PMID: 14977190) | 0.483 | n | 0.000102818 | |
| Clic5 | chloride intracellular channel 5 | 224796 | MGI:1917912 | y | 0.473 | n | 0.000419881 | |
| Ercc6 | excision repair cross-complementin | 319955 | MGI:1100494 | y* (PMID: 25762674) | 0.450 | n | 0.000971013 | |
| Slitrk6 | SLIT and NTRK-like family, membe | 239250 | MGI:2443198 | y | 0.447 | n | 0.000191487 | |
| Tomt | transmembrane O-methyltransferas | 791260 | MGI:3769724 | y | 0.435 | n | 0.000670323 | |
| Smpx | small muscle protein, X-linked | 66106 | MGI:1913356 | y* (PMID: 34722533) | 0.380 | n | 0.000207342 | |
| Eya1 | EYA transcriptional coactivator and | 14048 | MGI:109344 | y* (PMID: 18678597) | 0.306 | n | 0.00082903 | |
| Lhfpl5 | lipoma HMGIC fusion partner-like 5 | 328789 | MGI:1915382 | y | -0.343 | n | 0.000805895 | |
| Pou3f4 | POU domain, class 3, transcription | 18994 | MGI:101894 | y | -0.511 | n | 0.000240401 | |
| Grxcr1 | glutaredoxin, cysteine rich 1 | 433899 | MGI:3577767 | y | -0.521 | n | 0.000213385 | |
| Atp8a2 | ATPase, aminophospholipid transp | 50769 | MGI:1354710 | y* (PMID: 24413176) | -0.592 | n | 0.000673666 | |
| Axin1 | axin 1 | 12005 | MGI:1096327 | y | -0.656 | n | 0.000368589 | |
| Synj2 | synaptojanin 2 | 20975 | MGI:1201671 | y* (PMID: 21423608, 33100973) | -0.725 | n | 1.45E-05 | |
| Tbx18 | T-box18 | 76365 | MGI:1923615 | y | -0.738 | n | 4.31E-05 | |
| Cdkn1b | cyclin dependent kinase inhibitor 1E | 12576 | MGI:104565 | y | -0.896 | n | 1.48E-06 | |
| Otof | otoferlin | 83762 | MGI:1891247 | y | -0.938 | n | 6.18E-06 | |

*Manually updated based on indicated PMID.
