## Supplementary material for "CASZ1 regulates the rate at which outer hair cells mature and is required for hearing": Table S5

**Table S5. Genes annotated with actin-related and myosin-related GO terms and differentially expressed in organs of Corti of *Casz1fl/fl;Tg(Sox10-Cre)* versus WT mice.**

| Gene symbol | Gene name | Entrez gene ID | MGI gene ID | Annotated with 'actin binding', 'actin filament binding', 'myosin complex', or 'myosin II complex' GO terms? y/n | logFC *Casz1(fl/fl);Tg(Sox10-Cre)* vs WT | More than 2-fold difference in expression? y/n | FDR-adjusted P (Benjamini-Yekutieli method) | Previously reported association between gene defect and hearing loss or HC loss. Listed for genes with 2-fold difference in expression between *Casz1(fl/fl);Tg(Sox10-Cre)* vs WT. |
|---|---|---|---|---|---|---|---|---|
| Myl1 | myosin, light polypeptide 1 | 17901 | MGI:97269 | y | 6.895 | y | 3.59E-08 | *Myl1* overexpressor mouse line has not been produced or characterized for hearing or HC loss. |
| Myh13 | myosin, heavy polypeptide 13, s | 544791 | MGI:1339967 | y | 2.266 | y | 3.70E-07 | *Myh13* overexpressor mouse line has not been produced or characterized for hearing or HC loss. |
| Nod2 | nucleotide-binding oligomerizati | 257632 | MGI:2429397 | y | 2.209 | y | 1.82E-07 | Abnormally high NOD2 activity is associated with recurrent uveitis, dermatitis and arthritis but not hearing loss in humans (PMID: 22884558). |
| Cap1 | cyclase associated actin cytosk | 12331 | MGI:88262 | y | 1.784 | y | 1.77E-12 | *Cap1* overexpressor mouse line has not been produced or characterized for hearing or HC loss. |
| Coro2a | coronin, actin binding protein 2A | 107684 | MGI:1345966 | y | 1.655 | y | 8.73E-08 | *Coro2a* overexpressor mouse line has not been produced or characterized for hearing or HC loss. |
| Car7 | carbonic anhydrase 7 | 12354 | MGI:103100 | y* (PMID: 33719157) | 1.484 | y | 1.29E-08 | *Car7* overexpressor mouse line has not been produced or characterized for hearing or HC loss. |
| Fscn2 | fascin actin-bundling protein 2 | 238021 | MGI:2443337 | y | 1.004 | y | 1.90E-08 | Overexpression of EGFP-FSCN2 in HCs does not impair auditory function in mice (PMID: 29874122). |
| Nf2 | neurofibromin 2 | 18016 | MGI:97307 | y | -1.125 | y | 3.98E-09 | Cranial nerve VIII tumors cause hearing loss in *Nf2* cKO mice (PMID: 25113746). |
| Hpca | hippocalcin | 15444 | MGI:1336200 | y | -2.429 | y | 2.04E-07 | Normal hearing in *Hpca* KO mice (PMID: 36305825). |
| Myo3a | myosin IIIA | 667663 | MGI:2183924 | y | 0.987 | n | 1.17E-06 | |
| Myo1h | myosin 1H | 231646 | MGI:1914674 | y | 0.905 | n | 6.21E-06 | |
| Dmtn | dematin actin binding protein | 13829 | MGI:99670 | y | 0.897 | n | 1.57E-07 | |
| Myo15a | myosin XVA | 17910 | MGI:1261811 | y | 0.840 | n | 4.40E-06 | |
| Lmod1 | leiomodin 1 (smooth muscle) | 93689 | MGI:2135671 | y | 0.769 | n | 9.07E-05 | |
| Adcy8 | adenylate cyclase 8 | 11514 | MGI:1341110 | y | 0.731 | n | 0.000557841 | |
| Nebl | nebulette | 74103 | MGI:1921353 | y | 0.667 | n | 1.23E-05 | |
| Ablim1 | actin-binding LIM protein 1 | 226251 | MGI:1194500 | y | 0.626 | n | 1.77E-05 | |
| Myo18a | myosin XVIIIA | 360013 | MGI:2667185 | y | 0.585 | n | 6.21E-06 | |
| Xirp2 | xin actin-binding repeat containi | 241431 | MGI:2685198 | y | 0.562 | n | 4.99E-05 | |
| Twf2 | twinfilin actin binding protein 2 | 23999 | MGI:1346078 | y | 0.556 | n | 5.34E-06 | |
| Cap2 | cyclase associated actin cytosk | 67252 | MGI:1914502 | y | 0.515 | n | 7.02E-05 | |
| Abitram | actin binding transcription modu | 230234 | MGI:2677850 | y | 0.508 | n | 0.000212673 | |
| Map2 | microtubule-associated protein : | 17756 | MGI:97175 | y | 0.505 | n | 0.000226324 | |
| Vcl | vinculin | 22330 | MGI:98927 | y | 0.481 | n | 6.10E-05 | |
| Ncald | neurocalcin delta | 52589 | MGI:1196326 | y | 0.452 | n | 0.000277737 | |
| Pick1 | protein interacting with C kinase | 18693 | MGI:894645 | y | 0.448 | n | 7.75E-06 | |
| Misp | mitotic spindle positioning | 78906 | MGI:1926156 | y | 0.386 | n | 0.000515786 | |
| Lima1 | LIM domain and actin binding 1 | 65970 | MGI:1920992 | y | 0.356 | n | 0.000601212 | |
| Dbn1 | drebrin 1 | 56320 | MGI:1931838 | y | 0.336 | n | 0.000459513 | |
| Spire2 | spire type actin nucleation facto | 234857 | MGI:2446256 | y | 0.308 | n | 0.000752339 | |
| Trpv4 | transient receptor potential catic | 63873 | MGI:1926945 | y | 0.279 | n | 0.000955152 | |
| Calml4 | calmodulin-like 4 | 75600 | MGI:1922850 | y | -0.313 | n | 0.000343608 | |
| Dstn | destrin | 56431 | MGI:1929270 | y | -0.327 | n | 0.000600601 | |
| Tpm4 | tropomyosin 4 | 326618 | MGI:2449202 | y | -0.352 | n | 0.000314352 | |
| Myl9 | myosin, light polypeptide 9, regu | 98932 | MGI:2138915 | y | -0.362 | n | 3.37E-05 | |
| Arpc1b | actin related protein 2/3 comple: | 11867 | MGI:1343142 | y | -0.427 | n | 0.000228817 | |
| Ace | angiotensin I converting enzyme | 11421 | MGI:87874 | y | -0.430 | n | 0.000719821 | |

| | | | | | | | |
|---|---|---|---|---|---|---|---|
| Ajuba | ajuba LIM protein | 16475 | MGI:1341886 | y | -0.609 | n | 0.000162868 |
| Smtn | smoothelin | 29856 | MGI:1354727 | y | -0.622 | n | 0.000144272 |
| Wipf3 | WAS/WASL interacting protein f | 330319 | MGI:3044681 | y | -0.673 | n | 0.000840784 |
| Fhod3 | formin homology 2 domain conta | 225288 | MGI:1925847 | y | -0.827 | n | 2.24E-05 |
| Tpm2 | tropomyosin 2, beta | 22004 | MGI:98810 | y | -0.829 | n | 7.57E-06 |
| Sgip1 | SH3-domain GRB2-like (endoph | 73094 | MGI:1920344 | y | -0.914 | n | 4.40E-06 |
| Gas2l3 | growth arrest-specific 2 like 3 | 237436 | MGI:1918780 | y | -0.959 | n | 1.78E-07 |

*Manually updated based on indicated PMID.
