## Supplementary material for "CASZ1 regulates the rate at which outer hair cells mature and is required for hearing": Table S6

**Table S6. Primers and synthetic gene fragments used in this study.**

| Gene symbol or shRNA name | Forward primer (5' to 3') | Reverse primer (5' to 3') | DNA sequence for shRNA production or testing | Assay type | shRNA target sequence | Comments |
|---|---|---|---|---|---|---|
| Arhgap22 | GTCCAGAACTTGGCAACTGTC | TGAACCAGAGACGTGCCTCC | N/A | qRT-PCR | N/A | |
| Atp2a3 | TGAGATCACTGCCATGACTGG | TGGAGGCAAAGTTGTCATCGG | N/A | qRT-PCR | N/A | |
| Cacng2 | CTGCTGCCTCGAAGGGAACT | GATCACACTCAGGATCGGGA | N/A | qRT-PCR | N/A | |
| Cap1 | GAAAGCATCACACATGCCCTG | GGGTTTAGGTGCAGAGAATGG | N/A | qRT-PCR | N/A | |
| Car7 | CCCATCAATATCATATCCAGCC | CCCAGACACCACGGTTCTG | N/A | qRT-PCR | N/A | |
| Casz1 | GGAACTGTCTCCACTGTCAAG | AGCAGGCAGAGGGAACAGAG | N/A | genotyping PCR | N/A | |
| Casz1 | CCAGGCAGGCTTCAGGAGAA | GCTCATCCCTGGTGTTCTGG | N/A | qRT-PCR | N/A | |
| Chrna1 | GACATCACCTACCACTTCGTCA | GCGTCATCTTCTCCCCTGAGT | N/A | qRT-PCR | N/A | |
| Chrng | CAGCCTGAAGCAAGCCTCCC | CACTCCTCGTTCCCACTGTC | N/A | qRT-PCR | N/A | |
| Coro2a | CACAGAAGGGGATCGGTATC | CTGGCTCGATGAGGCTTTTGG | N/A | qRT-PCR | N/A | |
| Fgf8 | GGGGAAGCTAATTGCCAAGAG | CGCCGTGTAGTTGTTCTCCAG | N/A | qRT-PCR | N/A | |
| Galnt9 | CCCATGACAAACCCAGGAGT | GCTTTGCTGTTTCTCACCTCTC | N/A | qRT-PCR | N/A | |
| Grk1 | CCAGCAAGGACTTCTGTGAG | GGAATGAGCATCCCGGCTTC | N/A | qRT-PCR | N/A | |
| Hgf | GATCAGGACCATGTGAGGGAG | ATACCAGGACGATTTGGGATGG | N/A | qRT-PCR | N/A | |
| Insm2 | CGAGAGAAGCACCGGCTATG | GAGGGGTGGCACTTATTGATG | N/A | qRT-PCR | N/A | |
| Myl1 | TGTCCTCGCCACTCTGGGAG | GATGTGTTTGACAAAAGCTTCATAG | N/A | qRT-PCR | N/A | |
| Myo6 | GTGGTGTTCACTGTCAGTCATC | GCCGTCAATGCGTGGTTTGTG | N/A | qRT-PCR | N/A | |
| Nf2 | ACCAAGCCCACCTATCCACC | AGTCGCTTCATGTCCGTATCC | N/A | qRT-PCR | N/A | |
| Nsg2 | TCGCTGAATTTACGGTCACCA | GCTTGTGCTTGTAGACAAATCC | N/A | qRT-PCR | N/A | |
| Ocm | GAGCATCACGGACATTCTGAG | TCAAAGGTGTCTGGGTCTTGG | N/A | qRT-PCR | N/A | |
| Otof | TGCTCAACCCTGACAAGCCAG | ACCCGCAGCTCGTACTTCTTG | N/A | qRT-PCR | N/A | |
| Pitpnm1 | AACGTCACTTCCAACCACCGA | GACGTAGACATCCACCTTCTC | N/A | qRT-PCR | N/A | |
| Ppp1r14d | CCCAGGCAACAGAGAGCCT | CTTCAGGAGATGGTCCCTCC | N/A | qRT-PCR | N/A | |
| Ptgir | ATGGCTCGTTTGTACCGACCT | CTGAGTGAAGCCTCGGATCA | N/A | qRT-PCR | N/A | |
| Runx1t1 | CCTGTGGTGCTAGGCAACTCA | GTCAAAGTAGAGTTCACCAGTC | N/A | qRT-PCR | N/A | |
| Runx1t1 | CCTGGGCTTATACATGGTCATC | CCCACCTTGCTTATTCTTCTGG | N/A | PCR, 5' loxP site testing | N/A | |
| Runx1t1 | GGTTTCCATTGCTGAGTTACATC | AAGGCCCATGTTCTGGTCTTGT | N/A | PCR, 3' loxP site testing | N/A | |
| Cre | CGGTCTGGCAGTAAAAACTATC | CTACACCAGAGACGGAAATCC | N/A | genotyping PCR | N/A | |
| shRNA(Coro2a)1 | N/A | N/A | GGAGCGTCTTGGACGTTAActcctgacc caagTTGATGTCCAAGACGTTCCttttttt | shRNA expression vector production | gagGGAACGTCTTGGACATCAA | |
| shRNA(Coro2a)2 | N/A | N/A | GTGTGGAATCTGGGCATAActcctgacc caagTTGTGTCCAGGTTCCACACttttttt | shRNA expression vector production | atgGTGTGGAACCTGGACACAA | |
| shRNA(Coro2a)3 | N/A | N/A | GCGGGAGCATCTGCTATTActcctgacc caagTAGTAGCGGATGTTCCCGCttttttt | shRNA expression vector production | gagACGGGAACATCCGCTACTA | |
| shRNA(Coro2a)4 | N/A | N/A | ATGAGGTGAGCGTGGGGAActccagcc acaagTTCTCCATGCTTACCTCATttttttt | shRNA expression vector production | actATGAGGTAAGCATGGAGAA | |
| shRNA(Calb1)1 | N/A | N/A | GAATCCTACCTGTAGTCGTctccagcca caagATGACTGCAGGTGGGATTCttttttt | shRNA expression vector production | gcaGAATCCCACCTGCAGTCAT | |
| shRNA(Calb1)2 | N/A | N/A | GCAATGGGTACATAGGTGActcctgacc caagTCATCTATGTATCCGTTGCttttttt | shRNA expression vector production | atgGCAACGGATACATAGATGA | |
| Fusion of shRNA(Coro2a) targets, T2A, and Fluc | N/A | N/A | ATGGGACAGGGCctcgagCACAGAGG GAACGTCTTGGACATCAAATGGGTG ATGGTGTGGAACCTGGACACAAAGG ATAAGGGAGACGGGAACATCCGCTA CTATGAGGTAAGCATGGAGAAACCT gcatgcGgcagcggagagggcagaggaagcctg ctcacatgcggggacgtcgaggagaatcctggacc tatgGAAGATGCCAAAAACATTAAGAA GGGCCC… | shCoro2a-testing (reporter) vector production | gagGGAACGTCTTGGACATCAA [and] atgGTGTGGAACCTGGACACAA [and] gagACGGGAACATCCGCTACTA [and] actATGAGGTAAGCATGGAGAA | nt1-3: start ATG of Fluc; nt4-12: filler sequence; nt13-18: XhoI site used for subcloning the shRNA target sequence; nt19-126: shRNA target sequences; nt127-132: SphI site used for subcloning the shRNA target sequence; nt133-195: T2A coding sequence; nt196-onwards: continuation of Fluc sequence |

| | | | | | | |
|---|---|---|---|---|---|---|
| Fusion of shRNA(Calb1) targets, T2A, and Fluc | N/A | N/A | ATGGGACAGGGCctcgagGATGGCAACGGATACATAGATGAAAATGAGCTGGCAGAATCCCACCTGCAGTCATCTCTGgcatgcGgcagcggagagggcagaggaagcctgctcacatgcggggacgtcgaggagaatcctggacctatgGAAGATGCCAAAAACATTAAGAAGGGCCC… | shCalb1-testing (reporter) vector production | gcaGAATCCCACCTGCAGTCAT [and] atgGCAACGGATACATAGATGA | nt1-3: start ATG of Fluc; nt4-12: filler sequence; nt13-18: XhoI site used for subcloning the shRNA target sequence; nt19-78: shRNA target sequences; nt79-87: SphI site used for subcloning the shRNA target sequence; nt85-147: T2A coding sequence; nt148-onwards: continuation of Fluc sequence |
