## Supplementary material for "CASZ1 regulates the rate at which outer hair cells mature and is required for hearing": Table S7

**Table S7. Key resources.**

| REAGENT or RESOURCE | SOURCE | IDENTIFIER |
|---|---|---|
| **Antibodies** | | |
| Rabbit anti-RFP antibody | Rockland Immunochemicals, Inc. | Cat# 600-401-379; RRID:AB_2209751 |
| Rabbit anti-CALB1 antibody | Millipore | Cat# AB1778; RRID:AB_2068336 |
| Mouse anti-MYO7A antibody | Developmental Studies Hybridoma Bank, University of Iowa | Cat# MYO7A 138-1; RRID:AB_2282417 |
| Rabbit anti-MYO7A antibody | Proteus Biosciences | Cat# 25-6790; RRID:AB_10015251 |
| Mouse anti-Myc antibody | Developmental Studies Hybridoma Bank, University of Iowa | Cat# 9E10; RRID:AB_2266850 |
| Goat anti-OCM antibody | Santa Cruz | Cat# sc-7446; RRID:AB_2267583 |
| Mouse anti-acetylated-α-tubulin antibody | Sigma | Cat# T7451; RRID:AB_609894 |
| Alexa Fluor Plus 594-conjugated donkey anti-Mouse IgG Antibody | Themo Fisher Scientific | Cat# A32744; RRID:AB_2762826 |
| Alexa Fluor 568-conjugated donkey anti-goat IgG Antibody | Themo Fisher Scientific | Cat# A-11057; RRID:AB_2534104 |
| Alexa Fluor 488-conjugated donkey anti-rabbit IgG Antibody | Themo Fisher Scientific | Cat# A-21206; RRID:AB_2535792 |
| Alexa 594-conjugated donkey anti-rabbit antibody | Thermo Fisher Scientific | Cat# A-21207; RRID:AB_141637 |
| Alexa 488-conjugated donkey anti-mouse antibody | Thermo Fisher Scientific | Cat# A-21202; RRID:AB_141607 |
| **Bacterial and Virus Strains** | | |
| Recombinant AAV: (ss)AAV-Coro2a | This paper | N/A |
| Recombinant AAV: (ss)AAV-Myl1 | This paper | N/A |
| Recombinant AAV: (ss)AAV-Car7 | This paper | N/A |
| Recombinant AAV: (ss)AAV-mCherry | This paper | N/A |
| Recombinant AAV: (ss)AAV-Coro2a-IRES-mCherry | This paper | N/A |
| Recombinant AAV: (ss)AAV-Coro2a-Myc | This paper | N/A |
| Recombinant AAV: (ss)AAV-shRNA$_{Calb1}$ 1 | This paper | N/A |
| Recombinant AAV: (sc)AAV-shRNA$_{Calb1}$ 1 | This paper | N/A |
| Recombinant AAV: (sc)AAV-shRNA$_{Calb1}$ 2 | This paper | N/A |
| Recombinant AAV: (sc)AAV-shRNA$_{Coro2a}$ 1 | This paper | N/A |
| Recombinant AAV: (sc)AAV-shRNA$_{Coro2a}$ 2 | This paper | N/A |
| Recombinant AAV: (sc)AAV-shRNA$_{Coro2a}$ 3 | This paper | N/A |
| Recombinant AAV: (sc)AAV-shRNA$_{Coro2a}$ 4 | This paper | N/A |
| **Chemicals and Reagent** | | |
| Alexa Fluor 488 Phalloidin | Thermo Fisher Scientific | Cat# A12379 |
| Thermolysin | Sigma | Cat# T7902; CAS 9073-78-3 |
| FM1-43 Dye | Thermo Fisher Scientific | Cat# T3163 |
| DNase I | Worthington Biochemical Corp. | Cat# LK003170; CAS 9003-98-9 |
| B-27 Supplement (50X) | Thermo Fisher Scientific | Cat# 17504044 |
| N-2 Supplement (100X) | Thermo Fisher Scientific | Cat# 17502048 |
| Neurobasal-A Medium | Thermo Fisher Scientific | Cat# 10888022 |

| | | |
|---|---|---|
| Lipofectamine LTX with PLUS reagent | Thermo Fisher Scientific | Cat# 15338100 |
| TransIT-VirusGEN® Transfection Reagent | Mirus Bio | Cat# MIR 6700 |
| ProLong Glass Antifade Mountant | Thermo Fisher Scientific | P36984 |

Critical Commercial Assays

| | | |
|---|---|---|
| SsoAdvanced Universal SYBR® Green Supermix | Bio-Rad | Cat# 1725271 |
| RNeasy Micro Kit | Qiagen | Cat# 74004 |
| SuperScript™ IV First-Strand Synthesis System | Thermo Fisher Scientific | Cat# 18091050 |
| Dual-Luciferase Reporter Assay System | Promega | Cat# E1910 |

Deposited Data

| | | |
|---|---|---|
| RNA sequencing data | This paper | GSE300215 |

Experimental Models: Cell Lines

| | | |
|---|---|---|
| HEK293T cell | Takara Bio | Cat# 632273 |

Experimental Models: Organisms/Strains

| | | |
|---|---|---|
| Mouse: Casz1$^{fl/fl}$ | Mutant Mouse Resource and Research Center (MMRRC) | RRID:MMRRC_041176-UNC |
| Mouse: C57BL/6 | Charles River Laboratories | Cat# 027 |
| Mouse: Runx1t1$^{fl/fl}$ | GemPharmatech Co., Ltd. | N/A |
| Mouse: B6;CBA-Tg(Sox10-cre)1Wdr/J | The Jackson Laboratory | RRID:IMSR_JAX:025807 |
| Mouse: B6.Cg-*Shh*$^{tm1(EGFP/cre)Cjt/}$ J | The Jackson Laboratory | RRID:IMSR_JAX:005622 |
| Mouse: B6(Cg)-*Calb2*$^{tm1(cre)Zjh}$ /J | The Jackson Laboratory | RRID:IMSR_JAX:010774 |
| Mouse: FVB/N-*Gfi1*$^{+/Cre}$ | Yang et al., 2010 | N/A |

Oligonucleotides

| | | |
|---|---|---|
| Listed in Table S6 | IDT | N/A |

Recombinant DNA

| | | |
|---|---|---|
| pGL4.74 [hRluc/TK] vector | Promega | Cat# E692A |
| pGL4.54 [luc2] Vector | Promega | Cat# E5061 |
| pALD-X80 | Aldevron | Cat# 5017-10 |
| pFBAAVCAGmcsBgHpA | Viral Vector core facility, University of Iowa | Cat# G0463 |
| pscAAVmcsBgHpa | Viral Vector core facility, University of Iowa | Cat# G0345 |
| pUCmini-iCAP-PHP.eB | a gift from Viviana Gradinaru | Addgene plasmid # 103005 |
| pFBAAVCAG-Coro2a-BgHpA shuttle plasmid | This paper | N/A |
| pFBAAVCAG-Myl1-BgHpA shuttle plasmid | This paper | N/A |
| pFBAAVCAG-Car7-BgHpA shuttle plasmid | This paper | N/A |
| pFBAAVCAG-mCherry-BgHpA shuttle plasmid | This paper | N/A |
| pFBAAVCAG-Coro2a-IRES-mCherry-BgHpA shuttle plasmid | This paper | N/A |

| | | |
|---|---|---|
| pFBAAVCAG-Coro2a-Myc-BgHpA shuttle plasmid | This paper | N/A |
| pFBAAVU6-shRNA$_{Calb1}$ 1 shuttle plasmid | This paper | N/A |
| pscAAVU6-shRNA$_{Calb1}$ 1 shuttle plasmid | This paper | N/A |
| pscAAVU6-shRNA$_{Calb1}$ 2 shuttle plasmid | This paper | N/A |
| pscAAVU6-shRNA$_{Coro2a}$ 1 shuttle plasmid | This paper | N/A |
| pscAAVU6-shRNA$_{Coro2a}$ 2 shuttle plasmid | This paper | N/A |
| pscAAVU6-shRNA$_{Coro2a}$ 3 shuttle plasmid | This paper | N/A |
| pscAAVU6-shRNA$_{Coro2a}$ 4 shuttle plasmid | This paper | N/A |
| pGL4.54-Targets-of-all-shRNA$_{Coro2a}$ -Luc | This paper | N/A |
| pGL4.54-Targets-of-all-shRNA$_{Calb1}$ -Luc | This paper | N/A |

Software and Algorithms

| | | |
|---|---|---|
| Zen lite 2012 | Zeiss | https://www.zeiss.com/microscopy/en_us/products/microscope-software/zen.html RRID:SCR_013672 |
| DAVID | Huang da et al., 2009 | https://david.ncifcrf.gov/ RRID:SCR_001881 |
| GraphPad PRISM v.10.00 | GraphPad Prism | https://www.graphpad.com/scientific-software/prism/ RRID:SCR_002798 |
| STAR aligner v2.7.11a | Dobin et al., 2013 | https://github.com/alexdobin/STAR |
| edgeR | Robinson et al., 2009 | https://github.com/OliverVoogd/edgeR |
| Trimmomatic | Bolger et al., 2014 | http://www.usadellab.org/cms/index.php?page=trimmomatic |
| featureCounts | Liao et al., 2013 | https://toolshed.g2.bx.psu.edu/repository?repository_id=00d38f44b709f93a |
